# Adaptive coupling of chromosomal inversions with multilocus ecological, mating-bias, and hybrid-incompatibility genotypes facilitates sympatric speciation

**DOI:** 10.64898/2026.04.02.716010

**Authors:** Curtis Heyda, David Hernandez, Taylor Rios, Lea Ozdere, Zakia Sultana, John Lin

## Abstract

Chromosomal inversions (CIs) are widespread structural variants that suppress recombination and maintain favorable allele combinations as tightly linked supergenes. They are increasingly recognized for their contributions to local adaptation and speciation, yet their role in sympatric speciation under ongoing gene flow remains unresolved. Here, we develop mathematical models and computer simulations to examine how polygenic architectures and CI invasion jointly shape the evolution of reproductive isolation (RI) in a two-niche model under disruptive ecological selection. Extending previous frameworks, we incorporate multilocus ecological traits, multilocus mating-bias traits, and multilocus hybrid-incompatibility genotypes, and evaluate the invasion fitness and evolutionary consequences of CIs capturing different combinations of locally adaptive alleles. We show that the evolutionary impact of CI invasion depends critically on how polygenic trait structure influences hybrid production and the strength of barrier mechanisms. Increasing the number of ecological or hybrid-incompatibility loci strengthens disruptive ecological selection and postzygotic incompatibility selection by generating more unfit hybrids, whereas increasing the number of mating-bias loci weakens premating selection because mating-bias hybrids remain viable within the same niche. Accordingly, CIs capturing ecological or hybrid-incompatibility alleles tend to reduce the effective number of loci, diminish hybrid loss, and weaken existing RI, whereas CIs capturing mating-bias alleles strengthen premating isolation by reducing the effective number of mating-bias loci. Importantly, CIs that couple alleles across distinct barrier mechanisms exhibit elevated invasion fitness and generate synergistic reinforcement and positive feedback among premating isolation, postmating isolation, and ecological divergence. These findings reconcile contrasting theoretical predictions by demonstrating that CIs can either facilitate or constrain sympatric speciation depending on how they reshape effective locus number and barrier coupling, and provide a unified framework for understanding how structural genomic variants interact with polygenic architectures to influence the origin and stability of reproductive isolation.

## I. INTRODUCTION

Chromosoml inversions (CIs) are widespread structural variants found across the genomes of diverse taxa and are increasingly recognized as important contributors to adaptation and speciation [1-9]. By suppressing recombination within inverted regions, CIs can preserve favorable combinations of alleles and facilitate their joint inheritance as tightly linked “supergenes” [3, 10-12]. Inversions that capture locally adaptive allele combinations may therefore function as “adaptive cassettes” that spread among populations and facilitate adaptation across heterogeneous environments [13-16].

Beyond local adaptation, the capacity of CIs to suppress recombination is thought to be the key mechanism by which they contribute to the evolution of reproductive isolation (RI) in speciation [17, 18]. For example, in the sympatric species pair Drosophila pseudoobscura and D. persimilis, traits contributing to reproductive isolation map primarily or exclusively to chromosomal regions that differ by inversions, implicating an important role of CIs in the evolution of sympatric speciation [19-21]. In addition to recombination suppression, other inversion-associated mechanisms—such as gene disruption at breakpoints or reduced fitness of heterokaryotypes [2, 22]—may further contribute to speciation; however, empirical support for these alternative mechanisms remains limited [1, 2, 18, 23-26]. Despite extensive theoretical and empirical study, the conditions under which CIs promote, constrain, or otherwise reshape the evolution of RI remain incompletely understood [1, 27].

Chromosomal inversions have been studied since the 1920s [2]. Theoretical models have traditionally emphasized the role of CIs in allopatric and parapatric divergence, where partial geographic isolation allows locally adaptive alleles to accumulate before being protected from recombination by inversions [4, 5, 18, 28-33]. Despite extensive theoretical work, the role of chromosomal inversions in sympatric speciation— where ecological divergence and reproductive isolation evolve in the face of ongoing gene flow— remains less well resolved [5, 22, 27, 34].

The evolutionary advantage of recombination suppression is expected to be greatest in sympatry, where gene flow is substantial and divergence traits are polygenic. Under these conditions, CIs may promote RI by assembling small-effect alleles into supergenes of large phenotypic effect or by coupling ecological divergence with isolating mechanisms such as mating bias, habitat preference, or intrinsic hybrid incompatibilities, potentially functioning as “magic” or “pseudomagic” traits [35, 36].

However, prior modeling studies indicate that the evolutionary consequences of CI invasion during sympatric speciation are not straightforward [37]. By capturing locally adaptive alleles, an inversion may reduce maladaptive hybrid production and thereby weaken the disruptive ecological selection that drives divergence and RI [38]. Moreover, interactions between inversion-mediated recombination suppression and the polygenic architecture of ecological traits, assortative mating, or postzygotic incompatibilities can generate nonlinear and sometimes counterintuitive population dynamics [38]. These complexities underscore the need for models that explicitly incorporate multilocus traits to determine when CI invasion facilitates or constrains the evolution of RI.

To address this knowledge gap, we use mathematical modeling and computer simulations to investigate how polygenic architectures and CI invasion shape evolutionary dynamics during sympatric speciation. Building on a previously developed two-niche model of disruptive ecological selection and assortative mating [39], we extend the framework to incorporate multilocus ecological traits, multilocus mating-bias traits, and multilocus hybrid-incompatibility genotypes. We then model CI mutations that capture various combinations of ecological, mating-bias, and incompatibility alleles to evaluate their invasion fitness, equilibrium frequencies, and effects on the evolution of RI in sympatry.

Our objectives are threefold. First, we examine how polygenic architectures influence disruptive ecological selection, mating-bias selection, and intrinsic hybrid incompatibility selection in our model of sympatric speciation, and how they affect system convergence and the emergence of RI. Second, we analyze how the invasion of CIs capturing different combinations of alleles alters these selection mechanisms and reshapes evolutionary dynamics. Third, we examine how the capture and coupling of alleles underlying distinct isolating mechanisms alter CI invasion dynamics and generate reciprocal reinforcement among ecological divergence, premating isolation, and postzygotic incompatibilities. By systematically varying the number of gene loci, selection strengths, niche sizes, and other model parameters, we characterize the conditions under which CIs promote, weaken, or transform RI.

Our results show that the evolutionary consequences of CI invasion depend critically on how polygenic architectures affect hybrid production and the strength of selection across barrier mechanisms. Increasing the number of loci strengthens disruptive ecological selection and hybrid incompatibility selection by generating more unfit hybrids, whereas reducing locus number strengthens mating-bias selection because mating-bias hybrids remain viable within the same niche. Consequently, CIs capturing ecological or incompatibility alleles tend to weaken selection and diminish existing RI, while those capturing mating-bias alleles strengthen selection and enhance RI. Notably, CIs that couple alleles across distinct barrier processes consistently exhibit higher invasion fitness and generate stronger RI, underscoring the importance of barrier coupling in determining whether inversions promote or constrain speciation.

By integrating multilocus trait architectures with explicit modeling of CI invasion, our study delineates the conditions under which CIs can facilitate, constrain, or redirect the evolution of RI in sympatry. We demonstrate that the evolutionary consequences of inversion spread depend critically on how inversions alter the effective number of loci and the coupling among loci underlying ecological divergence, mating bias, and hybrid incompatibilities. In doing so, we reconcile previously contrasting theoretical predictions by showing that CIs can either weaken or reinforce divergence, depending on how they reshape both individual barrier processes and their interactions. More broadly, our results provide a unified conceptual framework for understanding how structural genomic variants interact with polygenic architectures and multiple isolating mechanisms to generate RI and influence the trajectory of speciation.

## II. METHODOLOGY

We extended the mathematical model of a two-allele mating-bias barrier developed in a prior study [39] by incorporating multilocus ecological, mating-bias, and hybrid-incompatibility genotypes and examining how the invasion of CIs into such polygenic systems may alter their dynamics to either facilitate or impede the evolution of RI. Using MATLAB (version R2021a) and its App Designer tool, we created interactive graphical user interface (GUI) applications that enable users to specify any number of gene loci underlying ecological, mating-bias, and hybrid-incompatibility traits, as well as to simulate the invasion of CIs into these premating and postmating barrier systems. Three GUI applications were constructed: one models multilocus ecological barriers that permit viable ecological hybrids; the other two model multilocus mating-bias barriers and multilocus hybrid-incompatibility barriers in the absence of viable ecological hybrids. The outcomes of these nonlinear models are visualized as phase portraits, providing a bird’s-eye view of the solution landscape and an intuitive understanding of the underlying population dynamics.

### 1 Mathematical model of a two-allele mating-bias barrier developed in a prior study

The models used in this study are modifications of an earlier two-mating-bias-allele, two-ecological-niche mathematical model without viable hybrids, developed in our previous work (see Fig 3) [39]. For context, a brief overview of the original model is presented here.

The flowchart in Fig 1 shows the life cycle of a hypothetical sympatric population used in our model. In each generation, individuals undergo ecological and sexual selection to produce offspring for the next generation. The ecosystem contains two distinct niche resources, *A* and *B*, generated by disruptive ecological selection. During the ecological selection phase, only individuals with genotypes adapted to one of the two niches (i.e., ecotype *A* or ecotype *B*) can access the niche-specific resources and survive.

**Fig 1.**
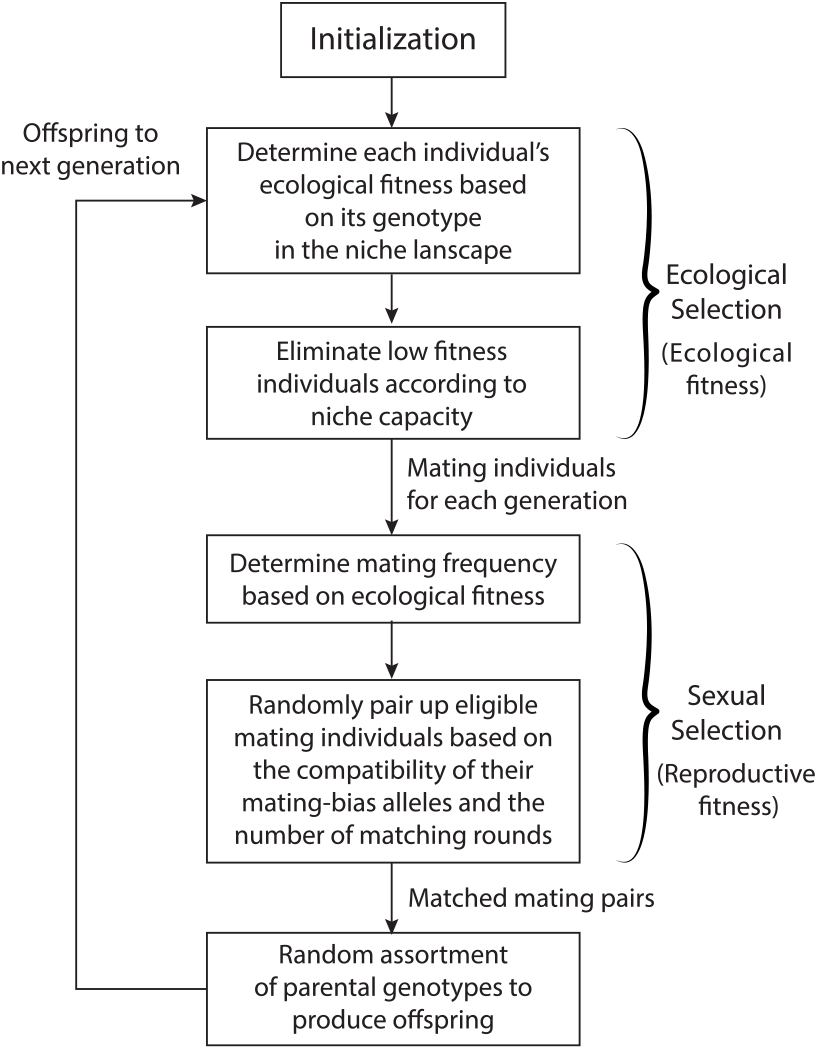
Flowchart showing the life cycle of a sympatric population used in the computer simulations.

Next, in the sexual selection phase (see flowchart in Fig 2), survivors of ecological selection encounter one another randomly to find mates. The outcome of each matching encounter is determined by the mating-bias alleles—*X* or *Y*—that individuals carry at a single gene locus. Compatibility between alleles is defined by the Matching Compatibility Table shown in Fig 4. Individuals with identical mating-bias alleles always match with a probability of 1, while individuals with different alleles match with a probability defined by the mating-bias value *α*, which ranges from 0 to 1.

**Fig 2.**
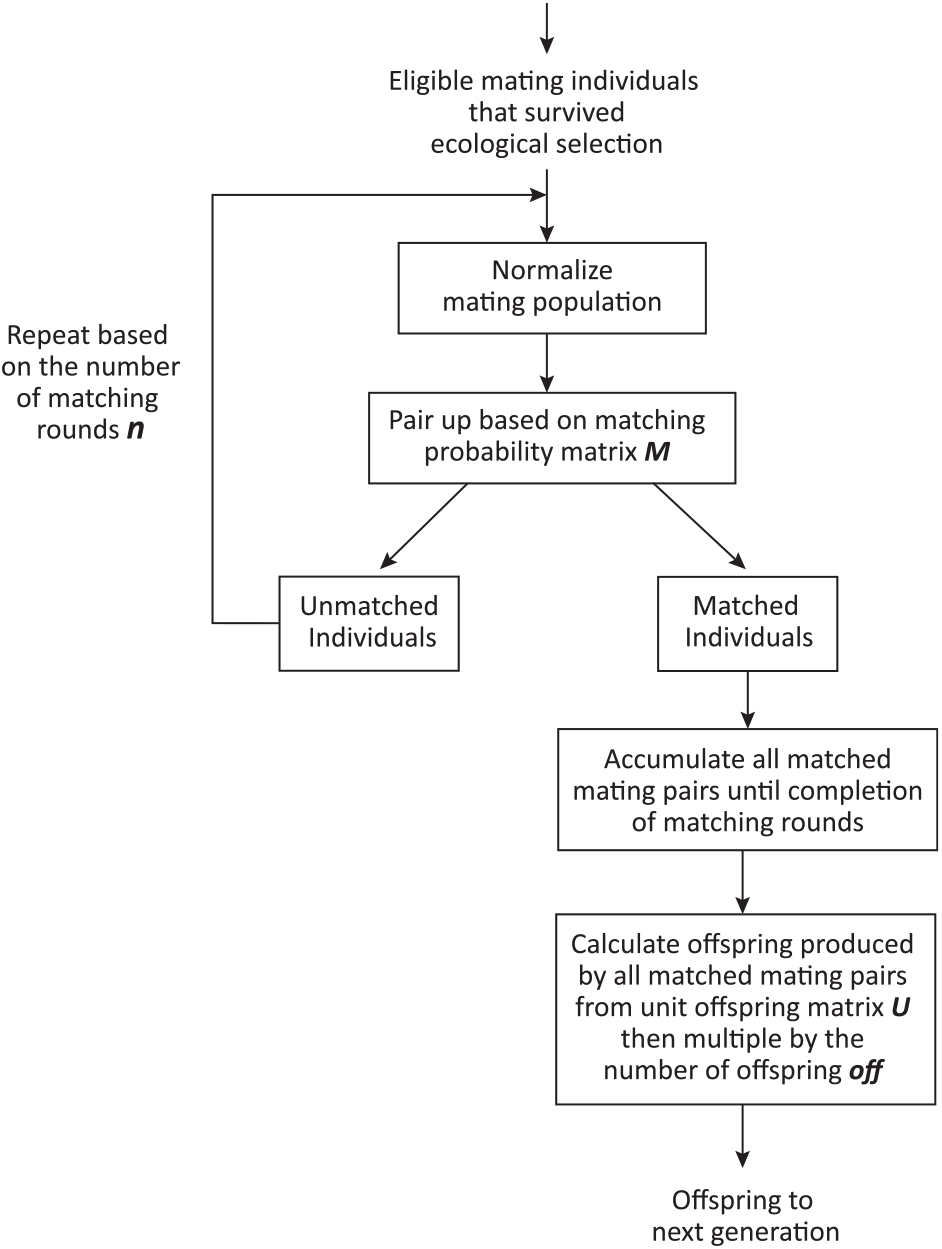
Flowchart showing the algorithm used to match eligible mating individuals and reproduce offspring.

**Fig 3.**
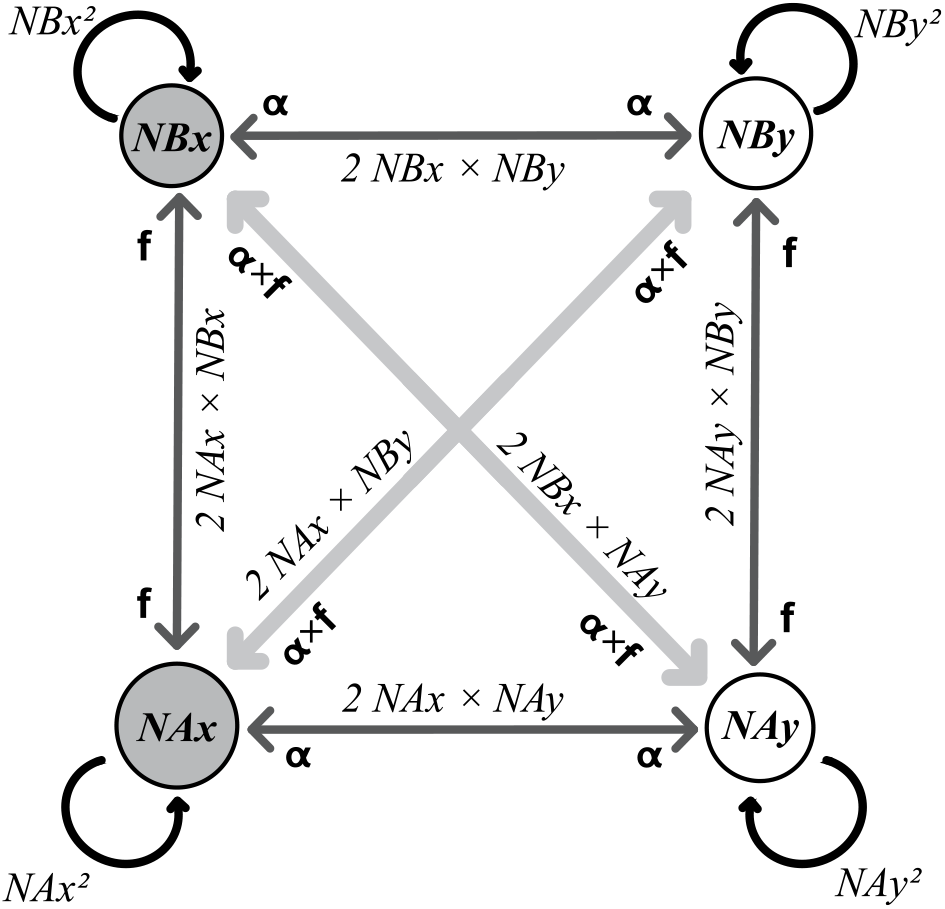
Mathematical model of a 2-mating-bias-allele, 2-ecological-niche sympatric ecosystem.

**Fig 4.**
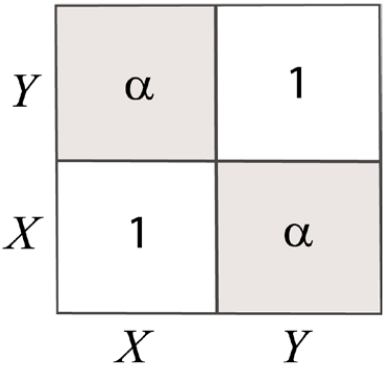
Matching Compatibility Table for mating-bias alleles. Same-allele individuals (*X* or *Y*) always match with a probability of 1, whereas different-allele individuals match with a probability of *α* (0 ≤ *α* ≤ 1).

Unmatched individuals may attempt to find a mate again in subsequent rounds, up to a maximum of *n* rounds. Those who fail to match after *n* rounds die without producing offspring, while matched pairs reproduce offspring through random assortment of their alleles at each gene locus. Thus, the number of matching rounds, *n*, reflects the cost of assortative mating, and the values of *α* and *n* together determine the strength of sexual selection.

Fig 3 shows the two-mating-bias-allele, two-niche mathematical model used to describe the population dynamics of the sympatric population during a matching round. The model includes four genotype groups—*NAx, NAy, NBx*, and *NBy*—representing the normalized population ratios of the different ecotypes and their associated mating-bias alleles. Because all parametric values in the model are normalized, *NAx* + *NAy* + *NBx* + *NBy* = 1.

As these genotype groups encounter one another randomly in sympatry, their pairwise encounter probabilities are summarized in the Encounter Probability Matrix shown in Fig 5. The elements of this matrix represent the encounter probabilities of all possible pairings and collectively sum to 1.

**Fig 5.**
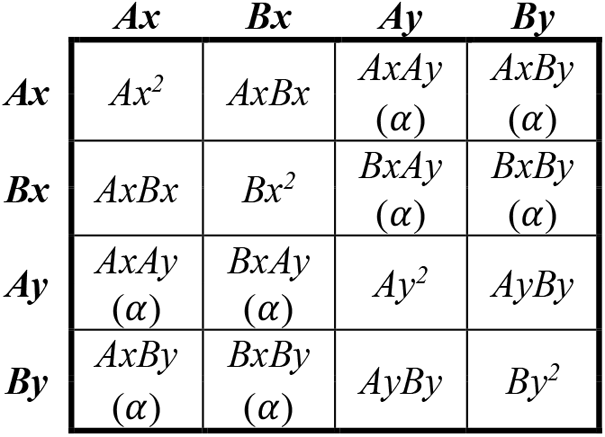
Encounter Probability Matrix (*M*) for a sympatric model with no viable hybrids. The variable *α* in parentheses serves as a reminder that inter-niche matching success is subsequently determined by multiplying the encounter probabilities by the mating-bias *α*.

After *n* matching rounds, all successfully matched genotype pairs are collected. These parental pairs reproduce by randomly assorting alleles at each gene locus. Fig 6 presents a Unit Offspring Matrix that lists the expected genotype ratios of offspring produced by different parental genotype combinations. In this matrix, the variable *f* denotes the “offspring return ratio,” which specifies the proportion of offspring that inherit the same genotype as their parents. The value of *f* ranges from 0 to 0.5 and serves as a measure of the strength of ecological selection.

**Fig 6.**
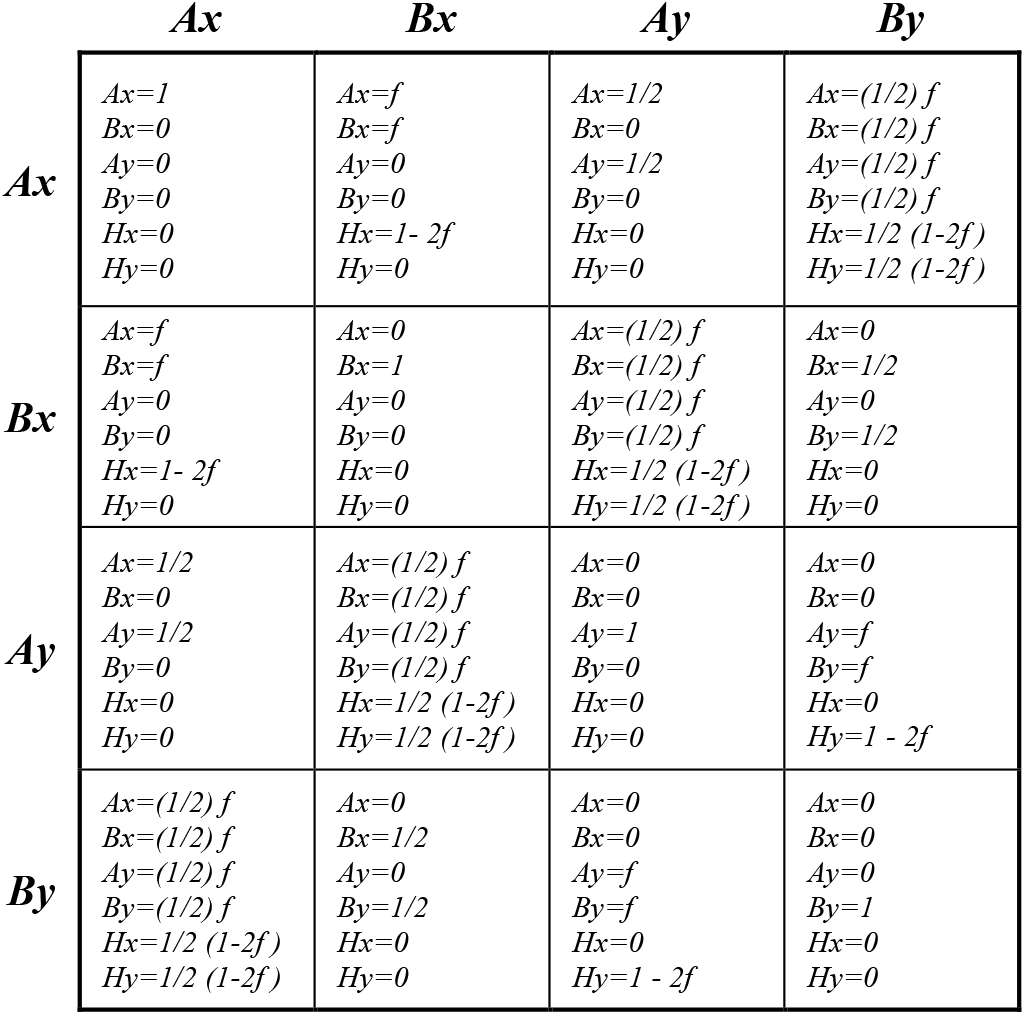
Unit Offspring Matrix (*U*).

The unit offspring ratios represent the normalized proportions of different offspring genotypes produced by each generation of matched parental pairs, under the assumption that each parent produces only one offspring to replace itself. The values in each cell of the Unit Offspring Matrix sum to 1. Because the ratios of all parental pairs are normalized (i.e., they sum to 1), the total offspring genotype ratios produced by the parental pairs also sum to 1. To compute the final offspring population sizes, the unit offspring ratios are multiplied by a fertility parameter, *off*, which specifies the actual number of offspring each parent produces.

Although the model assumes no viable hybrid offspring by default, scenarios involving viable hybrids supported by hybrid niche resources can still be represented by increasing the value of *f*. Raising the value of *f* decreases the effective strength of ecological selection.

To avoid the effect of “incumbent selection”—a phenomenon in which a dominant population can use its numerical advantage to eliminate a minority population through their attritive interactions—our mathematical model assumes that ecotypes in each niche reproduce sufficient offspring to fully occupy the niche’s carrying capacity after each mating generation [39]. Therefore, in our model, the normalized niche population ratios, *NA* and *NB*, are always fixed at the beginning of the sexual selection phase in each generation.

We then use computer applications to generate *Ax*/*Bx* phase portrait solutions of the model using user-defined parameter values (see Fig 7). In the phase portrait, *AX* represents the normalized ratio of niche-*A* ecotypes carrying the *X* allele, while *Ay* denotes those with the *Y* allele, such that *Ax* + *Ay* = 1. The same definitions apply to *Bx* and *By* for niche *B*. The phase portrait displays changes in these normalized genotype ratios over *g* generations.

**Fig 7.**
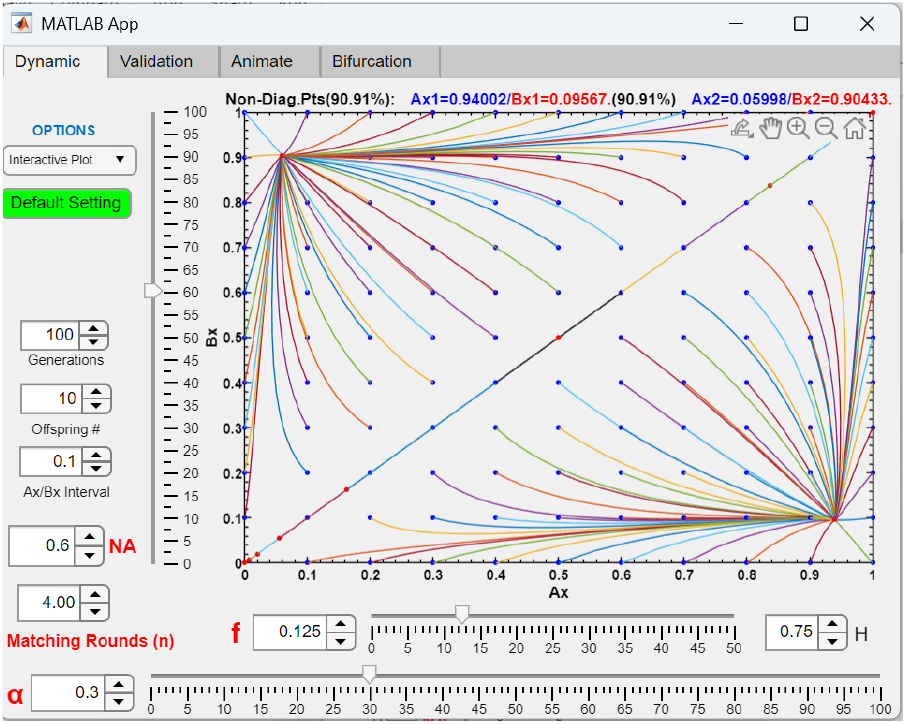
Representative phase portrait showing the population dynamics of a two-mating-bias-allele, two-niche model developed in a prior study. Given the parameter values *NA* = 0.6, *α* = 0.3, *f* = 0.125, and *n* = 4, the phase portrait’s vector field converges after *g* = 100 generations to fixed-point polymorphisms at *Ax*1/*Bx*1 = 0.94002/0.09567 and *Ax*2/*Bx*2 = 0.05998/0.90433, establishing premating RI between niche *A* and niche *B* ecotypes. In this case, the system is said to be “convergent” (i.e., its phase portrait converges to fixed points), in contrast to a “divergent” system, where the phase portrait does not converge to fixed points.

We classify a barrier’s phase portrait as either “convergent,” in which the vector field leads (converges) to fixed points, or “divergent,” where no fixed points are present. Assuming that *NAx* represents the largest genotype group in the model, maximum RI is achieved when *NAx* and *NBy* are large, while *NAy* and *NBx* are small. In the *Ax*/*Bx* phase portrait, this outcome corresponds to a fixed point located as close as possible to the bottom right corner, where *Ax* = 1 and *Bx* = 0.

### 2 Extended model of multilocus ecological genotypes with viable ecological hybrids

We developed a GUI application (MultiEcoCI) based on the mathematical framework illustrated in Fig 3, extending it to accommodate any specified number of ecological loci and to allow for the existence of viable ecological hybrids. This application enables us to investigate the invasion dynamics of a chromosomal inversion that captures an arbitrarily defined set of ecological alleles, with or without associated mating-bias alleles, and to examine how such an invasion alters the resulting population dynamics. The model incorporates ecological hybrid niche resources that sustain viable populations of hybrid genotypes within the system. As in the original framework, we assume that in each generation sufficient offspring of the pure parental ecotypes (i.e., niche *A* and niche *B* ecotypes) and hybrid ecotypes (i.e., ecological hybrids produced by matings between niche *A* and niche *B* ecotypes) are generated to fill all specified niche carrying capacities. Consequently, at the beginning of each mating generation, the population ratios of all niche ecotypes are fixed according to their respective niche carrying capacities. Because matings among viable hybrids regenerate the parental (niche *A* and niche *B*) ecotypes and determine the offspring return ratio, the variable *f* is no longer used to specify the offspring return ratio or to represent the strength of ecological disruption. Instead, these quantities are determined by the amount and distribution of hybrid niche resources. For mating-bias discrimination, individuals retain the same single-locus mating-bias genotype as in the original model (Fig 3), with mating-bias alleles *X* and *Y* and mating-bias value *α*.

Fig 8 shows the computer interface of the GUI application. The number of letters in the *CI Vector* input determines the number of gene loci for the ecological genotype. Each ecological gene locus has two alleles, *a* or *b*: the niche *A* ecotype carries all *a* alleles (e.g., *aaa*), the niche *B* ecotype carries all *b* alleles (e.g., *bbb*), and hybrid ecotypes contain mixtures of the two (e.g., *aba, bba*, etc.). The *a* and *b* alleles are distinct entities at different loci, even though the same letters are used for simplicity. In the example shown in Fig 8, the *CI Vector* is “*aa*0”, indicating three ecological loci, with the CI capturing the *a* alleles at the first two loci. The digit “0” in the third position indicates that no allele is captured at that locus. Each individual has one mating-bias gene locus, which has two alleles, *X* or *Y*. The input field *CI mating bias* specifies which mating-bias allele is captured by the CI—either *X* or *Y*—while “0” denotes no capture. Therefore, in the Fig 8 example, the composition of the CI is “*aa*0*X*.”

**Fig 8.**
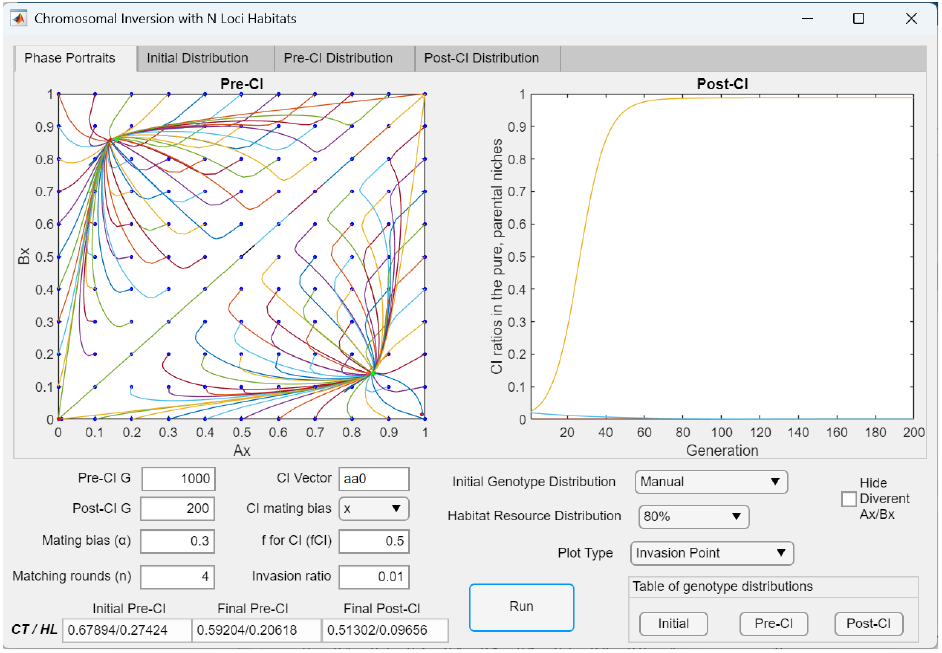
User interface of the application (MultiEcoCI) used to model multilocus ecological genotypes and the invasion dynamics of CIs. In this example, the application models a sympatric population with 80% of the habitat resources allocated to pure parental ecotypes (equally divided between niches *A* and *B*) and 20% allocated equally among hybrid niches to sustain viable hybrid ecotypes. The initial genotype and habitat resource distributions used are shown in Fig 9. Each individual carries three ecological gene loci (specified by the number of digits in the CI vector, which is three in this example) and one mating-bias locus. The *Invasion Point* plot option displays all initial *Ax*/*Bx* population ratios in the *Ax*/*Bx* phase portrait against the invading CI population ratio (which is 0.01 in this example). The figure illustrates the invasion dynamics of a chromosomal inversion (CI = *aa*0*X*) that captures two of the three locally adaptive ecological alleles in niche *A* and the *X* mating-bias allele. Before the CI invasion, the initial population ratios of the *Ax* and *Bx* ecotypes (blue dots) in the *Ax*/*Bx* phase portrait converge to a fixed point at *Ax*/*Bx* = 0.8606/ 0.1428 (green dot) after 1000 generations (*Pre-CI G*). Afterward, when the CI is introduced at an initial population ratio of 0.01, it is able to successfully invade niche *A* and stabilize at a population ratio of 0.9877 after 200 generations (*Post-CI G*). The CI invasion plot on the right shows this invasion trajectory. This invasion shifts the *Ax*/*Bx* fixed point to *Ax*/*Bx* = 0.9920/0.0161 (red dot), located closer to the lower-right corner of the *Ax*/*Bx* phase portrait, indicating stronger premating RI. Consequently, the total inter-niche offspring ratio (*CT*) and maladaptive hybrid loss (*HL*) progressively decline from the *Initial Pre CI* to *Final Post CI* stages (i.e., from *CT*/*HL* = 0.67894/ 0.27424 at *Initial Pre CI* to 0.59204/0.20618 at *Final Pre-CI*, and finally reaching 0.51302/0.09656 at *Final Post CI* after the CI invasion), producing ever stronger RI between the parental niche ecotypes. See text for a description of the parametric variables used in the interface.

The relative amount of niche resources available to each niche ecotype is specified by the input field *Habitat Resource Distribution*. Three options are available: *Manual, 90%, 80%*, and *Extreme*. Selecting the *Manual* option displays the popup table on the left in Fig 9, allowing the user to specify the normalized ratios of niche resources (niche carrying capacity) for each ecotype, with the requirement that all ratios must sum to 1. In this example, 80% of the total available resources are divided equally between the pure parental niche *A* and *B* ecotypes, while the remaining 20% are allocated equally among the six hybrid ecotypes. This distribution is equivalent to selecting the *80%* option for *Habitat Resource Distribution*. Similarly, selecting the *90%* option assigns 90% of the niche resources equally to the niche *A* and *B* ecotypes, while the remaining 10% are distributed equally among the hybrid ecotypes. The *Extreme* option allocates 50% of the total niche resources equally to niche *A* and niche *B*, with no resources available for hybrid niches.

**Fig 9.**
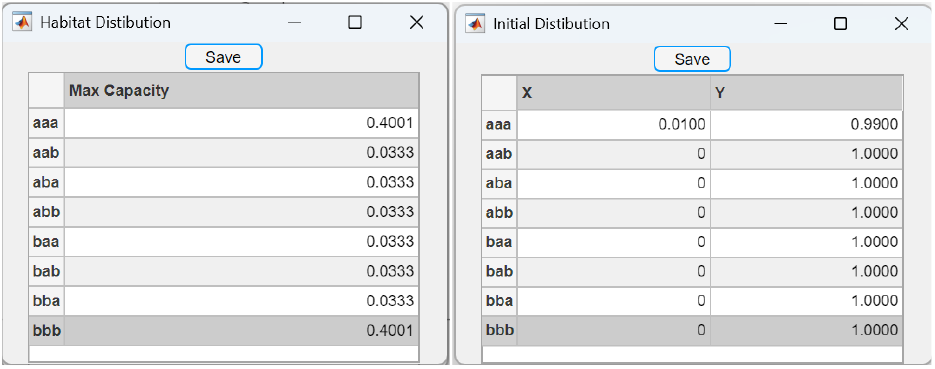
Manually setting the habitat resource distribution and initial genotype distribution in the GUI application (MultiEcoCI) used to model multilocus ecological genotypes. Selecting the *Manual* option for the habitat resource distribution in Fig 8 displays the popup table on the left, which allows the user to specify the normalized niche resource distribution (niche carrying capacities) for all possible ecological niches. Here, *aaa* and *bbb* represent the pure parental niches *A* and *B*, respectively, while the remaining entries correspond to hybrid niches. All niche resource ratios must sum to 1. As shown, the values set here correspond to the habitat resource distribution obtained by selecting the “80%” option. Similarly, selecting the *Manual* option for the initial genotype distribution displays the popup table on the right, which allows the user to define the starting ratios of the *X* and *Y* mating-bias alleles in each genotype niche—that is, the *X* and *Y* ratios in each niche population before running the pre-CI phase. The *X* and *Y* ratios in each niche must sum to 1. As shown, the values in the table depict a situation in which an *X* mutant allele arises in niche *A* with an initial population ratio of *Ax* = 0.01 and attempts to invade a sympatric population that otherwise possesses only the *Y* allele.

The mating-bias value (α) and the number of mating rounds (*n*) are the same variables defined in the two-allele model of the mating-bias barrier.

The program runs in two phases: the pre-CI phase followed by the post-CI phase. The pre-CI phase simulates the two-allele mating-bias barrier model before the emergence of a CI mutation. The *Pre-CI G* input specifies the number of generations to be run in this phase, while the *Initial Genotype Distribution* input defines the niche genotype distribution before the simulation (i.e., at generation 1). Selecting the *Manual* option in *Initial Genotype Distribution* displays the popup table on the right of Fig 9. This table allows the user to specify the initial *X* and *Y* allele ratios in each ecotype population. The two ratios must sum to 1 for each ecotype. In this example, the values specified in the table depict a situation in which a mutant *X* allele arises in niche *A* (with a population ratio of *Ax* = 0.01) and attempts to invade a sympatric population that carries only the *Y* allele.

In the post-CI phase, the program simulates the invasion dynamics of a CI mutation that arises in the population and examines its effects on genotype compositions and RI between the parental niche-*A* and niche-*B* ecotypes. The *Post-CI G* input specifies the number of generations to be run in this phase, while the *Invasion Ratio* input specifies the initial population ratio of the CI mutant before the simulation begins. The alleles captured by the CI mutant are defined by entries in the *CI Vector* and *CI mating bias* input fields (e.g., CI = *aa*0*X* in the Fig 8 example).

The variable *fCI* specifies the offspring return ratio of the captured CI alleles in heterokaryotype matings (i.e., when an individual carrying the CI mates with another individual that does not). This value is typically set to 0.5 if we ignore the rare occurrence of single crossovers in the inverted regions [22, 40], as well as recombination caused by gene flux from double crossovers and gene conversion [41]. In heterokaryotype matings, alleles at gene loci captured by the CI remain linked and cannot be broken up by recombination, whereas alleles at loci outside the inverted region can freely recombine and are assorted randomly in the model.

As an illustrative example, consider a mating between an individual in niche *A* with a *CI* = *aa*0*X* genotype (i.e., *aaaX*) and another individual from niche *B* with the *bbbY* genotype. If *fCI* = 0.5, half of their offspring will retain the aa alleles in the first two gene loci and the *X* allele, since this allele combination cannot be broken up by recombination within the CI. The other half will retain the *bb* alleles at the corresponding loci and the *Y* allele. Because the third locus is not captured by the CI, the resulting offspring genotype composition consists of equal proportions of *aaaX, aabX, bbaY*, and *bbbY*.

An *fCI* value less than 0.5 represents a situation in which a portion of the offspring produced by hetero-karyotype matings is lost, either due to single cross-over events or reduced viability of heterokaryotype hybrid offspring. In this case, the proportions of the viable offspring genotypes (*aaaX, aabX, bbaY*, and *bbbY*) are reduced by multiplying their original proportions by a factor of *fCI*/0.5.

The GUI application provides two plot options: *Invasion Point* and *Single Line*. Fig 8 shows the *Invasion Point* plot option. The phase portrait on the left displays changes in the *Ax*/*Bx* population ratios during the pre-CI and post-CI phases of the simulation. In the pre-CI phase, the phase portrait plots the vector-field trajectories of all possible permutations of the initial *Ax*/*Bx* population ratios (displayed as blue dots spaced at 0.1 intervals) over the specified pre-CI generations. The final pre-CI *Ax*/*Bx* population ratios are shown as green dots. In this example, after 1000 generations, all nondiagonal population vectors converge to fixed points on opposite sides of the diagonal line.

After a CI mutation arises at the start of the post-CI phase, the program plots the population vector trajectories of all final pre-CI *Ax*/*Bx* population ratios (green dots) against the initial CI mutant population ratio (0.01) over 200 post-CI generations. The final post-CI *Ax*/*Bx* population ratios are shown as red dots in the phase portrait.

The plot on the right of Fig 8 displays the CI population trajectories in the post-CI phase. It illustrates how, in this example, a small, adaptive CI population (CI = *aa*0*X*, initial population ratio 0.01) can invade and reach a population ratio of 98% in niche *A* after 60 generations. Consequently, in the *Ax*/*Bx* phase portrait, this invasion shifts the fixed point closer to the lower-right corner (from the green to the red dot), resulting in stronger RI between the parental niche ecotypes.

Fig 10 shows the *Ax*/*Bx* phase portrait solution when the *Single Line* plot option is selected in Fig 8. It illustrates how a higher-mating-bias *X* allele (*α* = 0.3) arising in niche *A* with an initial population ratio of 0.01 (as specified in the Initial Genotype Distribution table in Fig 9) is able to invade a population carrying only the *Y* mating-bias allele and reach a fixed-point polymorphism (green dot), thereby producing initial premating RI. Subsequently, a CI mutation (*CI* = *aa*0*X*) invades, reaches a high population ratio in niche *A*, and shifts the *Ax*/*Bx* fixed point closer to the lower-right corner of the phase portrait, resulting in even stronger RI.

**Fig 10.**
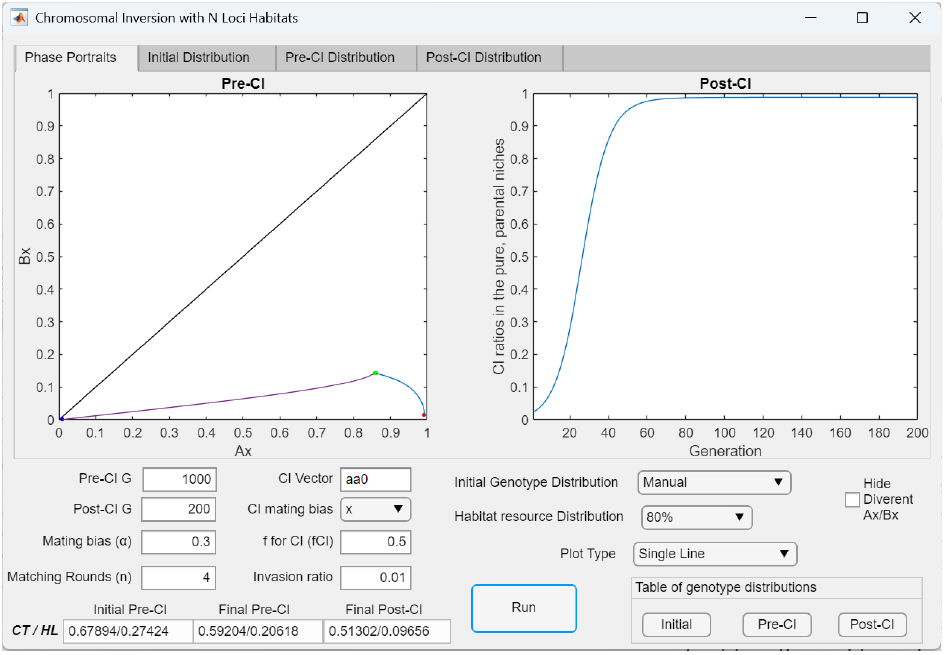
Single-line plots showing an *X* mutant mating-bias allele invading niche *A* and reaching a fixed-point polymorphism before an adaptive CI subsequently invades and further strengthens premating RI. In Fig 8, selecting the *Single Line* plot option displays the invasion trajectory of a high–mating-bias *X* mutant allele in niche *A* during the pre-CI phase, followed by the invasion trajectory of the CI mutant (*CI* = *aa*0*X*) during the post-CI phase. In the *Ax*/*Bx* phase portrait, the initial *Ax* population ratio (*Ax*/*Bx* = 0.01/0) is shown as a blue dot. It reaches a fixed-point polymorphism after 1000 generations, marked by the green dot. The subsequent invasion of the CI shifts this fixed point closer to the lower-right corner, marked by the red dot, indicating stronger premating RI.

Displayed at the bottom of Fig 10 are the *CT* and *HL* values corresponding to the initial pre-CI, final pre-CI, and final post-CI stages of the simulation. Here, *CT* represents the total ratio of offspring produced from inter-niche matings—including both hybrid and parental (niche *A* and niche *B*) offspring— relative to the total number of offspring produced per generation from both intra- and inter-niche matings. Thus, *CT* serves as a measure of the degree of premating RI between the parental niche ecotypes.

*HL* represents the total ratio of hybrid offspring loss relative to the total offspring produced. It is calculated by subtracting the total hybrid niche resource ratio (as specified in the Habitat Distribution table in Fig 9) from the total hybrid offspring ratio. *HL* therefore quantifies the proportion of hybrid offspring that exceed the available hybrid niche resources (niche carrying capacities) and fail to survive. As such, it reflects the strength of disruptive ecological selection—or the extent of maladaptive hybrid loss— that drives the evolution of reproductive barriers in sympatric speciation.

In Fig 10, the initial pre-CI *CT* and *HL* values correspond to the state before the invasion of the pre-CI mutant (*Ax* = 0.01), as specified in the Initial Genotype Distribution table in Fig 9. The final pre-CI and final post-CI values represent the *CT* and *HL* ratios at the end of the pre-CI and post-CI phases, respectively. As shown, both *CT* and *HL* decrease steadily following the invasion of the *Ax* and CI mutants, resulting in ever stronger RI between the parental niche ecotypes.

Selecting the *Initial Distribution* tab at the top of the user interface in Fig 10 displays the bar graph in Fig 11, which shows the initial genotype distribution before the pre-CI phase of the simulation. In this example, the displayed ratios correspond to the values specified in the Habitat Distribution Table in Fig 9. Similarly, selecting the *Pre-CI Distribution* tab displays the bar graph in Fig 12, showing the genotype distribution after the pre-CI phase of the simulation. Finally, selecting the *Post-CI Distribution* tab displays the bar graph in Fig 13, showing the final genotype distribution after the post-CI phase. In Fig 13, the blue bars represent the ratios of the *X* allele in uninverted genotypes, the red bars represent the ratios of the *Y* allele in uninverted genotypes, and the black bars represent the ratios of the CI in the genotypes. These ratios are displayed in the order *X, Y*, and CI for each genotype.

**Fig 11.**
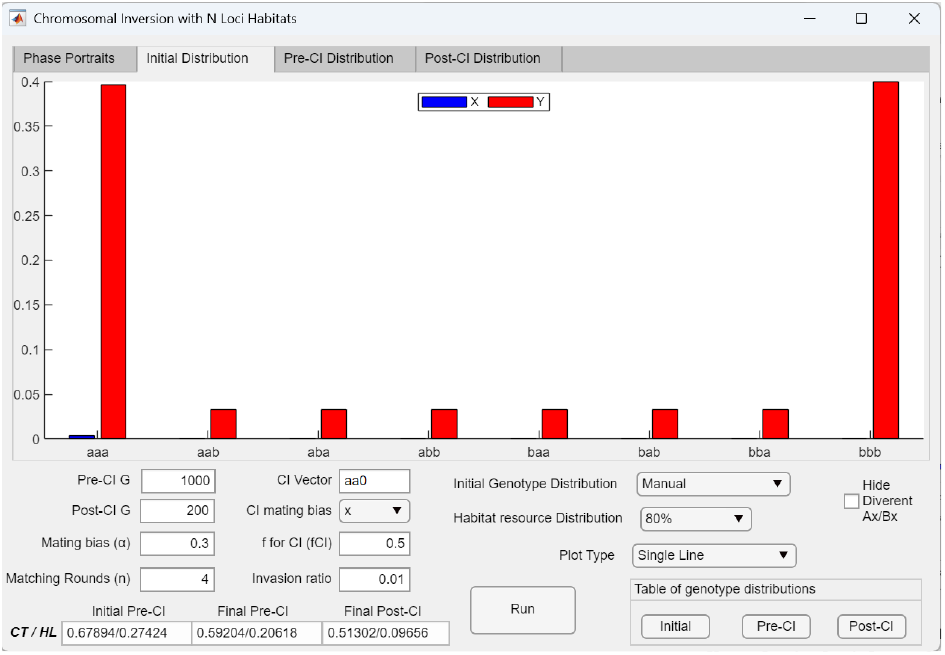
Bar graph displaying the user-specified habitat resource distribution in a model with three-locus ecological genotypes. Selecting the *Initial Distribution* tab at the top of the user interface in Fig 10 displays the specified niche resource distribution. In this example, the “80%” habitat resource option allocates 80% of the total niche resources equally between the parental niches *A* and *B*, while the remaining 20% are divided equally among all hybrid niches.

**Fig 12.**
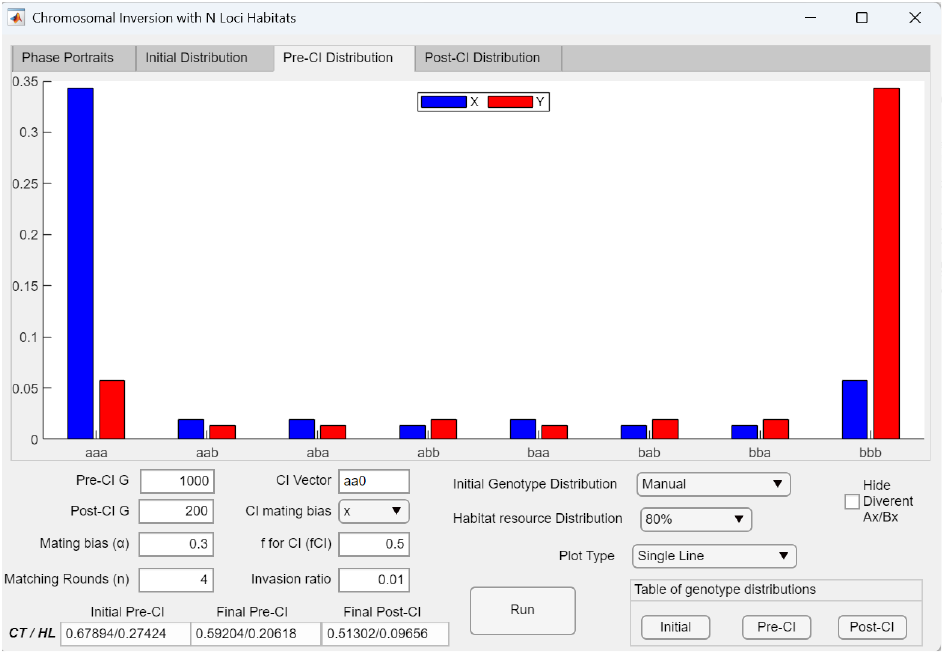
Bar graph showing the genotype compositions across all ecological niches after the pre-CI phase in a model with three-locus ecological genotypes. Selecting the *Pre-CI Distribution* tab at the top of the user interface in Fig 10 displays the genotype compositions in all ecological niches after 1000 pre-CI generations, prior to the invasion of the CI.

**Fig 13.**
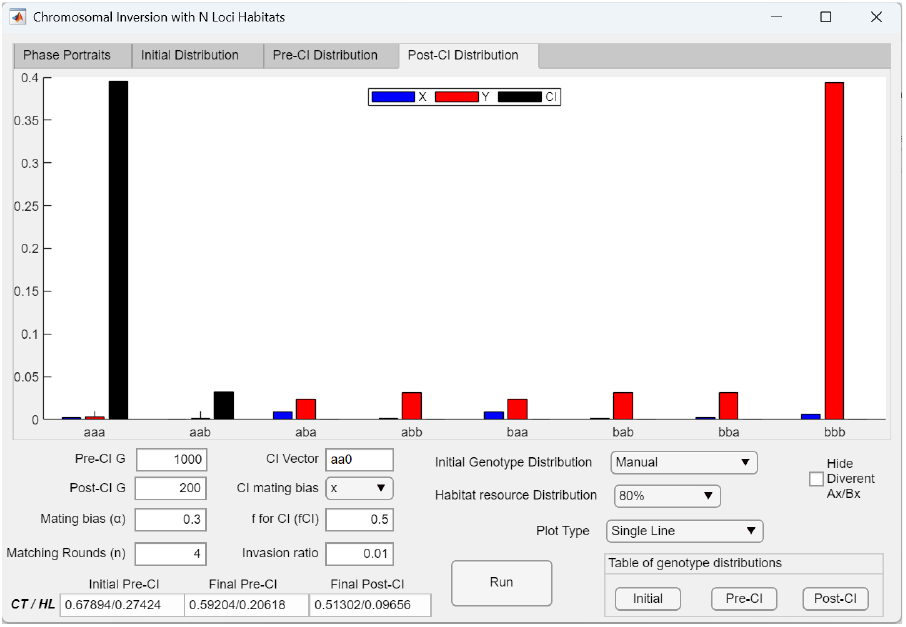
Bar graph showing the genotype compositions across all ecological niches after the post-CI phase in a model with three-locus ecological genotypes. Selecting the *Post-CI Distribution* tab at the top of the user interface in Fig 10 displays the genotype compositions across all ecological niches after 200 post-CI generations. The black bars in the graph represent the ratios of the CI in each niche, while the blue and red bars indicate the ratios of the *X* and *Y* alleles in the uninverted niche genotypes. As shown, the CI (*aa*0*X*) is able to invade niches *aaa* and *aab* and attain high ratios in a polymorphism. The CI does not reach fixation, however, because mating among viable hybrid populations continues to regenerate the uninverted niche *A* ecotype.

### 3 Extended model of multilocus mating-bias genotypes without viable ecological hybrids

We developed a GUI application (MultiSexCI) based on the mathematical framework illustrated in Fig 3, extending the model to accommodate any specified number of gene loci for the mating-bias genotypes. When the number of mating-bias loci exceeds one, hybrid mating-bias genotypes can arise within the niche-*A* and niche-*B* ecotype populations. Because no disruptive selection—either ecological or sexual—acts against mating-bias hybrids within the same ecological niche, all such mating-bias hybrids remain viable. Their presence increases the effective value of *α* in the Fig 3 model, thereby diminishing the strength of sexual selection and hindering the emergence of RI between niche ecotypes.

The application also models CI mutations that capture any arbitrarily defined set of mating-bias alleles. This makes it possible to examine the invasion dynamics of these CIs and assess how their invasion may alter system behavior—either promoting or constraining the development of RI.

For ecological selection, the model adopts the same framework used in the Fig 3 mathematical model. It assumes that the niche-*A* and niche-*B* ecotypes do not produce viable ecological hybrids and that the offspring return ratio (*f*) specifies the strength of disruptive ecological selection against ecological hybrids.

Fig 14 shows the computer interface of the GUI application. The number of letters in the CI vector input determines the number of gene loci for the mating-bias genotype. Each mating-bias gene locus carries one of two allele types, *X* or *Y*. The extreme parental genotypes consist entirely of *X*-type alleles (e.g., *xxx*) or entirely of *Y*-type alleles (e.g., *yyy*), while hybrid genotypes contain mixtures of the two (e.g., *xyx, yyx*, etc.). In this notation, the *x* and *y* alleles represent distinct entities at different loci, even though the same letters are used for simplicity. In the example shown in Fig 14, the CI vector is “*xx*0”, indicating three mating-bias gene loci, with the CI capturing the *x* alleles at the first two loci. The digit “0” in the third position indicates that no allele is captured at that locus. Each individual also carries either the niche-*A* or niche-*B* ecotype, as the model assumes that ecological hybrids are not viable.

**Fig 14.**
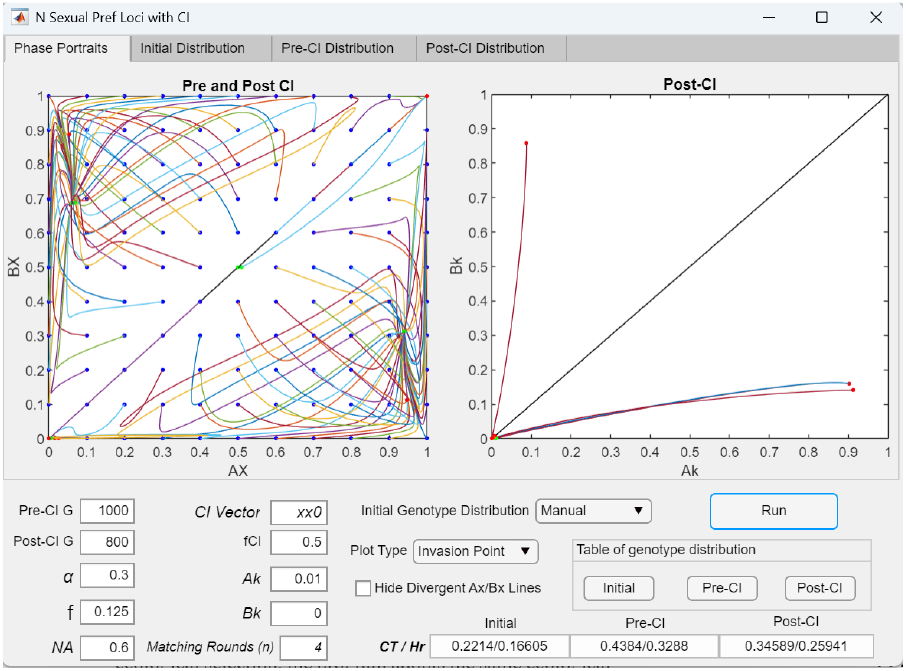
User interface of the application (MultiSexCI) used to model multilocus mating-bias genotypes and the invasion dynamics of CIs that capture mating-bias alleles. The application models a sympatric population with multilocus mating-bias genotypes. For ecological selection, it adopts the same framework as the model without viable ecological hybrids shown in Fig 3. The variables *f* (offspring return ratio), *NA* (normalized niche *A* carrying capacity), and *n* (number of matching rounds) are defined as in Fig 3. For sexual selection, the number of letters in the *CI vector* specifies the number of gene loci in the mating-bias genotypes. Each locus carries either the *X* or *Y* allele, and the CI captures either the *X* or *Y* allele—or “0” to indicate no capture. The variable *α* specifies the maximum mating bias between the pure genotypes (e.g., *xxx* and *yyy* in this three-locus example). Mating biases between different genotypes are determined by their genetic distances and are listed in the Matching Compatibility Table (Fig 15). The variable *fCI* specifies the offspring return ratio of the CI. *Ak* and *Bk* are the initial CI mutant population ratios in niches *A* and *B*, respectively. *Pre*-*CI G* and *Post*-*CI G* specify the number of generations simulated in the pre-CI and post-CI phases. Selecting the *Manual* option in *Initial Genotype Distribution* opens the table in Fig 16, allowing users to define the initial genotype distribution before the pre-CI phase begins. The figure displays the *Invasion Point* plot option. In the *AX*/*BX* phase portrait, *AX* is calculated as the normalized ratio of the pure *X* genotype (*xxx*) in niche *A* relative to the total of pure *X* and *Y* genotypes in that niche, excluding all hybrid genotypes (Eq. 1). *BX* is calculated in the same way for niche *B* (Eq. 2). Given the specified parameters, the *AX*/*BX* phase portrait shows all initial population ratios in the vector field (blue dots) converging to fixed points (green dots) during the pre-CI phase. At the start of the post-CI phase, a CI mutant (CI = *xx*0), represented by allele *K* with an initial population ratio of *Ak* = 0.01, successfully invades and reaches a fixed-point polymorphism (red dot at *Ak*/*Bk* = 0.9125/0.1412) in the *Ak*/*Bk* phase portrait. Consequently, the final pre-CI *AX*/*BX* fixed point in niche *A* (green dot at *AX*/*BX* = 0.9407/0.3086) shifts toward the lower-right corner (red dot at *AX*/*BX* = 0.9416/0.1014), indicating stronger RI between the niche-*A* and niche-*B* ecotypes. The pre-CI *AX*/*BX* phase portrait shows an invasion-resistant pattern.

The matching success between individuals of different mating-bias genotypes is defined in a Matching Compatibility Table. Fig 15 shows a representative example for a three-locus system. In the table, mating-bias genotypes are arranged according to their genetic similarity (genetic distance). In Fig 14, the variable *α* specifies the maximum mating bias attainable in the system (e.g., *α* = 0.3 in the Fig 15 table), which occurs between encounters of the extreme parental genotypes (*xxx* and *yyy*) located at the lower-left and upper-right corners of the table. The matching probabilities increase linearly as genotype pairings move from the corners toward the diagonal, where identical mating-bias genotypes always match with a probability of 1 (*α* = 1).

**Fig 15.**
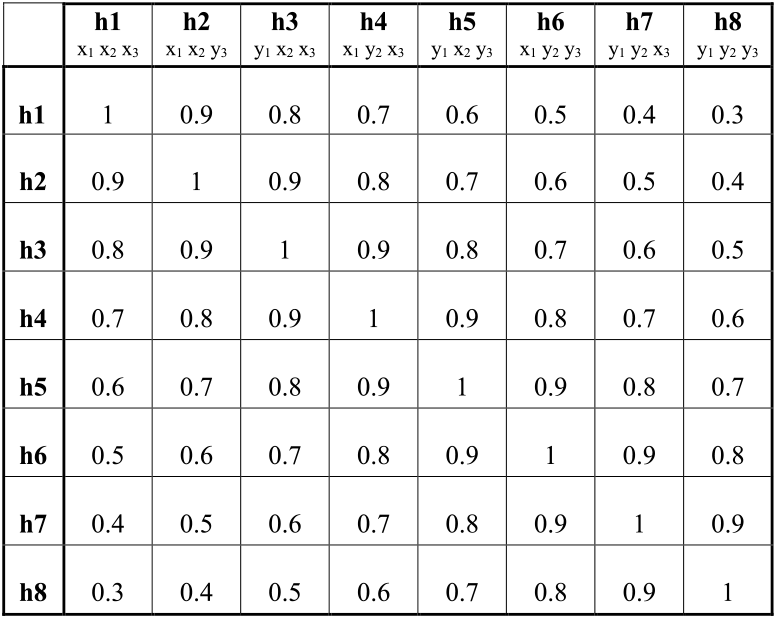
Matching Compatibility Table for three-locus mating-bias genotypes. Each mating-bias genotype consists of three gene loci, with each locus carrying either the *X* or *Y* allele. In the table, *h*1 to *h*8 represent the eight possible genotype permutations, ordered by their genetic distances from one another. The pure parental genotypes are *h*1 (*xxx*) and *h*8 (*yyy*), while the remaining entries represent hybrid genotypes. In this example, *α* = 0.3 denotes the maximum mating bias in the system; it also represents the mating bias between the extreme parental genotypes *h*1 (*xxx*) and *h*8 (*yyy*). Mating-bias values increase linearly toward the diagonal entries, where identical genotypes always match with a probability of 1.

Like the GUI application for multilocus ecological genotypes, the GUI application for multilocus mating-bias genotypes operates in two phases: the pre-CI phase followed by the post-CI phase. In Fig 14, the variables *Pre*-*CI G* (number of generations in the pre-CI phase), *Post*-*CI G* (number of generations in the post-CI phase), *n* (number of matching rounds), *Ak* and *Bk* (initial CI population ratios in niches *A* and *B*), and *fCI* (offspring return ratio of the CI) are defined in the same way as in the ecological genotype model shown in Fig 8. Because no ecological hybrid niche resources are present in the multilocus mating-bias genotype model, the variable *NA* in Fig 14 specifies the normalized niche carrying capacity for niche *A*, while the carrying capacity for niche *B* (*NB*) is 1 −*NA*.

Selecting the *Manual* option in *Initial Genotype Distribution* displays the popup table shown in Fig 16, which allows the user to specify the initial mating-bias genotype distributions before the pre-CI phase of the simulation. In this example, all individuals in niche *A* carry the *xxx* genotype, and all individuals in niche *B* carry the *yyy* genotype. This setup could represent a scenario of secondary contact, in which two allopatrically isolated populations that have evolved different mating biases come into sympatry.

**Fig 16.**
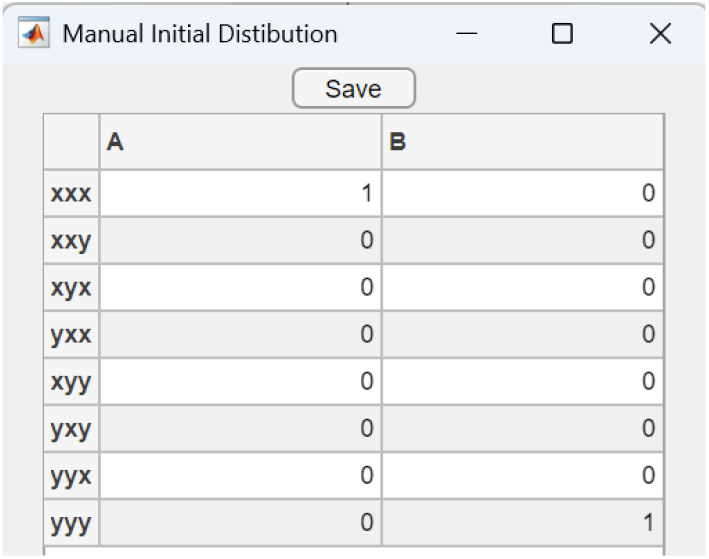
Initial Genotype Distribution Table for three-locus mating-bias genotypes. Selecting the *Manual* option for *Initial Genotype Distribution* displays a popup table that allows the user to specify the initial mating-bias genotype distribution in each niche before running the pre-CI phase of the simulation. In this table, all ratios in the two vertical columns—representing the normalized genotype distributions in niches *A* and *B*—must sum to 1.

The GUI application provides two plot options: *Invasion Point* and *Single Line*. Fig 14 shows the *Invasion Point* plot option. In the *AX*/*BX* phase portrait, the values of *AX* (*x*-axis) and *BX* (*y*-axis) are calculated as follows.

For *AX*, all hybrid mating-bias genotypes present in niche *A* are ignored, and only the pure, parental extreme genotypes are considered (e.g., *xxx* and *yyy* in a three-locus system). *AX* is then defined as the normalized ratio of the pure *X* genotype in niche *A* (Eq. 1):

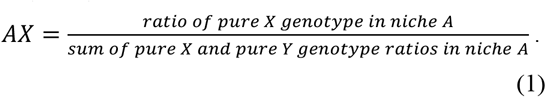

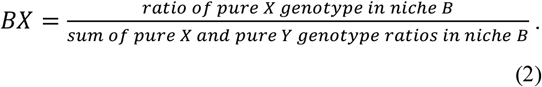

It follows that the normalized ratio of the pure *Y* genotype in niche *B* is: *BY* = 1 − *BX*.

It is important to note that the *AX*/*BX* phase portrait is not an exact projection but rather a lower-dimensional snapshot of the population dynamics that may be occurring in the higher-dimensional phase portrait determined by the number of mating-bias gene loci. For example, a complete visualization of a three-locus system would require a 14-dimensional phase portrait (2^*N*+1^ − 2, where *N* is the number of loci). Nevertheless, plotting the ratios of pure, extreme mating-bias genotypes in each niche could provide useful insight into the degree of random assortment and premating RI between the niche ecotypes.

We also expect that convergence to a fixed point in this reduced two-dimensional *AX*/*BX* phase portrait corresponds to convergence to a fixed point in the full, higher-dimensional phase portrait that includes all hybrid genotype permutations. However, the reverse is not necessarily true—nonconvergence in the *AX*/*BX* phase portrait does not preclude the existence of a fixed point consisting solely of hybrid genotypes in the higher-dimensional phase portrait. Nonetheless, by selecting the *Initial Distribution, Pre-CI Distribution*, and *Post-CI Distribution* tabs in the *Single Line* plot (Fig 17), users can visualize the complete mating-bias genotype distributions at different stages of the simulation (Figs 18–20).

**Fig 17.**
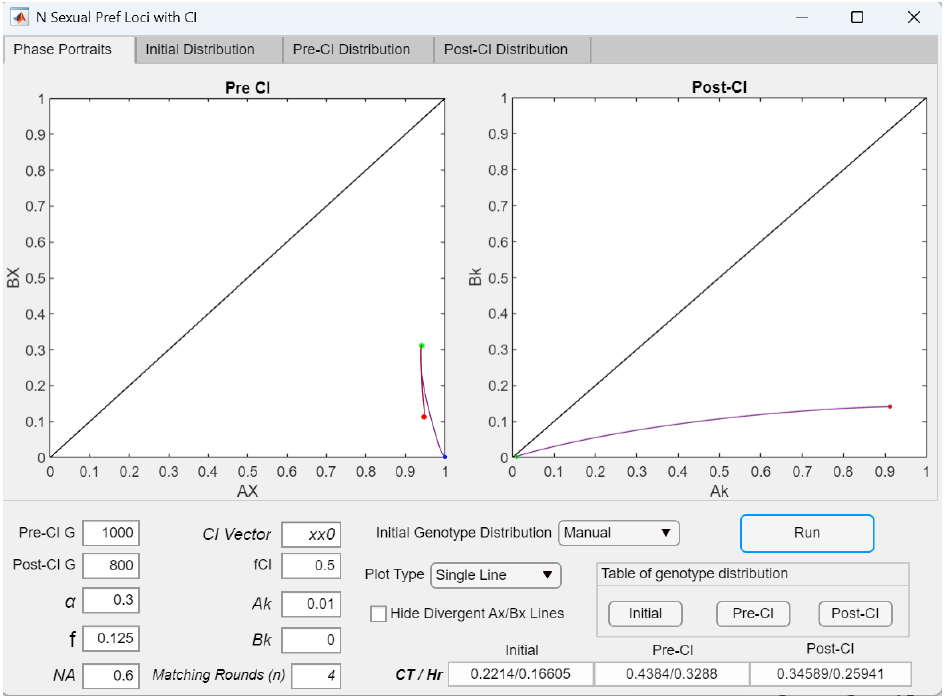
Single-line plots showing how the presence of mating-bias hybrids decreases RI, whereas the invasion of a CI capturing high mating-bias alleles increases RI. When the *Single Line* plot option is selected in Fig 14, the *AX*/*BX* phase portrait plots the single-line vector trajectory of the initial genotype ratio (*AX*/*BX* = 1/0, as specified in Fig 16) across the pre-CI and post-CI phases of the simulation. Similarly, the *Ak*/*Bk* phase portrait plots the single-line invasion trajectory of the CI mutation (i.e., the *K* mutant allele in niche *A*). Displayed at the bottom of the figure are the *CT* and *Hr* values at different stages of the simulation. The variable *CT* (total inter-niche offspring) is defined as in Fig 10 and serves as a measure of inter-niche mating and premating RI. The variable *Hr* represents the total ratio of ecological hybrid offspring produced from inter-niche matings. Because ecological hybrids are not viable in this model, *Hr* is equivalent to *HL* in Fig 10 and reflects the degree of ecological hybrid loss that drives the evolution of RI between niche ecotypes. As shown, *CT* and *Hr* increase after the pre-CI phase because the emergence of viable mating-bias hybrid genotypes reduces the effective mating bias and weakens sexual selection, thereby diminishing premating RI. In the post-CI phase, however, the invasion of the adaptive CI (*CI* = *xx*0) reduces the effective number of mating-bias loci and limits the production of mating-bias hybrids. This shifts the final pre-CI *AX*/*BX* fixed point (green dot) toward the lower-right corner of the phase portrait (red dot), resulting in stronger RI between niche-*A* and niche-*B* ecotypes.

**Fig 18a.**
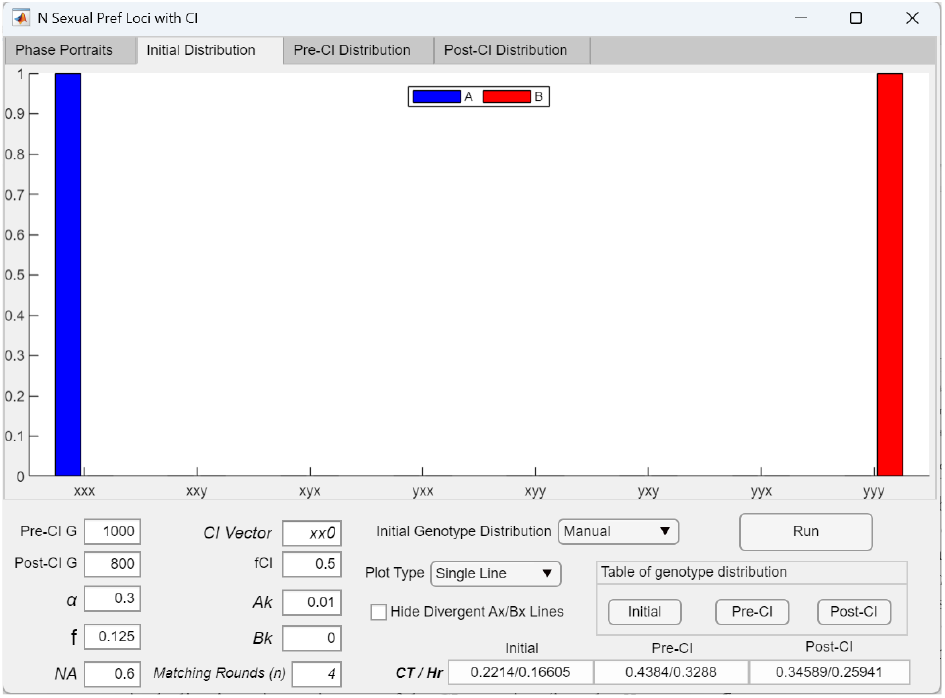
Bar graph showing the user-specified initial mating-bias genotype distribution before the pre-CI phase in a model with three-locus mating-bias genotypes. Selecting the *Initial Distribution* tab at the top of the user interface in Fig 17 displays the mating-bias genotype distributions in the two ecological niches, *A* and *B*, as specified by the values in the Initial Genotype Distribution Table shown in Fig 16.

**Fig 18b.**
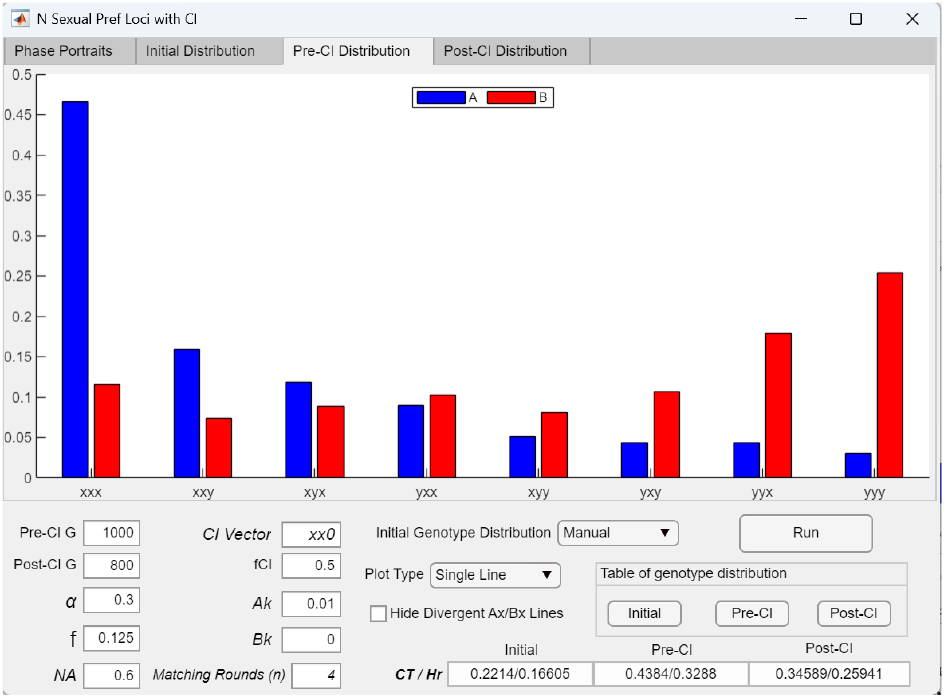
Bar graph showing the mating-bias genotype compositions across all ecological niches after the pre-CI phase in a model with three-locus mating-bias genotypes. Selecting the *Pre-CI Distribution* tab at the top of the user interface in Fig 17 displays the mating-bias genotype compositions in niches *A* and *B* after 1000 pre-CI generations, before the invasion of the CI. Blue bars represent the niche *A* ecotype, and red bars represent the niche *B* ecotype.

**Fig 18c.**
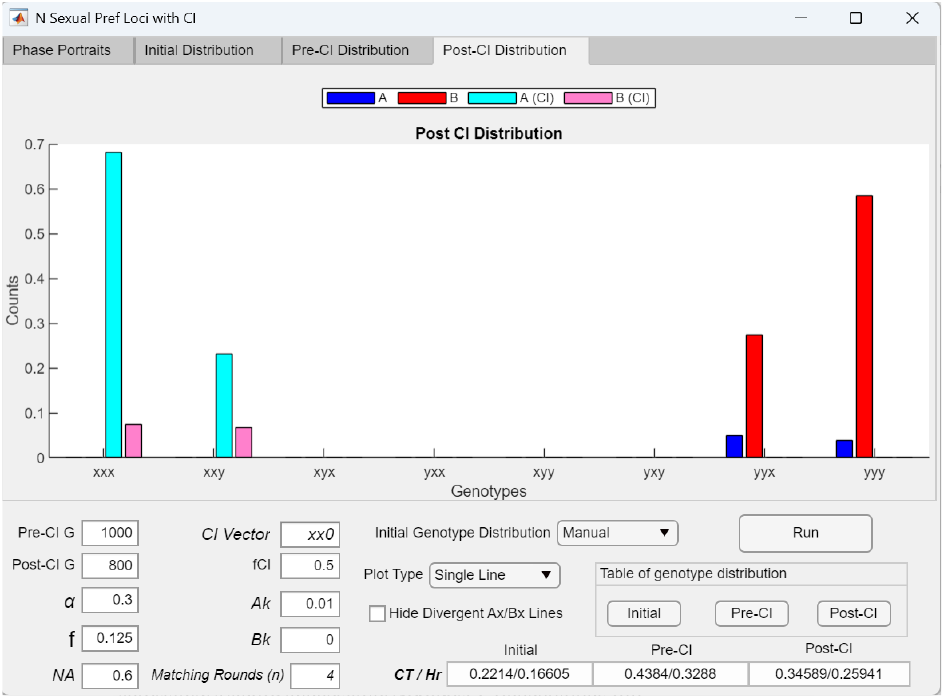
Bar graph showing the genotype compositions across all ecological niches after the post-CI phase in a model with three-locus mating-bias genotypes. Selecting the *Post-CI Distribution* tab at the top of the user interface in Fig 17 displays the mating-bias genotype compositions across the *A* and *B* niches after 800 post-CI generations. In the bar graph, the cyan bars represent the ratios of mating-bias genotypes carrying the CI in niche *A*, and the magenta bars represent those carrying the CI in niche *B*. The blue and red bars indicate the ratios of uninverted (non-CI) mating-bias genotypes in niches *A* and *B*, respectively. As shown, the CI (CI = *xx*0) mutation successfully invades and reaches high ratios in the niche-*A* genotypes *Axxx* and *Axxy*, which primarily carry high-bias *X* alleles, while in niche *B*, the uninverted genotypes *Byyy* and *Byyx*, which mainly carry *Y* alleles, dominate. As a result, the CI invasion enhances the divergent assortment of mating-bias genotypes between the two niches, increasing the proportions of opposite extreme genotypes (*X*-biased in niche *A* and *Y*-biased in niche *B*) while reducing the production of intermediate mating-bias hybrids. This divergence strengthens the effective mating bias and increases RI between the niche ecotypes.

During the pre-CI phase of the simulation, the *AX*/*BX* phase portrait plots the vector-field trajectories of all possible permutations of *AX*/*BX* initial population ratios (blue dots), spaced at 0.1 intervals, over the specified pre-CI generations. Each initial population is generated by varying *AX* and *BX* from 0 to 1, at 0.1 intervals, to produce 121 possible combinations of pure, extreme mating-bias genotypes (e.g., *AX* = *Axxx* and *BX* = *Bxxx*, where *BX* = 1 − *BY* and *BY* = *Byyy*), while keeping all hybrid genotype ratios unchanged as specified in the Initial Genotype Distribution table (Fig 16). In calculating these initial populations, the total ratios of pure genotypes (*xxx* and *yyy*) allocated to each niche— namely, the total population proportions of *AX* + *AY* in niche *A* and *BX* + *BY* in niche *B*, as specified in Fig 16—remain fixed; only the relative ratios of *AX* to *AY* and of *BX* to *BY* within each niche are varied.

As shown in the three-locus mating-bias genotype example in Fig 14, the presence of viable mating-bias hybrids produces an invasion-resistant pattern in the *AX*/*BX* phase portrait: initial population ratios near the origin cannot invade and fall back toward the origin, whereas ratios outside this region converge to fixed-point polymorphisms (green dots).

At the start of the post-CI phase, a CI mutation (*CI* = *xx*0) arises in niche *A* with an initial population ratio of *Ak*/*Bk* = 0.01/0. Here *Ak* is the ratio of the niche-*A* ecotype carrying the CI, and *Bk* is the ratio of the niche-*B* ecotype carrying the CI. During the post-CI phase, the program plots all final pre-CI *AX*/*BX* population ratios (green dots) against this initial *Ak* population ratio over the specified post-CI generations. The resulting final post-CI *AX*/*BX* population ratios are shown as red dots. The CI mutant successfully invades and reaches fixed-point polymorphism in the *Ak*/*Bk* phase portrait. Consequently, the final pre-CI *AX*/*BX* fixed point (green dot) shifts to a new post-CI fixed point (red dot) closer to the lower-right corner of the *AX*/*BX* phase portrait, indicating stronger premating RI between the niche ecotypes.

Fig 17 displays the single-line *AX*/*BX* and *Ak*/*Bk* phase portrait solutions when the *Single Line* plot option is selected in Fig 14. The *AX*/*BX* phase portrait shows the trajectory of the initial mating-bias genotype populations (as specified in the Initial Genotype Distribution Table in Fig 16) during the pre-CI phase, while the *Ak*/*Bk* phase portrait shows the invasion trajectory of the initial CI mutant population (*Ak* = 0.01) during the post-CI phase.

The *CT* and *Hr* values displayed at the bottom of Fig 17 represent, respectively, the total offspring ratio produced from inter-niche matings (*CT*) and the ecological hybrid offspring ratio (*Hr*) at different stages of the simulation. Similar to *CI* and *HL* in the multilocus ecological genotype model (Fig 10), *CT* quantifies the extent of inter-niche mating and the strength of premating RI, while *Hr* quantifies the degree of ecological hybrid loss that drives the evolution of reproductive barriers.

In Fig 17, *CT* and *Hr* initially increase following the pre-CI phase because the emergence of viable mating-bias hybrids weakens sexual selection and reduces RI. In the post-CI phase, however, the invasion of the locally adaptive CI (CI = *xx*0) reduces the effective number of mating-bias loci, lowers the production of mating-bias hybrids, and strengthens overall RI between the niche ecotypes.

Selecting the *Initial Distribution* tab at the top of the user interface in Fig 17 displays the bar graph in Fig 18a, which shows the initial genotype distribution before the pre-CI phase of the simulation. In this example, the displayed ratios correspond to the values specified in the Initial Genotype Distribution Table in Fig 16. Similarly, selecting the *Pre-CI Distribution* tab displays the bar graph in Fig 18b, showing the mating-bias genotype distribution after the pre-CI phase of the simulation. Finally, selecting the *Post-CI Distribution* tab displays the bar graph in Fig 18c, showing the final mating-bias genotype distribution after the post-CI phase. In Fig 18c, the blue bars represent the ratios of niche-*A* ecotypes with uninverted (non-CI) mating-bias genotypes, and the red bars represent the ratios of niche-*B* ecotypes with uninverted genotypes. The cyan bars represent the ratios of niche-*A* ecotypes carrying the CI, while the magenta bars represent the ratios of niche-*B* ecotypes carrying the CI. These ratios are displayed in the order *A*(*non*-*CI*), *B*(*non*-*CI*), *A*(*CI*), and *B*(*CI*) for each mating-bias genotype.

### 4 Two-allele model of multilocus postzygotic barriers

Lastly, our GUI applications were modified to investigate the coupling dynamics of a two-allele model of postzygotic barriers in sympatric populations under disruptive ecological selection. Our analyses focus on how these dynamics change when postzygotic isolation is mediated by multiple gene loci and when CIs capture different combinations of incompatibility alleles.

Consider two ecotype populations, *A* and *B*, that have diverged in allopatry and subsequently come into secondary contact in sympatry. Individuals carry two population-specific background loci, each with two alleles. During geographic isolation, ecotype-*A* individuals predominantly accumulated mutually compatible alleles *p*1 and *p*2, whereas ecotype-*B* individuals predominantly accumulated compatible alleles *q*1 and *q*2. We assume that there are no premating barriers and that alleles are inherited by random assortment at each gene locus. Upon secondary contact, matings between ecotype-*A* and ecotype-*B* individuals generate offspring with six possible multilocus genotypes: *p*1*p*2, *p*1*q*1, *p*1*q*2, *p*2*q*1, *p*2*q*2, and *q*1*q*2. If mismatched allele combinations (*p*1*q*1, *p*1*q*2, *p*2*q*1, or *p*2*q*2) interact through negative epistasis, hybrid genotypes carrying these combinations experience reduced viability.

We classify genotypes carrying *p*1*p*2 as possessing incompatibility allele *P* and those carrying *q*1*q*2 as possessing incompatibility allele *Q*. All mixed *p*–*q* genotypes are then treated as hybrid offspring produced by matings between individuals carrying *P* and *Q* alleles. Under this formulation, interactions between background alleles *P* and *Q* generate hybrid incompatibilities through the same mechanism that produces postzygotic isolation in the Bateson– Dobzhansky–Muller (BDM) model [42-47].

We can construct a Hybrid Incompatibility Table shown in Fig 19 that is similar to the matching compatibility table in Fig 4 to specify the degree of hybrid viability produced by matings between individuals carrying the *P* and *Q* alleles. Crosses between individuals carrying the same incompatibility allele (*P* × *P* or *Q* × *Q*) produce fully viable offspring, whereas crosses between individuals carrying different incompatibility alleles (*P* × *Q*) produce offspring with viability reduced by a factor *v*, which we refer to as the viability ratio and which ranges from 0 to 1. The parameter *v* therefore quantifies the strength of postzygotic reproductive isolation, with lower values corresponding to stronger incompatibilities and reduced hybrid survival.

**Fig 19.**
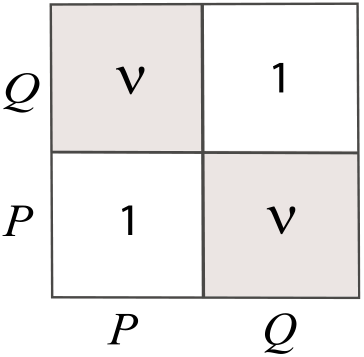
Hybrid Incompatibility Table for background *P* and *Q* alleles. Individuals carrying the same allele (*P* or *Q*) produce viable offspring with probability 1, whereas matings between individuals carrying different alleles produce viable offspring with probability *v* (0 ≤ *v* ≤ 1).

From the parallelism between the premating and postmating barrier models, it follows that they can be treated as mathematically equivalent in our computational implementation (see Appendix for derivation). Accordingly, our existing GUI applications can be used to simulate postzygotic incompatibility by substituting variables *X* and *Y* with *P* and *Q*, replacing the mating-bias parameter *α* with the viability ratio *v*, and setting the number of mating rounds (*n*) to one. This restriction to a single mating round reflects the key difference between the two processes: under mating-bias selection, unmatched individuals may still mate in subsequent rounds when *n* is greater than one, whereas under hybrid incompatibility selection, inviable hybrid offspring (occurring with probability 1 − *v* ) are eliminated after the first mating event and have no further opportunity to survive or reproduce. Additionally, in the absence of premating barriers, all individuals in a sympatric population match in the first matching round (*n* = 1), and no unmatched individuals remain for additional matching rounds.

Fig 20 shows that, after substituting the relevant variables in Fig 7 and setting *n* to one, a region of initial *Ap*/*Bp* population ratios within the lower-right quadrant of the phase portrait converges to a stable fixed point under the specified parameter values. This result indicates that, during secondary contact, disruptive ecological selection can increase the nonrandom assortment of hybrid-incompatibility alleles between two allopatrically diverged ecotypes by driving the system toward a fixed-point polymorphism nearer the lower-right corner of the phase portrait. Consequently, postzygotic RI between niche ecotypes is strengthened despite the presence of gene flow.

**Fig 20.**
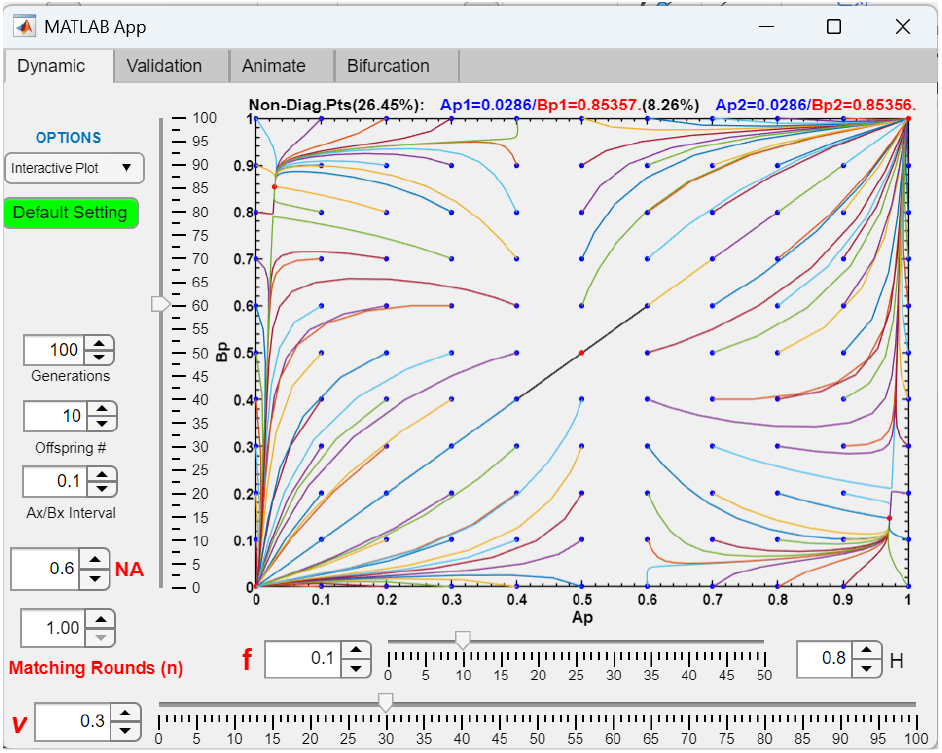
*Ap*/*Bp* phase-portrait dynamics under divergent incompatibility alleles. The phase portrait shows that when ecotype-*A* and ecotype-*B* populations already carry high frequencies of different incompatibility alleles (*P* and *Q*, respectively), initial *Ap*/*Bp* population ratios within the lower-right quadrant converge to a stable fixed point. For the specified parameter values, disruptive ecological selection can drive *Ap*/*Bp* ratios that start farther from the lower-right corner toward a fixed point closer to it, thereby strengthening postzygotic RI between niche ecotypes. In contrast, when *Ap*/*Bp* ratios are already near the lower-right corner, gene flow can shift the equilibrium to a fixed point farther from this region, resulting in weaker postzygotic RI.

Similarly, our existing GUI application for multilocus mating-bias alleles can be adapted to model postzygotic RI caused by multilocus incompatibility alleles. This is achieved by substituting *P* for *X* and *Q* for *Y*, replacing the maximum mating-bias parameter *α* with an analogous minimum viability ratio *v*, and setting the number of matching rounds to one (*n* = 1). Under this implementation, offspring genotypes first experience intrinsic incompatibility selection determined by *v* before undergoing extrinsic disruptive ecological selection governed by *f* (see Appendix for a description of the modified algorithm).

The modified GUI application (MultiCompCI) displays the *AP*/*BP* phase portrait, which tracks the population dynamics of the extreme-*P* and extreme-*Q* parental genotypes (i.e., *pppp* and *qqqq*), analogous to the *AX*/*BX* phase portrait used for the extreme-*X* and extreme-*Y* mating-bias genotypes. This same application is also used to investigate the invasion dynamics of chromosomal inversions that capture different combinations of hybrid-incompatibility alleles.

## III. RESULTS

Using the three GUI applications described in the Methodology section (MultiEcoCI, MultiSexCI, and MultiCompCI), we examined how multilocus ecological, mating-bias, and hybrid-incompatibility genotypes influence the population dynamics of the two-niche, two-allele model (Fig 3) developed in our previous study [39]. We further investigated how the invasion of a chromosomal inversion (CI) capturing locally adaptive ecological, mating-bias, and hybrid-incompatibility alleles in such polygenic systems may either facilitate or hinder the development of reproductive isolation (RI) between sympatric niche ecotypes under disruptive ecological selection.

### I. Systems with multilocus ecological genotypes and single-locus mating-bias genotypes

We first present the results obtained using the MultiEcoCI application to examine the effects of incorporating an arbitrary number of ecological gene loci and viable hybrid populations. To focus solely on the pre-CI dynamics, the post-CI phase of the simulation was disabled by setting the number of post-CI generations (*Post*-*CI GPn*) to zero. The proportions of hybrid populations were specified in the Habitat Resource Distribution Table (Fig 9), where hybrid niche resources were included to support viable hybrid genotypes. In subsequent simulations, the post-CI phase was enabled, and CI mutations capturing different combinations of ecological alleles were introduced to examine their invasion dynamics and effects on system behavior.

#### 1 The effects of having multilocus ecological genotypes and viable ecological hybrids (without CI invasion) on one-locus mating-bias barriers

By setting the number of post-CI generations (*Post*-*CI GPn*) to zero, we can examine the effects of having any specified number of ecological gene loci and the presence of viable ecological hybrids in the model. In general, viable ecological hybrids sustained by hybrid niche resources increase the effective offspring return ratio (*f*) in the Fig 3 model, thereby reducing disruptive ecological selection and weakening RI between the parental niche *A* and *B* ecotypes. This effect is illustrated in Fig 21. Compared with the example shown in Fig 8, when hybrid niche resources and viable hybrid offspring are removed from the system in Fig 21, the *Ax*/*Bx* fixed point shifts closer to the lower-right corner of the phase portrait, and the total inter-niche offspring ratio (*CT*) decreases, indicating reduced inter-niche mating and stronger RI between niche ecotypes.

**Fig 21.**
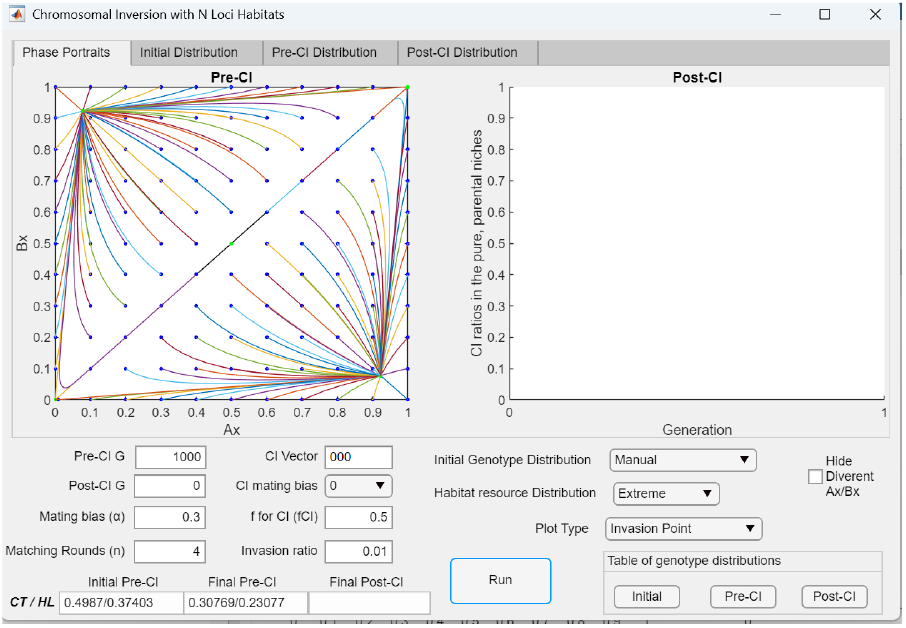
Eliminating viable ecological hybrids strengthens premating RI. Selecting the *Extreme* option for *Habitat Resource Distribution* in Fig 8, while keeping all other parameter values unchanged during the pre-CI phase of the simulation, removes all hybrid niche resources. As a result, compared with Fig 8, the *Ax*/*Bx* fixed point shifts closer to the lower-right corner of the phase portrait, indicating stronger RI. The final pre-CI total inter-niche offspring ratio (*CT*) decreases from *CT* = 0.59204 in Fig 8 to *CT* = 0.30769 after the elimination of viable hybrids, reflecting reduced inter-niche mating and enhanced premating RI between the niche ecotypes.

As illustrated in Fig 22, under disruptive ecological selection against hybrids, increasing the number of ecological gene loci (from three in Fig 8 to four in Fig 22) tends to lower the effective value of *f* and lead to stronger premating RI between niche ecotypes.

**Fig 22.**
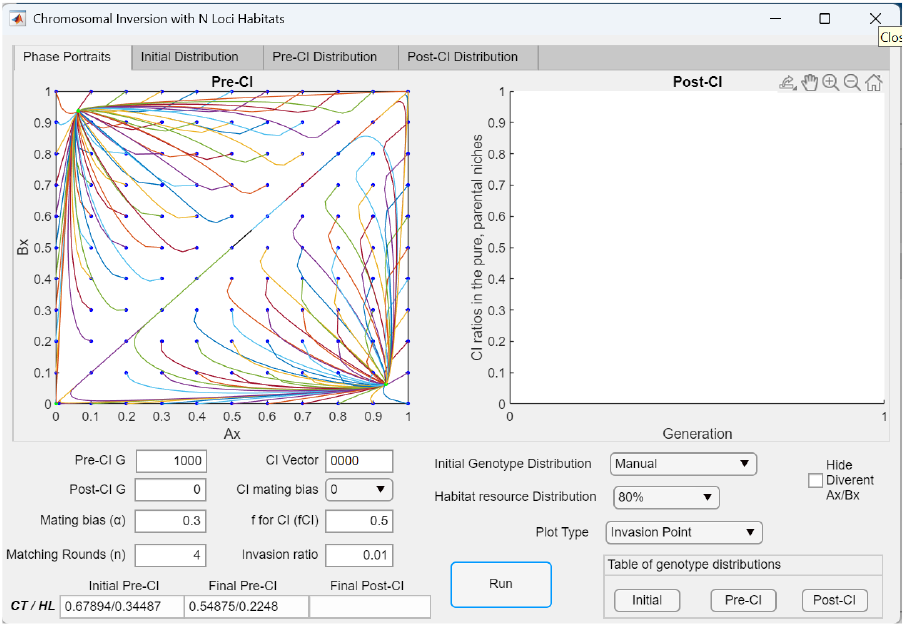
Increasing the number of ecological gene loci strengthens premating RI. As shown, increasing the number of gene loci for the ecological genotype in Fig 8 from three to four, while keeping all other parameter values unchanged during the pre-CI phase, shifts the *Ax*/*Bx* fixed point closer to the lower-right corner of the phase portrait, indicating greater assortment of mating-bias alleles between the niche ecotypes and stronger RI. The final pre-CI total inter-niche offspring ratio (*CT*) decreases from *CT* = 0.59204 in Fig 8 to *CI* = 0.54875, reflecting reduced inter-niche mating and enhanced premating RI.

#### 2 Invasion of a CI in a two-niche, two-mating-bias-allele system with multilocus ecological genotypes

We next set the number of post-CI generations (*Post*-*CI Gen*) to a value greater than zero to examine the invasion dynamics of a CI arising in niche *A* and its effects on system behavior. In a previous study, we investigated the effects of having CIs in the model without viable ecological hybrids shown in Fig 3 [39]. The current application extends that framework, enabling comparison between our earlier results and those obtained from the present model incorporating multilocus ecological genotypes and viable hybrids. We present the results in two parts: first, when the CI captures only ecological alleles, and second, when it captures both ecological and mating-bias alleles.

##### 2.1 Invasion of a CI capturing only ecological alleles

First, we examine the invasion dynamics of a CI that captures only locally adaptive ecological alleles without capturing any mating-bias allele. This is implemented by selecting “0” for *CI mating bias* in the user interface.

Fig 23a shows the *Invasion Point* plot of a CI mutant (initial *Ak* ratio = 0.01) that captures all locally adaptive ecological alleles in niche *A* in a system with three-locus ecological genotypes (*CI* = *aaa*). Because the favorable ecological alleles linked within the CI cannot be broken up by recombination, niche-*A* genotypes carrying the CI gain a fitness advantage over uninverted genotypes in the same niche, allowing the CI to invade and rise to near fixation in niche *A*. However, since *fCI* = 0.5, no hybrid offspring are produced in inter-niche heterokaryotype matings between niche-*A* ecotypes carrying the CI and niche-*B* ecotypes. Without maladaptive hybrid loss to drive or sustain the mating-bias barrier, the *Ax*/*Bx* phase portrait becomes divergent, eliminating the preexisting premating RI. This outcome is illustrated in the *Single Line* plot shown in Fig 23b, where after 200 post-CI generations the *X* allele is lost from the population (see also the post-CI genotype distribution bar graph in Fig 23c). Following the successful CI invasion, *CT* increases from 0.54875 to 0.68, reflecting increased inter-niche mating.

**Fig 23a.**
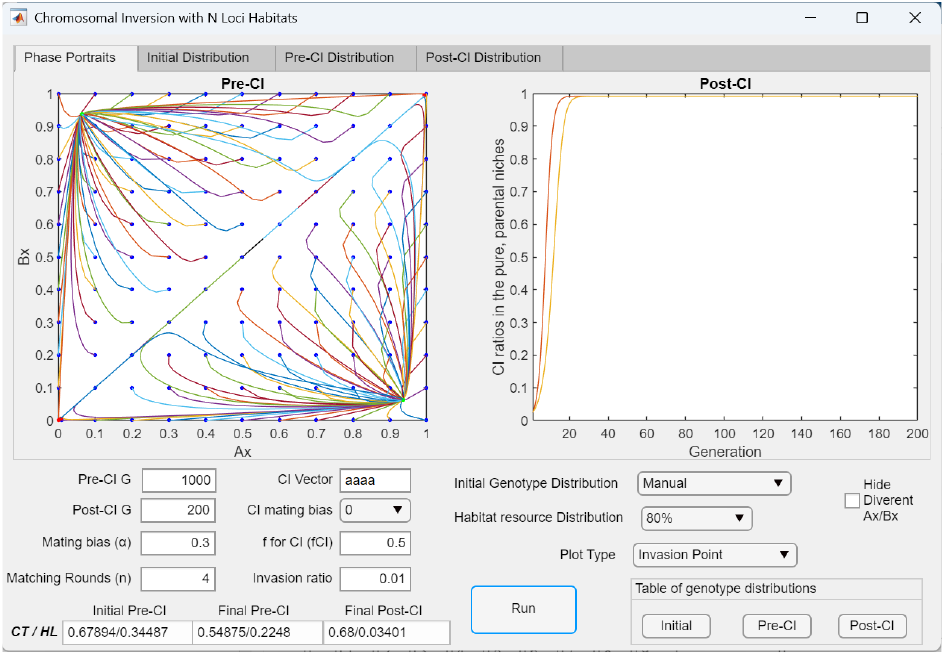
An *Invasion Point* plot showing that in a system with four-locus ecological genotypes, invasion of a CI capturing all ecological alleles—but not a mating-bias allele—causes the *Ax*/*Bx* phase portrait to become divergent and destroys pre-established premating RI. Following the pre-CI establishment of premating RI in the four-locus system shown in Fig 22, invasion of a CI mutant capturing all ecological alleles (*aaaa*), but not a mating-bias allele, removes heterokaryotype hybrid loss from the system, thereby rendering the *Ax*/*Bx* phase portrait divergent and eliminating the previously established RI.

**Fig 23b.**
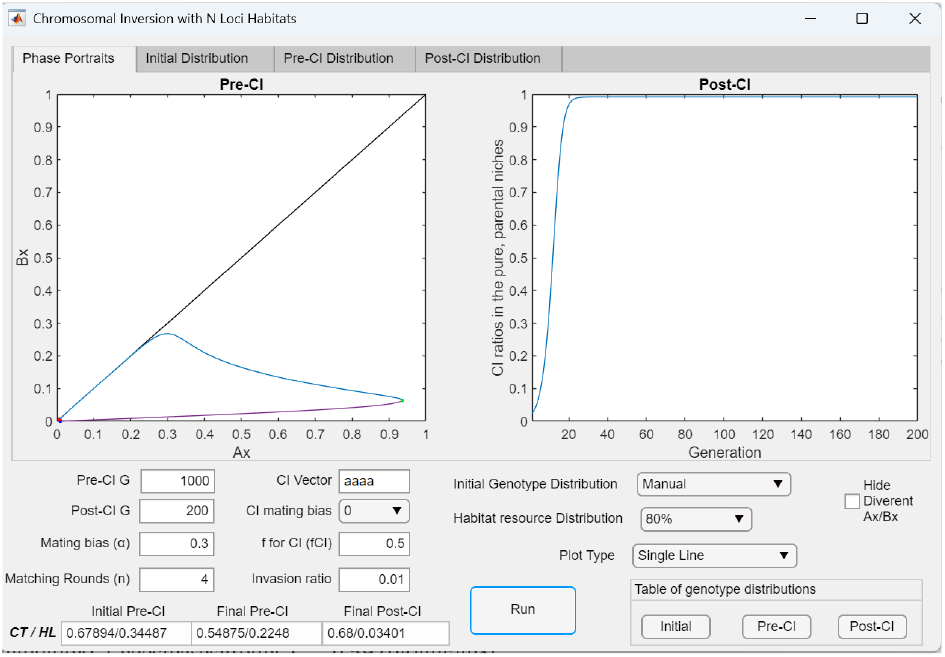
A *Single Line* plot showing that the invasion of a CI mutation capturing all ecological alleles—but not a mating-bias allele—causes a system with four-locus ecological genotypes to become divergent. A single-line plot of the phase portrait in Fig 23a shows the invasion of a CI mutant (*aaaa*) capturing all ecological alleles, but not a mating-bias allele. As the CI rises to near fixation in niche *A*, it eliminates the fixed point established in the pre-CI phase, leading to the extinction of the *X* mating-bias allele.

**Fig 23c.**
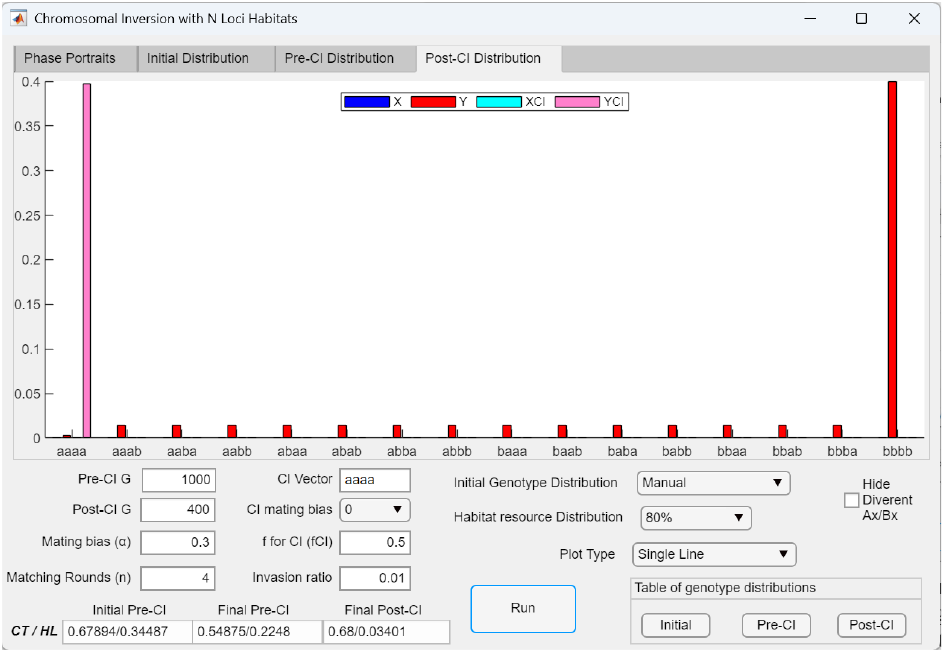
Post-CI bar graph showing that invasion of a CI mutation capturing all ecological alleles—but not a mating-bias allele—causes a system with four-locus ecological genotypes to become divergent and eliminates the *X* mating-bias allele. The post-CI bar graph shows the genotype distribution after the invasion of the CI (*aaaa*) in Fig 23b. After 400 post-CI generations, the CI rises to near fixation in niche *A*, and the *X* mating-bias allele is eliminated from the system.

When a CI captures only a subset of locally adaptive ecological alleles within a niche, its invasion fitness is reduced, yet hybrid offspring production in inter-niche mating is restored. The resulting maladaptive hybrid offspring loss reintroduces disruptive ecological selection in the system, which may allow weaker premating RI to persist. This pattern is shown in the single-line plot in Fig 24a, where the CI captures only two of the three locally adaptive ecological alleles in niche *A*. Successful invasion of the CI shifts the preexisting *Ax*/*Bx* fixed point away from the lower-right corner of the phase portrait, leading to weakened premating RI. The post-CI genotype distribution in Fig 24b shows differential assortment of the *X* and *Y* mating-bias alleles across niches. Compared to Fig 23b, the CI invasion plot in Fig 24a shows that a CI capturing a partial set of ecological alleles invades more slowly and stabilizes at a lower frequency in niche *A* than a CI capturing all ecological alleles, indicating reduced invasion fitness.

**Fig 24a.**
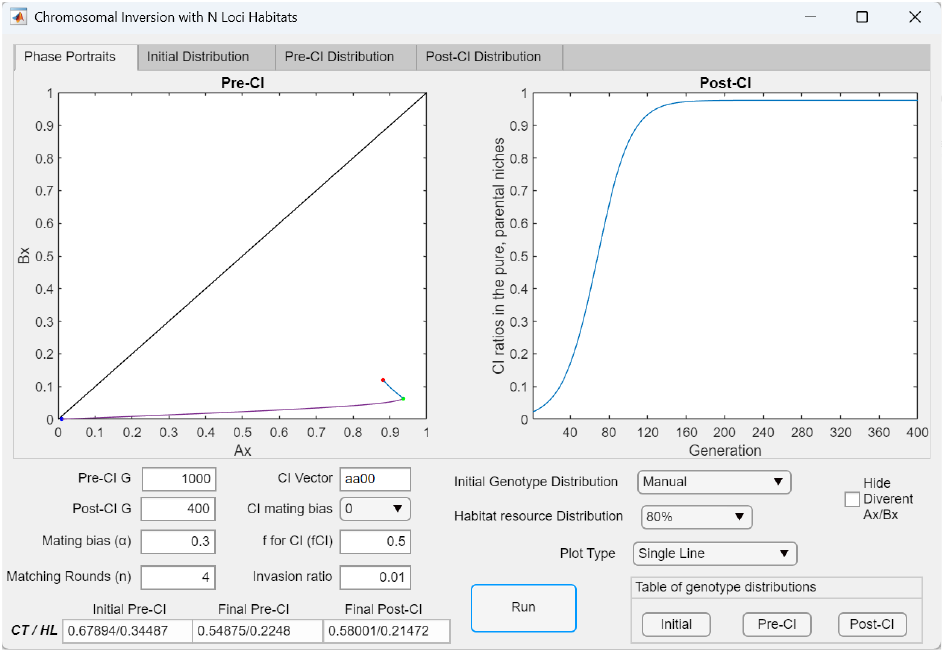
In a system with four-locus ecological genotypes, the invasion of a CI that captures only a partial set of locally adaptive ecological alleles—but not a mating-bias allele—partially restores heterokaryotype hybrid loss but still leads to weakened premating RI. In contrast to the example in Figs 23a and 23b, where the CI captures all niche-*A* alleles, the CI here captures only two of the four ecological alleles (CI = *aa*00), resulting in partial restoration of heterokaryotype hybrid loss. However, its invasion still produces a weakened premating RI, shifting the *Ax*/*Bx* fixed point away from the lower-right corner of the phase portrait (from *Ax*/*Bx* = 0.9364/0.0636 to *Ax*/*Bx* = 0.8809/0.1192). Consequently, *CT* increases from 0.54875 to 0.5800 after the post-CI phase of the simulation, indicating increased inter-niche mating. Because this CI captures only a subset of adaptive ecological alleles, its invasion fitness is reduced relative to that of a CI capturing all ecological alleles. This is reflected in the invasion plot, where the CI invades more slowly and ultimately stabilizes at a lower frequency ratio in niche *A* than in the invasion plots shown in Figs 23a and 23b.

**Fig 24b.**
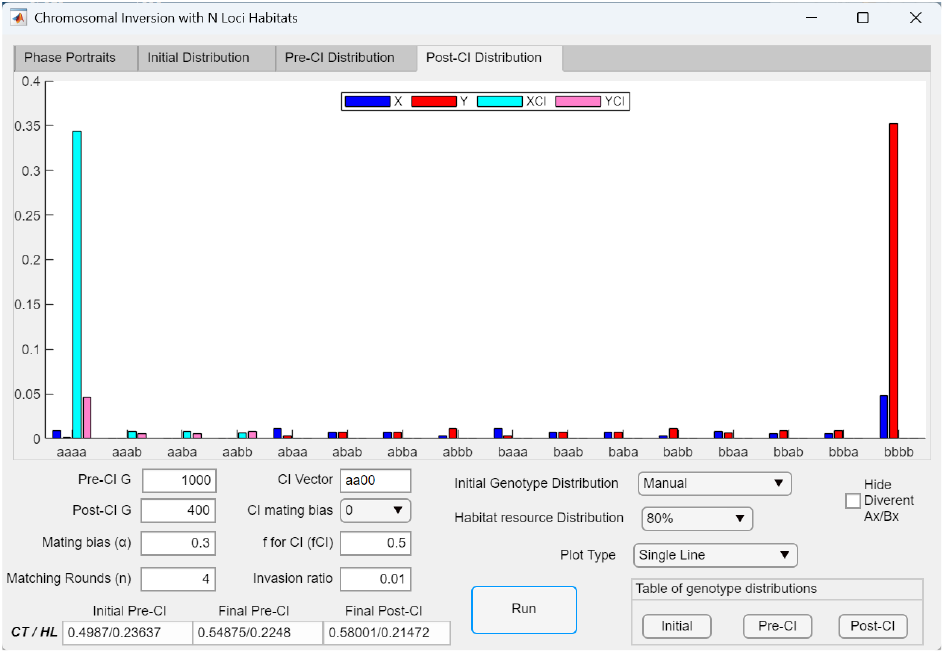
Post-CI genotype distribution following the invasion of a CI (aa00) that captures only a partial set of locally adaptive ecological alleles and no mating-bias allele. The post-CI bar graph shows the genotype distribution after the invasion of the CI (*aa*00) in Fig 24a. After 400 post-CI generations, differential assortment of the *X* and *Y* mating-bias alleles between niches *A* and *B* is re-established, resulting in weaker premating RI.

In Fig 25, the CI captures only one of the three locally adaptive ecological alleles in niche *A* (CI = *a*00). A CI capturing a single ecological allele cannot affect the effective value of *f* in inter-niche mating or alter the system’s overall dynamics. The fitness advantage of a CI mutant arises from its ability to preserve a favorable combination of adaptive alleles that would otherwise be broken up by recombination during inter-niche mating—a condition that requires capturing multiple adaptive alleles. Because the CI in Fig 25 captures only one adaptive allele, it provides no additional fitness advantage to genotypes carrying the inversion relative to their same-niche counterparts lacking it and therefore cannot invade. As a result, the preexisting *Ax*/*Bx* fixed point and premating RI remain unchanged between the final pre-CI and post-CI stages.

**Fig 25.**
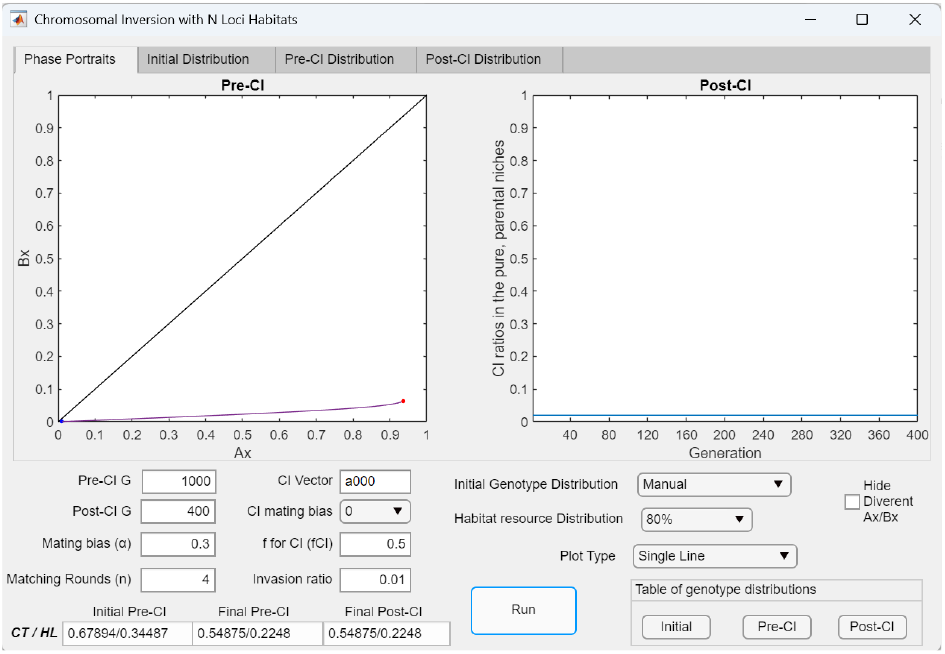
In a system with four-locus ecological genotypes, a CI capturing only one adaptive ecological allele—but not a mating-bias allele—has no effect on hybrid loss or pre-established RI. In the Fig 24a example, if the CI instead captures only a single locally adaptive allele in niche *A* (CI = *a*000) without capturing a mating-bias allele, it has no effect on the existing hybrid loss or the established premating RI. The CI has no fitness advantage over same-niche individuals lacking the inversion and therefore cannot invade.

##### 2.2 Invasion of a CI capturing both ecological alleles and a mating-bias allele

Next, our study examined the invasion of a CI mutant that captures locally adaptive ecological alleles together with a mating-bias allele. We first analyzed the case in which the CI captures all of the locally adaptive alleles, followed by the case in which it captures only a partial subset of these ecological alleles.

The invasion dynamics of a CI that captures all locally adaptive ecological alleles and a mating-bias allele depend on the relative sizes of niches *A* and *B*. Fig 26 shows the post-CI invasion dynamics of a CI mutant that captures all locally adaptive ecological alleles together with the prevalent mating-bias allele in niche *A* (CI = *aaaX*) in a system with three ecological loci. Under the *80% Habitat Resource Distribution* setting (*NA*: *Hybrids*: *NB* = 0.4: 0.2: 0.4), the invasion of the CI mutant destroys the pre-established premating RI from the pre-CI phase. As a result, the *Ax*/*Bx* phase portrait becomes divergent, and the *Y* mating-bias allele is eliminated.

**Fig 26.**
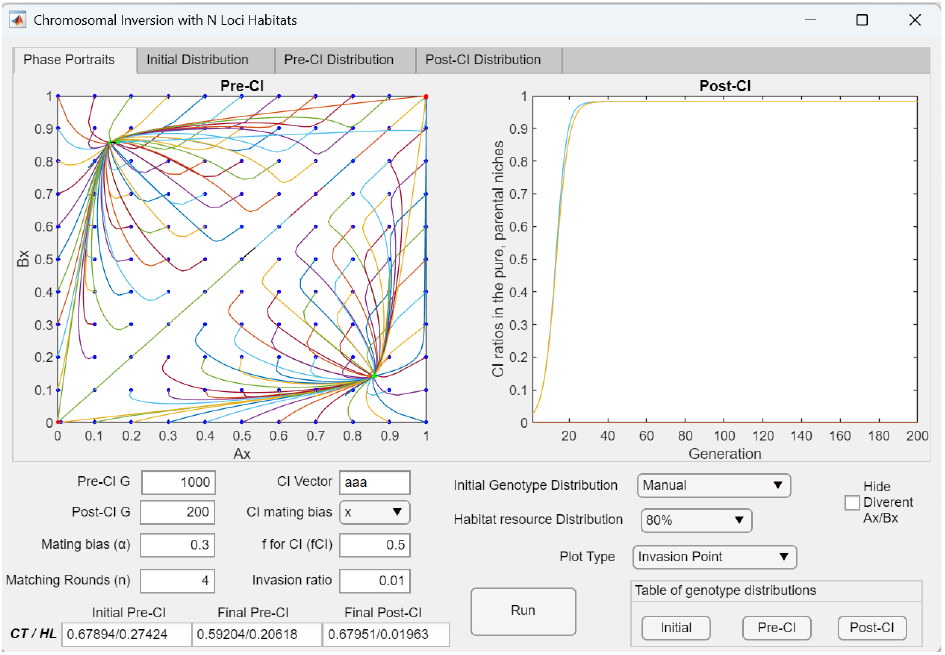
In a system with three-locus ecological genotypes, invasion of a CI capturing all adaptive ecological alleles and the prevalent mating-bias allele in niche *A* nonetheless causes the system to become divergent. In this example, the CI captures all locally adaptive ecological alleles and the dominant *X* mating-bias allele in niche *A* (CI = *aaaX*). The CI invades successfully and rises to near fixation in niche *A*. However, because no hybrid offspring are produced in inter-niche heterokaryotype matings between niche-*A* ecotypes carrying the CI and niche-*B* ecotypes, the effective value of *f* = 0.5, and there is no maladaptive hybrid loss to drive or maintain the *Ax*/*Bx* mating-bias barrier. Consequently, the *Ax*/*Bx* phase portrait becomes divergent, and the *Y* mating-bias allele is eliminated, while the *X* allele cannot be eliminated because it is linked with the niche-*A* ecotype through the CI.

the same outcome is observed when niche *A* is larger than niche *B*. As illustrated in Fig 28, when the habitat resource distribution shown in Fig 27 is modified so that *NA* > *NB* (*NA*: *Hybrids*: *NB* = 0.6: 0.2: 0.2), the invasion of the CI mutant (*CI* = *aaaX*) likewise destroys the pre-established premating RI and renders the *Ax*/*Bx* phase portrait divergent.

**Fig 27.**
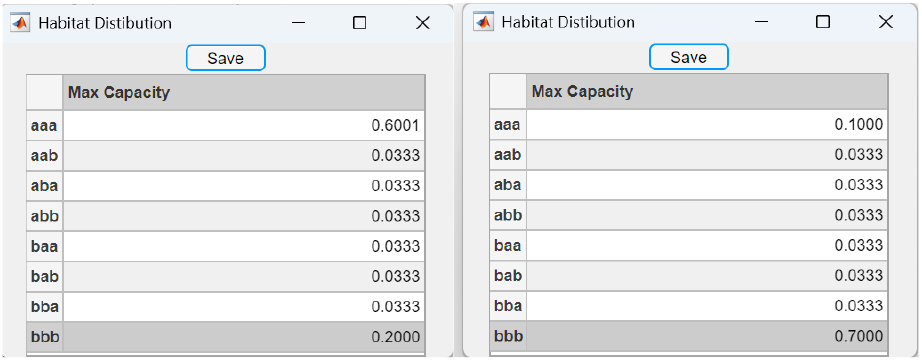
Two Habitat Distribution Tables specifying unequal carrying capacities for niche *A* and niche *B* in a system with three-locus ecological genotypes. The table on the left shows an example where *NA* > *NB* (0.6 versus 0.2), and the table on the right shows the opposite case where *NB* > *NA* (0.7 versus 0.1). In both cases, the hybrid habitat distribution corresponds to the 80% option in *Habitat Resource Distribution*, where all hybrid niche resources sum to 20%.

**Fig 28.**
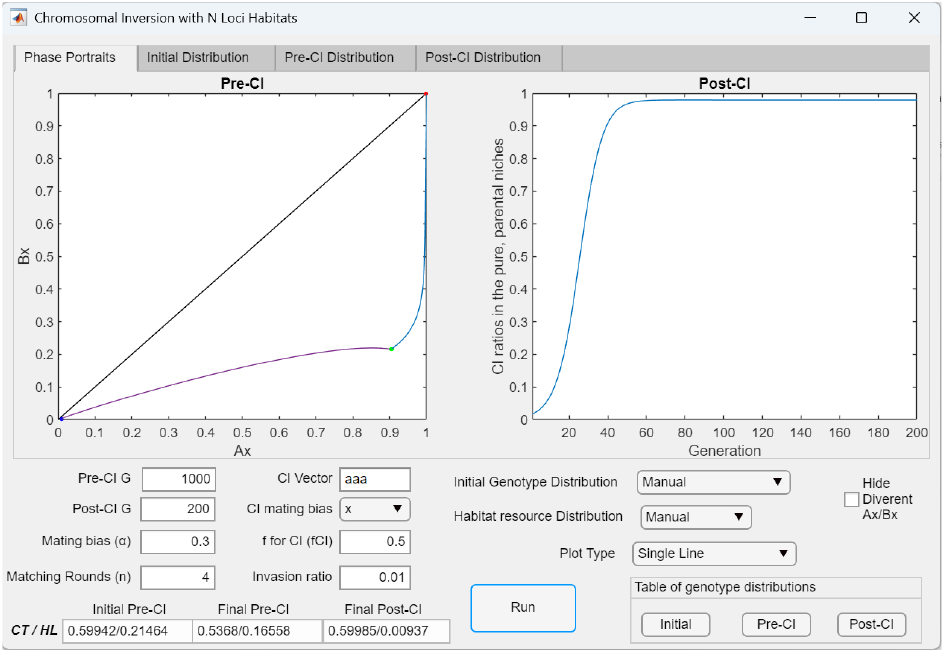
In a system with three-locus ecological genotypes, when *NA* > *NB*, invasion of a CI capturing all adaptive ecological alleles and the prevalent mating-bias allele *X* in niche *A* causes the system to become divergent. Here, the Habitat Resource Distribution in Fig 26 (the 80% option with equal *NA*: *NB* ratios of 0.4:0.4) is replaced by the distribution shown in the left table of Fig 27, where *NA* > *NB* (0.6: 0.2). The resulting single-line plot shows that, following the invasion of the CI (CI = *aaaX*), the *Ax*/*Bx* fixed point is eliminated, and the *Y* mating-bias allele is lost after the post-CI phase of the simulation.

In contrast, when niche *B* is larger than niche *A*, the outcome is different, and the *Ax*/*Bx* phase portrait remains convergent. As shown in Fig 29a, when *NB* > *NA* (*NA*: *Hybrids*: *NB* = 0.1: 0.2: 0.7), the CI mutant capturing all ecological alleles (*CI* = *aaaX*) is able to invade and shift the pre-CI *Ax*/*Bx* fixed point toward the lower-right corner of the phase portrait. The post-CI genotype distribution in Fig 29b shows that the *X* allele is able to hitchhike with the ecologically adaptive CI and rise to near fixation in niche *A*. However, the CI invasion also increases the *X* allele ratio in niche *B*, resulting in little net change in premating RI, as reflected by nearly unchanged pre- and post-CI *CT* values. The post-CI *HL* decreases because fewer hybrid offspring are produced through inter-niche heterokaryotype matings. Nonetheless, when *α* exceeds 3.1 in Fig 29a, the weakened mating bias leads to more inter-niche mating, which then gives the *X* allele in niche *B* a sufficient fitness advantage to eliminate the *Y* allele and causes the system to become divergent.

**Fig 29a.**
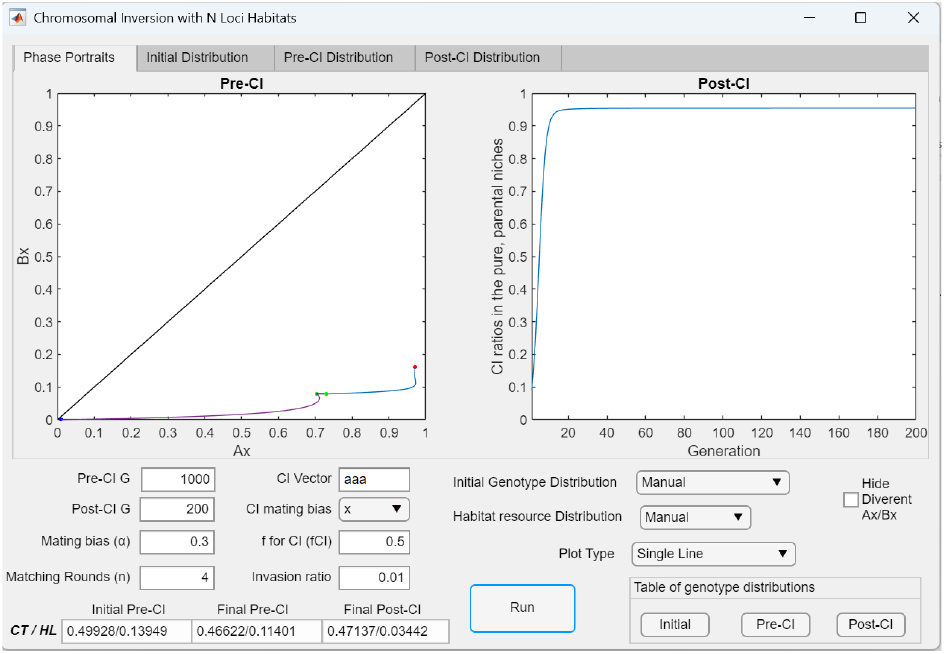
In a system with three-locus ecological genotypes, when *NA* < *NB*, invasion of a CI capturing all adaptive ecological alleles and the prevalent mating-bias allele *X* in niche *A* maintains system convergence. The Habitat Resource Distribution in Fig 26 (the 80% option with equal *NA*: *NB* ratios of 0.4:0.4) is replaced by the distribution shown in the right table of Fig 27, where *NA* < *NB* (0.1:0.7). The resulting single-line plot shows that, following the invasion of the CI (CI = *aaaX*), the *Ax*/*Bx* fixed point is not eliminated but shifts closer to the lower-right corner of the phase portrait.

**Fig 29b.**
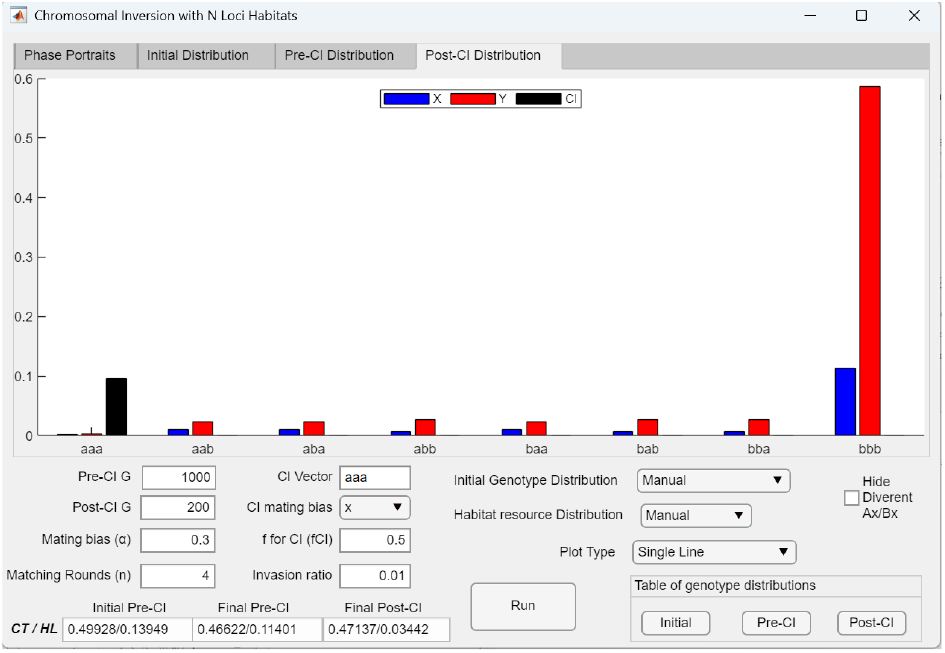
Bar graph showing the post-CI genotype distribution after the invasion of a CI (CI = *aaaX*) when *NA* < *NB* in a system with three-locus ecological genotypes. The post-CI genotype distribution corresponding to Fig 29a shows that the more numerous *Y* alleles in niche *B* cannot eliminate the *X* allele in niche *A* because *X* is linked to the niche-*A* ecotype within the CI and cannot be removed from niche *A* through recombination. Consequently, the *X* allele is able to hitchhike with the ecologically adaptive alleles in the CI and rise to near fixation in niche *A*. However, the CI invasion also increases the *X* allele ratio in niche *B*, resulting in little net change in premating RI, with *CT* remaining nearly identical after the invasion. Post-CI *HL* decreases because fewer hybrid offspring are produced in inter-niche heterokaryotype matings.

When the CI captures only a partial subset of locally adaptive ecological alleles together with the prevalent mating-bias allele, the relative sizes of niches *A* and *B* do not appear to influence the outcome. In such cases, the system remains convergent following CI invasion. This is demonstrated in the examples shown in Figs 30a and 30b. In both scenarios (*NA* > *NB* and *NB* > *NA*), a CI mutant capturing two of the three locally adaptive ecological alleles together with the prevalent mating-bias allele (*CI* = *aa*0*X*) is able to invade the pre-established *Ax*/*Bx* barrier system, shift the pre-CI fixed point toward the lower-right corner of the *Ax*/*Bx* phase portrait, and strengthen the existing premating RI.

**Fig 30a.**
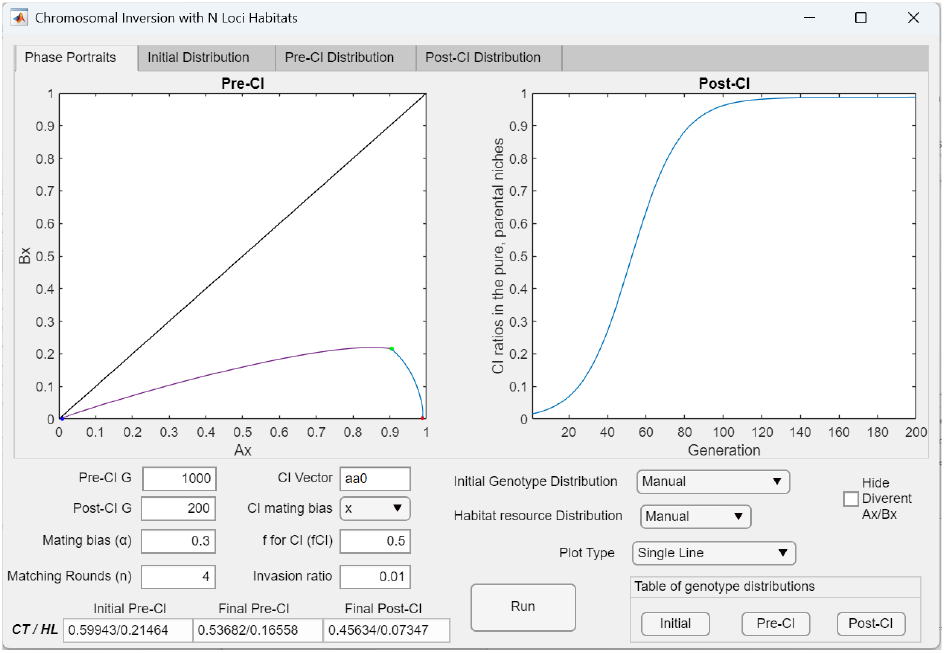
In a system with three-locus ecological genotypes, when *NA* > *NB*, the invasion of a CI capturing only a partial set of adaptive ecological alleles and the prevalent mating-bias allele *X* in niche *A* strengthens RI. If the CI in Fig 28 captures only a subset of the adaptive ecological alleles in niche *A* (CI = *aa*0*X*), hybrid offspring are again produced from inter-niche matings, and disruptive ecological selection is restored. The resulting maladaptive hybrid loss drives the evolution of mating-bias barriers to minimize this loss. Because the high-mating-bias *X* allele can hitchhike with adaptive ecological alleles within the CI, it gains an augmented fitness advantage to invade and rises to near fixation in niche *A*, thereby enhancing premating RI between the niche ecotypes. Following CI invasion, both *CT* and *HL* decrease.

**Fig 30b.**
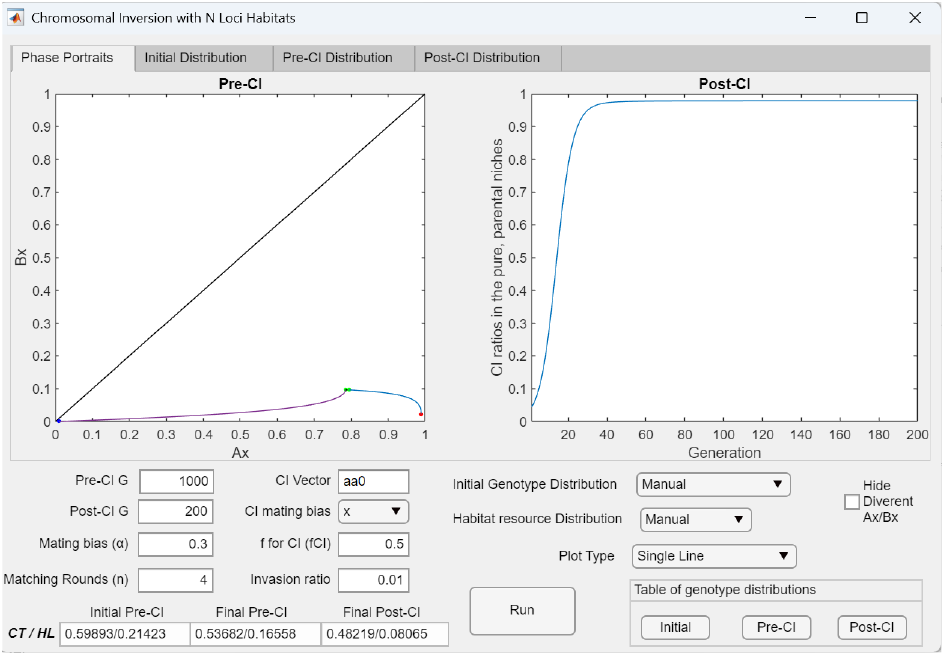
In a system with three-locus ecological genotypes, when *NA* < *NB*, the invasion of a CI capturing a partial set of adaptive ecological alleles and the prevalent mating-bias allele *X* in niche *A* strengthens RI. When the habitat *NA*: *NB* ratios in Fig 30a are reversed (i.e., *NA*: *NB* = 0.2: 0.6) so that *NA* < *NB*, with the hybrid distribution unchanged, successful invasion of the CI (CI = *aa*0*X*) restores hybrid offspring production and disruptive ecological selection, leading to stronger premating RI through reinforcement. As shown, the CI invasion shifts the *Ax*/*Bx* fixed point closer to the lower-right corner of the phase portrait, resulting in stronger premating RI. Both *CT* and *Hr* decrease after the CI invasion.

In Figs 31a and 31b, the CI mutant captures only one of the three locally adaptive ecological alleles together with the prevalent mating-bias allele (*CI* = *a*00*X*) in a pre-established *Ax*/*Bx* system with a fixed-point polymorphism. Although capturing only a single adaptive ecological allele does not by itself provide a fitness advantage over noninverted same-niche counterparts (see Fig 25), linkage with the locally prevalent mating-bias allele (*X*) anchors the mating-bias allele in niche A. This coupling generates a synergistic invasion fitness advantage, enabling the CI to invade and reach fixation in niche *A*, thereby strengthening the pre-existing premating RI.

**Fig 31a.**
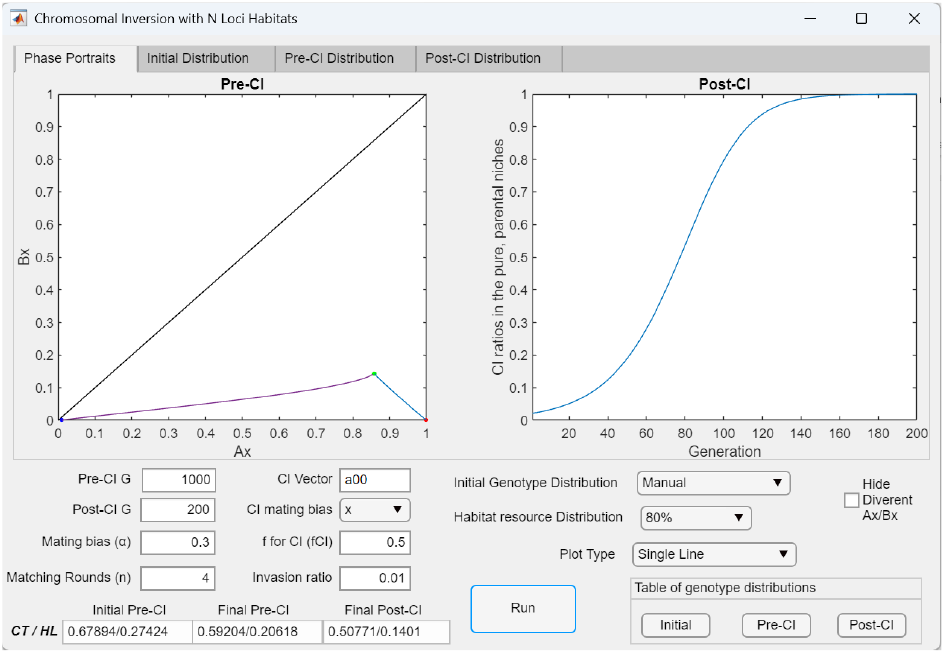
In a system with three-locus ecological genotypes, the invasion of a CI capturing a single adaptive ecological allele and the prevalent mating-bias allele *X* in niche *A* strengthens RI. A CI capturing only one ecological allele does not reduce hybrid production or affect disruptive ecological selection. Therefore, it gains no fitness advantage in inter-niche mating and has the same fitness as its same-niche counterparts without the CI. However, when the CI also captures an adaptive mating-bias allele along with the single ecological allele, their linkage association can generate a strong synergistic advantage. As shown in the *Invasion Point* plot, in a system with three loci for ecological genotypes, such a CI (CI = *a*00*X*) can readily invade and rise to fixation with its linked *X* allele in niche *A*, shift the *Ax*/*Bx* fixed point to the lower-right corner of the phase portrait, and produce strong premating RI between the niche ecotypes.

**Fig 31b.**
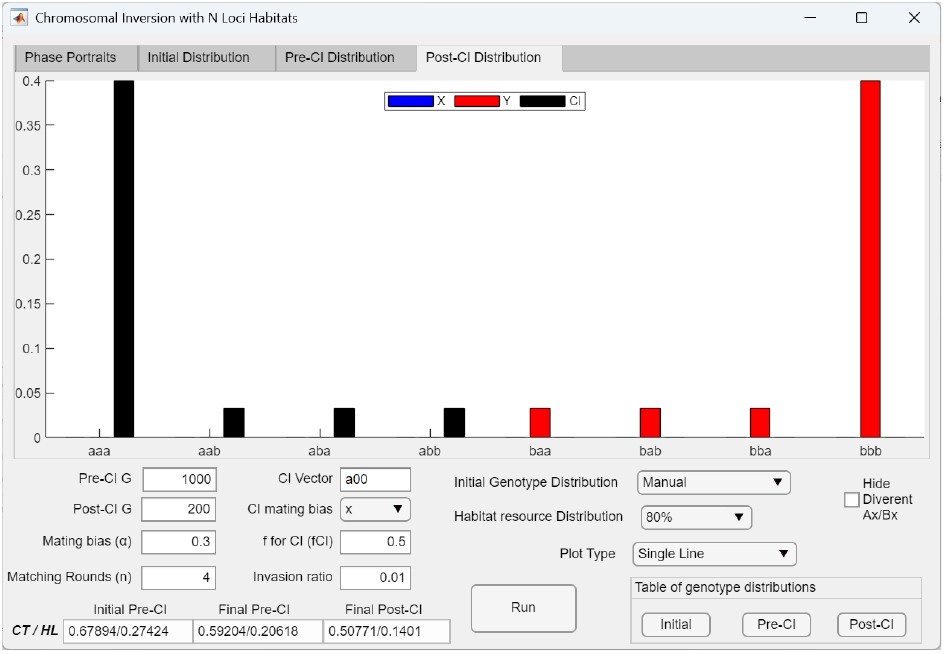
Bar graph showing the post-CI genotype distribution after the invasion of a CI (CI = *a*00*X*) in a system with three-locus ecological genotypes. The bar graph displays the post-CI genotype distribution from Fig 31a. It shows that the CI (CI = *a*00*X*) has successfully invaded and become fixed in all genotypes carrying the captured *a* allele.

**Fig 31c.**
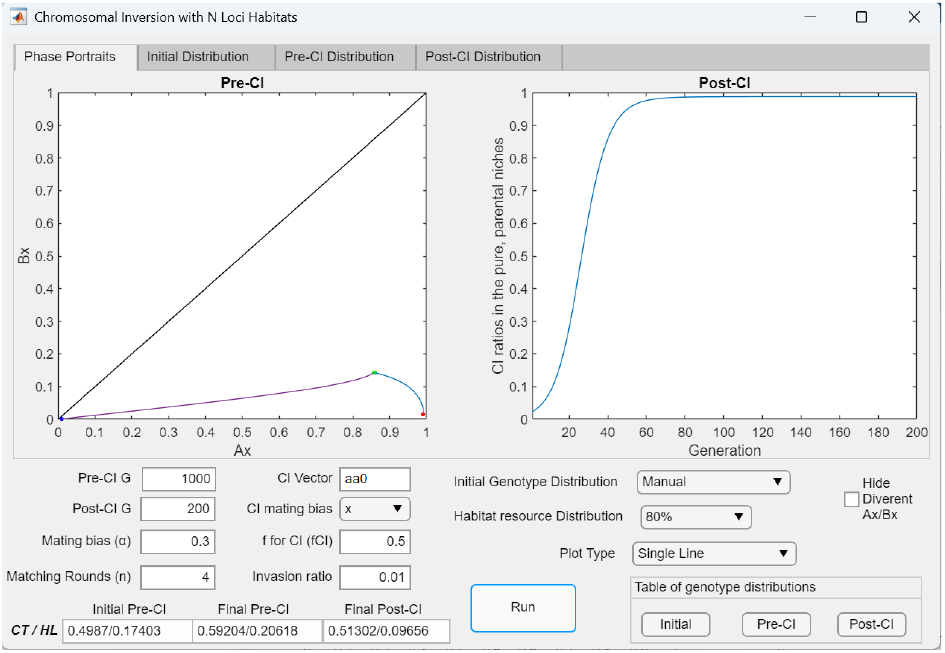
In a system with three-locus ecological genotypes, a CI capturing two of the three locally adaptive ecological alleles together with the prevalent mating-bias allele *X* in niche *A* (*CI* = *aa*0*X*) generates weaker premating RI and reduced hybrid loss compared to a CI capturing only a single ecological allele. Using the same parametric values as in Fig 31a, where the CI captures only one adaptive ecological allele (*CI* = *a*00*X*), invasion of the CI mutant capturing two of three ecological alleles (*CI* = *aa*0*X*) produces weaker premating RI, as indicated by a higher post-CI *CT* value (0.51302 versus 0.50771), because the CI does not rise to fixation. Conversely, post-CI hybrid loss is lower (*HL* = 0.09656 versus 0.1401), as capturing more adaptive ecological alleles results in fewer hybrid offspring from inter-niche matings.

**Fig 31d.**
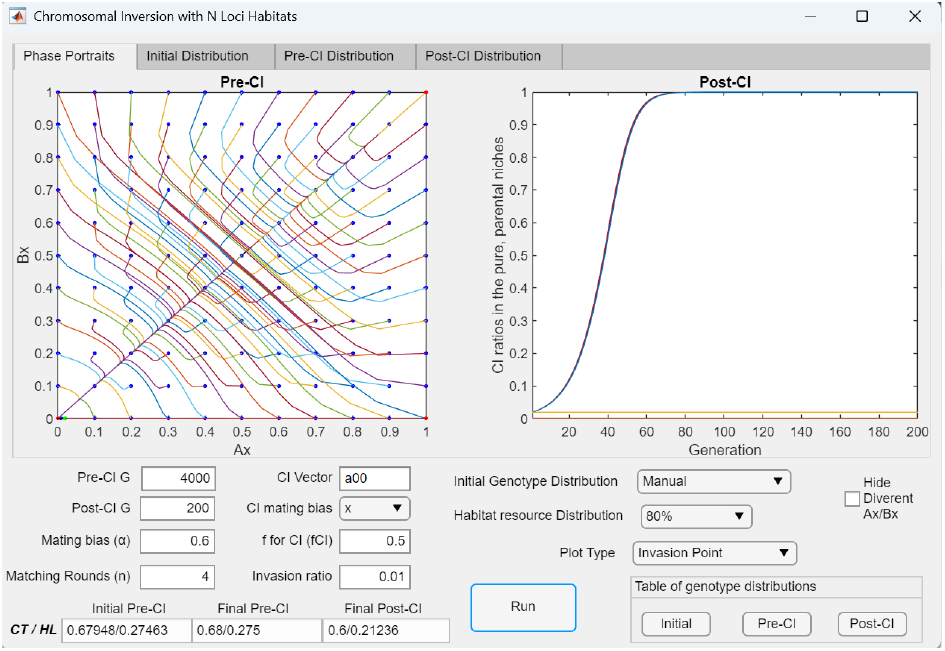
In a system with three-locus ecological genotypes, the invasion of a CI capturing a single adaptive ecological allele together with a high-mating-bias allele in niche *A* can generate fixed-point polymorphism and premating RI even when the *Ax*/*Bx* phase portrait is divergent. In this *Invasion Point* plot, increasing the value of *α* in Fig 31a from 0.3 to 0.6, while keeping all other relevant parameters constant, causes the pre-CI *Ax*/*Bx* phase portrait to become divergent, eliminating fixed points. As a result, in the pre-CI phase, an *X* mating-bias mutant arising in niche *A* (*Ax* = 0.01) with *α* = 0.6 exhibits weakened mating bias and cannot invade. Nonetheless, during the post-CI phase, a mutant CI that captures both the *X* allele and a locally adaptive ecological allele (CI = *a*00*X, Ak* = 0.01) can invade despite the weak mating bias of the *X* allele. As the CI rises to fixation in niche *A*, it carries the *X* allele with it. In the post-CI *Ax*/*Bx* phase portrait, this dynamic is reflected by the *AN* mutant population invading from near the origin and becoming fixed at the lower-right corner, thereby producing premating RI. In our simulations, such a CI can invade at an *α* value as high as 0.999.

In our simulations, the CI mutant (*CI* = *a*00*X*) can invade even when *α* is as high as 0.999 to establish an *AN*/*BN* fixed point at *Ax*/*Bx* = 1/0 and generate premating RI. Thus, by varying *α* between 0 and 1 (0 < *α* < 1), such a CI—which captures one ecological allele together with a high mating-bias allele—can always invade to produce any level of premating RI from 0% to 100% between niche ecotypes. This result appears to hold in systems with any number of ecological gene loci greater than one.

As demonstrated in Fig 31c, compared to systems in which the CI captures more than one adaptive ecological allele, a CI capturing only a single adaptive allele does not diminish disruptive ecological selection. Consequently, it more effectively drives the linked mating-bias allele toward fixation and tends to generate stronger premating RI, despite producing a higher proportion of hybrid offspring due to capturing fewer adaptive ecological alleles.

The high invasion fitness of a CI that couples a single locally adaptive ecological allele with a mating-bias allele may enable it to invade a divergent *Ax*/*Bx* system and generate premating RI. This is demonstrated by the example in Fig 31d. On their own, a CI capturing only one locally adaptive ecological allele (*CI* = *a*00) and a weak mating-bias allele X (*α* = 0.6) cannot invade a divergent *Ax*/*Bx* phase portrait. However, when these alleles are linked within a CI (*CI* = *a*00*X*), the CI can function as a magic trait that acquires synergistic fitness advantage over its same-niche ecotypes lacking the inversion. Consequently, the CI is able to invade successfully, leading to fixed-point polymorphism in the *Ax*/*Bx* phase portrait and premating RI.

In Fig 32, a CI mutant with magic-trait properties that captures more than one locally adaptive ecological allele (*CI* = *aa*0*X*) exhibits greater invasion fitness than a CI that captures only a single adaptive allele (*CI* = *a*00*X*). This is reflected by the shorter generation time (*T*_95_) required for the CI to reach its steady-state frequency. Capturing a larger number of locally adaptive ecological alleles provides the CI with a greater fitness advantage over noninverted same-niche ecotypes, thereby accelerating its invasion to near fixation in niche *A*.

**Fig 32.**
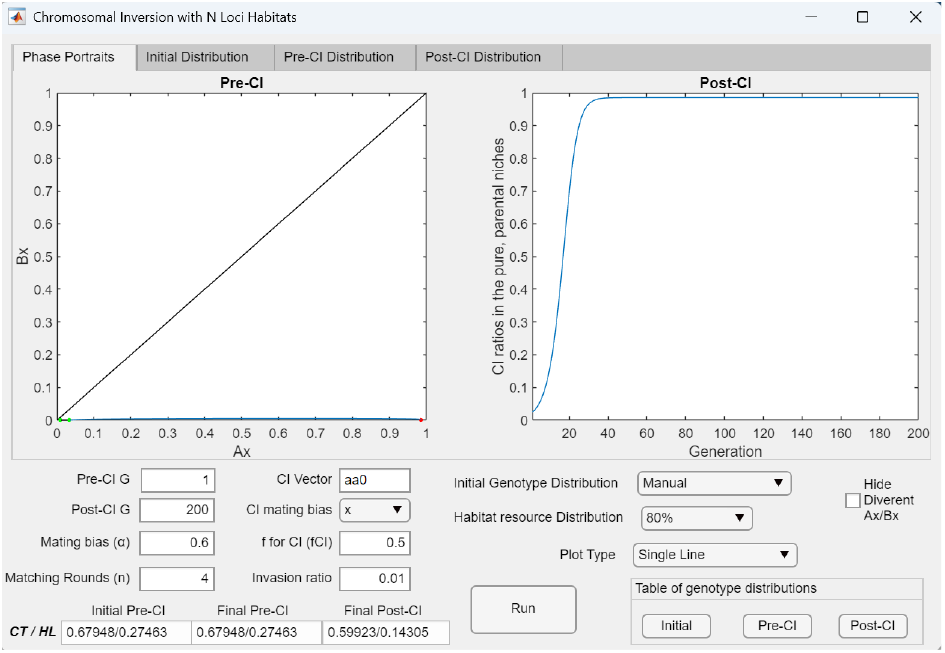
In a system with three-locus ecological genotypes, a CI mutant capturing two adaptive ecological alleles together with a high-mating-bias allele in niche *A* forms a magic trait with synergistically increased invasion fitness, even when the mating bias is weak. With all other parameters held constant, the pre-CI generation in Fig 31d is set to one, and the CI vector is specified as *aa*0*X* to simulate the invasion dynamics of a CI mutant capturing a partial set (2 of 3) of adaptive ecological alleles and the *X* mating-bias allele in a sympatric population carrying only the *Y* allele. This CI mutant acts as a magic trait, exhibiting a synergistic fitness advantage that enables invasion despite the weak mating bias (*α* = 0.6). Without linkage to the adaptive ecological alleles, the *X* allele alone cannot invade; however, when captured within the CI, it can hitchhike to near fixation in niche *A*. The time (*T*_95_) for the mutant CI (*CI* = *aa*0*X*) to reach 95% of its steady-state frequency is 28 generations— shorter than the *T*_95_ observed when the CI captures only one ecological allele (60 generations for *CI* = *a*00*X* in Fig 31d)—indicating that capturing more adaptive ecological alleles confers greater invasion fitness.

As illustrated by the example in Fig 31c, a CI that captures more locally adaptive alleles acquires higher invasion fitness; however, it also tends to produce weaker RI (reflected in higher *CT*). This general pattern is likewise confirmed in our simulations of a system with four ecological gene loci: a CI capturing more locally adaptive alleles together with the locally favored mating-bias allele invades more rapidly due to its enhanced invasion fitness, yet tends to reach a lower equilibrium frequency and to produce weaker RI (higher *CT*) along with reduced residual hybrid loss (lower *HL*).

Conversely, if the CI captures a locally maladaptive ecological allele, its fitness is reduced relative to same-niche ecotypes without the inversion. This outcome is shown in Fig 33, where a CI arising in niche *A* but capturing a maladaptive allele originating from niche *B* (*CI* = *a*0*bX*) exhibits lower fitness than its noninverted counterparts in niche *A* and cannot invade.

**Fig 33.**
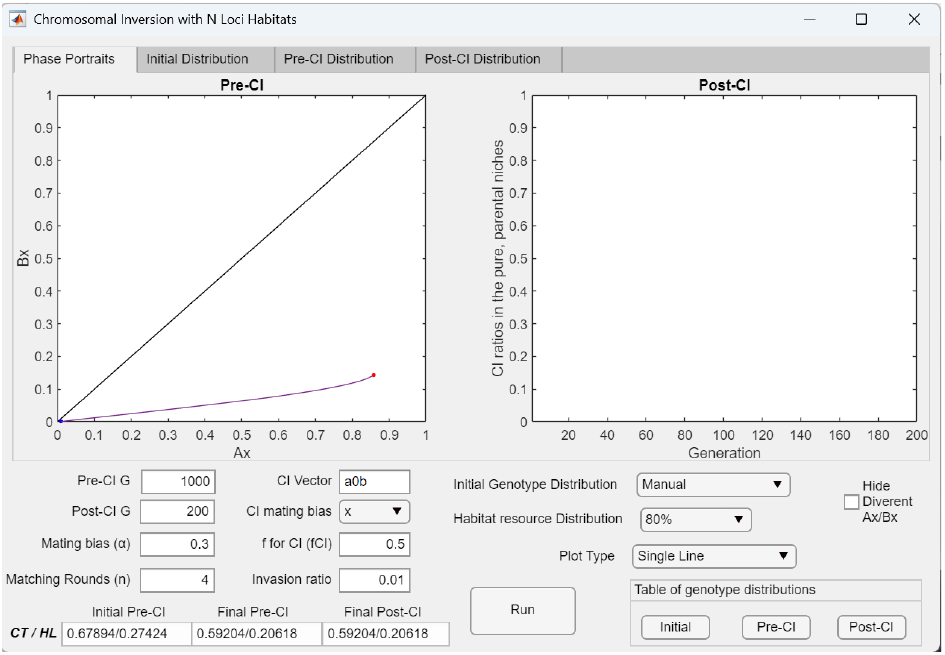
In a system with three-locus ecological genotypes, the invasion fitness of a CI decreases when it captures locally maladaptive ecological alleles. If the CI in Fig 31a (CI = *a*00*X*) captures a locally maladaptive allele (*b*) in niche *A* (CI = *a*0*bX*), its fitness is reduced when nonrandom assortment of ecological and mating-bias alleles has been established across niches after the pre-CI phase. As a result, the CI has lower fitness than its same-niche counterparts in niche *A* and cannot invade.

With 40% hybrid habitat resources (Fig 34b), the results shown in Figs 34a and 34c demonstrate that a mutant CI capturing a partial set of locally adaptive ecological alleles together with the locally prevalent mating-bias allele (*CI* = *aa*0*X*) has greater invasion fitness than a CI capturing the same adaptive ecological alleles without the mating-bias allele (*CI* = *aa*0). This difference is reflected by the shorter invasion time (*T*_95_) and the final steady-state frequency of the CI capturing both allele types. These outcomes imply the existence of positive selection favoring the linkage of locally adaptive ecological and mating-bias alleles within a low-recombination region of the genome.

**Fig 34a.**
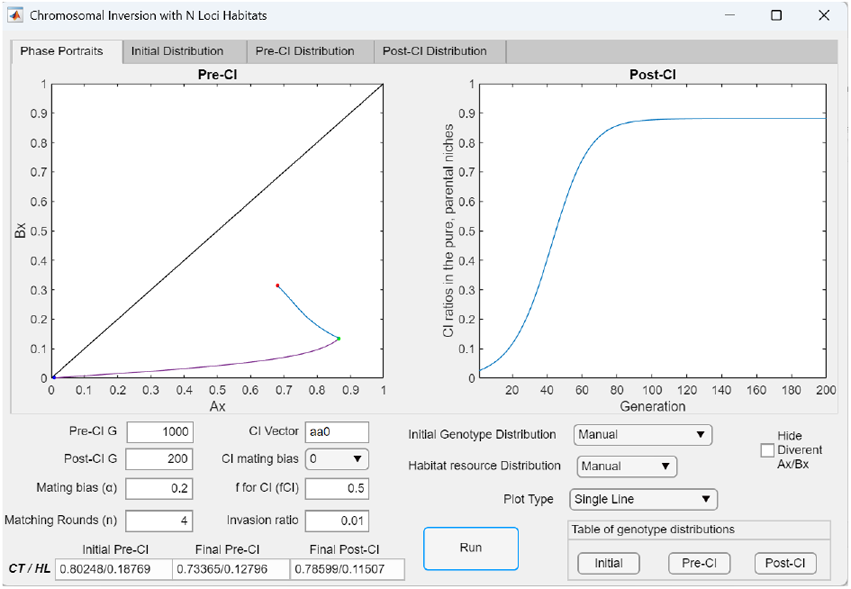
In a system with three-locus ecological genotypes and 40% hybrid habitat resources, the invasion fitness and final steady-state frequency of a CI capturing only locally adaptive ecological alleles depend on the number of adaptive alleles it captures. Using the Initial Genotype Distribution and Habitat Resource Distribution tables shown in Fig 34b, the *Invasion Point* plot shows that in the post-CI phase, an invading CI mutant (CI = *aa*0) requires 74 generations to reach 95% of its final steady-state ratio (0.8812) in a polymorphism with uninverted ecotypes in niche *A*.

**Fig 34b.**
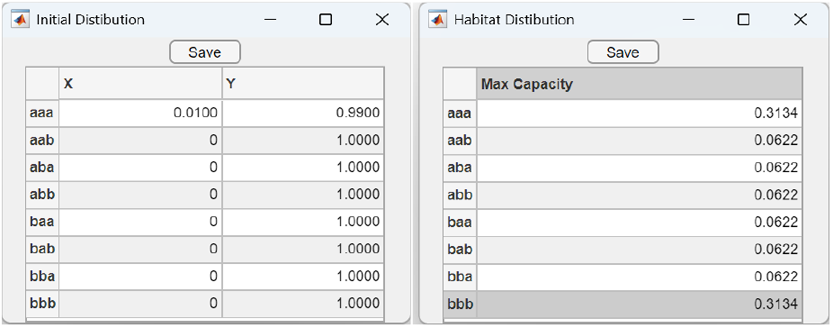
The Initial Genotype Distribution Table and Habitat Resource Distribution Table used in Fig 34a and Figs 34c–34e. The Initial Genotype Distribution Table (left) shows the initial population ratio of a mutant *X* allele arising in niche *A* (*Ax* = 0.01) within a sympatric population otherwise carrying only the *Y* mating-bias allele. The Habitat Resource Distribution Table (right) shows that 60% of the habitat resources are allocated equally between niches *A* and *B*, while the remaining 40% are distributed equally among hybrid niches.

**Fig 34c.**
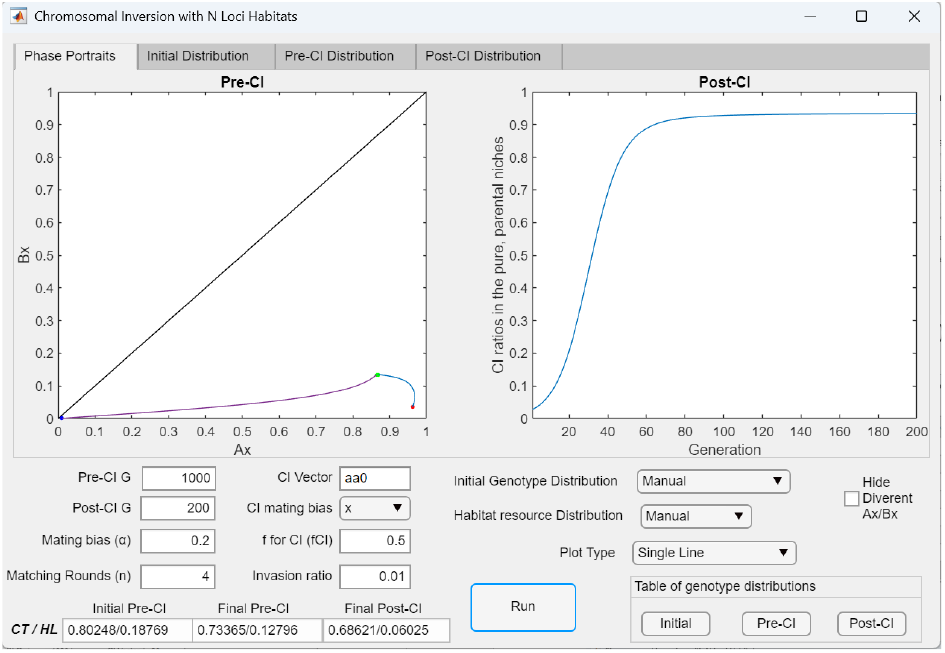
In a system with three-locus ecological genotypes and 40% hybrid habitat resources, a CI capturing both locally adaptive ecological alleles and the prevalent mating-bias allele exhibits synergistically enhanced invasion fitness, allowing it to reach a higher frequency more rapidly in a polymorphism with uninverted same-niche ecotypes. When the CI in Fig 34a also captures the locally prevalent mating-bias allele *X* (CI = *aa*0*X*), while all other parameters remain unchanged, the linkage between favorable ecological and mating-bias traits causes the CI to behave like a magic trait with augmented invasion fitness. In the *Invasion Point* plot, this is reflected by the shorter time required to reach 95% of its steady-state frequency (60 generations) and the higher equilibrium ratio it attains (0.9331) in polymorphism with uninverted same-niche ecotypes. The invasion of the CI also produces greater reductions in the post-CI *CT* and *HL* values, indicating stronger reproductive isolation between the niche-*A* and niche-*B* ecotypes.

**Fig 34d.**
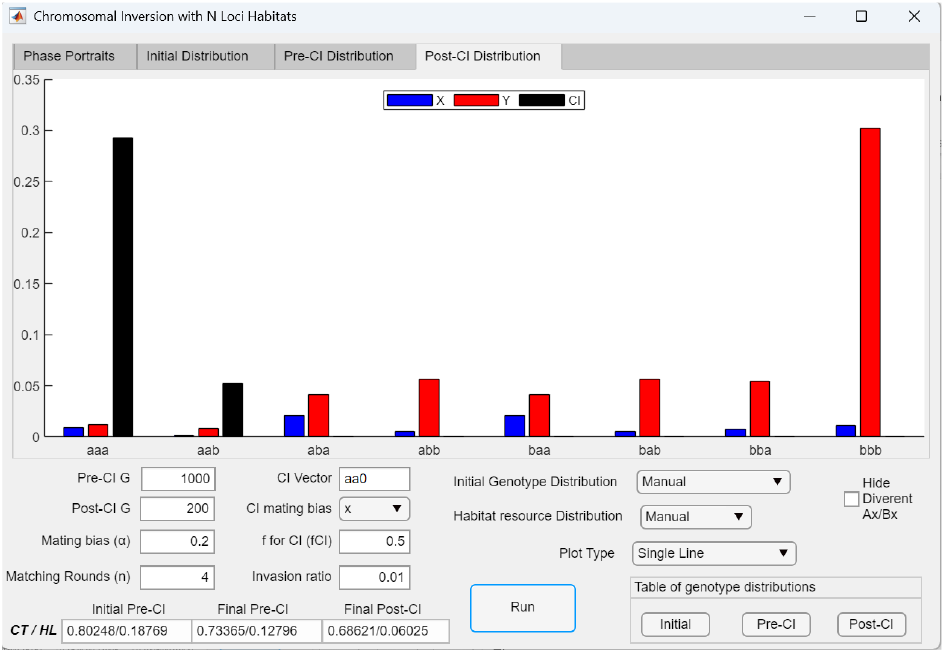
Post-CI genotype distribution bar graph showing that when a large viable hybrid population exists, hybrids can reproduce among themselves to regenerate the uninverted parental genotypes and prevent an invading CI from reaching fixation. Selecting the Post-CI Distribution tab in Fig 34c displays the steady-state genotype distribution after CI invasion. As shown, the invading CI population (CI = *aa*0*X*) coexists in a polymorphism with uninverted niche-*A* ecotypes that are continuously replenished through hybrid matings. A balance between migration and selection maintains this polymorphism and prevents fixation of the CI.

**Fig 34e.**
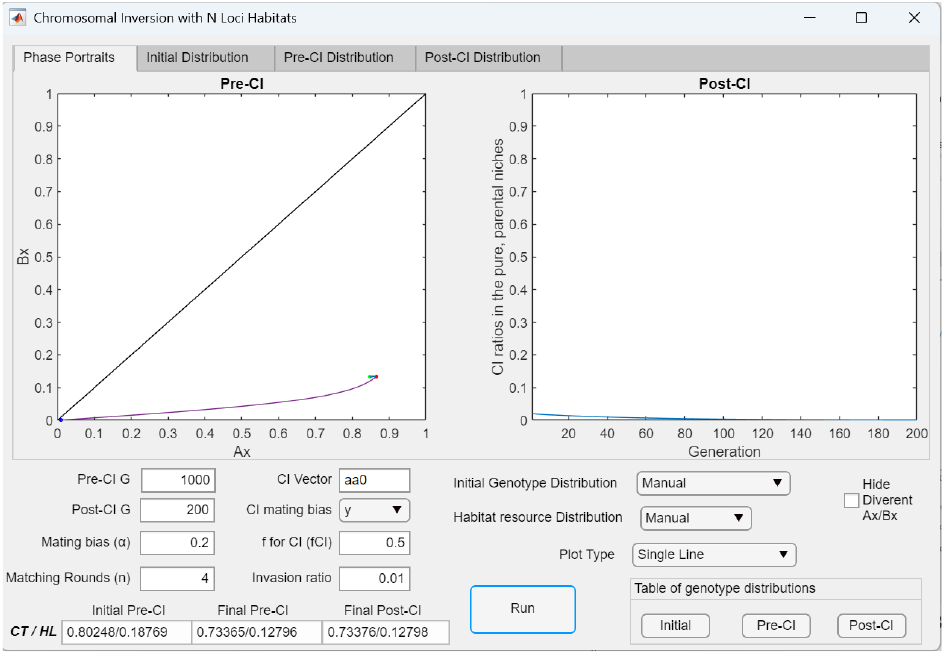
In a system with three-locus ecological genotypes and 40% hybrid habitat resources, a CI capturing locally adaptive ecological alleles but not the prevalent mating-bias allele exhibits reduced invasion fitness. Once nonrandom assortment of locally adaptive ecological and mating-bias alleles has been established across niches, capturing the less prevalent mating-bias allele in a niche decreases the fitness of the CI. The *Invasion Point* plot shows that if the CI in Fig 34c captures the *Y* allele instead of the *X* allele (CI = *aa*0*Y*), it experiences reduced fitness and cannot invade.

Another key observation from the CI invasion plots in Figs 34a and 34c and the post-CI genotype distribution shown in Fig 34d is that, when hybrid habitat resources support the existence of viable ecological hybrids, the invading CIs tend to persist in a polymorphism with uninverted ecotypes in their adaptive niche (niche *A*). This occurs because hybrid matings can continually regenerate the uninverted parental ecotypes. As a result, a balance between immigration and selection maintains the polymorphism between inverted and uninverted genotypes in the parental niche and prevents the CIs from reaching fixation.

Fig 34e shows that when an ecologically adaptive CI mutant captures the wrong mating-bias allele, it suffers reduced fitness and cannot invade. In this example, the CI mutant (*CI* = *aa*0*Y*) captures the prevalent mating-bias allele (*Y*) in niche *B* instead of the prevalent allele (*X*) in niche A, after nonrandom assortment of mating-bias alleles across niches has already been established. This mismatch reduces the CI’s fitness and prevents its invasion.

Lastly, the examples in Figs 35a–35c illustrate that when large populations of viable ecological hybrids are present, hybrid matings can regenerate substantial numbers of uninverted parental ecotypes. Consequently, a large hybrid swarm may overwhelm and ultimately eliminate the CI population from the polymorphism between inverted and uninverted genotypes in the parental niche.

**Fig 35a.**
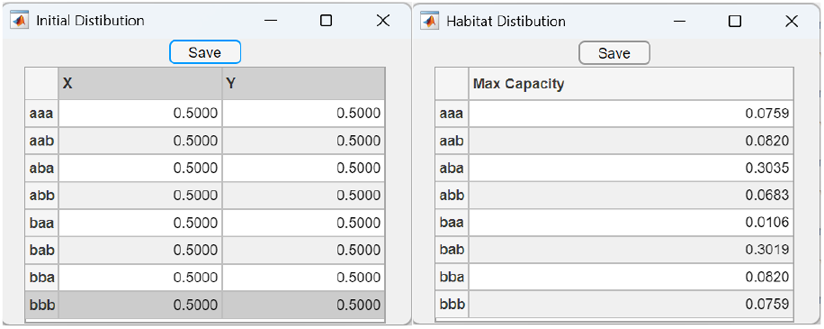
The Initial Genotype Distribution Table and Habitat Resource Distribution Table used in Figs 35b and 35c. The Initial Genotype Distribution Table (left) shows equal starting population ratios of *X* and *Y* alleles across all ecological niches in a sympatric population with three-locus ecological genotypes. The Habitat Resource Distribution Table (right) indicates that approximately 15% of the habitat resources are allocated equally between niches *A* and *B*, while the remaining 85% are distributed among hybrid niches. The exact habitat resource distribution is shown in the bar graph in Fig 35b.

**Fig 35b.**
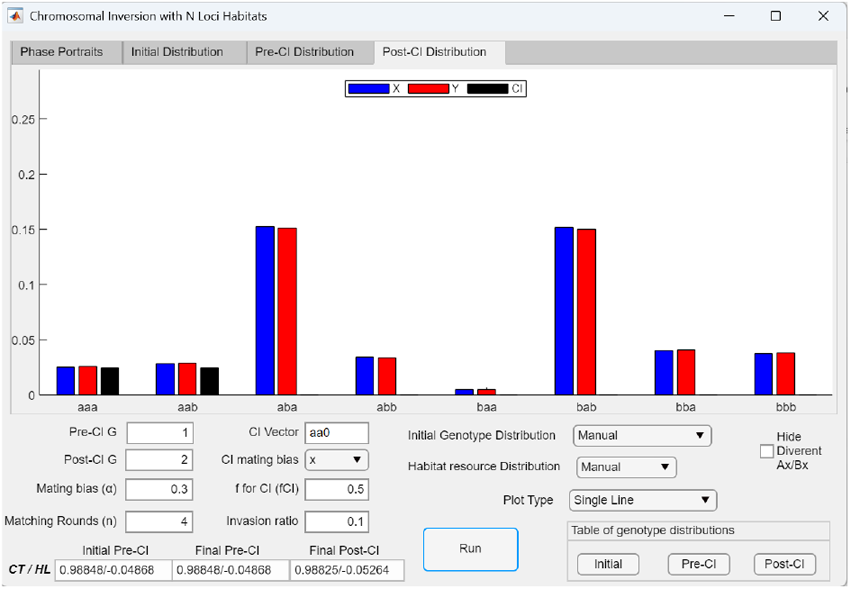
Bar graph showing the initial genotype distribution based on the habitat and genotype distribution tables in Fig 35a, with an initial CI mutant (CI = *aa*0*X*) population ratio of 0.1. With the pre-CI generation set to 1 and the post-CI generation set to 2, the *Post-CI Distribution* bar graph shows a predominance of hybrid niche resources (85% of total resources), while the remaining 15% is shared equally between the parental niche habitats *A* and *B*.

**Fig 35c.**
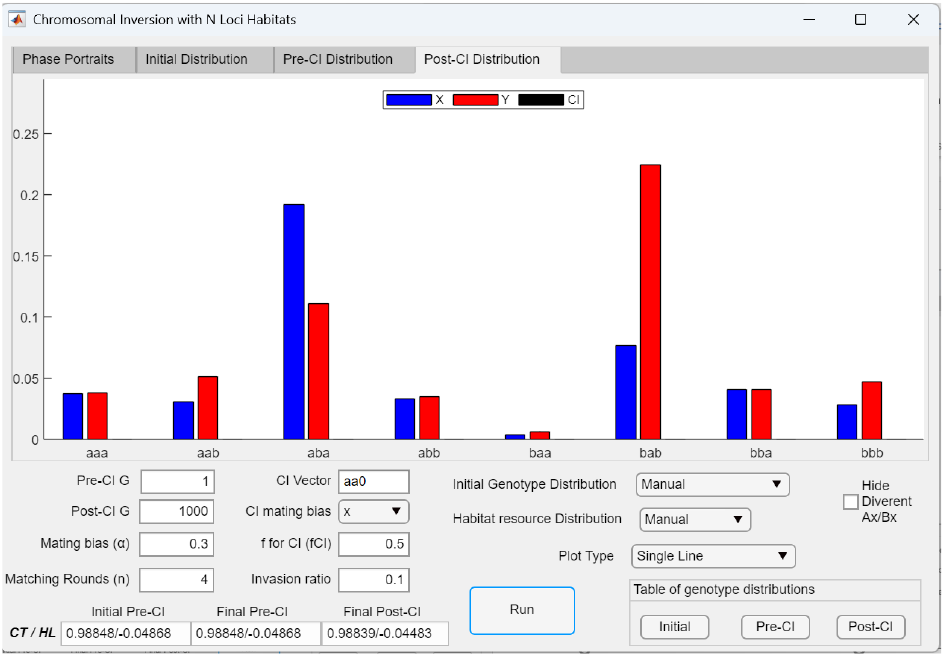
*Post-CI genotype distribution* bar graph showing that when a large viable hybrid population exists, the initial CI population in genotypes near the niche-*A* ecotype is eliminated by a hybrid swarm. Using the same parameter values as in Fig 35b, after 1,000 post-CI generations, the initial CI population (ratio = 0.1) is completely eliminated due to extensive mating and backcrossing within the large hybrid population, which continuously regenerates the uninverted parental genotypes.

### II. SYSTEMS WITH MULTILOCUS MATING-BIAS GENOTYPES

The GUI application MultiSexCI was used to examine how incorporating an arbitrary number of mating-bias gene loci influences system dynamics. To focus solely on pre-CI dynamics, the post-CI phase of the simulation was disabled by setting the number of post-CI generations (*Post*-*CI Gen*) to zero. In subsequent simulations, the post-CI phase was enabled, and CI mutations capturing different combinations of mating-bias alleles were introduced to investigate their invasion dynamics and effects on overall system behavior.

#### 1 The effects of having multilocus mating-bias genotypes (without CI invasion)

We first examined how varying the number of mating-bias gene loci affects population dynamics in the *AX*/*BX* phase portrait. To do so, the post-CI phase was disabled by setting *Post*-*CI G* = 0. Fig 36a shows the *AX*/*BX* phase-portrait solution of the mathematical model in Fig 3, where the mating-bias genotype is determined by a single gene locus and ecological selection is modeled by a two-niche system without viable hybrids. The strength of disruptive ecological selection is specified by the offspring return ratio *f*, which ranges from 0 to 0.5, with lower values indicating stronger ecological selection. As shown in Fig 36a, under the specified parametric conditions, the *AX*/*BX* phase portrait converges to a fixed point, thereby producing premating RI.

**Fig 36a.**
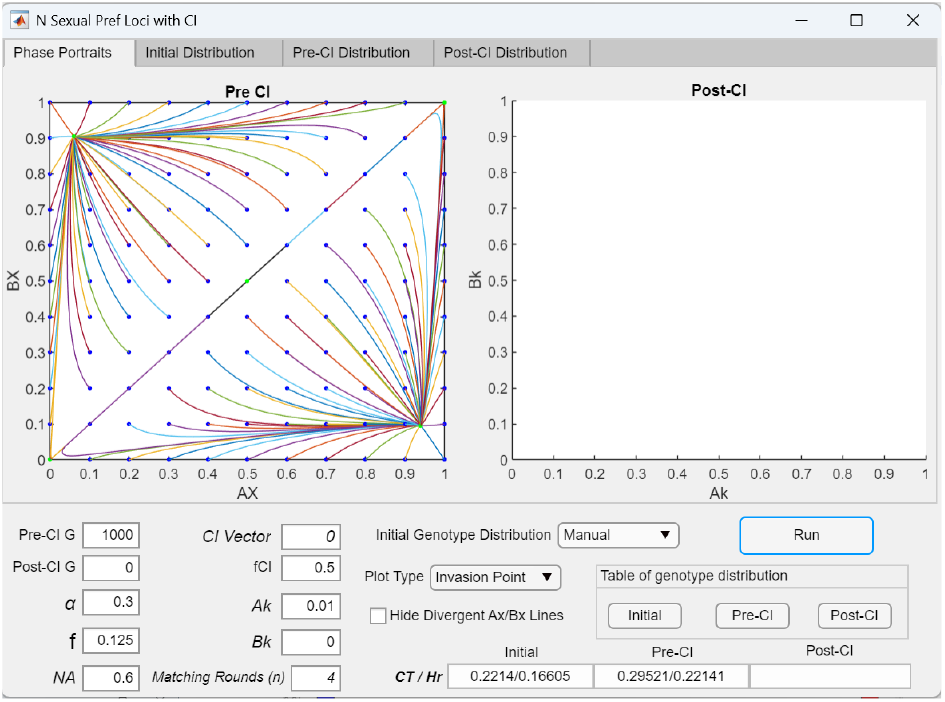
In a two-niche model without viable ecological hybrids, a single gene locus for mating-bias genotypes produces a globally convergent phase portrait. With the post-CI generation set to zero (i.e., no CI present), the GUI application (MultiSexCI) is used to examine how different numbers of mating-bias gene loci affect the two-niche model shown in Fig 3. With a single mating-bias locus (*CI* = 0) and *Pre*-*CI G* = 1000 generations, the *AX*/*BX* phase portrait displays a globally convergent pattern with a fixed-point polymorphism at *AX*/*BX* = 0.94002/0.09567 (see also Fig 7).

**Fig 36b.**
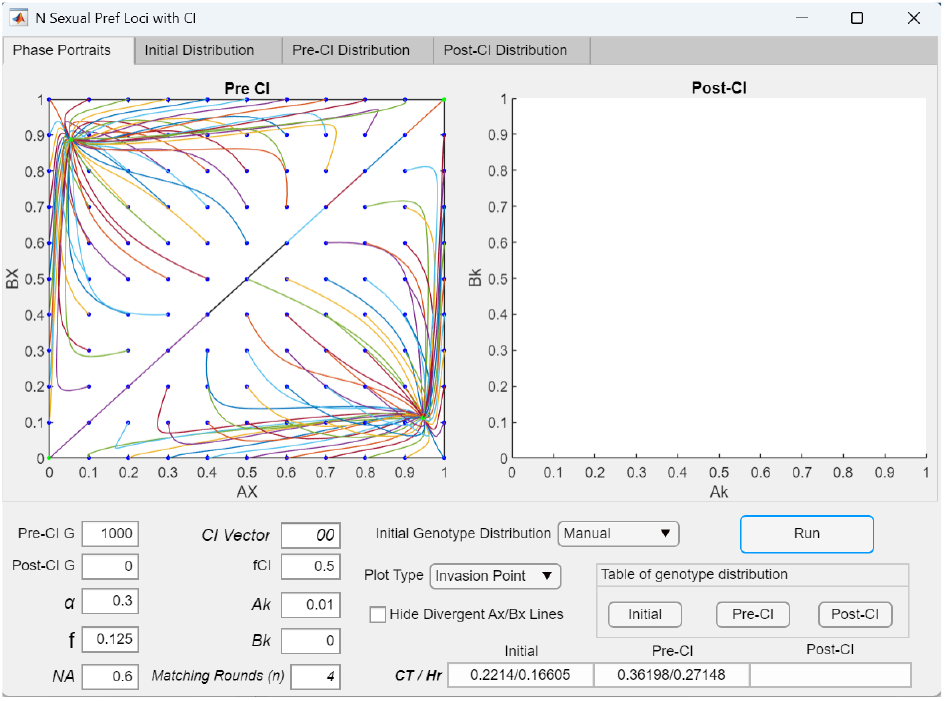
In a two-niche model without viable ecological hybrids, having two gene loci for mating-bias genotypes produces weaker premating RI than having a single gene locus. Increasing the number of mating-bias loci in Fig 36a from one to two (*CI* = 00) still yields a globally convergent *AX*/*BX* phase portrait. However, compared to the one-locus case in Fig 36a, the *AX*/*BX* fixed point shifts further away from the lower-right corner of the phase portrait (to *AX*/*BX* = 0.94620/0.11213), and the fixed-point *CT*/*Hr* values increase from *CT*/*Hr* = 0.29521/0.22142 to 0.36198/0.27148, indicating weaker premating RI.

**Fig 36c.**
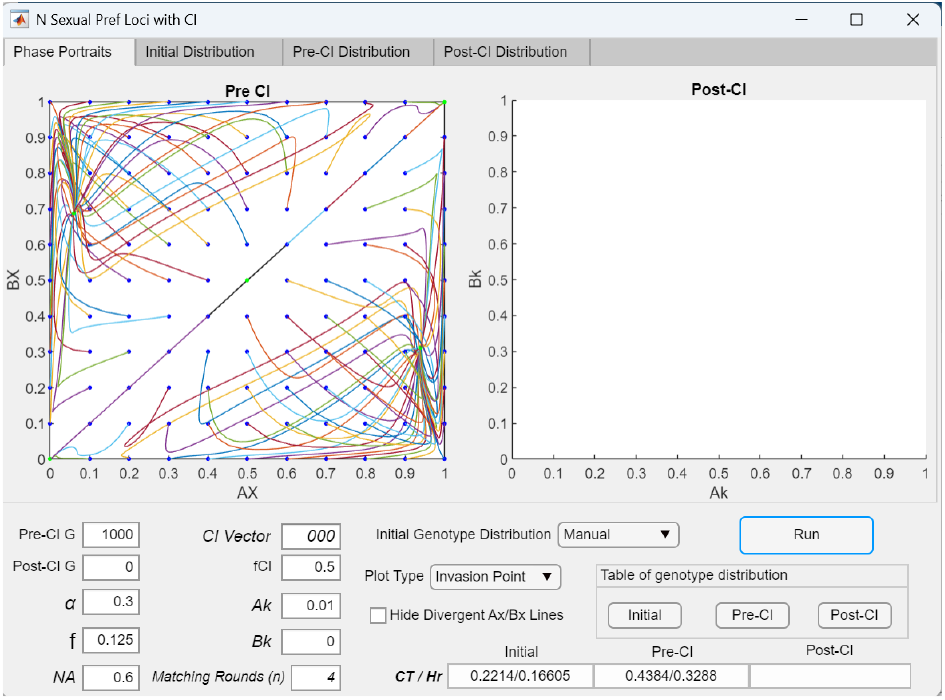
In a two-niche model without viable ecological hybrids, having three gene loci for mating-bias genotypes causes an invasion-resistant pattern to emerge in the phase portrait. Increasing the number of mating-bias loci in Fig 36b from two to three (*CI* = 000) produces an invasion-resistant pattern in the *AX*/*BX* phase portrait. The *AX*/*BX* fixed point shifts even farther from the lower-right corner of the phase portrait, and the fixed-point *CT*/*Hr* values also increase.

**Fig 36d.**
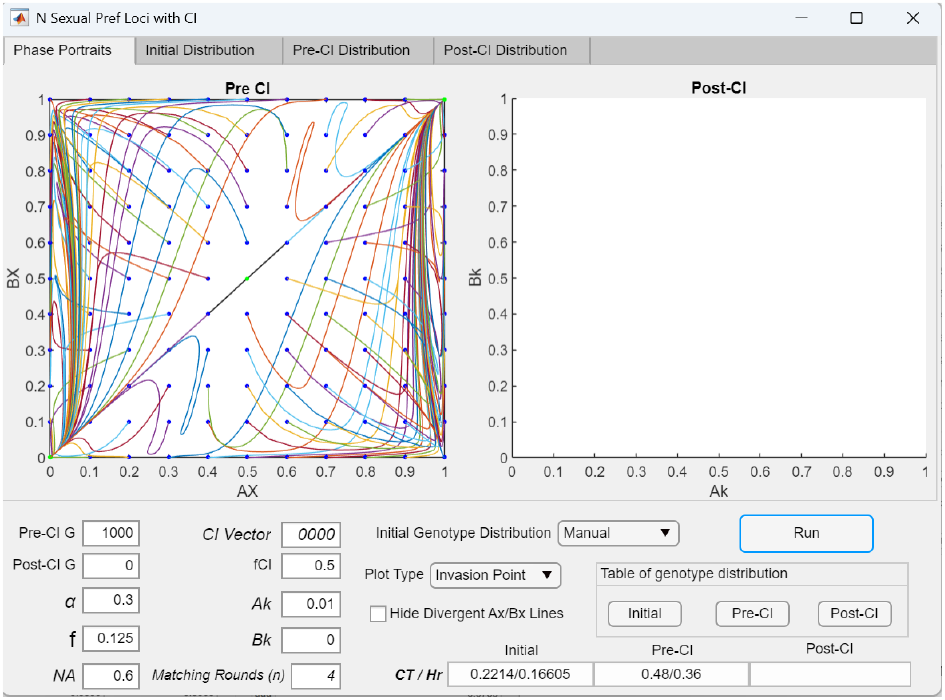
In a two-niche model without viable ecological hybrids, having four gene loci for mating-bias genotypes causes the phase portrait to become divergent. Increasing the number of mating-bias loci in Fig 36c from three to four (*CI* = 0000) results in a divergent *AX*/*BX* phase portrait with no fixed points and no premating RI between the niche-*A* and niche-*B* ecotypes.

**Fig 36e.**
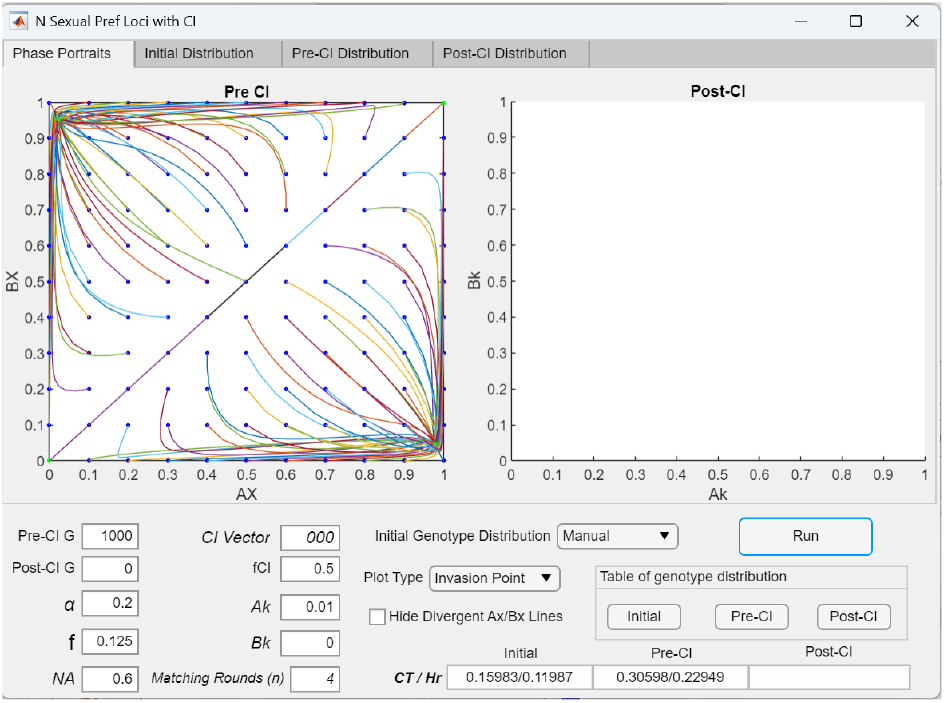
In a two-niche model without viable ecological hybrids and with three gene loci for mating-bias genotypes, increasing the maximum mating bias produces stronger premating RI. Reducing *α* from 0.3 to 0.2 in the three-locus case shown in Fig 36c eliminates the invasion-resistant pattern in the *AX*/*BX* phase portrait. With the stronger mating bias, the *AX*/*BX* fixed point shifts closer to the lower-right corner, and the fixed-point *CT*/*Hr* values decrease from 0.4384/0.3288 to 0.30598/0.22949, reflecting stronger premating RI.

**Fig 36f.**
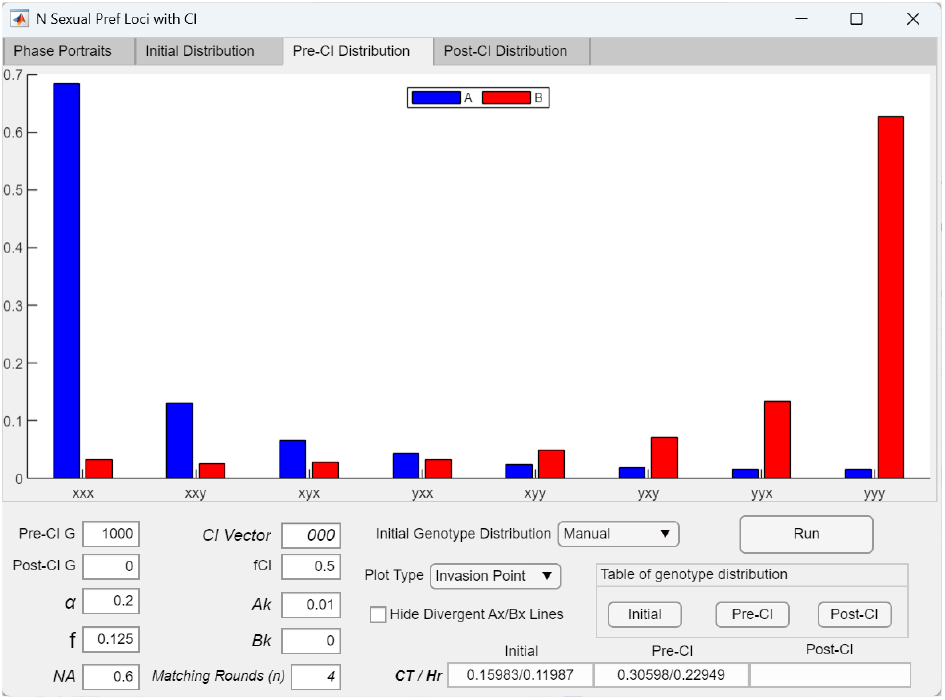
Bar graph showing the fixed-point mating-bias genotype distribution in a system with three mating-bias loci. The *Pre-CI Distribution* bar graph displays the genotype distribution at the *AX*/*BX* fixed point shown in Fig 36e. Because *α* is nonzero, viable mating-bias hybrids are produced, but their distribution is skewed toward the like-kind parental genotypes (i.e., concentrated near the *xxx* and *yyy* ends). This skew reflects nonrandom assortment of incompatible mating-bias genotypes from opposite ends of the Matching Compatibility Table (Fig 15), resulting in premating RI between niche ecotypes.

**Fig 36g.**
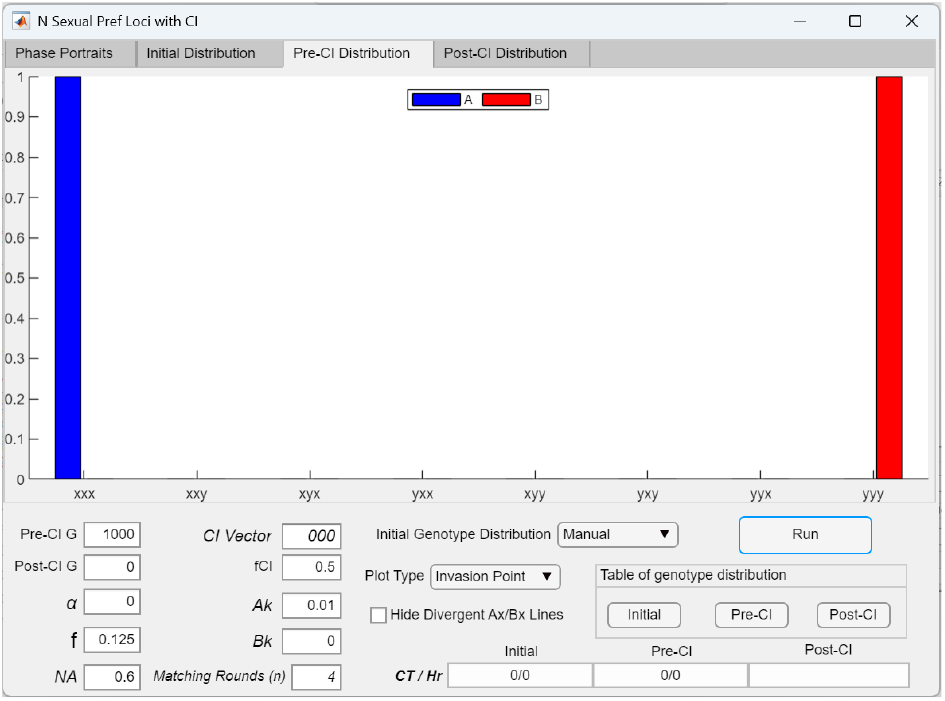
Bar graph showing that when *α* = 0, no viable hybrids exist in the fixed-point mating-bias genotype distribution in a system with three mating-bias loci. When *α* in Fig 36e is set to zero, the *Pre-CI Distribution* bar graph shows that all hybrid genotypes are eliminated, leaving only the extreme parental mating-bias genotypes at the resulting fixed point *AX*/*BX* = 1/0.

**Fig 36h.**
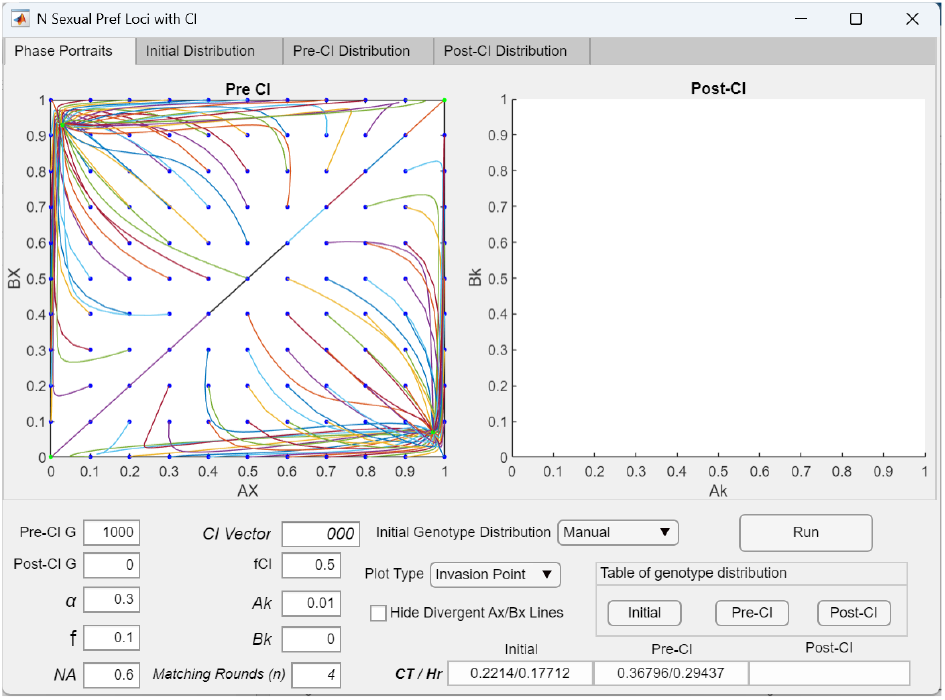
In a two-niche model without viable ecological hybrids and with three gene loci for mating-bias genotypes, increasing the strength of disruptive ecological selection produces stronger premating RI. Reducing *f* from 0.125 to 0.1 in the three-locus case shown in Fig 36c eliminates the invasion-resistant pattern in the *AX*/*BX* phase portrait. With stronger disruptive ecological selection (lower *f*), the *AX*/*BX* fixed point shifts closer to the lower-right corner, and the fixed-point *CT*/*Hr* values decrease from 0.4384/0.3288 to 0.36796/0.29437, reflecting stronger premating RI.

In Figs 36b–36d, we incrementally increase the number of mating-bias gene loci from one to four by adjusting the number of allelic positions in the *CI Vector* input, while keeping all other parametric values identical to those in Fig 36a. The results show that as the number of mating-bias loci increases from one to two and then three, the *AX*/*BX* fixed point progressively shifts away from the lower-right corner of the phase portrait. This shift is accompanied by increased pre-CI *CC* and *HP* values at the fixed points, indicating weakened premating RI as the number of loci increases. With two mating-bias loci, the *AX*/*BX* phase portrait in Fig 36b remains globally convergent, but when the number of loci is increased to three, an invasion-resistant pattern emerges in the *AX*/*BX* phase portrait in Fig 36c, which prevents invasion by small mutant populations near the origin. Further increasing the number of mating-bias loci to four in Fig 36d causes the *AX*/*BX* phase portrait to become globally divergent, with no fixed point and no premating RI.

It should be noted that the *AX*/*BX* phase portrait only tracks changes in the population ratios of the extreme parental mating-bias genotypes (e.g., *xxx* and *yyy*, or *xxxx* and y*yyy*). To determine whether a fixed point may exist at hybrid genotypes elsewhere in the higher-dimensional genotype space beyond the *AX*/*BX* dimension, the post-CI genotype distribution must be examined using the *Post-CI Distribution* tab in the interface. Across all simulation runs, the results consistently indicate that at most one fixed point— represented by a single peak in the genotype distribution—exists within the full high-dimensional phase portrait defined by the number of mating-bias loci.

In Figs 36e–36g, we investigate how varying the maximum mating-bias value *α* in the Matching Compatibility Table (Fig 15) affects system dynamics. As shown in Fig 36e, reducing *α* from 0.3 to 0.2 in a system with three mating-bias loci increases the mating bias between the extreme genotypes, facilitates system convergence, and shifts the fixed point closer to the lower-right corner of the *Ax*/*Bx* phase portrait, resulting in stronger premating RI. This reduction in *α* also removes the invasion-resistance pattern that the *AX*/*BX* phase portrait in Fig 36c exhibits when *α* = 0.3.

A comparison of the pre-CI genotype distributions in Fig 36f, where *α* = 0.2, and Fig 18b, where *α* = 0.3, shows that as *α* decreases, the genotype distribution shifts toward the extreme ends and fewer intermediate hybrids are produced. Correspondingly, both *CT* and *Hr* values decline, indicating stronger RI. In general, as the *AX*/*BX* fixed point shifts closer to the lower-right corner of the phase portrait—reflecting increased premating RI—the genotype distribution becomes more concentrated at the extremes and leaves fewer intermediate hybrids.

When *α* is reduced to zero, no mating-bias hybrid offspring are produced. As shown in Fig 36g, only the extreme mating-bias genotypes (*xxx* and *yyy*) remain at the fixed point. Both *CT* and *Hr* are zero, and premating RI is complete.

In Fig 36h, strengthening disruptive ecological selection by reducing the value of *f* in Fig 36c from 0.125 to 0.1 produces the same effect as reducing the mating-bias parameter *α*. This reduction in *f* also removes the invasion-resistance pattern observed in Fig 36c and shifts the *AX*/*BX* fixed point closer to the lower-right corner of the phase portrait.

Fig 37a demonstrates that increasing the number of matching rounds (*n*) can reduce the invasion-resistant region in the *AX*/*BX* phase portrait in systems with multiple mating-bias loci, just as in the case of systems with a single mating-bias locus.

**Fig 37a.**
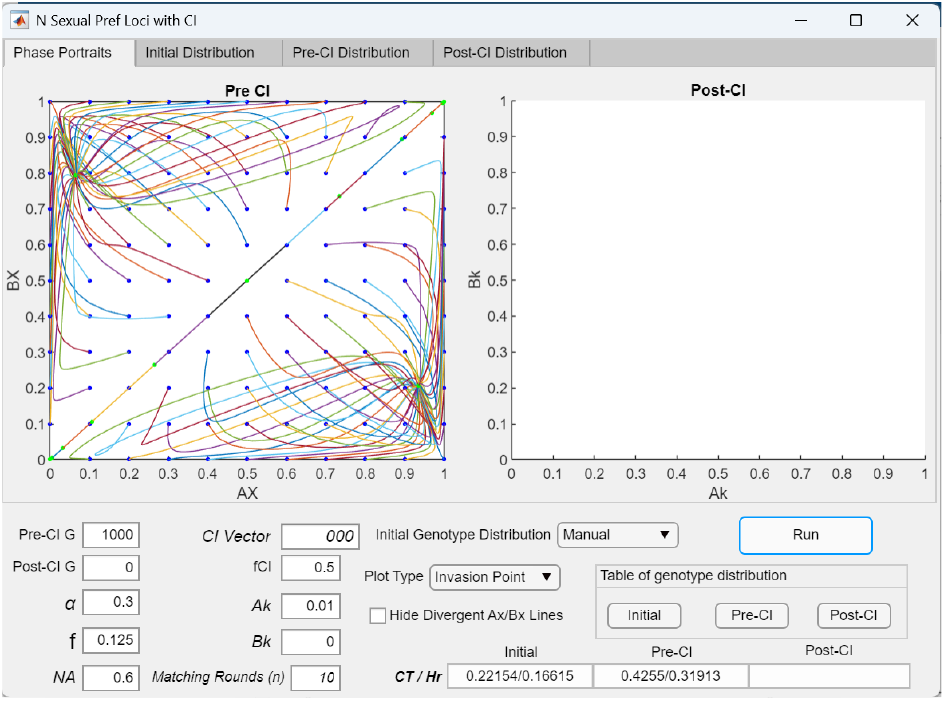
Increasing the number of matching rounds (*n*) reduces the invasion-resistant region in the *AX*/*BX* phase portrait of a system with three mating-bias loci. When the number of matching rounds in Fig 36c is increased from *n* = 4 to *n* = 10, while keeping all other parametric values unchanged, the invasion-resistant region in the *AX*/*BX* phase portrait is reduced toward the origin.

**Fig 37b.**
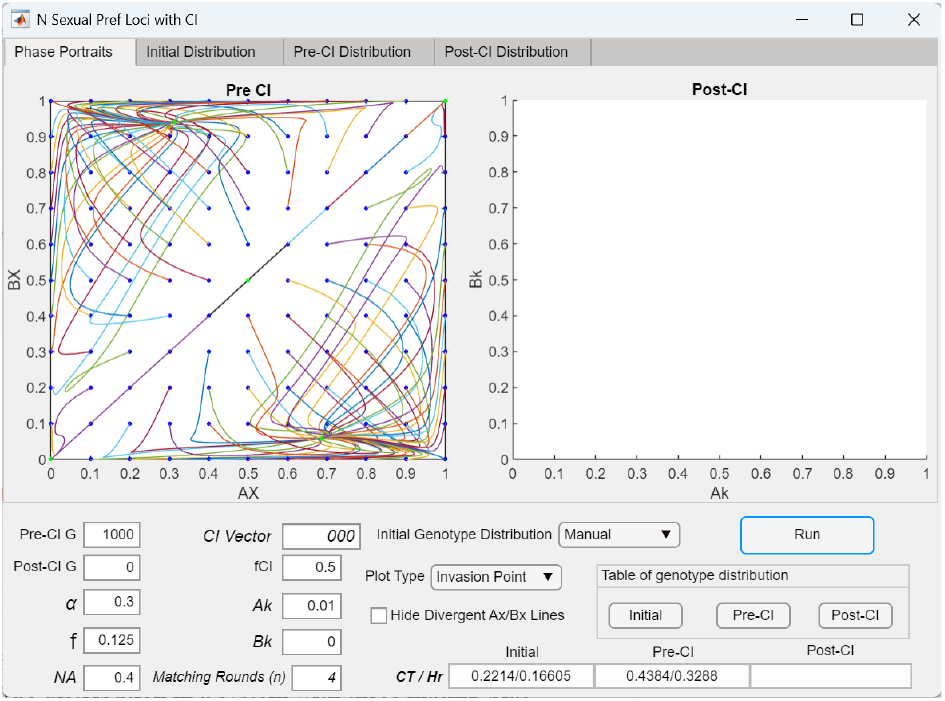
Decreasing the size of niche *A* (*NA*) reduces the invasion-resistant region along the *A*-axis in the *AX*/*BX* phase portrait of a system with three mating-bias loci. When the niche-*A* size in Fig 36c is reduced from *NA* = 0.6 to *NA* = 0.4, while keeping all other parametric values unchanged, the invasion-resistant region contracts away from the *x*-axis. This creates a wedge-shaped area lying along the *x*-axis and pointing toward the origin, within which invasion is possible for initial *AX*/*BX* population ratios falling inside this region. As *NA* is further reduced toward 0.3, the wedge-shaped region becomes increasingly compressed toward the *x*-axis, and the fixed point shifts closer to the *x*-axis and farther from the lower-right corner of the phase portrait. When *NA* ≤ 0.3, the wedge-shaped region disappears entirely, and the *AX*/*BX* phase portrait becomes divergent. Similarly, increasing *NA* greater than 0.5 produces a comparable effect along the *y*-axis, and the phase portrait becomes globally divergent once *NA* ≥ 0.7.

Similarly, as illustrated in Fig 37b, reducing the niche size *NA* may open a wedge-shaped region along the x-axis within the invasion-resistant pattern, which enables invasion by extreme-*X* (*AX*) mutant populations. However, there appears to be a lower threshold of *NA* below which the *AX*/*BX* phase portrait becomes divergent, as well as an upper threshold above which divergence also occurs. These effects arise symmetrically around *NA* = 0.5. More generally, simulations with different numbers of mating-bias loci also show that *NA* = 0.5 yields the greatest reduction in the invasion-resistant region, and outside a symmetric range around this value, the system becomes divergent.

#### 2 Invasion of a CI capturing mating-bias alleles in a two-niche system with multilocus mating-bias genotypes

The invasion dynamics of a CI capturing mating-bias alleles in the multilocus mating-bias model are examined by specifying the CI Vector and setting the Post-CI G to be greater than zero.

The example in Fig 38 shows that when *α* = 0 in a system with three mating-bias gene loci, no mating-bias hybrids are produced in the pre-CI phase. Under these conditions, a CI capturing mating-bias alleles cannot invade because the invasion fitness of such a CI depends on its ability to reduce the effective number of mating-bias loci and suppress the production of intermediate mating-bias hybrids, thereby enhancing premating RI and reducing maladaptive ecological hybrid loss. When no intermediate mating-bias hybrids are present to be eliminated, there is no selective advantage for the CI to invade.

**Fig 38.**
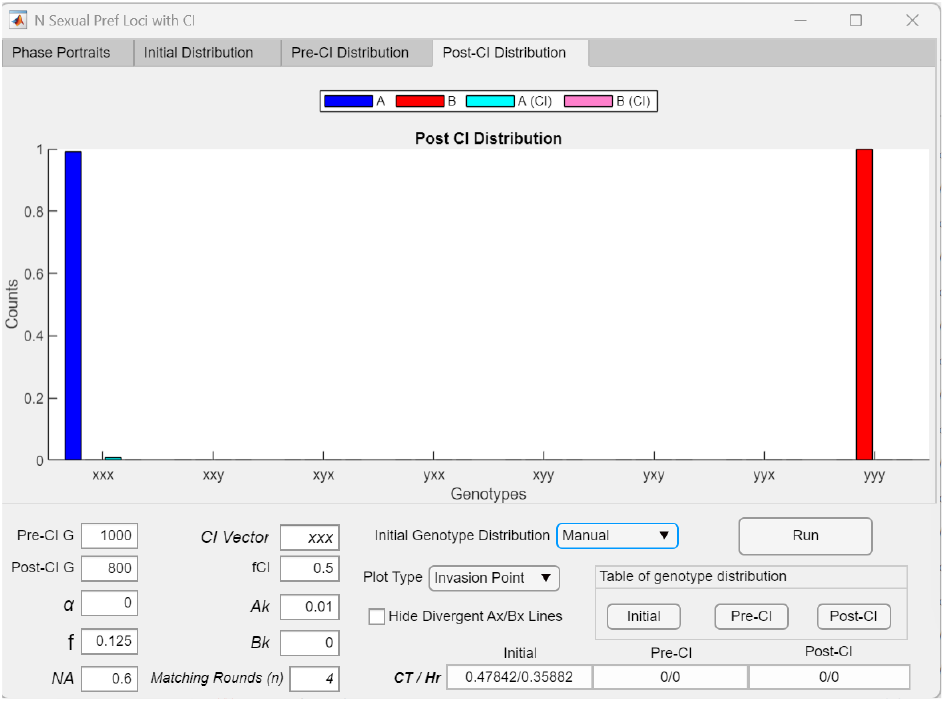
*Post-CI Distribution* bar graph showing that an adaptive CI (*CI* = *xxx*) cannot invade when *α* = 0 and no mating-bias hybrids exist in a system with three mating-bias loci. In Fig 36g, setting *α* = 0 eliminates all mating-bias hybrids at the *AX*/*BX* fixed point. When *α* = 0, an adaptive CI mutant (*CI* = *xxx*) cannot invade because its invasion fitness depends on reducing the production of mating-bias hybrid offspring—by lowering the effective number of mating-bias loci experienced by the system—and thereby increasing premating RI to reduce maladaptive ecological hybrid loss. When *α* = 0 and *CT*/*Hr* = 0/0, maximum mating bias has already been achieved, which leaves no room for a CI to further enhance it.

In Fig 39a, increasing the value of *α* from 0 to 0.3 in Fig 38 restores ecological hybrid loss due to disruptive ecological selection. Under the same parametric conditions as in Fig 36c, the pre-CI *AX*/*BX* phase portrait again displays an invasion-resistant pattern that prevents invasion by small extreme-*X* mutant populations near the origin. However, in the presence of hybrid loss, a CI mutant capturing all extreme-*X* mating-bias alleles (*CC* = *xxx*) gains enhanced invasion fitness, as genotypes carrying the CI experience reduced ecological hybrid offspring loss in inter-niche matings compared to same-niche counterparts without the inversion. This fitness advantage enables the CI to overcome the invasion-resistant pattern and successfully invade, resulting in a post-CI *AX*/*BX* fixed point that lies closer to the lower-right corner of the phase portrait and yields stronger premating RI. As shown in Fig 39b, the post-CI genotype distribution consists exclusively of extreme mating-bias genotypes, with no intermediate hybrids. This outcome demonstrates that, by reducing the effective number of mating-bias gene loci from three to one, the CI is able to eliminate the production of mating-bias hybrids and facilitate system convergence.

**Fig 39a.**
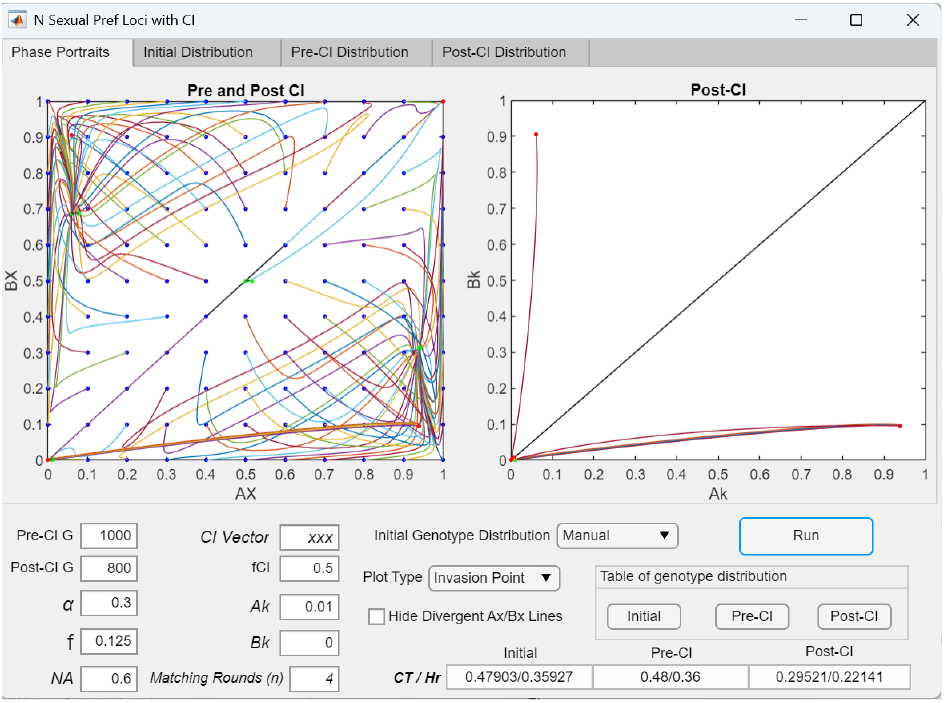
When ecological hybrid loss is present (*α* = 0. 3), an adaptive CI mutant (*CI* =*xxx*) can overcome the invasion-resistant pattern in the *AX*/*BX* phase portrait and invade to establish fixed-point polymorphism and premating RI. Increasing *α* in Fig 38 from 0 to 0.3 restores ecological hybrid loss due to disruptive ecological selection, causing the pre-CI *AX*/*BX* phase portrait to once again display the invasion-resistant pattern shown in Fig 36c. Under these conditions, hybrid loss allows a locally adaptive CI mutant (*CI* = *xxx, Ak* = 0.01), arising from a small extreme-*X* mutant population (*Axxx* = 0.01, as specified in Fig 40), to overcome the invasion-resistant pattern and successfully invade. This results in a post-CI *AX*/*BX* fixed point that lies closer to the lower-right corner of the phase portrait and produces stronger premating RI.

**Fig 39b.**
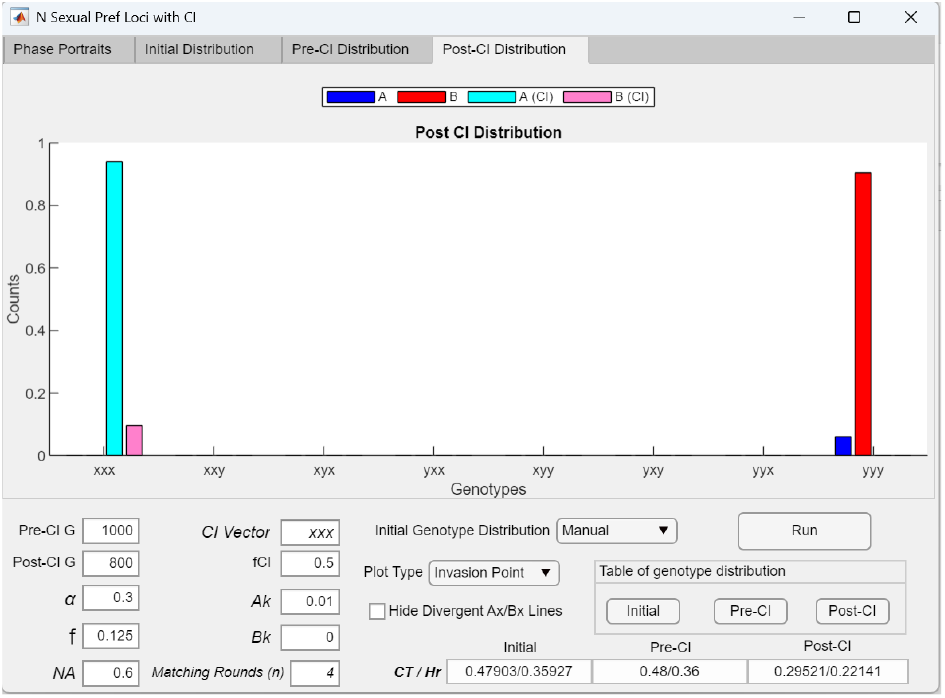
*Post-CI Distribution* bar graph showing the genotype distribution following the invasion of an adaptive CI mutant (*CI* = *xxx*) in a system with three mating-bias loci. The bar graph displays the genotype distribution after the CI invasion shown in Fig 39a. As illustrated, the CI reduces the effective number of mating-bias loci from three to one and suppresses the production of intermediate mating-bias hybrids, leading to premating RI, as reflected by reduced post-CI *CT* and *Hr* values.

The Initial Genotype Distribution Table in Fig 40 illustrates the scheme used to specify the initial extreme-*X* mutant population from which a CI mutant arises. This small population provides the *x*-type alleles at each mating-bias locus for the CI to capture. In general, the extreme-*X* population is set to a ratio of 0.01, and the remainder of the population consists solely of the extreme-*Y* genotype (*AX* = 0.01, *AY* = 0.99, *BX* = 0, *BY* = 1).

**Fig 40.**
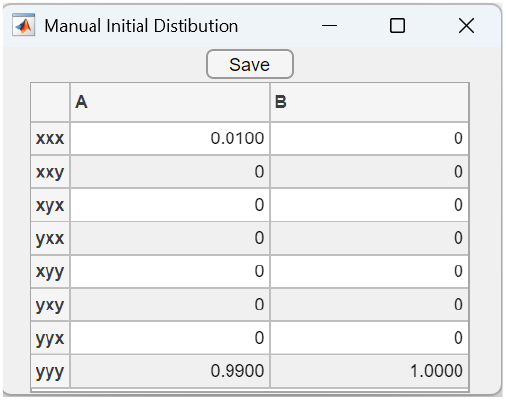
An Initial Genotype Distribution Table specifying the presence of a small population of extreme-*X* mating-bias genotype in a two-niche system otherwise fixed for the extreme-*Y* genotype. In this example for a system with three mating-bias loci, a small population of extreme-*X* genotype (*xxx*) arises in the niche-*A* ecotype (*Axxx* = 0.01) within a sympatric population composed entirely of the extreme-*Y* genotypes (*Ayyy* and *Byyy*). A CI mutant (*Ak* = 0.01) that captures all or part of the available *x*-type alleles at the mating-bias loci is assumed to originate from this initial extreme-*X* population located near the origin of the *AX*/*BX* phase portrait. The same initialization scheme is applied to systems with different numbers of mating-bias loci. In such cases, the extreme-*X* population in niche *A* is set to 0.01 (*AX* = 0.01), and the remaining population is assigned the extreme-*Y* genotype (*AY* = 0.99, *BX* = 0, *BY* = 1).

Fig 41a shows that a CI mutant capturing three of four x-type mating-bias alleles (*CI* = *xxx*0) and arising in niche *A* is able to invade a divergent *AX*/*BX* system that has no pre-CI fixed point and is composed entirely of the extreme-*Y* genotype. However, as shown in Fig 41b, the CI does not become fixed at the extreme-*X* genotype (*Axxxx*). Instead, it ultimately fixes at the hybrid mating-bias genotype *Axxxy*.

**Fig 41a.**
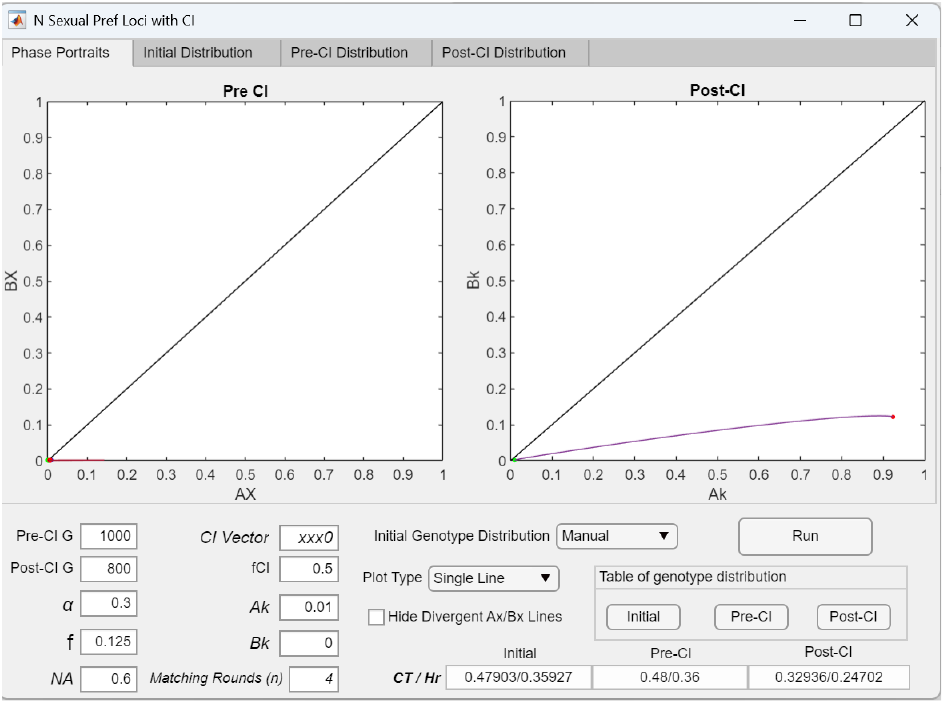
A CI mutant (*CI* = *xxx*0) capturing a partial set of extreme mating-bias alleles successfully invades and becomes fixed at a hybrid mating-bias genotype in a system with four mating-bias loci. Given the parametric values in Fig 36d, the pre-CI *AX*/*BX* phase portrait is divergent, with no fixed points. However, as shown in this *Single Line* plot, a CI mutant capturing three of the four *X* alleles (*CI* = *xxx*0) can nonetheless invade and establish premating RI, though it ultimately fixes at the hybrid genotype *xxxy* rather than the extreme genotype *xxxx* (see the post-CI distribution in Fig 41b). This occurs because the *x*-type allele at the fourth locus is eliminated early in the invasion due to the small initial CI population size and the weak mating bias *α*. Consequently, the invading CI mutation (Ak = 0.01) can only fix at the hybrid genotype *xxxy*. Because the *AX*/*BX* phase portrait tracks only the extreme genotypes (*xxxx* and *yyyy*), the post-CI *AX*/*BX* population ratio converges to the origin in the *AX*/*BX* phase portrait.

**Fig 41b.**
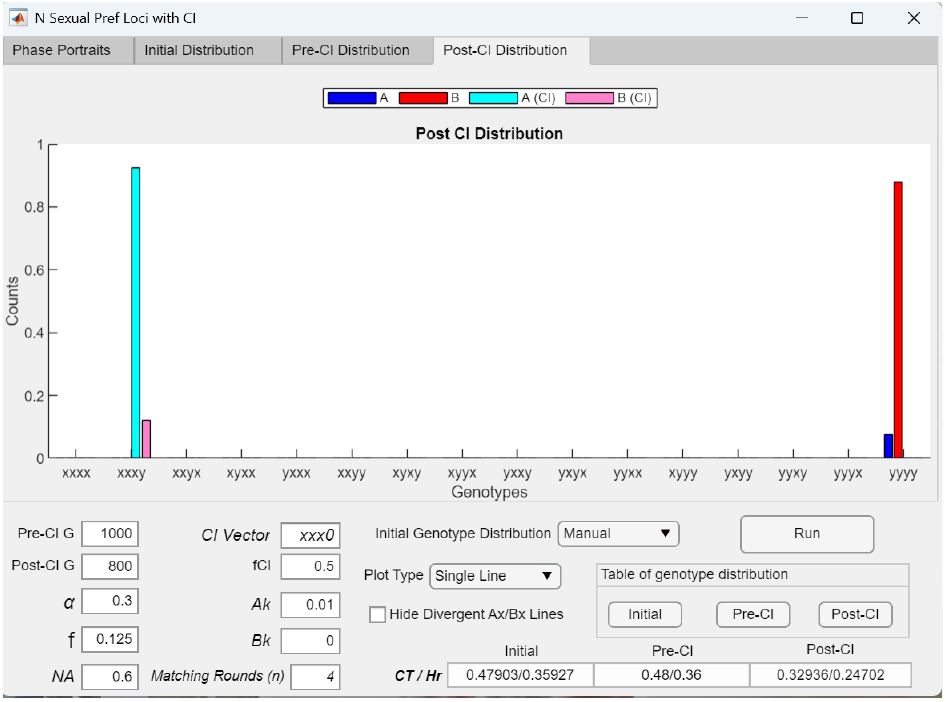
*Post-CI Distribution* bar graph showing the genotype distribution following the invasion of an adaptive CI mutant (*CI* = *xxx*0) in a system with four mating-bias loci. The bar graph displays the genotype distribution after the CI invasion shown in Fig 41a. As illustrated, the invading CI becomes fixed at the hybrid genotype *xxxy*.

**Fig 41c.**
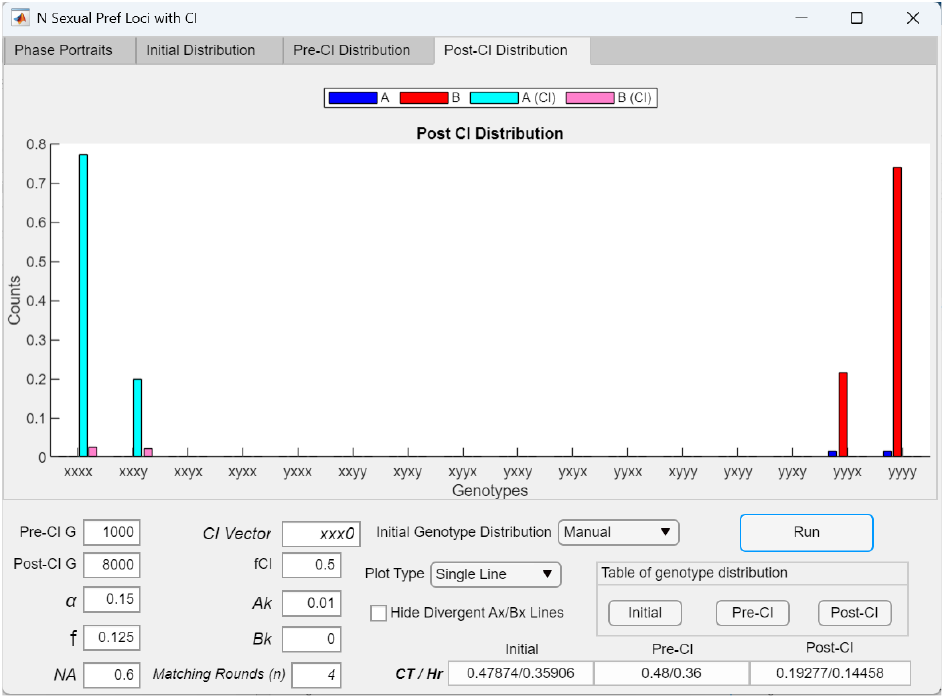
Bar graph showing the post-CI genotype distribution following invasion of an adaptive CI mutant (*CI* = *xxx*0) under increased maximum mating bias (*α* = *a*. 0.15) in a system with four mating-bias loci. Decreasing *α* from 0.3 to 0.15 in Fig 41b shifts the predominant genotype in niche *A* to the extreme-*X* type (*Axxx*) rather than *Axxy*, as seen in Fig 41b, thereby resulting in stronger premating RI following CI invasion.

**Fig 41d.**
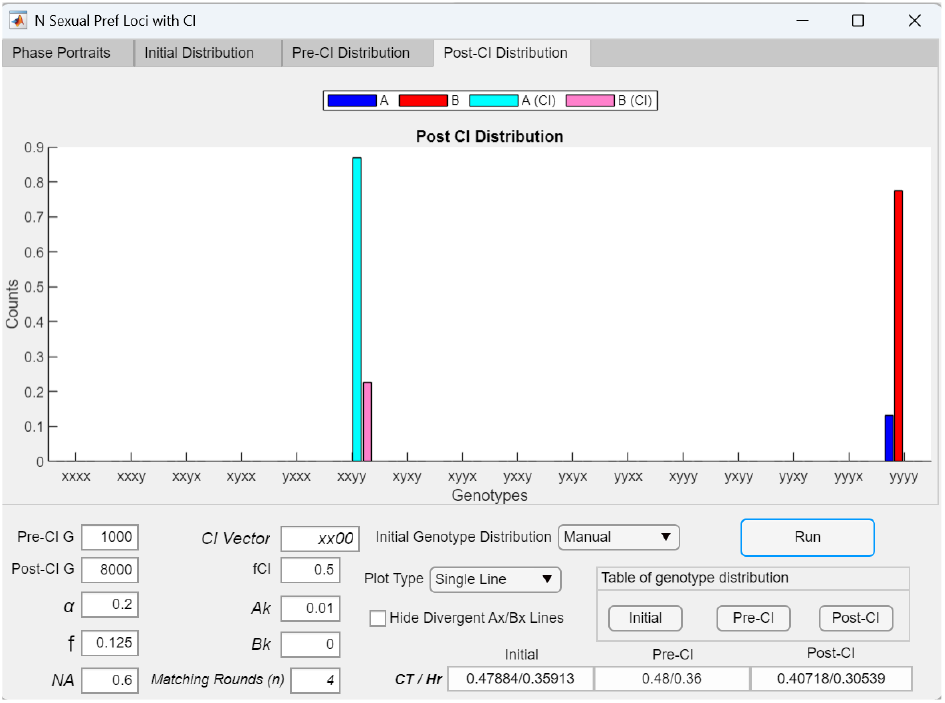
A CI mutant (CI = *xx*00) capturing a partial set of extreme mating-bias alleles successfully invades and becomes fixed at a hybrid mating-bias genotype when *α* = 0. 2 in a system with four mating-bias loci. Under the parametric conditions shown in Fig 41a, when *α* = 0.3, a CI mutant that captures two of the four *x*-type mating-bias alleles in niche *A* (*CI* = *xx*00) is unable to invade. However, if *α* is reduced from 0.3 to 0.2, the CI successfully invades and establishes premating RI, although it ultimately becomes fixed at the hybrid genotype *Axxyy*.

**Fig 41e.**
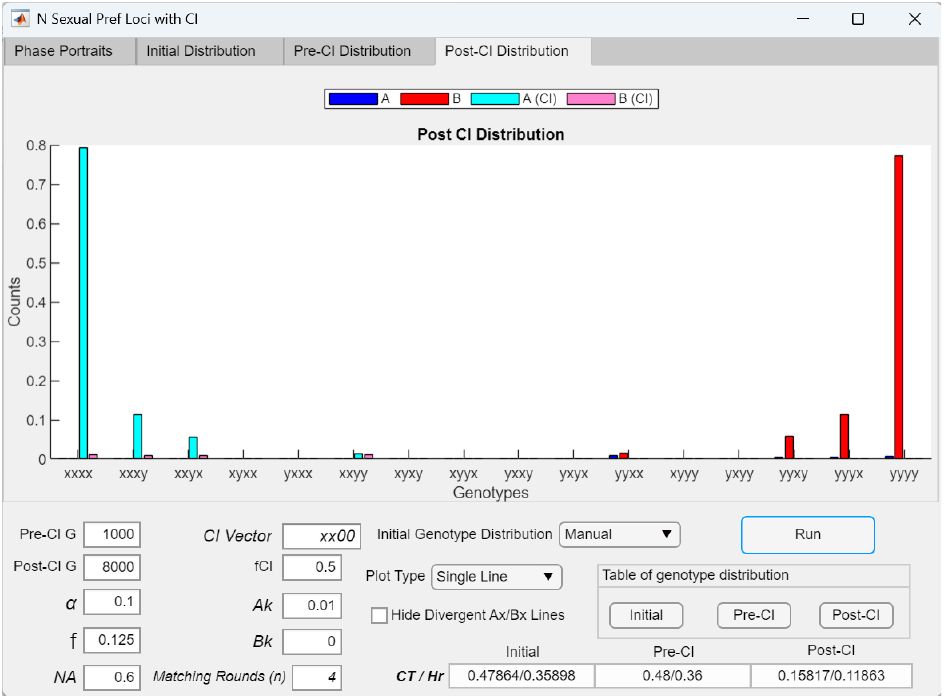
*Post-CI Distribution* bar graph showing the genotype distribution following the invasion of an adaptive CI mutant (*CI* = *xxx*0) when *α* is decreased to 0.1 in a system with four mating-bias loci. Further decreasing *α* in Fig 41d from 0.2 to 0.1 establishes the extreme-*X* genotype (*xxxx*) as the predominant mating-bias genotype in niche A.

A likely explanation is that when the *Axxxx* mutant first arises in niche *A*, which is initially composed entirely of the *Ayyyy* genotype, CI introgression through intra-niche matings rapidly generates a substantial *Axxxy* population. The low mating bias between *Axxxx* and *Axxxy* increases intra-niche matings and reduces inter-niche matings for *Axxxx*. At *α* = 0.3, the mating bias favoring *Axxxx* in inter-niche matings is too weak to offset its low frequency, allowing the much larger *Axxxy* population to gain greater fitness from more inter-niche matings despite its weaker inter-niche mating bias. Consequently, *Axxxy* eliminates *Axxxx* early in the invasion, before *Axxxx* can increase in size.

However, when the maximum mating bias between the extreme genotypes is strengthened, the outcome changes. As shown in Fig 41c, reducing *α* from 0.3 to 0.15 strengthens the mating bias between *Axxxx* and *Byyyy*, restoring the inter-niche mating advantage of the *Axxxx* genotype. This enhanced advantage enables the extreme-*X* genotype (*Axxxx*) to become predominant in niche *A* following CI invasion, resulting in stronger premating RI.

Similar invasion dynamics are observed when the invading CI mutant captures only two of the four extreme-*X* mating-bias alleles (*CC* = *xx*00). Under the same parametric conditions as in Fig 41a, this CI (*CI* = *xx*00) is unable to invade the divergent *AX*/*BX* system when *α* = 0.3. However, as shown in Fig 41d, reducing *α* from 0.3 to 0.2 allows the CI to invade and eventually become fixed at the hybrid genotype *Axxxy*. Further reducing *α* to 0.1 in Fig 41e enables the extreme-X genotype (*Axxxx*) to become predominant in niche *A* following CI invasion, resulting in stronger premating RI.

In general, a CI mutant that captures only a single mating-bias allele gains no fitness advantage over its same-niche counterparts without the inversion and cannot invade. This is because, to benefit from suppressing the breakdown of favorable allele combinations, a CI must capture more than one advantageous allele to preserve their linkage and epistatic effects. This outcome is demonstrated in Fig 42a, where a CI mutant capturing only one of three *x*-type mating-bias alleles (*CI* = *X*00) in a three-locus system fails to invade. Consequently, as shown in Fig 42b, the post-CI genotype distribution remains unchanged from the pre-CI phase, and the post-CI *CT* and *Hr* values are identical to their pre-CI values.

**Fig 42a.**
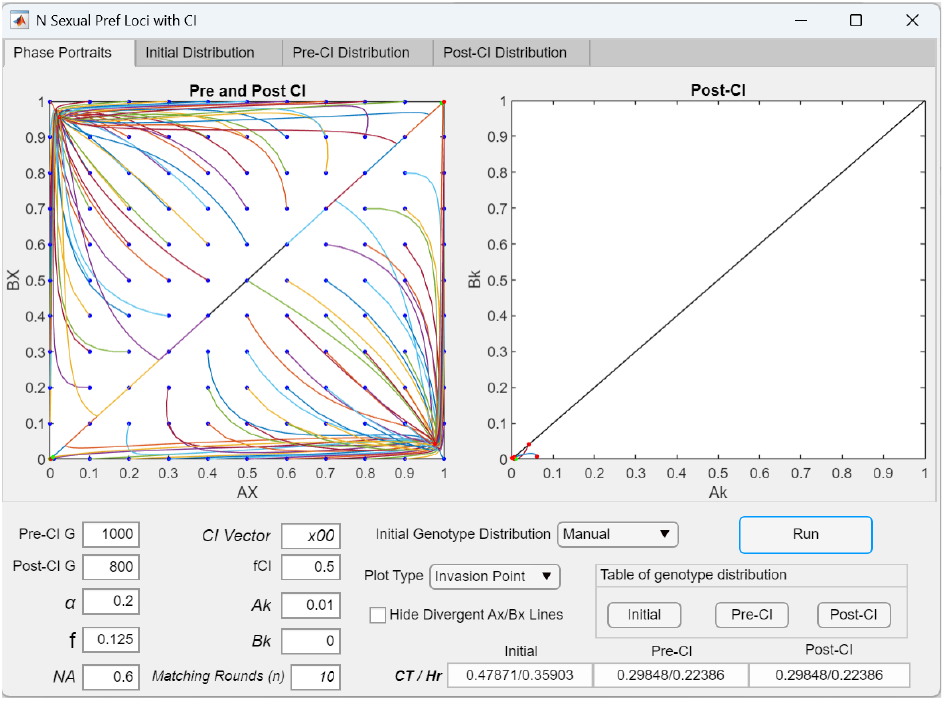
A CI mutant (*CI* = *x*00) capturing only a single mating-bias allele acquires no fitness advantage and cannot invade. Given the parametric values shown, the pre-CI *AX*/*BX* phase portrait is globally convergent with a fixed point at *AX*/*BX* = 0.9772/0.0430 in a system with three mating-bias loci. In the post-CI phase, a CI capturing only one locally prevalent mating-bias allele (*CI* = *x*00) gains no fitness benefit, because preserving a favorable association of linked alleles requires capturing more than one allele. As a result, the CI has no invasion advantage and cannot invade.

**Fig 42b.**
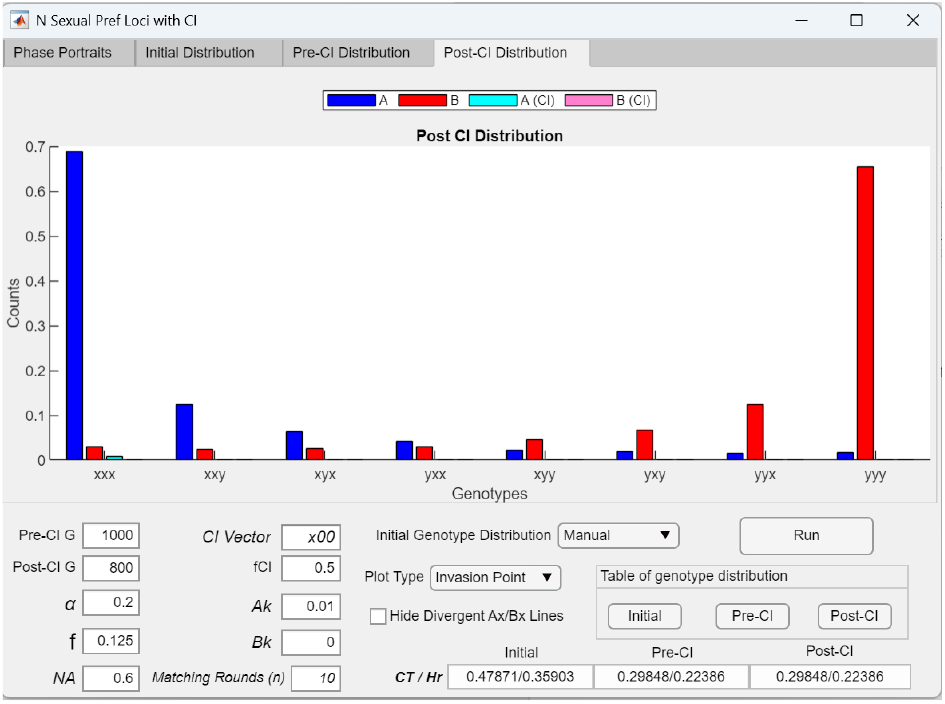
*Post-CI Distribution* bar graph showing the genotype distribution after a CI mutant (*CI* = *x*00) capturing only a single locally favored mating-bias allele fails to invade. The bar graph displays the genotype distribution following the failed CI invasion shown in Fig 42a. As shown, the distribution remains unchanged from the pre-CI *AX*/*BX* fixed-point distribution present before the CI mutant appeared.

**Fig 42c.**
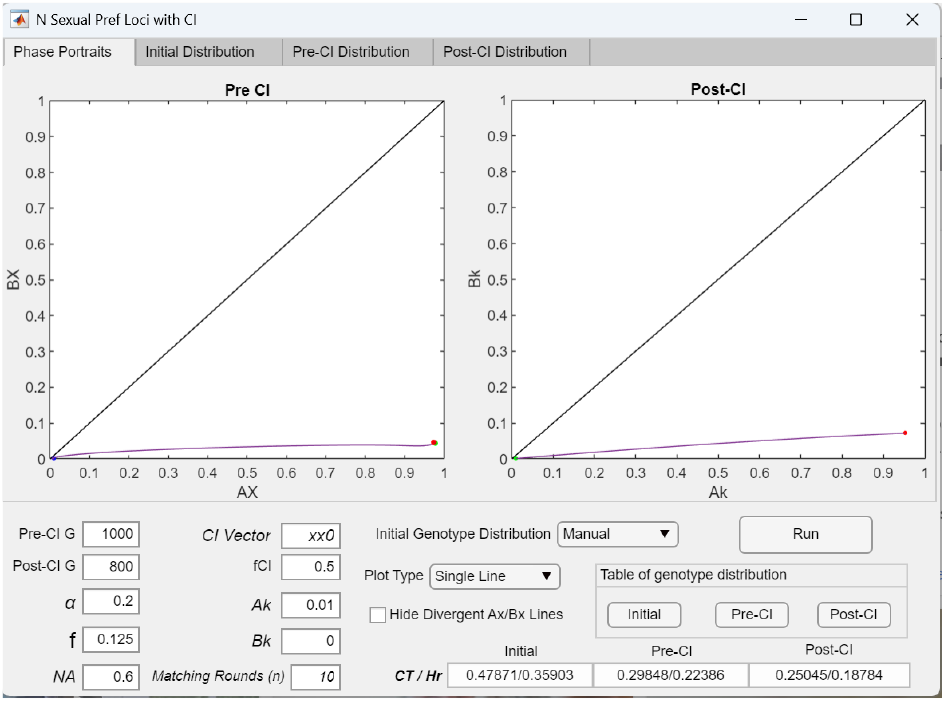
A CI mutant (*CI* = *xx*0) capturing two of the three locally prevalent mating-bias alleles in a convergent system with an *AX*/*BX* fixed-point polymorphism gains a fitness advantage that allows it to invade and strengthen existing premating RI. Using the same parametric values as in Fig 42a, a CI mutant that captures two locally favored mating-bias alleles (*CI* = *xx*0) benefits from the linkage among these alleles and gains an invasion fitness advantage. As the CI spreads through the niche-*A* population, it produces stronger premating RI, as indicated by reduced post-CI *CT* and *Hr* values. The time required to reach 95% of the steady-state *Ak* value (0.9504) is 488 generations (*T*_95_ = 488).

**Fig 42d.**
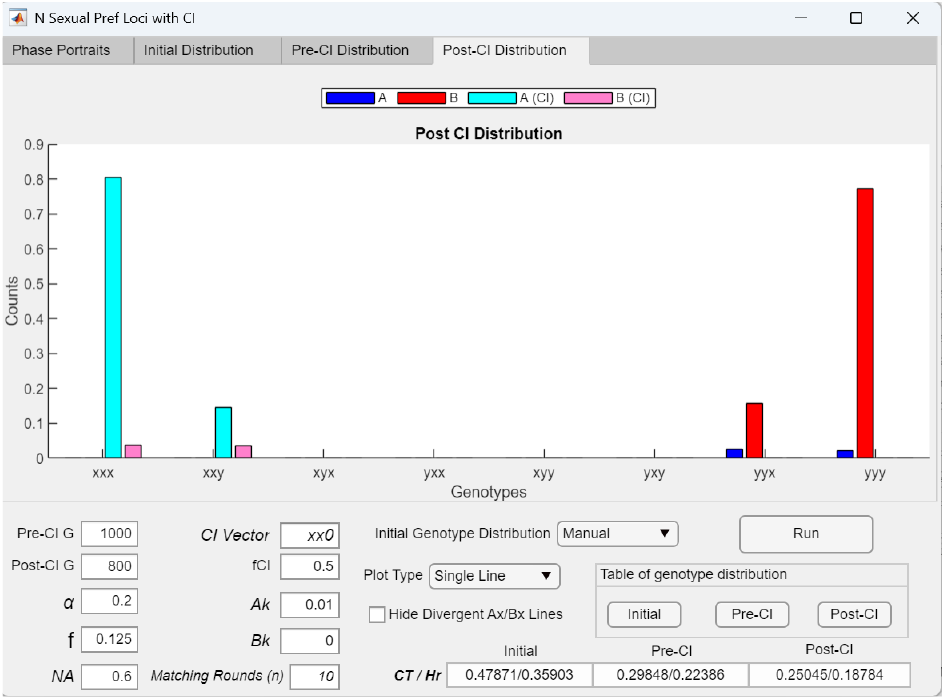
*Post-CI Distribution* bar graph showing the genotype distribution after a CI mutant (*CI* = *xx*0) invades and strengthens premating RI. The bar graph displays the genotype distribution following the CI invasion shown in Fig 42c. It reveals nonrandom assortment toward the extreme genotypes, with smaller hybrid populations persisting near those extremes.

**Fig 42e.**
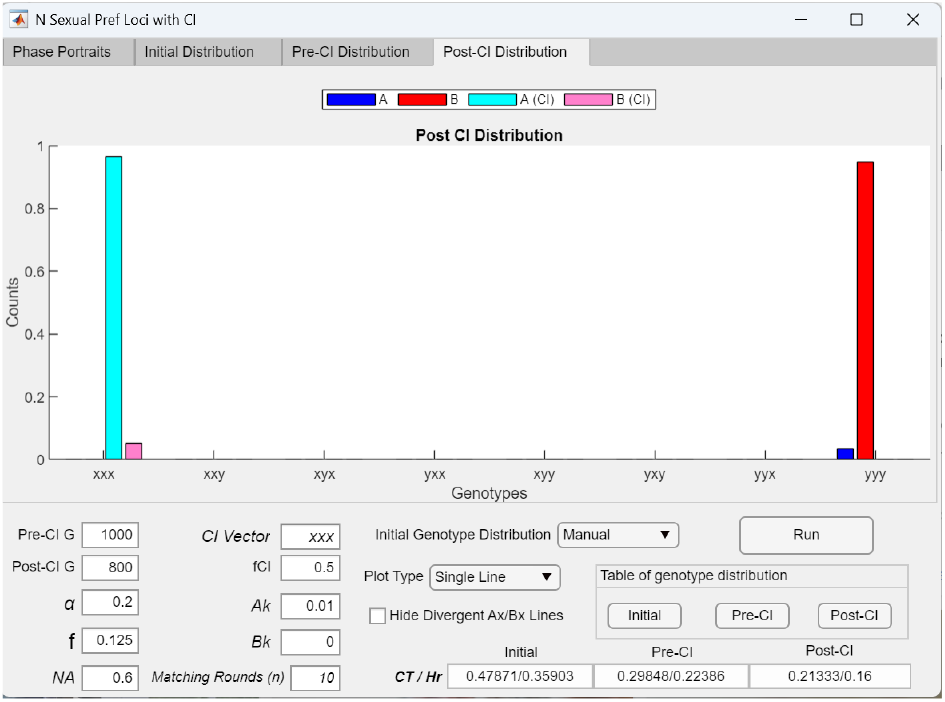
*Post-CI Distribution* bar graph showing the genotype distribution after a CI mutant capturing all locally prevalent mating-bias alleles (*CI* = *xxx*) invades and produces stronger premating RI. When the CI mutant in Fig 42c instead captures all three locally prevalent mating-bias alleles (*CI* = *xxx*), the post-CI genotype distribution shows that opposing mating-bias genotypes persist only at the extreme ends of the distribution, with all intermediate hybrids eliminated. The further reduction in post-CI *CT* and *Hr* values reflects the stronger premating RI achieved by this CI invasion. The time required to reach 95% of the steady-state *Ak* value (0.9646) is 310 generations (*T*_95_ = 310). Compared to the CI that captures only two adaptive mating-bias alleles (*CI* = *xx*0), the CI capturing all three (*CI* = *xxx*) attains a higher steady-state *Ak* value in a shorter time, confirming that CI invasion fitness increases with the number of locally favorable mating-bias alleles captured.

In Fig 42c, when the CI captures two of the three extreme-*X* mating-bias alleles (*CI* = *xx*0), recombination suppression becomes effective, allowing the CI to invade successfully and establish stronger premating RI. The post-CI genotype distribution shown in Fig 42d reveals increased nonrandom assortment of mating-bias alleles across the two niches and a reduction in mating-bias hybrids compared to the distribution in Fig 42b. The CI becomes fixed in niche *A* at mating-bias genotypes closer to the extreme end, with the *Axxx* genotype becoming predominant. Following CI invasion, both post-CI *CT* and *Hr* values decrease relative to their pre-CI values, indicating strengthened premating RI.

Fig 42e shows the post-CI genotype distribution following the invasion of a CI that captures all three extreme-*X* mating-bias alleles (*CI* = *xxx*). Invasion of the CI effectively reduces the number of mating-bias gene loci experienced by the system from three to one and yields a post-CI distribution composed exclusively of extreme genotypes, with no intermediate hybrids. Consequently, the reduction in post-CI *CT* and *Hr* values is maximal.

Compared to the case where the CI captures only two alleles (*CI* = *xx*0), capturing all three alleles (*CI* = *xxx*) further increases the CI’s invasion fitness, allowing it to reach a higher steady-state frequency in niche *A* and to do so more rapidly. This is evidenced by a lower *T*_95_ value, defined as the number of generations required for the CI to reach 95% of its final steady-state frequency. These results demonstrate that a CI capturing a greater number of favorable mating-bias alleles possesses greater invasion fitness.

Figs 43a and 43b show a system with four-locus mating-bias genotypes under parametric conditions that allow system convergence. In Fig 43a, during the pre-CI phase, a small extreme-*X* mutant population (*AX* = 0.01) is able to invade a population composed entirely of extreme-*Y* genotypes, resulting in a fixed-point polymorphism in the *AX*/*BX* phase portrait and the establishment of premating RI. Subsequently, in the post-CI phase, a CI mutant capturing all extreme-*X* mating-bias alleles (*CI* = *xxxx*) is able to invade, shifting the *AX*/*BX* fixed point further toward the lower-right corner of the phase portrait and thereby producing stronger premating RI.

**Fig 43a.**
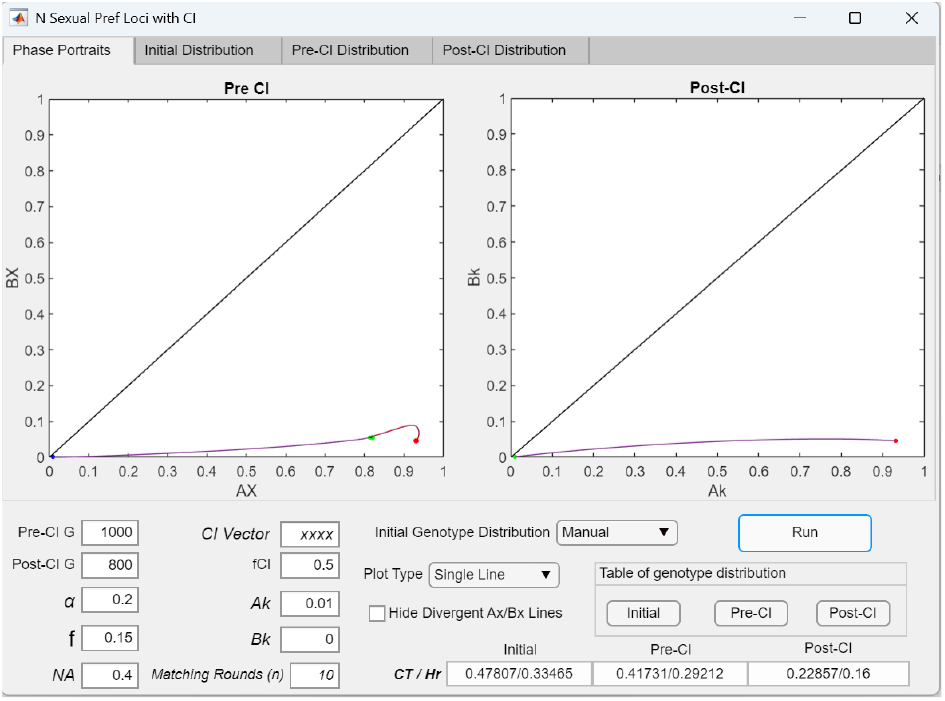
A CI mutant (*CI* = *xxxx*) capturing all locally prevalent mating-bias alleles in a convergent *AX*/*BX* system with four mating-bias loci gains a fitness advantage that allows it to invade and strengthen existing premating RI. Given the parametric values shown, the pre-CI *AX*/*BX* phase portrait is globally convergent with a fixed point at *AX*/*BX* = 0.8126/0.0554. In the post-CI phase, a CI capturing all locally favored mating-bias alleles (*CI* = *xxxx*) acquires an invasion fitness advantage and spreads in the niche-*A* population. This CI invasion strengthens existing premating RI, as reflected by reduced post-CI *CT* and *Hr* values.

**Fig 43b.**
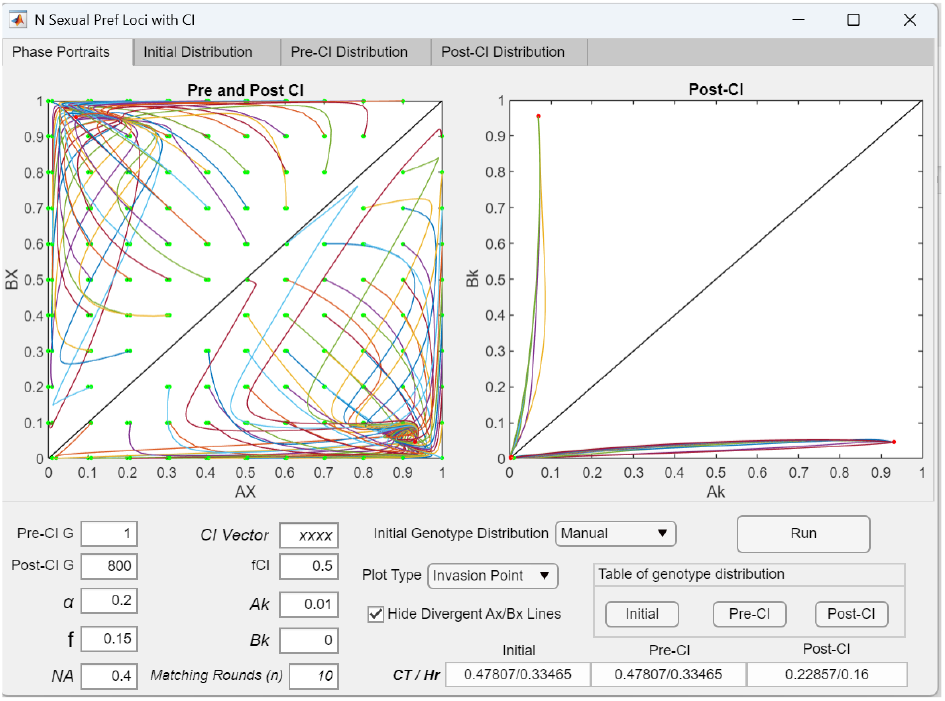
A CI mutant (*CI* = *xxxx*) successfully invades a population under disruptive ecological selection that carries only the *yyyy* mating-bias genotype, resulting in strong premating RI. Using the same parametric values as in Fig 43a but eliminating the pre-CI phase by setting *Pre*-*CI G* = 1, the invading mutant is now a CI containing linked *X* alleles (*CI* = *xxxx*) rather than a genotype of unlinked *X* alleles (*AX* = *Axxxx*), as is the case in the pre-CI phase of Fig 43a. As shown, the CI invasion reproduces the same *AX*/*BX* fixed point and post-CI *CT* and *Hr* values as in Fig 43a, yielding immediate strong premating RI.

**Fig 43c.**
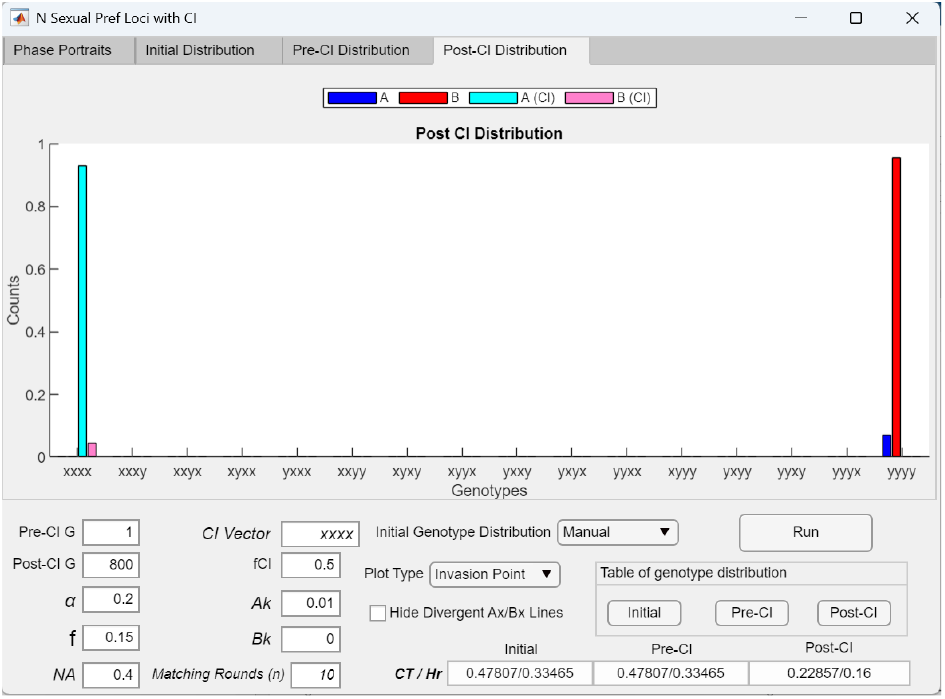
*Post-CI Distribution* bar graph showing the genotype distribution after a CI mutant capturing all locally prevalent mating-bias alleles (CI = *xxxx*) invades and produces stronger premating RI. The bar graph displays the post-CI genotype distribution following the CI invasion shown in Fig 43b. It is identical to the post-CI genotype distribution in Fig 43a. In both cases, invasion of the CI reduces the effective number of mating-bias gene loci to one.

In Fig 43b, the pre-CI phase is omitted by setting *Pre*-*CI G* = 1. Under these conditions, the CI mutant (*CI* = *xxxx, Ak* = 0.01) is able to invade directly into a population consisting solely of the extreme-*Y* genotype. The *AX*/*BX* phase portrait converges to the same fixed-point solution as in Fig 43a and produces the same level of premating RI.

Fig 43c shows the post-CI genotype distribution following CI invasion (*CI* = *xxxx, Ak* = 0.01) in Figs 43a and 43b. In both cases, invasion of the CI reduces the effective number of mating-bias gene loci to one.

In Fig 44a, increasing the value of *α* from 0.2 to 0.25 causes the four-locus system shown in Fig 43a to become divergent. Under these conditions, the pre-CI *AX*/*BX* phase portrait exhibits a globally divergent pattern that prevents invasion by a high-mating-bias mutant (*AX*) with an initial population ratio of *AX* = 0.01.

**Fig 44a.**
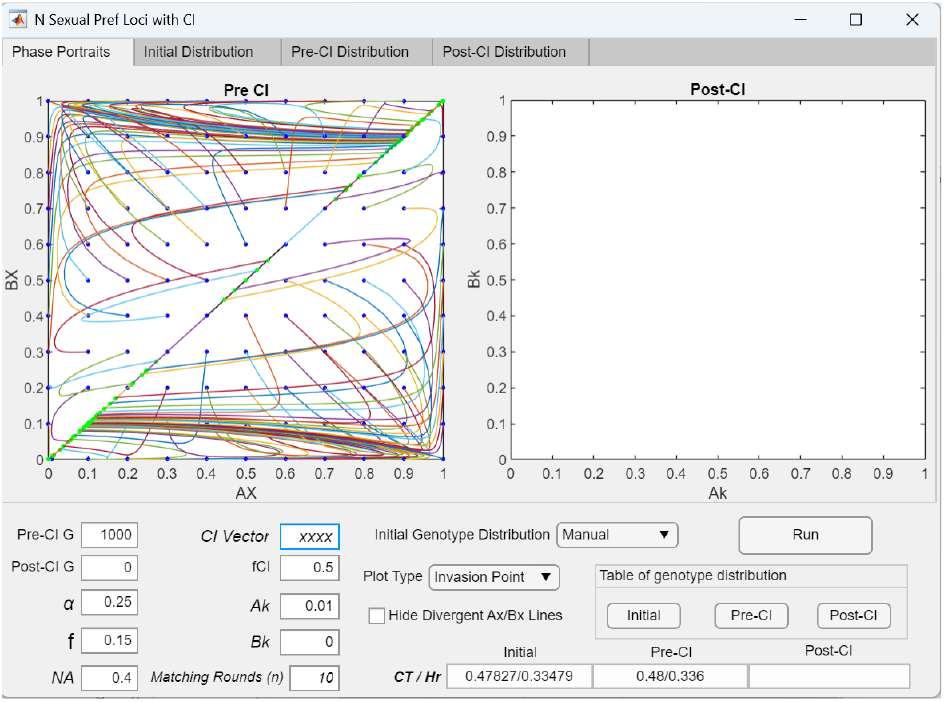
Weakening mating bias by increasing *α* produces a divergent *AX*/*BX* phase portrait in a system with four mating-bias loci. With the parametric values used in Figs 43a and 43b, the pre-CI *AX*/*BX* phase portrait is globally convergent and contains fixed points. However, increasing *α* from 0.2 to 0.25 weakens the mating bias sufficiently to render the *AX*/*BX* phase portrait globally divergent, eliminating all fixed points. This outcome is shown by setting *Post*-*CI G* = 0, which displays only the pre-CI *AX*/*BX* phase portrait and removes the post-CI phase from the simulation.

**Fig 44b.**
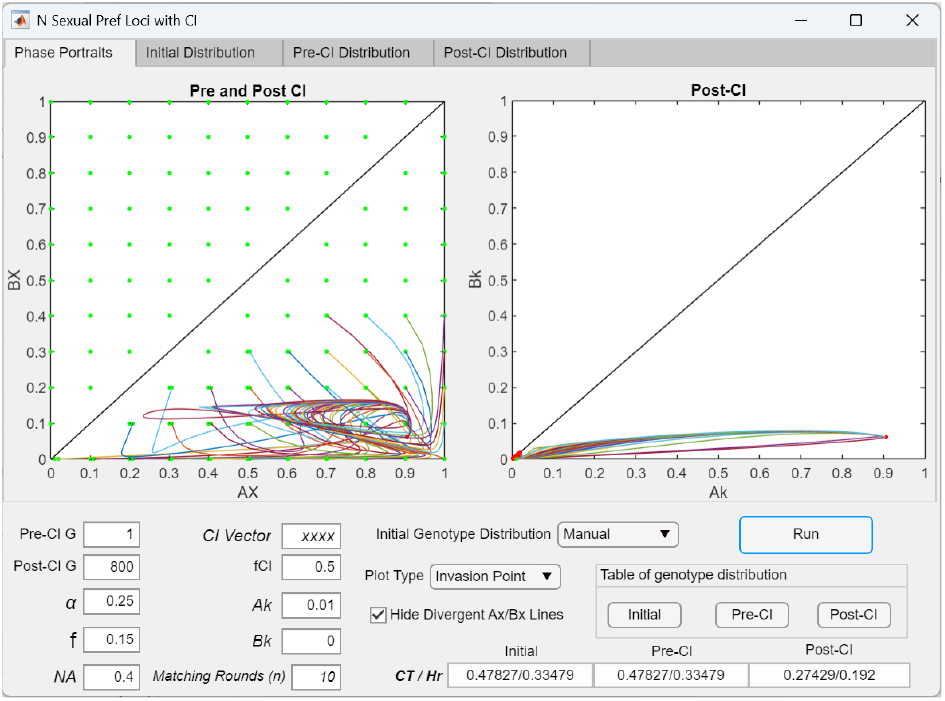
A CI mutant (*CI* = *xxxx*) can invade a divergent system with four mating-bias loci to generate an *AX*/*BX* fixed-point polymorphism and premating RI. In Fig 44a, the pre-CI *AX*/*BX* phase portrait is globally divergent. However, using the same parametric values but bypassing the pre-CI phase by setting *Pre*-*CI G* = 1, the post-CI simulation shows that a CI carrying four linked *X* alleles (*CI* = *xxxx*) can nonetheless invade a population composed entirely of the *yyyy* genotype and establish a fixed-point polymorphism at *AX*/*BX* = 0.9064/0.0628, resulting in reduced post-CI *CT* and *Hr* values. In the application interface, selecting the *Hide Divergent Ax/Bx Lines* option removes vector trajectories that terminate on the diagonal and displays only those that converge to the fixed point. As shown, a wedge-shaped region that includes the origin appears at the lower portion of the *AX*/*BX* phase portrait. This region identifies the range of initial *AX*/*BX* population ratios that permit CI invasion to produce fixed-point convergence and premating RI.

**Fig 44c.**
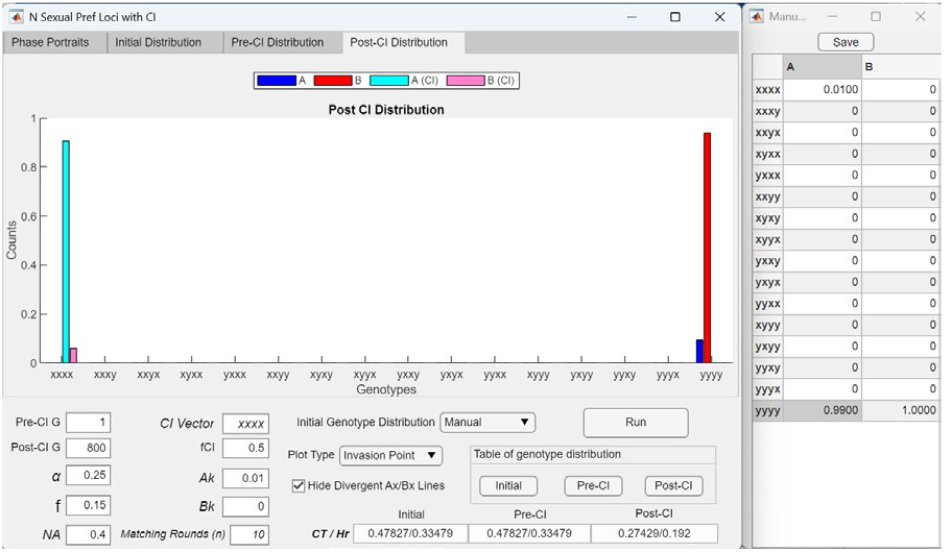
Post-CI genotype distribution showing that invasion of a CI mutant capturing all extreme-*X* alleles (*CI* = *xxxx*) reduces the effective number of mating-bias gene loci from four to one in a system with four-locus mating-bias genotypes, resulting in enhanced RI. The bar graph displays the post-CI genotype distribution from Fig 44b. Following invasion of the locally adaptive CI (*CI* = *xxxx*), the effective number of mating-bias gene loci experienced by the system is reduced from four to one, which facilitates convergence to a fixed point in the *Ax*/*Bx* phase portrait and produces RI. The post-CI genotype distribution resembles that of a system with one mating-bias locus, with no intermediate mating-bias hybrids present.

**Fig 44d.**
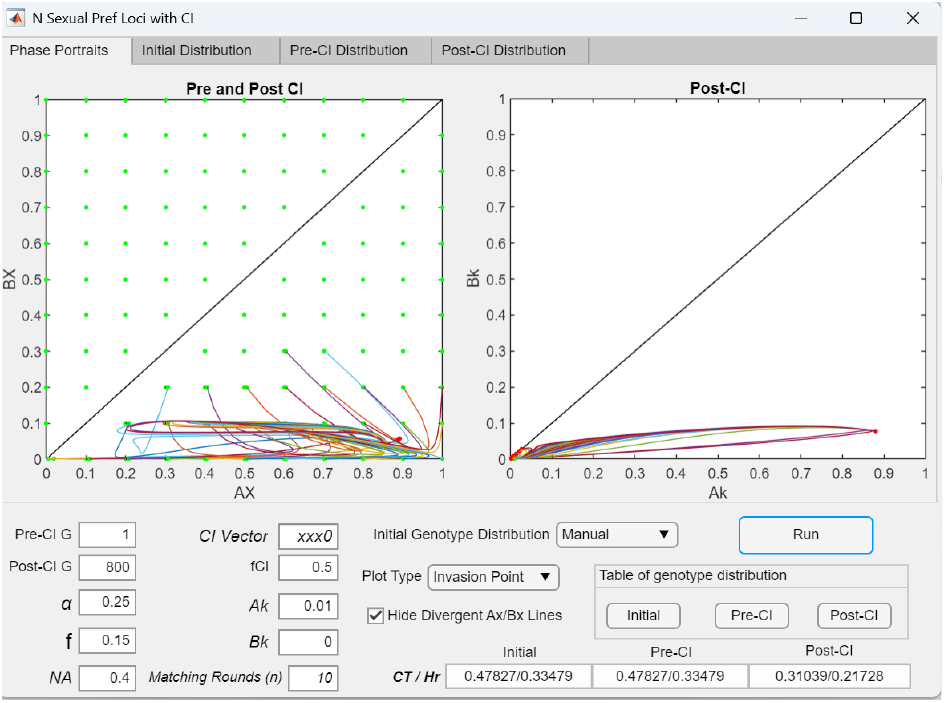
Decreasing the number of extreme-*X* mating-bias alleles captured by an invading CI mutant (*CI* = *xxx*0) reduces the region of initial *AX*/*BX* population ratios that allow invasion. If the CI mutant in Fig 44b captures only three of the four *N*-type mating-bias alleles (*CI* = *xxx*0), the wedge-shaped region at the bottom of the *AX*/*BX* phase portrait that permits invasion becomes further compressed toward the x-axis.

**Fig 44e.**
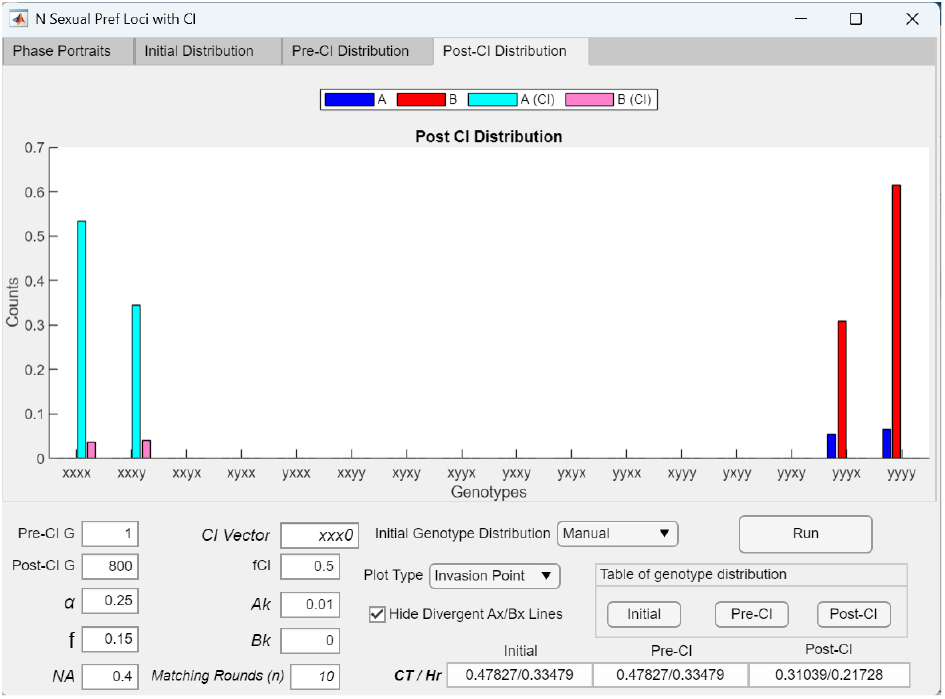
A post-CI genotype distribution showing that invasion of a CI mutant capturing three out of four extreme-*X* alleles (*CI* = *xxx*0) generates intermediate mating-bias hybrids and produces weaker RI. The bar graph displays the post-CI genotype distribution from Fig 44d. Following invasion of the locally adaptive CI (*CI* = *xxx*0), the effective number of mating-bias gene loci experienced by the system is reduced from four to two. Compared to the case where the CI captures all four loci (Fig 44b), capturing fewer loci results in the production of more mating-bias hybrids and weaker RI. The post-CI genotype distribution resembles that of a system with two mating-bias loci.

**Fig 44f.**
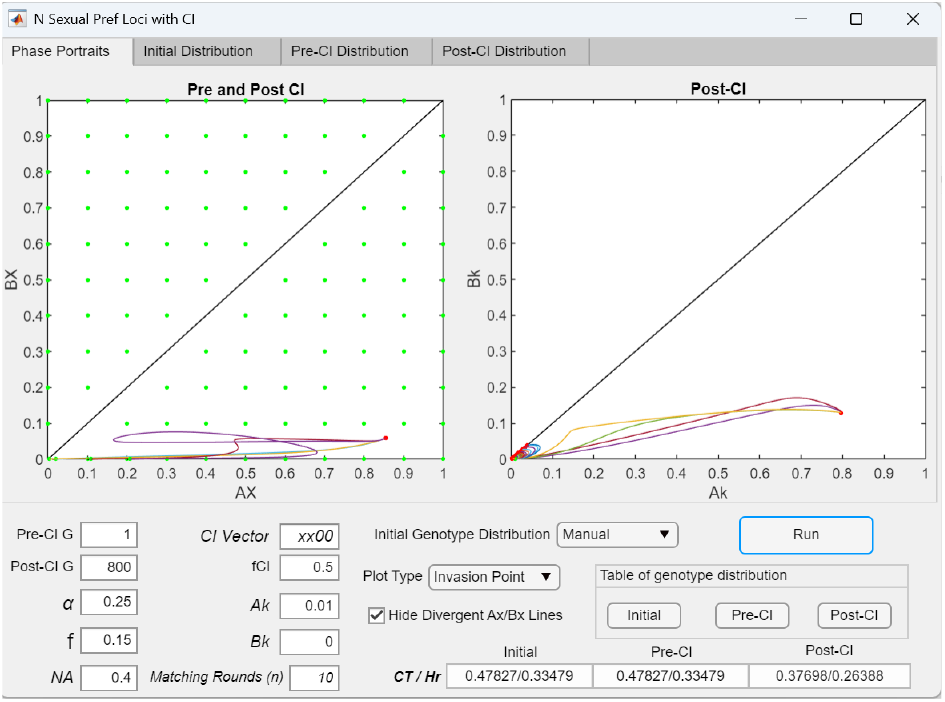
Further reducing the number of extreme-*X* mating-bias alleles captured by a mutant CI (*CI* = *xx*00) restricts the initial *AX*/*BX* population ratios that allow invasion to a narrow band along the *A*-axis. As shown, when the CI captures only two of the four *X*-type alleles (*CI* = *xx*00), successful invasion occurs only for initial *AX* values between 0 and 0.2 along the *x*-axis of the *AX*/*BX* phase portrait. Compared to cases where more *X*-type alleles are captured (Figs 44b and 44c), the higher post-CI *CT* and *Hr* values indicate that weaker premating RI is generated.

**Fig 44g.**
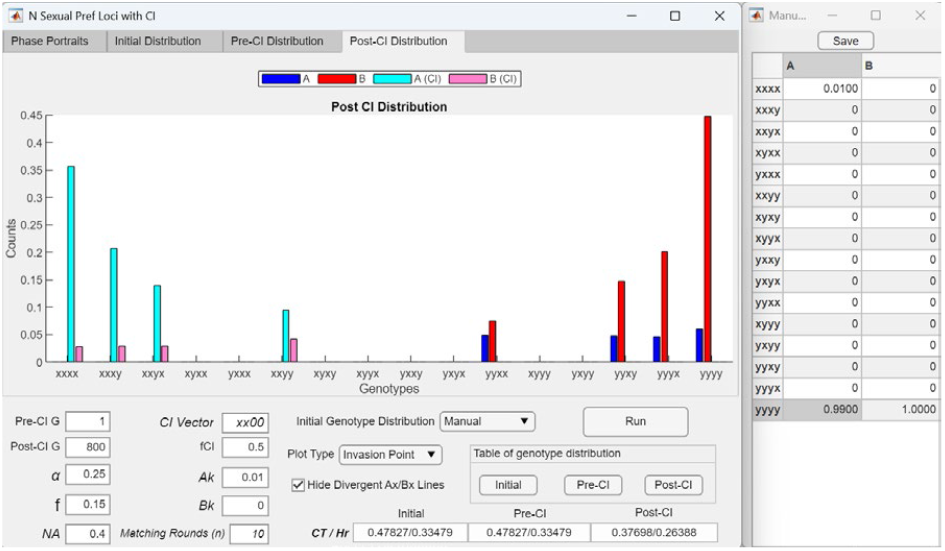
A post-CI genotype distribution showing that invasion of a CI mutant capturing two out of four extreme-*X* alleles (*CI* = *xx*00) generates intermediate mating-bias hybrids and produces weaker RI. The bar graph displays the post-CI genotype distribution from Fig 44f. Following invasion of the locally adaptive CI (*CI* = *xx*00), the effective number of mating-bias gene loci experienced by the system is reduced from four to three. Capturing fewer loci results in the production of more mating-bias hybrids and weaker RI. The post-CI genotype distribution resembles that of a system with three mating-bias loci.

**Fig 44h.**
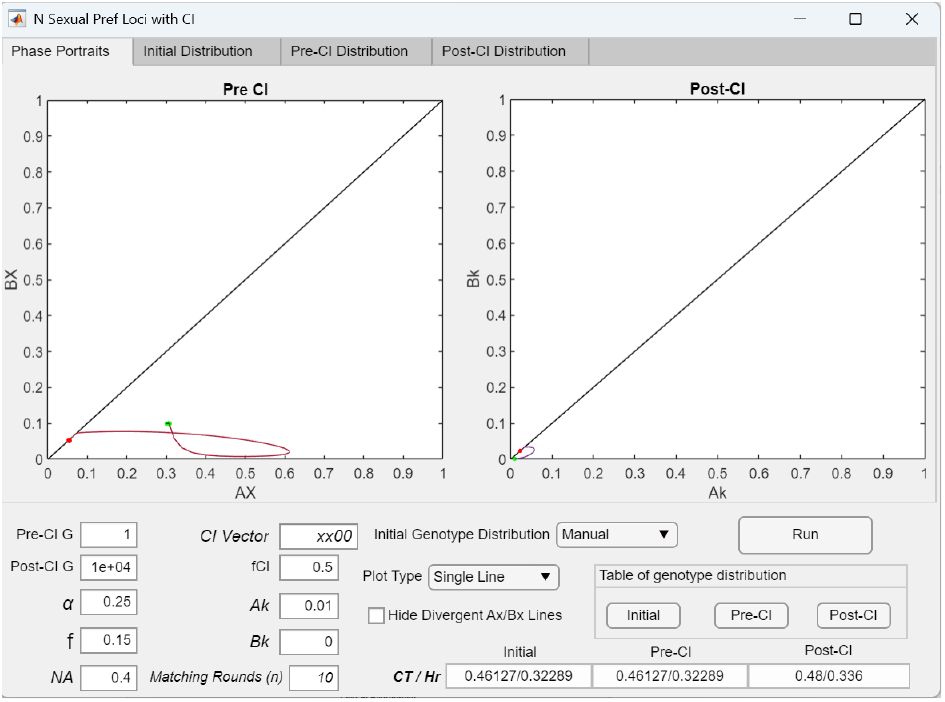
When the initial *AX*/*BX* population ratios fall outside the region in the *AX*/*BX* phase portrait that allows a CI mutant (*CI* = *xx*00) to invade and converge to a fixed point, the CI ultimately fails to invade and all genotypes containing the *x*-type alleles are eliminated. In this example, the initial *AX*/*BX* population ratios are set to be *AX*/*BX* = 0.3/0.1 (see the Initial Genotype Distribution Table in Fig 44i), which lies outside the wedge-shaped region in Fig 44f that permits invasion by a CI mutant (*CI* = *xx*00, *Ak* = 0.01) arising within this initial *AX*/*BX* population. Consequently, the CI fails to invade, and the population vector is driven toward the diagonal line in the phase portrait, eventually collapsing toward the origin.

**Fig 44i.**
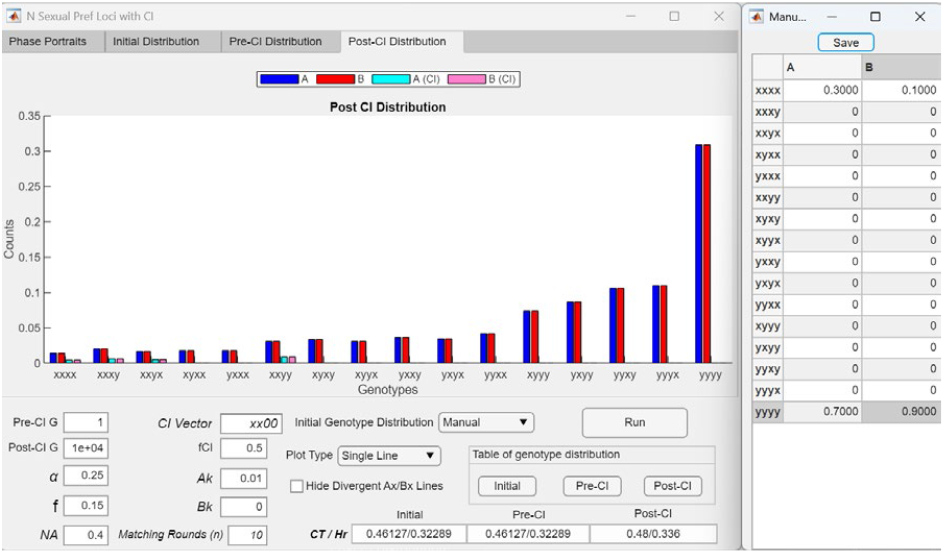
Post-CI genotype distribution graph showing genotypes closer to the extreme-*Y* genotype eliminating those closer to the extreme-*X* genotype. The bar graph displays the post-CI genotype distribution from Fig 44h after 10,000 post-CI generations. The increasingly skewed distribution reflects the progressive elimination of genotypes carrying more *N*-type alleles by those carrying more *x*-type alleles. With the initial *AX*/*BX* population ratios set to 0.3/0.1, as specified in the Initial Genotype Distribution Table on the right, a CI mutant (*CI* = *xx*00) is unable to invade. Eventually, all genotypes containing *x*-type alleles are driven to extinction, and the population becomes fixed for the extreme-*Y* genotype.

Nevertheless, as demonstrated in Fig 44b, a CI mutant capturing all extreme-*X* mating-bias alleles (*CI* = *xxxx, Ak* = 0.01) can still invade this divergent system and establish an *AX*/*BX* fixed-point polymorphism and premating RI. This occurs because the CI is able to reduce the effective number of mating-bias gene loci experienced by the system from four to one, thereby restoring convergence. As a result, the post-CI genotype distribution shown in Fig 44c consists exclusively of the extreme mating-bias genotypes (*xxxx* and *yyyy*), with no intermediate hybrids.

A further observation is that the range of initial *AX*/*BX* population ratios from which the CI mutant (*CI* = *xxxx*) can invade appears to be confined to a wedge-shaped region near the base of the *AX*/*BX* phase portrait. This region can be visualized by selecting the *Hide Divergent Ax/Bx Lines* option in the interface, which removes trajectories that terminate on the diagonal line and displays only the non-divergent portion of the phase portrait.

Figs 44d–44g examine how invasion dynamics change as the CI mutant in Fig 44b captures progressively fewer extreme-*X* mating-bias alleles. In Fig 44d, the CI captures three of four *x*-type alleles (*CI* = *xxx*0), and its invasion reduces the effective number of mating-bias loci experienced by the system from four to two. Accordingly, the post-CI genotype distribution in Fig 44e resembles that of a system with only two mating-bias loci. Similarly, in Fig 44f, when the CI captures two alleles (*CI* = *xx*00), the effective number of loci is reduced from four to three, and the post-CI genotype distribution in Fig 44g contains a total of eight genotypes at the fixed point, consistent with a three-locus system. The phase portraits in Figs 44d and 44f also show that capturing fewer mating-bias alleles causes the wedge-shaped region permitting CI invasion to contract progressively toward the *x*-axis. This region disappears entirely when the CI captures only a single *x* allele (*CI* = *x*000), which offers no fitness advantage for the CI to invade (see Fig 42a).

Figs 44h and 44i show how the phase-portrait dynamics change when the initial *AX*/*BX* population ratios fall outside the wedge-shaped region in Fig 44f that permits invasion by the CI mutant (*CI* = *xx*00). As illustrated by the phase-portrait trajectory in Fig 44h, the higher mating bias of the *AX* population initially allows it to increase during the early post-CI phase. However, once the *AX* ratio exceeds approximately 0.6, the increased production of intermediate hybrids shifts the genotype distribution toward genotypes carrying more *y*-type alleles, which begin to displace those carrying more *x*-type alleles. As a result, the *AX*/*BX* population vector turns toward the diagonal line in the phase portrait and gradually converges toward the origin. Over time, the post-CI genotype distribution shown in Fig 44i becomes increasingly skewed in favor of genotypes containing more *y*-type alleles. Ultimately, all genotypes carrying *x*-type alleles are eliminated, and the population becomes fixed for the extreme-*Y* genotype.

So far, Figs 44a–44i examine the invasion of a mutant CI capturing mating-bias alleles arising from a small extreme-*X* population (*AX* = 0.01) near the origin and invading a divergent system composed entirely of extreme-*Y* genotypes. In Figs 45a–45d, the CI instead originates directly from the origin (*AX*/ *BX* = 0/0). Because no *x*-type alleles exist at the uncaptured loci of the CI vector, those positions are automatically occupied by *y*-type alleles. As a result, only two mating-bias genotypes are possible in the system, and CI invasion invariably reduces the effective number of mating-bias loci to one.

**Fig 45a.**
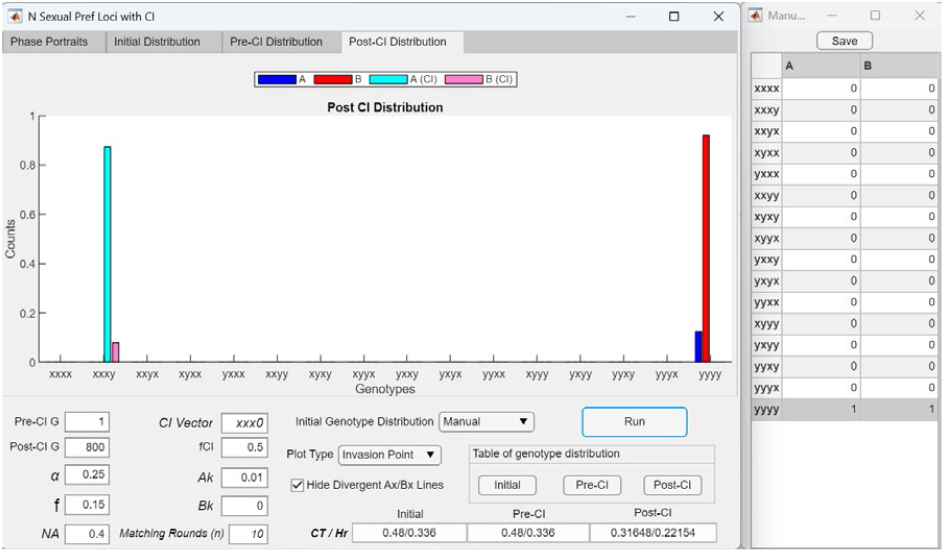
Invasion of a CI mutant (CI = *xxx*0) arising from the origin reduces the number of mating-bias loci experienced by the system from four to one. Unlike the case illustrated in Figs 44d and 44e, where the CI mutant arises from a small extreme-*X* population near the origin (*Axxxx* = 0.01), here the CI mutant (*CI* = *xxx*0) originates directly from the origin of the phase portrait, where *AX*/*BX* = 0/0 (see the Initial Genotype Distribution Table on the right), and is therefore equivalent to a CI mutant of the form *C* = *Axxxy*. Consequently, the post-CI genotype distribution resembles that of a system with one mating-bias locus, consisting of the extreme genotypes *xxxy* and *yyyy* with no intermediate hybrids. However, because the mating bias between genotypes *xxxy* and *yyyy* is weaker than the maximum mating bias (*α* = 0.25) between the extreme-*X* (*xxxx*) and extreme-*Y* (*yyyy*) genotypes, the resulting post-CI *CT* and *Hr* values are higher than those shown in Figs 44c and 44e, indicating weaker premating RI.

**Fig 45b.**
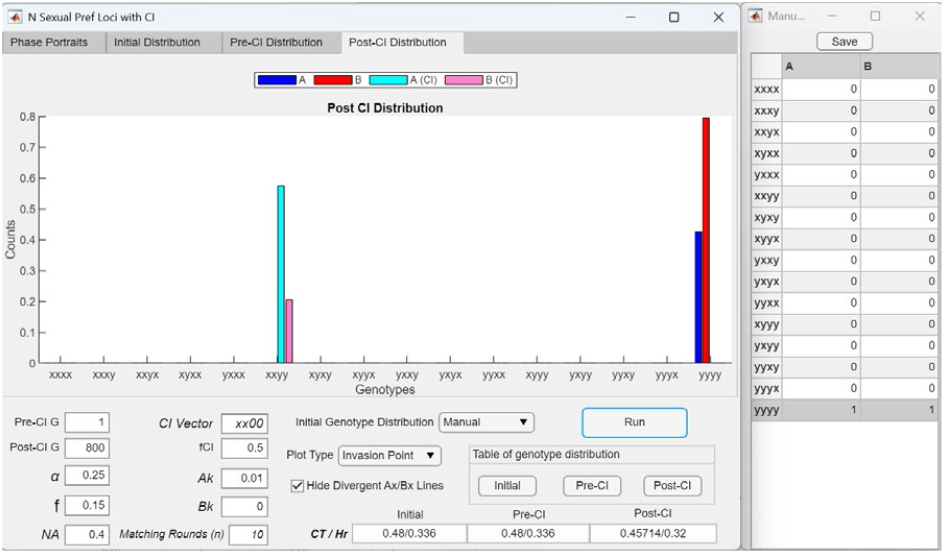
Invasion of a CI mutant (*CI* = *xx*00) arising from the origin reduces the number of mating-bias loci experienced by the system from four to one. The bar graph shows the post-CI genotype distribution following invasion by a CI mutant (*Axx*00) that originates at the origin of the *AX*/*BX* phase portrait and is equivalent to *CI* = *Axxyy*. The post-CI distribution consists only of the genotypes *xxyy* and *yyyy*, with no intermediate mating-bias hybrids. Thus, invasion of the CI reduces the effective number of mating-bias loci experienced by the system from four to one. However, because the mating bias between *xxyy* and *yyyy* is weaker than the maximum mating bias observed between the extreme genotypes (*xxxx* and *yyyy*), the resulting premating RI is correspondingly weaker.

**Fig 45c.**
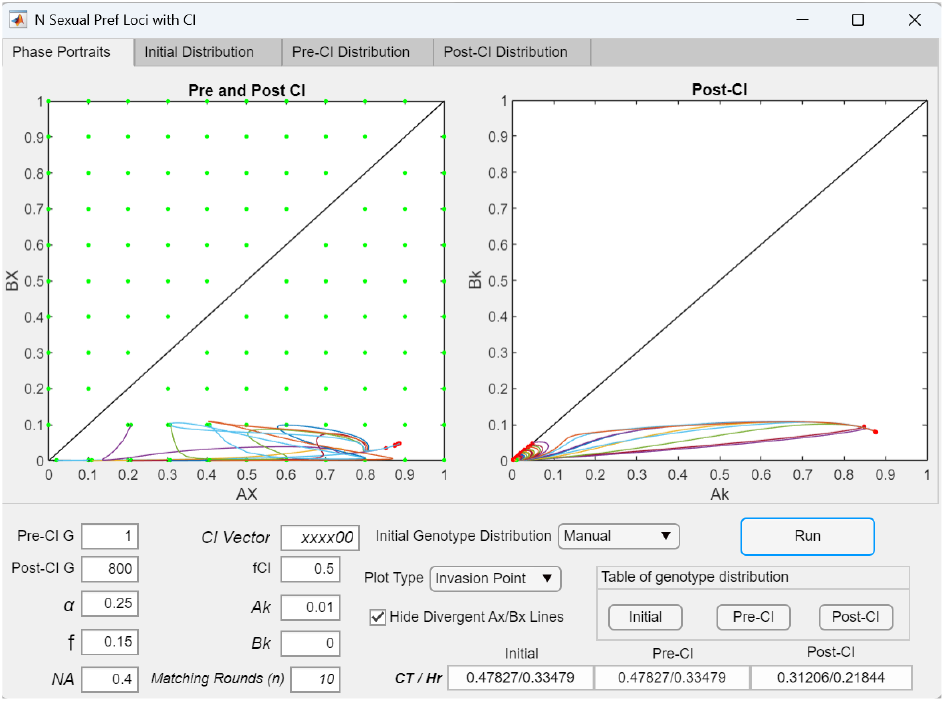
Invasion of a CI mutant (*CI* = *xxxx*00) arising from the origin and invading a system composed solely of six-locus extreme-*Y* genotypes. A CI mutant (*CI* = *xxxx*00) emerging from the origin of the *AX*/*BX* phase portrait is able to invade a system with six mating-bias loci that is composed entirely of extreme-*Y* genotypes. Although systems with such a large number of mating-bias loci are typically divergent, this CI can still invade and establish premating RI. The *AX*/*BX* phase portrait also delineates the region near the *x*-axis that permits invasion by the CI.

**Fig 45d.**
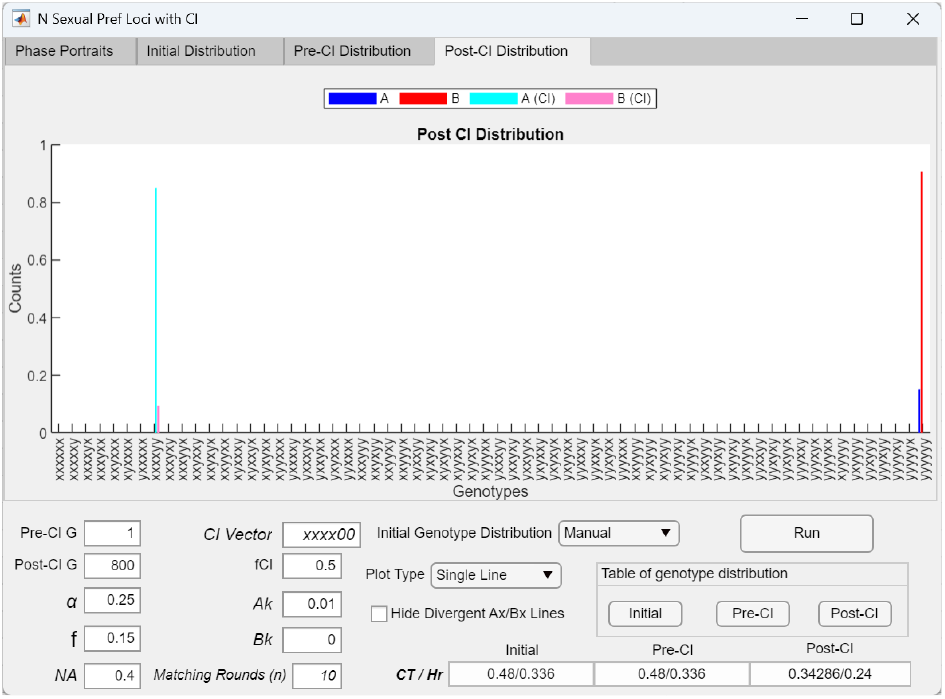
Post-CI genotype distribution graph showing that invasion by a CI mutant (*CI* = *Axxxx*00) arising from the origin reduces the effective number of mating-bias loci experienced by the system from six to one. The bar graph displays the post-CI genotype distribution from Fig 45c, where only the genotypes *xxxxyy* and *yyyyyy* remain at the fixed point. The enhanced invasion fitness of the CI likely results from its ability to reduce the effective number of mating-bias loci experienced by a multilocus system to one.

**Fig 45e.**
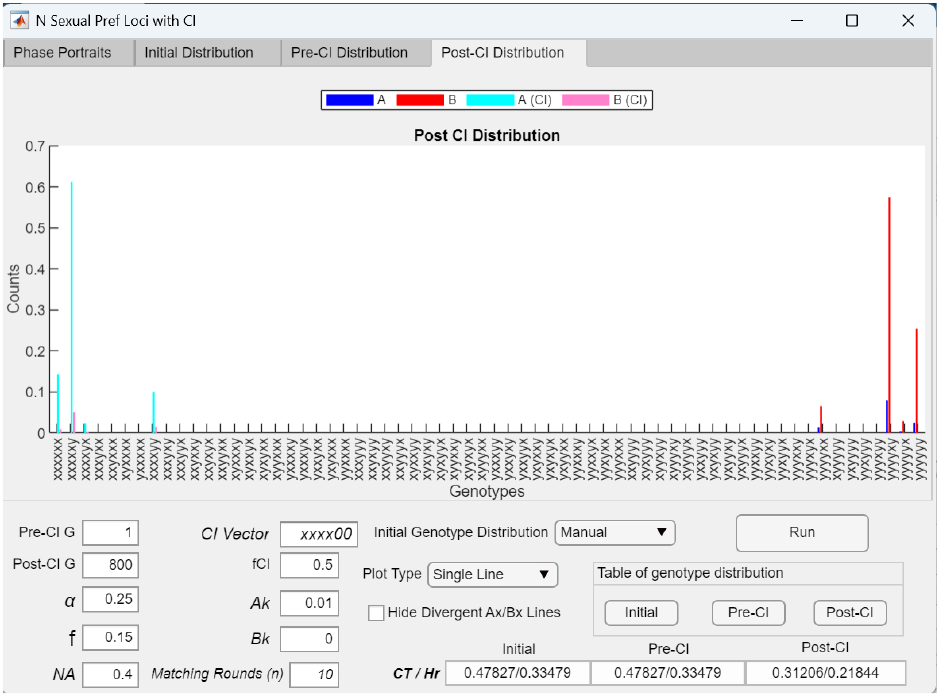
Post-CI genotype distribution graph showing that invasion by a CI mutant (*CI* = *Axxxx*00) arising from an initial extreme-*XX* population (*Axxxxxx* = 0.01) reduces the effective number of mating-bias loci experienced by the system from six to three. The bar graph shows the post-CI genotype distribution when a CI mutant (CI = *Axxxx*00) arises from an initial population of *Axxxxxx* = 0.01, rather than from the origin (*AX*/*BX* = 0/0) as in Fig 45d. The resulting post-CI distribution contains eight mating-bias genotypes, which is equivalent to a system with three mating-bias loci. These eight genotypes include all possible permutations of *NN*- and *NN*-type alleles at the uncaptured mating-bias loci in the CI vector (i.e., the “0” positions).

In Fig 45a, invasion of a CI mutant (*CI* = *xxx*0) arising from the origin into a system with four mating-bias loci and initially fixed for the extreme-*Y* genotype produces a post-CI distribution containing only *xxxy* and *yyyy* genotypes, with no intermediate hybrids between them. This outcome is equivalent to that of a system with a single mating-bias locus. A similar outcome occurs in Fig 45b, where the CI mutant (*CI* = *xx*00) is functionally equivalent to *CI* = *xxyy*. Again, only two genotypes (*xxyy* and *yyyy*) remain after invasion, with no intermediate hybrids, and the effective number of mating-bias loci is reduced to one. In both cases (Figs 45a and 45b), reducing the effective number of loci to one eliminates intermediate hybrid formation and promotes CI invasion and convergence.

However, the premating RI generated by such invasions tends to be weaker compared to cases where the CI captures all extreme-*X* alleles. This is because the mating bias between the post-CI hybrid genotype and the extreme-*Y* genotype (e.g., *xxxy* vs *yyyy* or *xxyy* vs *yyyy*) is lower than *α*, the maximum mating bias between the extreme-*X* and extreme-*Y* genotypes (*xxxx* and *yyyy*). Therefore, CI mutants that capture only a partial set of extreme-*X* alleles generate weaker RI than CI mutants that capture all extreme-*X* alleles.

The examples in Figs 45c and 45d show that a CI mutant (*CI* =*xxxx*00) arising from the origin can successfully invade a divergent system with six mating-bias loci and establish an *AX*/*BX* fixed-point polymorphism and premating RI. Ordinarily, systems with such a high number of mating-bias loci are highly resistant to invasion by alternative high-mating-bias genotypes without the invasion advantage conferred by a CI. As shown in Fig 45c, a narrow region near the *x*-axis in the *AX*/*BX* phase portrait permits invasion by the CI arising from the origin. Following invasion, the post-CI genotype distribution in Fig 45d indicates that the effective number of mating-bias loci experienced by the system is reduced from six to one. However, because the mating bias between the CI genotype and the extreme-*Y* genotype depends on the number of *x*-type alleles captured by the CI, the resulting premating RI is weaker when fewer *x*-type alleles are captured. The greater the number of *N*-type alleles captured by the CI, the stronger the effective mating bias and the resulting RI.

For comparison, Fig 45e shows the post-CI genotype distribution of the same CI mutant (*CI* = *Axxxx*00) when it arises from a small initial extreme-*X* population (*Axxxxxx* = 0.01), rather than from the origin. In this case, the effective number of mating-bias loci is reduced from six to three instead of one. As demonstrated by the presence of a restricted invasion zone in the *AX*/*BX* phase portrait in Fig 45c and by the differing post-CI distribution in Fig 45e, the final invasion outcome of a CI mutant in a divergent system is highly sensitive to the number, location, and initial frequency of *x*-type alleles accompanying the CI at its inception.

### III. Systems with multilocus ecological genotypes and single-locus hybrid-incompatibility genotypes

We first examine how population dynamics in the one-locus-equivalent model of hybrid incompatibility depend on the viability ratio (*v*) and the strength of ecological selection (*f*) in the absence of premating barriers (*α* = 1). We then introduce premating barriers to assess how mating bias (*α* < 1) modifies an established postzygotic barrier.

We next adapt the MultiEcoCI application to examine how incorporating multiple ecological loci and viable hybrid populations alters system dynamics. This is done by replacing the mating-bias parameter (*α*) with the viability ratio (*v*) and fixing the number of matching rounds (*n*) to one, as described in the Methodology section. Pre-CI dynamics are examined by setting the number of post-CI generations to zero. Subsequently, the post-CI phase is enabled and CI mutations capturing different combinations of ecological alleles are introduced to evaluate their invasion dynamics and effects on system behavior.

Finally, we use the MultiCompCI application to investigate how having multiple loci underlying hybrid incompatibility affects system dynamics and how the invasion of CIs capturing various extreme-type incompatibility alleles shapes the evolutionary outcomes of the system.

#### 1 Interaction dynamics of ecological, mating-bias, and hybrid-incompatibility alleles in single-locus-equivalent models

By replacing *α* with *v*, mating-bias alleles *X* and *Y* with incompatibility alleles *P* and *Q*, and setting *n* = 1, the GUI program in Fig 7—originally used to simulate the population dynamics of the mathematical model in Fig 3—can also be used to compute the population dynamics of a one-locus-equivalent model of hybrid incompatibility in the absence of premating barriers (see Methodology).

As shown in Fig 46a, when *f* = 0.5, disruptive ecological selection is absent and all individuals encounter one another randomly and mate freely in a single panmictic population. Under these conditions, the *Ap*/*Bp* phase portrait is divergent for all values of *v*. Provided that *v* < 1, the more prevalent hybrid incompatibility allele (*P* or *Q*) eliminates the less prevalent allele in the population through attritive interactions. In Fig 46b, when *f* < 0.5, weak disruptive ecological selection (i.e., relatively large *f*) combined with weak incompatibility selection (large *v*) still produces a divergent *Ap*/*Bp* phase portrait.

**Fig 46a.**
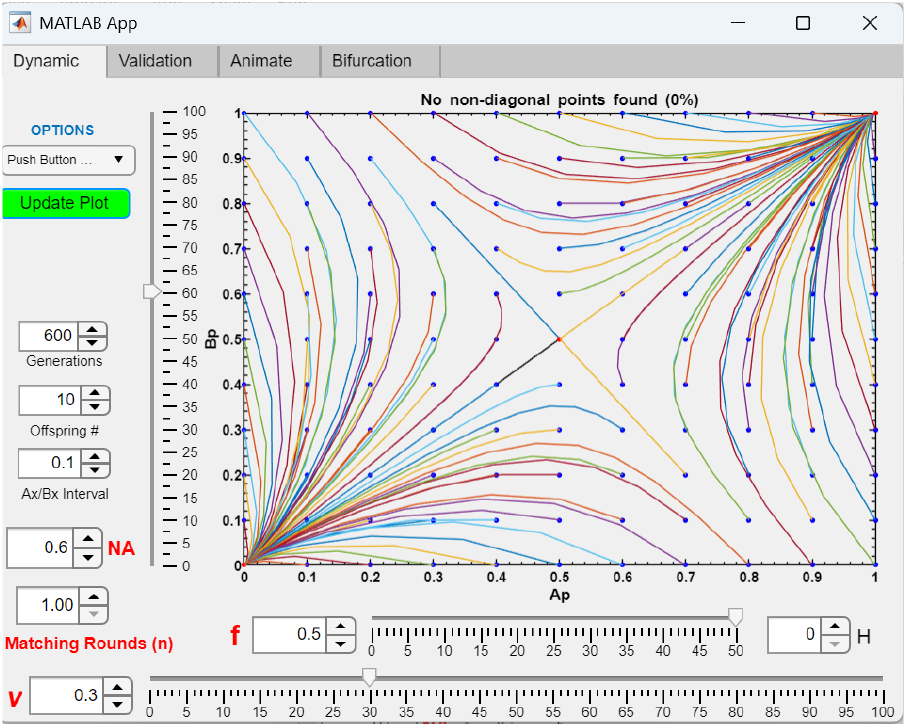
In a two-niche two-allele model of postzygotic hybrid incompatibility, no system convergence is possible in the absence of disruptive ecological selection. Without disruptive ecological selection (*f* = 0.5) and premating barriers (*α* = 1), no value of *v* can create fixed-point convergence in the *Ap*/*Bp* phase portrait. In the panmictic population, the more numerous *P* or *Q* allele eventually eliminates the less numerous, incompatible allele.

**Fig 46b.**
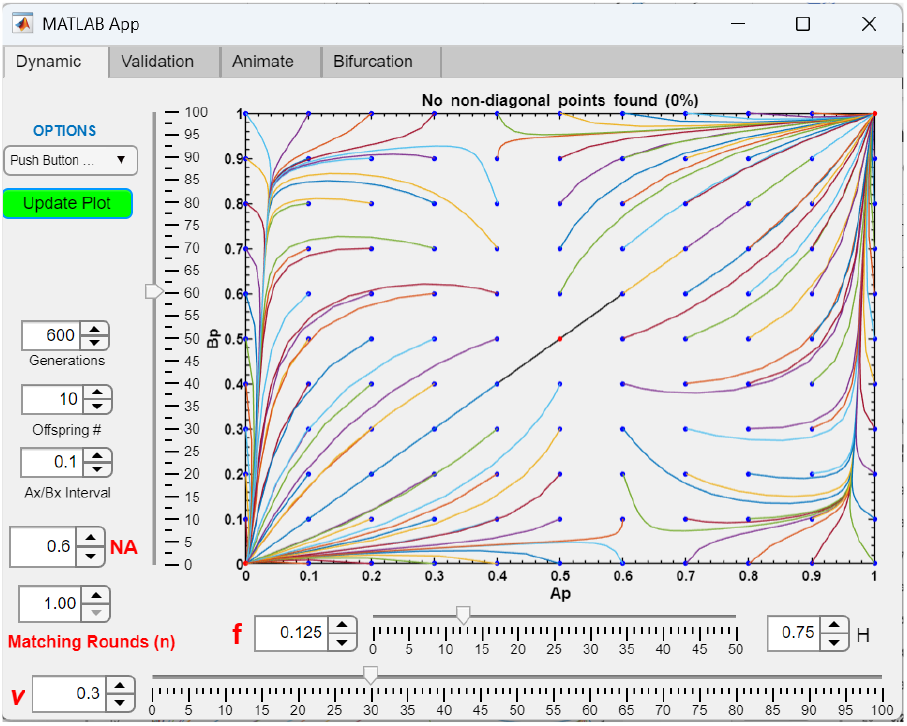
When the strength of disruptive ecological selection or hybrid incompatibility is weak, the system remains divergent. When *f* = 0.125, *v* = 0.3, no fixed-point convergence occurs in the *Ap*/*Bp* phase portrait.

**Fig 46c.**
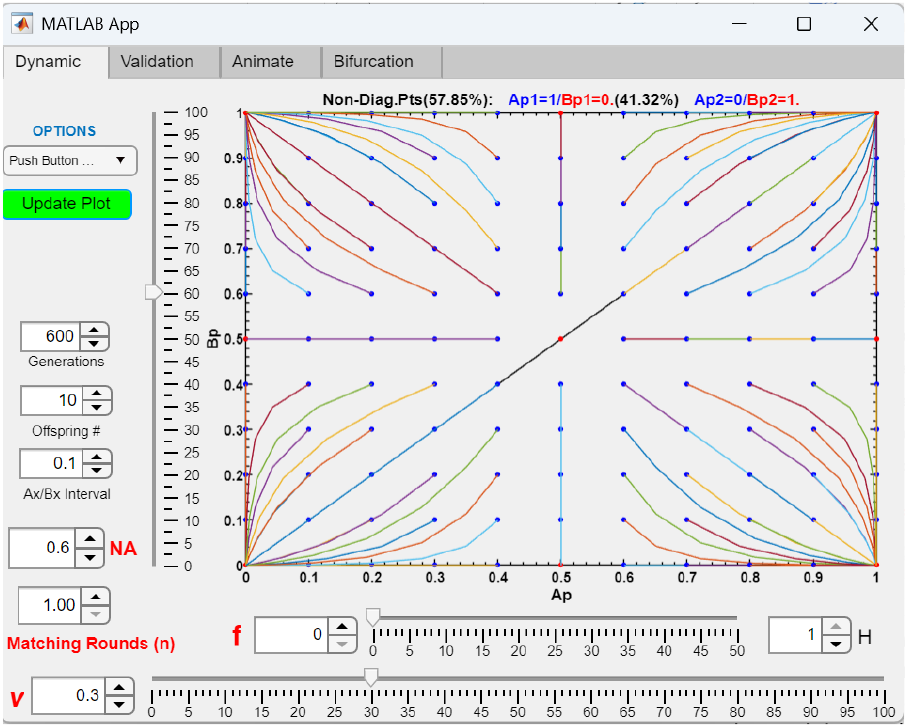
Complete disruptive ecological selection causes the more prevalent hybrid incompatibility allele in each niche to eliminate the less prevalent incompatible allele. When disruptive ecological selection is complete (*f* = 0), inter-niche mating produces no viable offspring, effectively isolating the two niche ecotypes. Under these conditions, the more prevalent incompatibility allele (*P* or *Q*) in each niche eliminates the less frequent incompatible allele. This outcome occurs for all values of *v* in the range 0 ≤ *v* < 1.

**Fig 46d.**
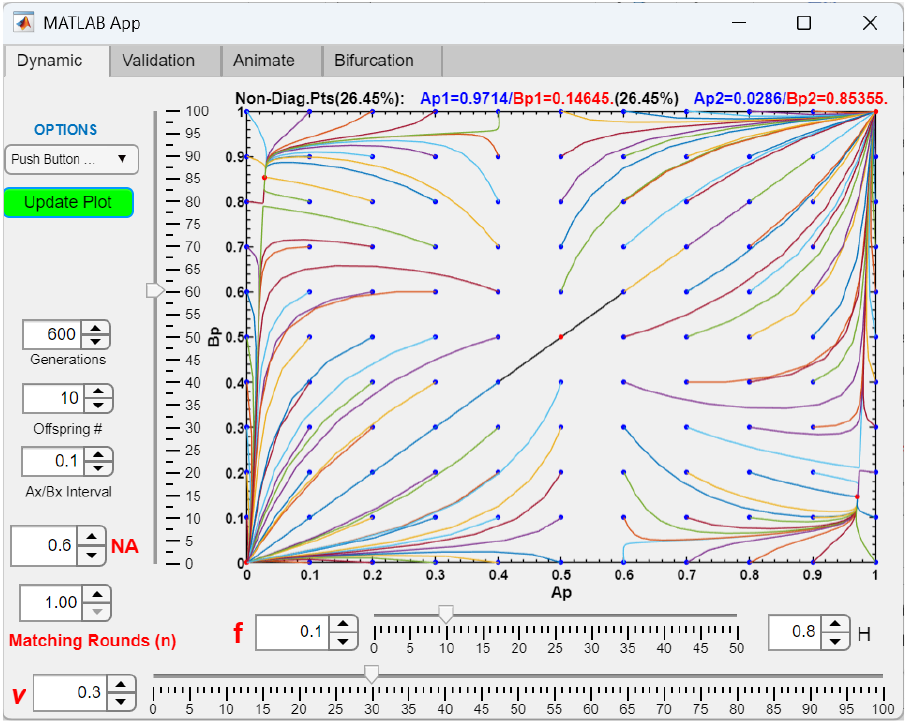
When disruptive ecological selection is sufficiently strong, a fixed point emerges in the lower-right quadrant of the *Ap*/*Bp* phase portrait. In Fig 46b, strengthening disruptive ecological selection by reducing *f* to 0.1 produces the fixed point in the lower-right quadrant of the phase portrait. For *v* = 0.3, such a fixed point appears for *f* ≤ 0.1, marking the transition between system divergence (Fig 46b) and complete isolation (Fig 46c, when *f* = 0). Lower values of *f* move the fixed point closer to the lower-right corner.

**Fig 46e.**
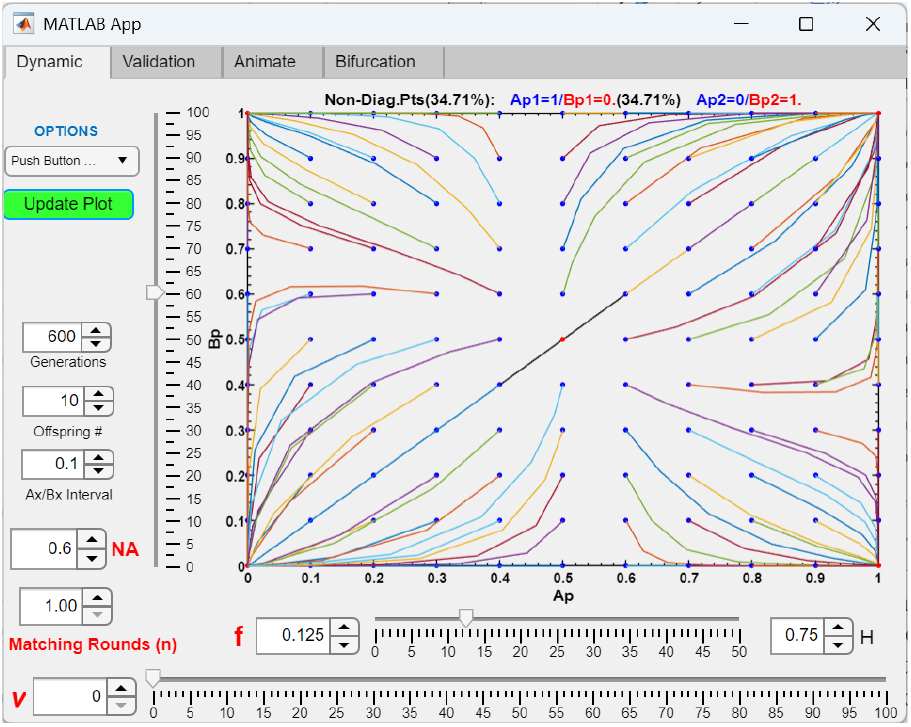
When hybrid incompatibility is complete, the *Ap*/*Bp* phase portrait resembles that observed under complete isolation between niches. When *v* = 0, such that no viable hybrids are produced due to complete intrinsic incompatibility, the resulting *Ap*/*Bp* phase portrait resembles that of complete isolation between niche ecotypes (Fig 46c). Consequently, in each quadrant of the phase portrait, the more prevalent incompatibility allele (*P* or *Q*) in each niche eliminates the less frequent incompatible allele.

**Fig 46f.**
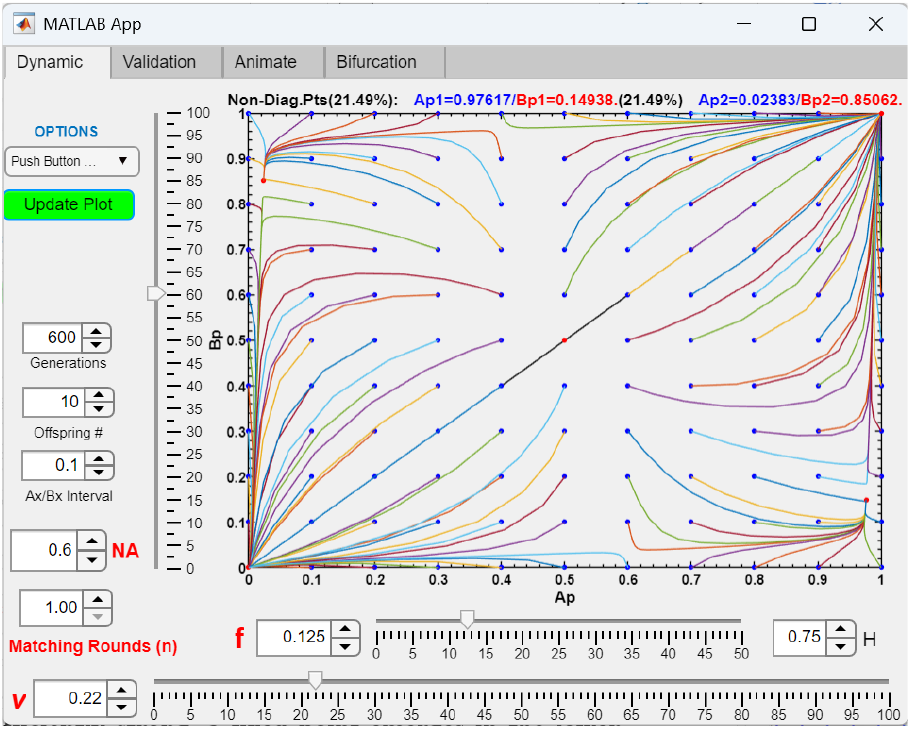
When hybrid incompatibility selection is sufficiently strong, a fixed point emerges in the lower-right quadrant of the *Ap*/*Bp* phase portrait. In Fig 46b, strengthening hybrid incompatibility selection by reducing *v* to 0.22 produces the fixed point in the lower-right quadrant of the phase portrait. For *f* = 0.125, the fixed point appears for *v* ≤ 0.22, marking the transition between system divergence (Fig 46b) and complete hybrid inviability (Fig 46e, when *v* = 0). As *v* decreases further, the fixed point shifts toward the lower-right corner.

**Fig 46g.**
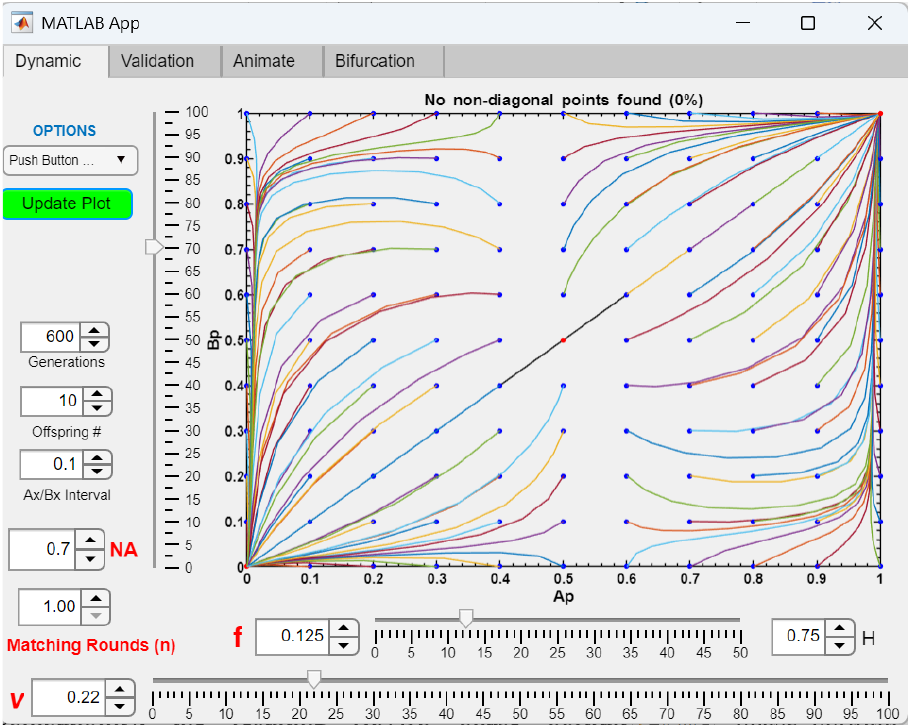
A symmetric range of *NA* values around *NA* = 0.5 permits the emergence of a fixed point in the lower-right quadrant of the *Ap*/*Bp* phase portrait. In Fig 46f, increasing *NA* to 0.7 eliminates the fixed point in the *Ap*/*Bp* phase portrait, causing the system to become divergent. Keeping all other parameter values the same, fixed-point convergence disappears when *NA* lies outside the range 0.3 < *NA* < 0.7.

In contrast, in Fig 46c, when disruptive ecological selection is complete (*f* = 0), no viable offspring (hybrid or parental) are produced from inter-niche matings, and no gene flow occurs between niche populations. Consequently, the more prevalent incompatibility allele eliminates the less prevalent allele within each niche for all values of *v* in the range 0 ≤ *v* < 1.

During the transition from the divergent phase portrait in Fig 46b (*f* = 0.125) to the complete isolation phase portrait in Fig 46c (*f* = 0), a fixed point appears in the lower-right quadrant of the Ap/Bp phase portrait (Fig 46d) once disruptive ecological selection becomes sufficiently strong (*f* ≤ 0.1).

Similarly, when hybrid incompatibility is complete (*v* = 0), no viable incompatible hybrid offspring are produced. The resulting *Ap*/*Bp* phase portrait shown in Fig 46e resembles that in Fig 46c. (The asymmetry in the phase portrait arises from a nonzero value of *f*, which reduces offspring from inter-niche matings between individuals carrying the same incompatibility allele, and from unequal niche sizes that determine the relative frequency of inter-niche matings.) Because incompatibility alleles diverge across niches, gene flow between niche ecotypes is greatly reduced, as only offspring from inter-niche matings that recreate the parental *P* or *Q* genotypes can survive. As a result, most offspring arise from intra-niche matings rather than inter-niche matings, allowing the more prevalent incompatibility allele to eliminate the less prevalent allele within each niche.

During the transition from the divergent *Ap*/*Bp* phase portrait in Fig 46b (*v* = 0.3) to that in Fig 46e (*v* = 0), an intermediate but sufficiently low value of *v* (*v* ≤ 0.22) produces a fixed point in the lower-right quadrant of the *Ap*/*Bp* phase portrait (Fig 46f).

Together, disruptive ecological selection (parameter *f*) and hybrid incompatibility selection (parameter *v*) act complementarily to promote fixed-point convergence in the *Ap*/*Bp* phase portrait. Weak disruptive ecological selection can be compensated by strong hybrid incompatibility selection, and vice versa. Furthermore, stronger disruptive ecological selection (lower *f*) or stronger hybrid incompatibility selection (lower *v*) shifts an existing *Ap*/*Bp* fixed point closer to the lower-right corner of the phase portrait, resulting in stronger postzygotic RI between niche ecotypes.

The effects of varying niche size (*NA*) on system convergence are illustrated in Fig 46g. There appears to be a symmetric range of *NA* values around *NA* = 0.5 that permits the emergence of *Ap*/*Bp* fixed points (0.3 < *NA* < 0.7 in the Fig 46g example). Outside this range, *Ap*/*Bp* fixed-point convergence is not observed.

Next, we used a previously developed GUI program to investigate how premating barriers mediated by mating biases reinforce postzygotic barriers established through the formation of a stable fixed point in the lower-right quadrant of the *Ap*/*Bp* phase portrait.

Fig 47a shows that, in the absence of premating barriers, a stable fixed point is produced in the *Ap*/*Bp* phase portrait when *f* = 0.125 and *v* = 0.22. Similarly, Fig 47b shows that, in the absence of hybrid incompatibility, fixed-point convergence in the *Ax*/ *Bx* phase portrait—and thus premating RI generated by the mating-bias barrier—occurs when *f* = 0.125 and *α* = 0.3.

**Fig 47a.**
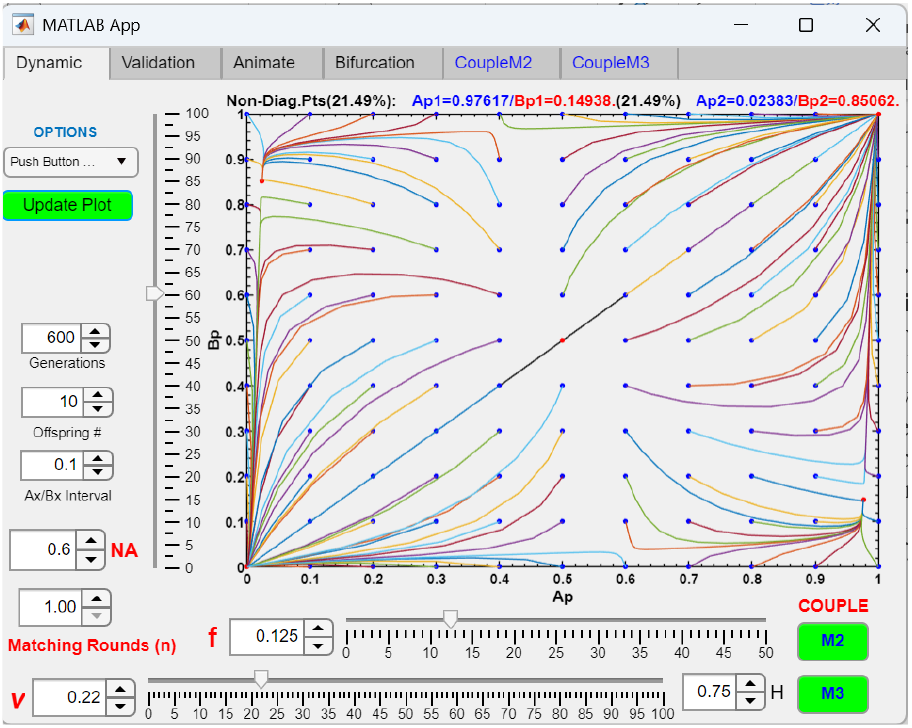
In the absence of premating barriers, a stable fixed point appears in the lower-right quadrant of the *Ap*/*Bp* phase portrait when the parameters *f* and *v* are sufficiently low. A GUI application was developed to simulate the coupling of a two-allele mating-bias model with a two-allele hybrid incompatibility model. As shown, when premating mating-bias barriers are absent, a fixed point occurs at *Ap*/*Bp* = 0.97617/0.14938 for *v* = 0.22 and *f* = 0.125. The resulting Ap/Bp phase portrait is identical to that shown in Fig 46f.

**Fig 47b.**
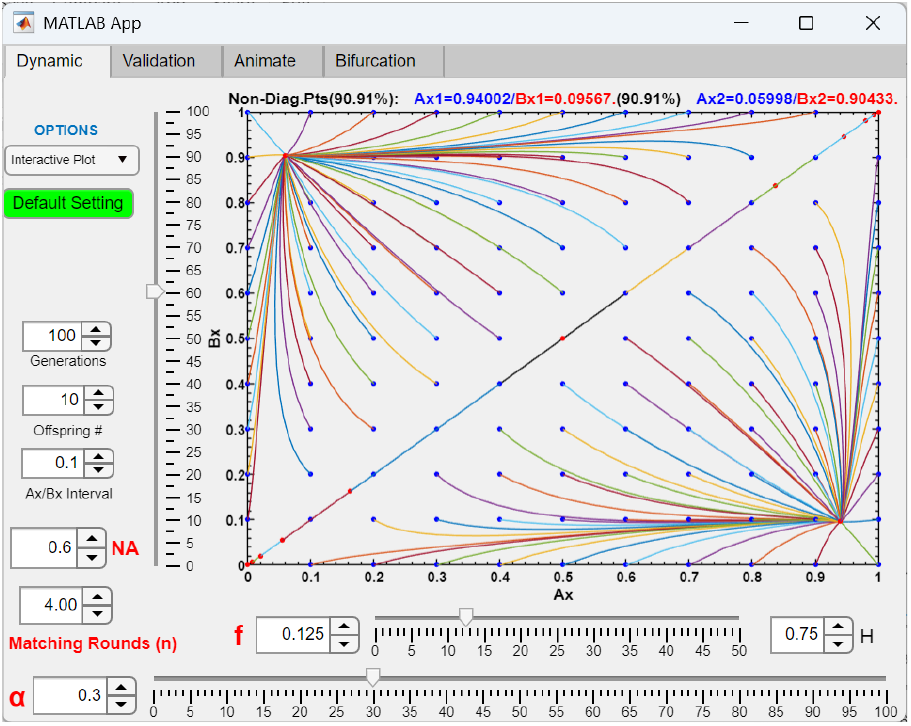
In the absence of hybrid incompatibility barriers, stable fixed points appear in the *Ax*/*Bx* phase portrait of the mating-bias barrier when the parameters *f* and *α* are sufficiently low. Under these conditions (*α* = 0.3, *f* = 0.125), a stable fixed point occurs at *Ax*/*Bx* = 0.94002/0.09567.

**Fig 47c.**
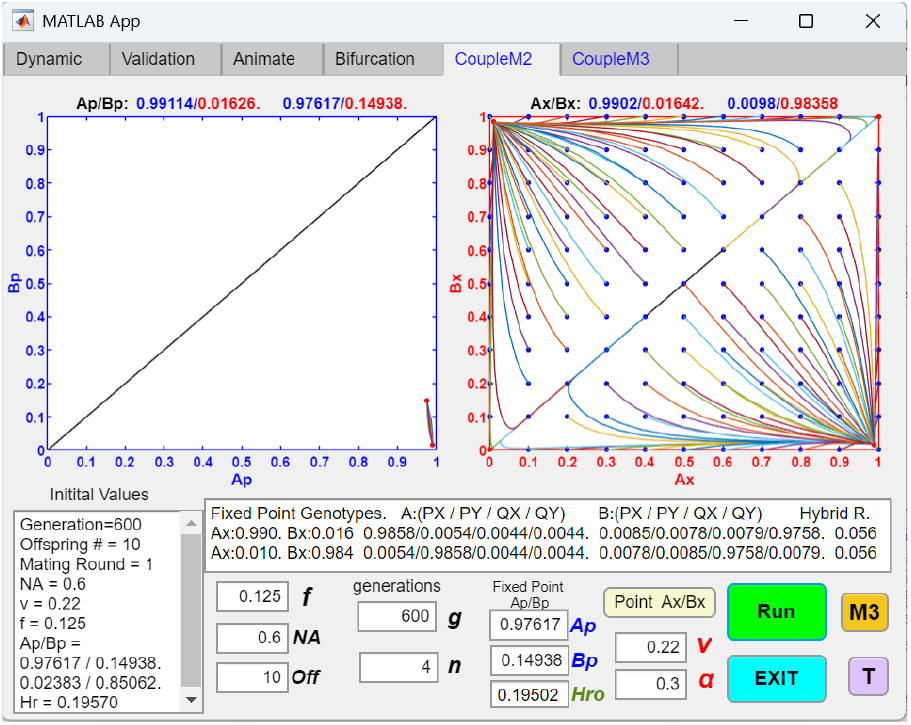
Positive feedback between premating mating-bias barriers and postzygotic incompatibility barriers strengthens both barriers and increases overall RI. In the GUI display, the hybrid-incompatibility barrier in Fig 47a first establishes postzygotic RI by generating a stable fixed point at *Ap*/*Bp* = 0.97617/0.14938. The resulting hybrid loss due to intrinsic incompatibility augments disruptive ecological selection against hybrid offspring from inter-niche matings. Consequently, reinforcement facilitates the emergence of the premating mating-bias barrier shown in Fig 47b. In the *Ax*/*Bx* phase portrait, a mutant *X* allele arising in niche *A* successfully invades at the established *Ap*/*Bp* fixed-point population and reaches a stable polymorphic equilibrium with the *Y* allele, producing premating RI. Phase-portrait analyses further show that premating isolation strengthens the postzygotic barrier by shifting its fixed point toward the lower-right corner of the *Ap*/*Bp* phase portrait (from *Ap*/*Bp* = 0.97617/0.14938 to 0.99114/0.01626), thereby increasing postzygotic RI between niche ecotypes. In turn, the strengthened postzygotic isolation further increases the premating mating-bias barrier via classical reinforcement, shifting the *Ax*/*Bx* fixed point toward the lower-right corner of the Ax/Bx phase portrait (from *Ax*/*Bx* = 0.94002/0.09567 to 0.9902/ 0.01642). Together, premating and postzygotic barriers mutually reinforce one another through positive feedback, resulting in stronger overall RI.

**Fig 47d.**
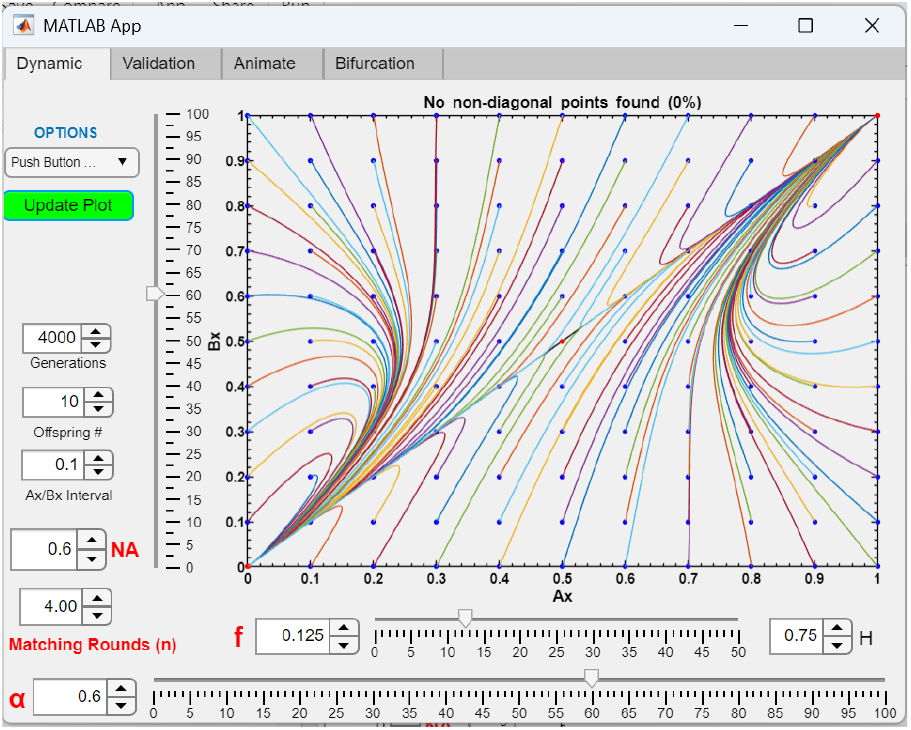
In the absence of hybrid incompatibility barriers, when the parameters *f* or *α* are sufficiently high, the system becomes divergent and no stable fixed points occur in the *Ax*/*Bx* phase portrait. Increasing the value of *α* in Fig 47b from 0.3 to 0.6 causes the phase portrait to become divergent, eliminating the fixed points.

**Fig 47e.**
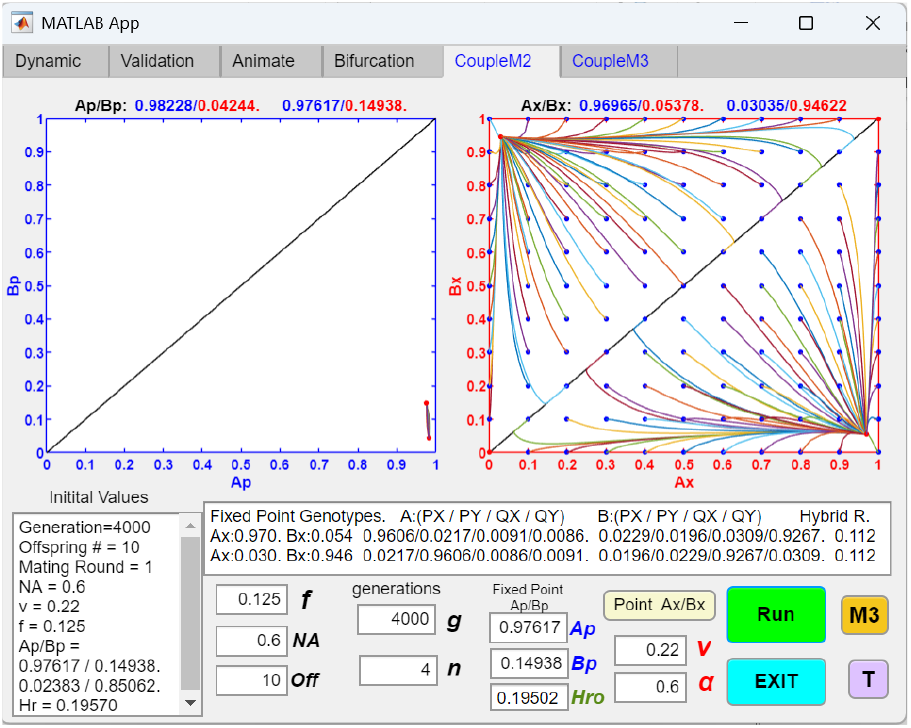
Intrinsic postzygotic RI can add to disruptive ecological selection and intensify hybrid loss from inter-niche matings, thereby facilitating the evolution of premating RI by reinforcement. In the GUI display, the hybrid-incompatibility barrier in Fig 47a first establishes postzygotic RI by generating a stable fixed point at *Ap*/*Bp* = 0.97617/0.14938. The resulting hybrid loss due to intrinsic incompatibility adds to existing disruptive ecological selection against hybrid offspring from inter-niche matings. The increased overall selection against hybrids relaxes the parametric conditions required for a subsequent mating-bias barrier to invade via reinforcement. Consequently, the mating-bias barrier shown in Fig 47d is able to establish system convergence and premating RI to reduce postmating inter-niche hybrid loss, whereas it would not be able to do so on its own. The resulting premating and postmating RI then reinforce one another through positive feedback to produce strong overall RI.

Fig 47c demonstrates that when the postzygotic hybrid-incompatibility barrier shown in Fig 47a is established prior to the premating mating-bias barrier, it facilitates the emergence of the premating barrier via classical reinforcement to reduce ongoing hybrid loss. After both barriers are established, the *Ax*/*Bx* and *Ap*/*Bp* fixed points shift closer to the lower-right corners of their respective phase portraits. This pattern indicates positive reinforcement between the premating mating-bias barrier and the postmating hybrid-incompatibility barrier, strengthening both barriers and increasing overall RI between niche ecotypes.

In Fig 47d, a weak mating-bias barrier with *f* = 0.125 and *α* = 0.6 cannot establish fixed-point convergence on its own. However, as shown in Fig 47e, by intensifying selection against inter-niche hybrid offspring, the presence of a preexisting postzygotic barrier can facilitate the establishment of such a weak premating barrier via reinforcement. This occurs because, in the mathematical model shown in Fig 3, ecological and sexual selection act complementarily to promote system convergence. Thus, adding intrinsic hybrid inviability to disruptive ecological selection increases inter-niche hybrid loss and lowers the effective value of *f*, making fixed-point convergence in the *Ax*/*Bx* phase portrait more readily attainable. After establishment, reciprocal reinforcement and positive feedback between premating and postzygotic barriers generate strong overall RI.

We also modified a GUI application originally developed for a two-allele, single-locus habitat-preference model in an open system [48] to investigate how premating barriers mediated by habitat preference reinforce postzygotic barriers formed through the establishment of a fixed point in the lower-right quadrant of the *Ap*/*Bp* phase portrait.

Fig 48a shows that, in the absence of hybrid incompatibility, fixed-point convergence in the habitat-preference *Ak*/*Bk* phase portrait—and thus premating RI generated by habitat preference—occurs when *f* = 0.125 and the habitat-preference parameters are *θ*1 = *θ*2 = 0.6.

**Fig 48a.**
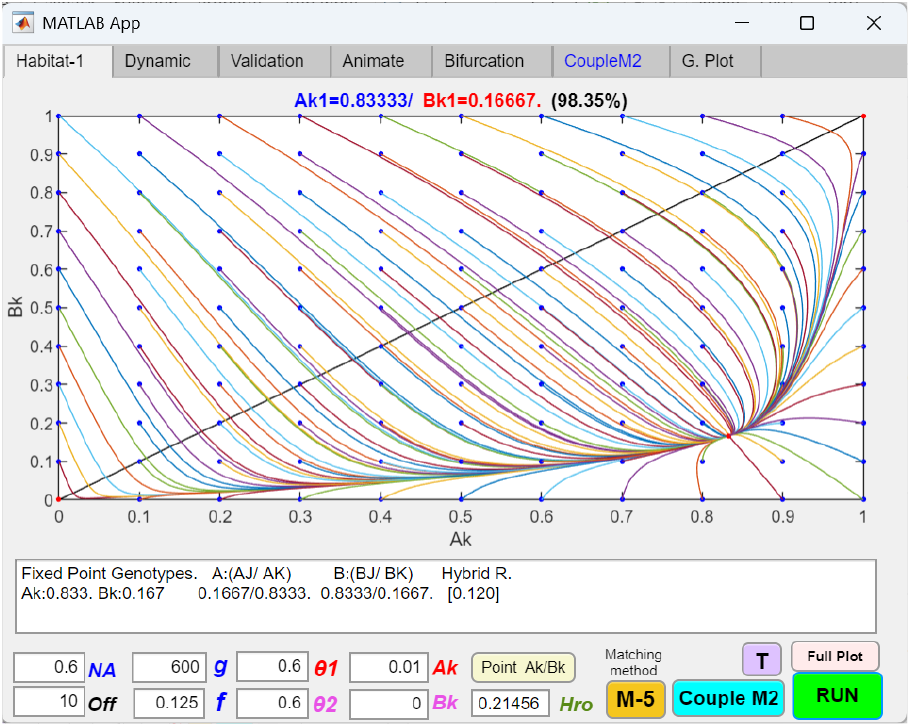
In the absence of hybrid incompatibility barriers, a stable fixed point appears in the *Ak*/*Bk* phase portrait of a habitat-preference barrier. A GUI application originally developed for a two-allele, single-locus habitat-preference model in an open system was modified to simulate coupling between habitat-preference and hybrid incompatibility barriers. Parameters *θθ*1 and *θθ*2 represent the strengths of habitat preference associated with alleles *K* and *J*, respectively. The *K* allele causes niche-*A* ecotypes carrying *K* to remain in niche *A* with probability 1 − *θ*1 and has no effect in niche-*B* ecotypes. Conversely, the *J* allele causes niche-*B* ecotypes carrying *J* to remain in niche *B* with probability 1 − *θθ*2 and has no effect in niche-*A* ecotypes. Under the parameter values shown, the *Ak*/*Bk* phase portrait converges to a stable equilibrium at *Ak*/*Bk* = 0.83333/0.16667.

**Fig 48b.**
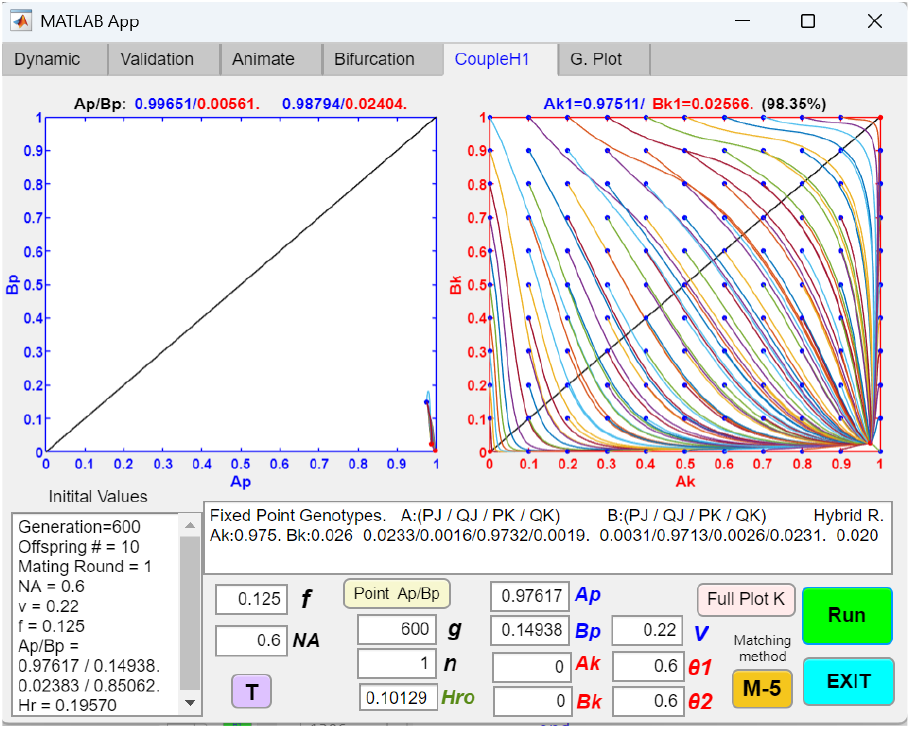
Positive feedback between premating habitat-preference barriers and postzygotic incompatibility barriers strengthens both barriers and increases overall RI. In the GUI display, the hybrid-incompatibility barrier in Fig 47a first establishes postzygotic RI by generating a stable fixed point at *Ap*/*Bp* = 0.97617/0.14938. Consequently, reinforcement facilitates the emergence of the premating habitat-preference barrier shown in Fig 48a. In the *AA*/*BA* phase portrait, a mutant *Ak* allele is able to invade at the established *Ap*/*Bp* fixed-point population and reach a stable fixed-point polymorphism to produce premating RI. Phase-portrait analyses show that premating habitat-preference isolation strengthens the postzygotic barrier by shifting its fixed point toward the lower-right corner of the *Ap*/*Bp* phase portrait (from *Ap*/*Bp* = 0.97617/0.14938 to 0.99651/0.00561), thereby increasing postzygotic RI between niche ecotypes. In turn, the strengthened postzygotic isolation further increases the premating habitat-preference barrier via classical reinforcement, shifting the *Ak*/*Bk* fixed point toward the lower-right corner of the *Ak*/*Bk* phase portrait (from *AA*/*BA* = 0.83333/0.16667 to 0.97511/ 0.02566). Together, premating and postzygotic barriers mutually reinforce one another through positive feedback, resulting in stronger overall RI.

Fig 48b shows that when the postzygotic hybrid-incompatibility barrier in Fig 47a is established prior to the premating habitat-preference barrier in Fig 48a, it facilitates the emergence of the habitat-preference barrier via reinforcement. After both barriers are established, the *Ak*/*Bk* and *Ap*/*Bp* fixed points shift closer to the lower-right corners of their respective phase portraits. This coordinated shift indicates reciprocal reinforcement between the premating habitat-preference barrier and the postzygotic hybrid-incompatibility barrier, strengthening both barriers and increasing overall RI between niche ecotypes.

Fig 49a demonstrates that when the postzygotic hybrid-incompatibility barrier in Fig 47a becomes established prior to the emergence of a second, identical postzygotic incompatibility barrier, the fixed points of both barriers (*Ap*1/*Bp*1 and *Ap*2/*Bp*2) shift toward the lower-right region of their respective phase portraits. This shift indicates positive reinforcement between the two postzygotic barriers, synergistically strengthening both barriers and increasing overall postmating RI between niche ecotypes.

**Fig 49a.**
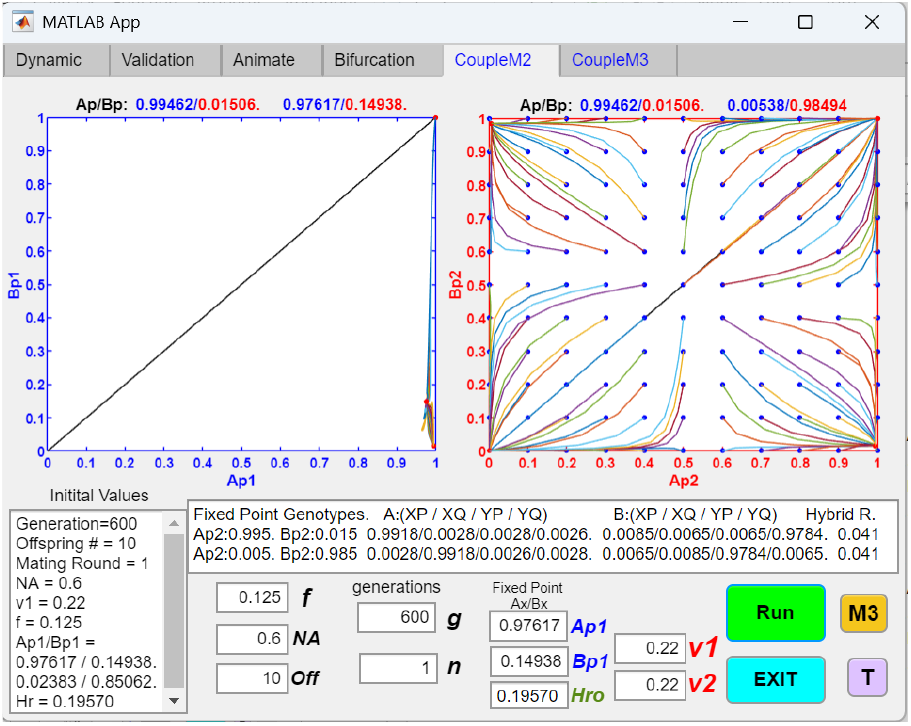
Positive feedback between two postzygotic hybrid-incompatibility barriers mutually strengthens each barrier and increases overall postmating RI. The GUI application shown in Fig 47c was modified to model coupling between two distinct postzygotic barriers. In this simulation, both barriers were assigned the same parameter values as in Fig 47a (*v*1 = 0.22 for barrier 1; *v*2 = 0.22 for barrier 2). The first postzygotic barrier (Fig 47a) initially establishes postmating RI by producing a stable fixed point at *Ap*1/*Bp*1 = 0.97617/0.14938. The second hybrid-incompatibility barrier subsequently emerges at this established *Ap*1/*Bp*1 fixed point. Phase-portrait analysis shows that the first barrier strengthens the second by shifting its fixed point toward the lower-right region of the *Ap*2/*Bp*2 phase portrait (from 0.97617/0.14938 to 0.99462/ 0.01506), thereby increasing postzygotic RI between niche ecotypes. Reciprocally, the strengthened second barrier reinforces the first, shifting its fixed point toward the lower-right region of the *Ap*1/*Bp*1 phase portrait (from 0.97617/ 0.14938 to 0.99462/0.01506). Together, the two postzygotic barriers form a positive feedback loop that increases overall postmating RI.

**Fig 49b.**
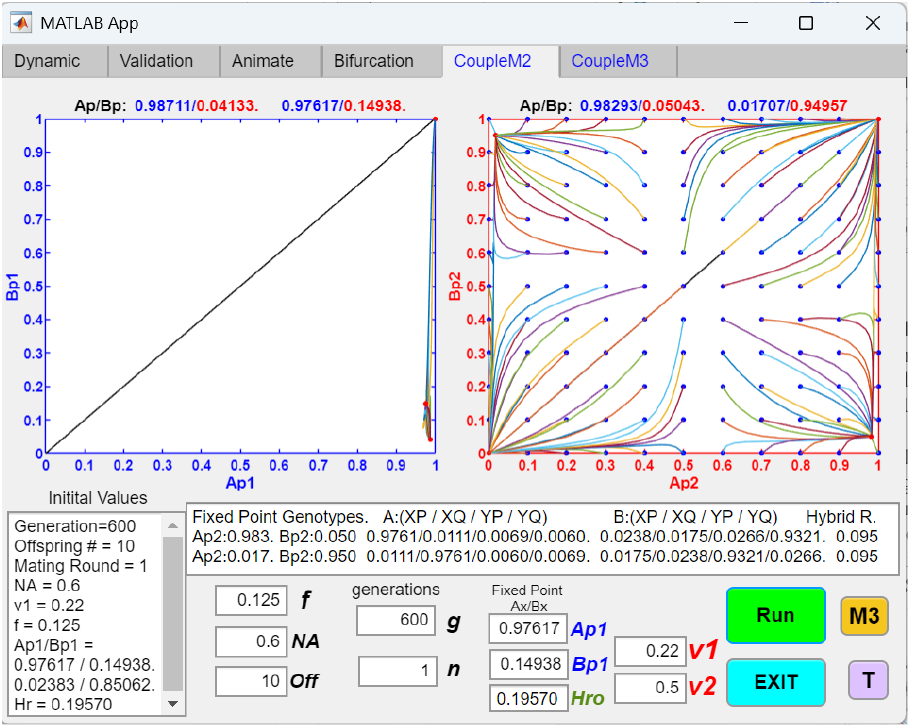
An existing postzygotic barrier increases the effective strength of selection against inter-niche hybrids, enabling the emergence of a second, weaker postzygotic barrier. When the viability ratio for the second postzygotic barrier in Fig 49a is increased from *v*2 = 0.22 to *v*2 = 0.5, the presence of the first postzygotic barrier still allows the second barrier to establish a stable fixed point in the *Ap*2/*Bp*2 phase portrait. In contrast, a weak postzygotic barrier with *v*2 = 0.5 cannot produce fixed-point convergence when acting alone. By increasing the effective strength of disruptive selection against inter-niche hybrid offspring, the first postzygotic barrier relaxes the conditions required for establishment of the second postzygotic barrier.

The example shown in Fig 49b further illustrates that the establishment of an initial postzygotic barrier can facilitate the emergence of a second, weaker postzygotic barrier by increasing the effective strength of disruptive selection against inter-niche hybrid offspring. In this scenario, the second barrier can arise only in the presence of the first barrier, as it is unable to achieve fixed-point convergence independently. Consequently, both barriers become reinforced, leading to stronger overall postmating RI.

#### 2 The effects of having multilocus ecological genotypes and viable ecological hybrids (without CI invasion) on one-locus hybrid-incompatibility barriers

We modified the GUI application MultiEcoCI to investigate how multilocus ecological genotypes and the presence of viable ecological hybrids influence a one-locus-equivalent hybrid incompatibility model (see Methodology). As expected, increasing the strength of disruptive ecological selection enhances postzygotic hybrid-incompatibility selection, consistent with the results shown in Figs 46b–46d.

As illustrated in Figs 50a–50b, during the pre-CI phase of the simulation (*Post* - *CI G* = 0), viable ecological hybrids weaken disruptive ecological selection and shift the *Ap*/*Bp* fixed point away from the lower-right corner, leading to reduced postzygotic RI. In contrast, in Fig 50c, increasing the number of ecological loci strengthens disruptive ecological selection and shifts the *Ap*/*Bp* fixed point toward the lower-right corner, resulting in stronger postzygotic RI.

**Fig 50a.**
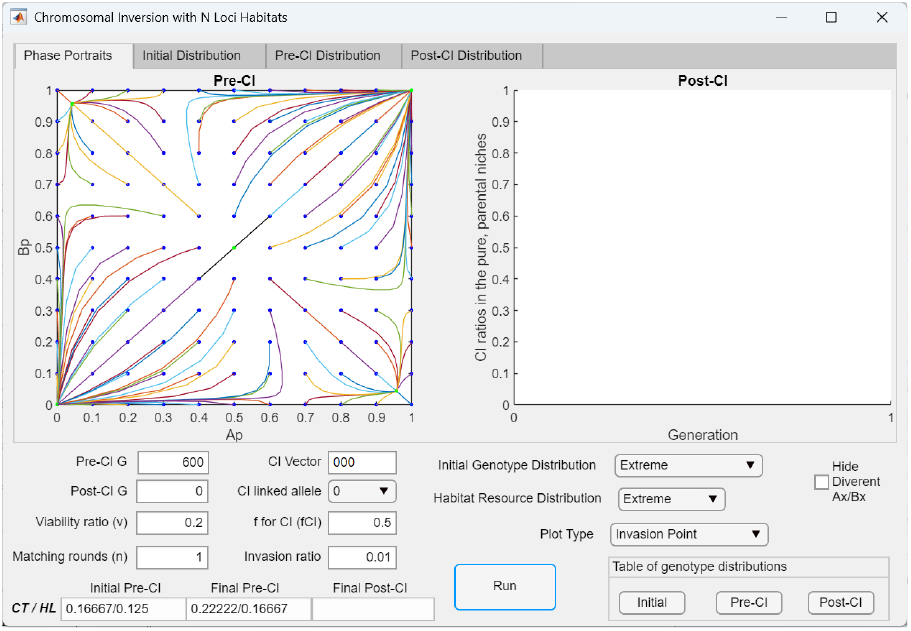
A fixed point appears in the lower-right quadrant of the *Ap*/*Bp* phase portrait for a one-locus-equivalent two-allele model of hybrid incompatibility with three ecological gene loci. In this model, the niche-*A* and niche-*B* ecotypes are specified by three loci. The habitat resource distribution is set to the *Extreme* option, in which resources are equally divided (50:50) between the parental ecotypes (*aaa* and *bbb*), with no resources available to support hybrid ecotypes. Matings between genotypes carrying different incompatibility alleles (*P* or *Q*) produce no viable hybrids, and the strength of incompatibility selection is governed by the viability ratio *v*. When *v* is sufficiently low (*v* = 0.2 in this example), indicating strong incompatibility selection, an *Ap*/*Bp* fixed point emerges in the lower-right region of the phase portrait.

**Fig 50b.**
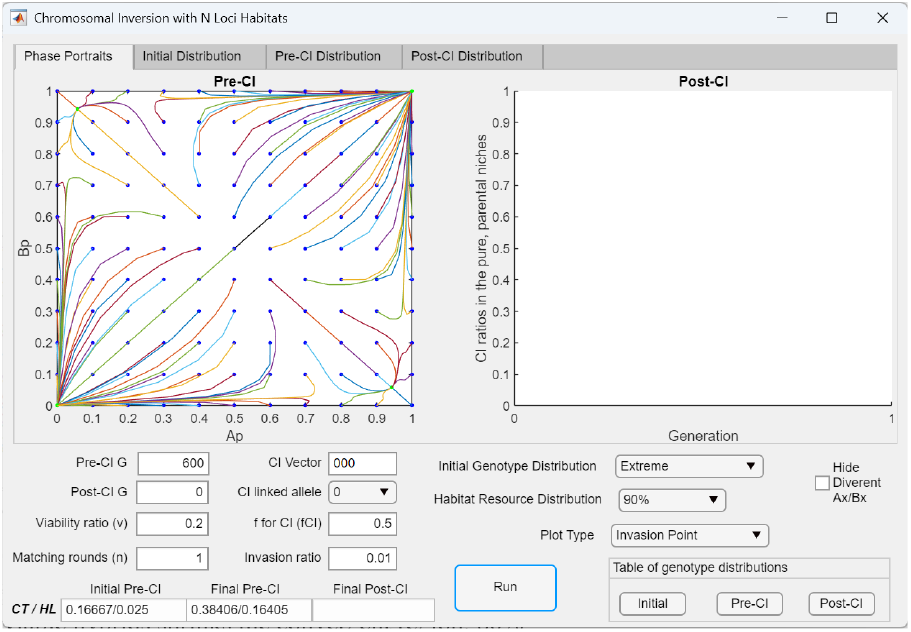
Viable ecological hybrids weaken disruptive ecological selection, shifting the *Ap*/*Bp* fixed point away from the lower-right corner and shrinking the region of convergence in the *Ap*/*Bp* phase portrait. In Fig 50a, selecting the *90%* option for the habitat resource distribution allocates 10% of niche resources to support viable hybrid ecotypes and results in weaker disruptive ecological selection (a higher value of *f*). This change shrinks the region of convergence in the lower-right quadrant of the *Ap*/*Bp* phase portrait and shifts the fixed point away from the lower-right corner (from *Ap*/*Bp* = 0.9566/0.0438 to 0.9419/0.0581), indicating weaker postzygotic RI between niche ecotypes. Selecting the *80%* option for habitat distribution eliminates both the region of convergence and the fixed point.

**Fig 50c.**
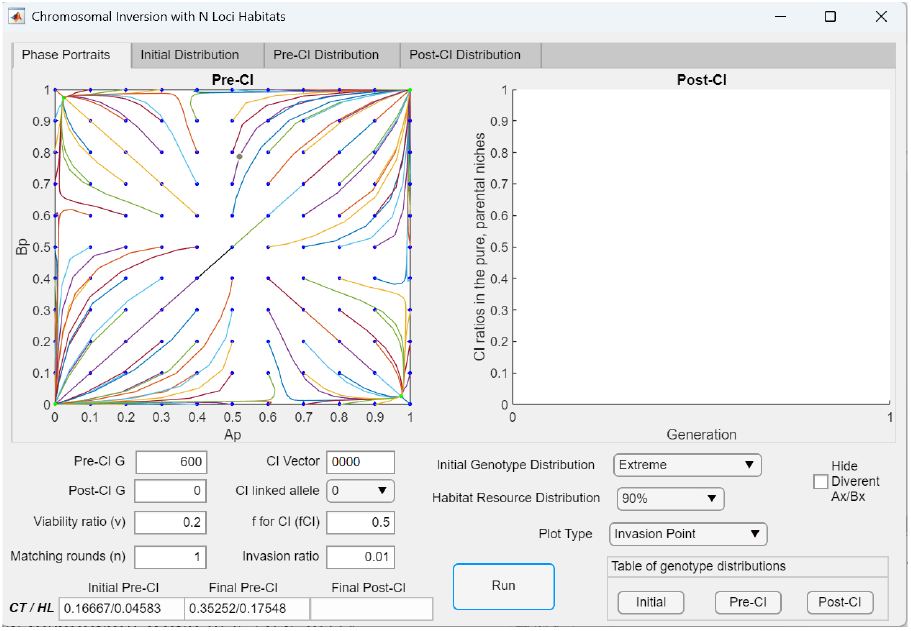
Increasing the number of ecological gene loci strengthens disruptive ecological selection and increases postzygotic RI, whereas decreasing the number of ecological gene loci produces the opposite effect. In Fig 50a, increasing the number of ecological gene loci from three to four strengthens disruptive ecological selection (a lower value of *f*). This change expands the region of convergence in the lower-right quadrant of the *Ap*/*Bp* phase portrait and shifts the *Ap*/*Bp* fixed point closer to the lower-right corner, indicating stronger postzygotic RI between niche ecotypes. Conversely, reducing the number of ecological gene loci from three to two eliminates both the region of convergence and the fixed point.

#### 3 Invasion of a CI in a two-niche, two-hybrid-incompatibility-allele system with multilocus ecological genotypes

By setting the number of post-CI generations (*Post*-*CI G*) to values greater than zero, we used the modified MultiEcoCI application to examine the invasion dynamics of CI arising in niche *A* and its effects on system behavior. We present the results in two parts: first, when the CI captures only ecological alleles, and second, when it captures both ecological and hybrid-incompatibility alleles.

##### 3.1 Invasion of a CI capturing only ecological alleles

First, we examine the invasion dynamics of a CI that captures only locally adaptive ecological alleles without capturing any hybrid-incompatibility allele. This is implemented by selecting “0” for *CI linked allele* in the user interface.

As illustrated by the example in Fig 51, the invasion of a CI capturing only ecological alleles reduces the effective number of ecological gene loci and weakens disruptive ecological selection, leading to either weakened postzygotic RI or system divergence and elimination of the *Ap*/*Bp* fixed point.

**Fig 51.**
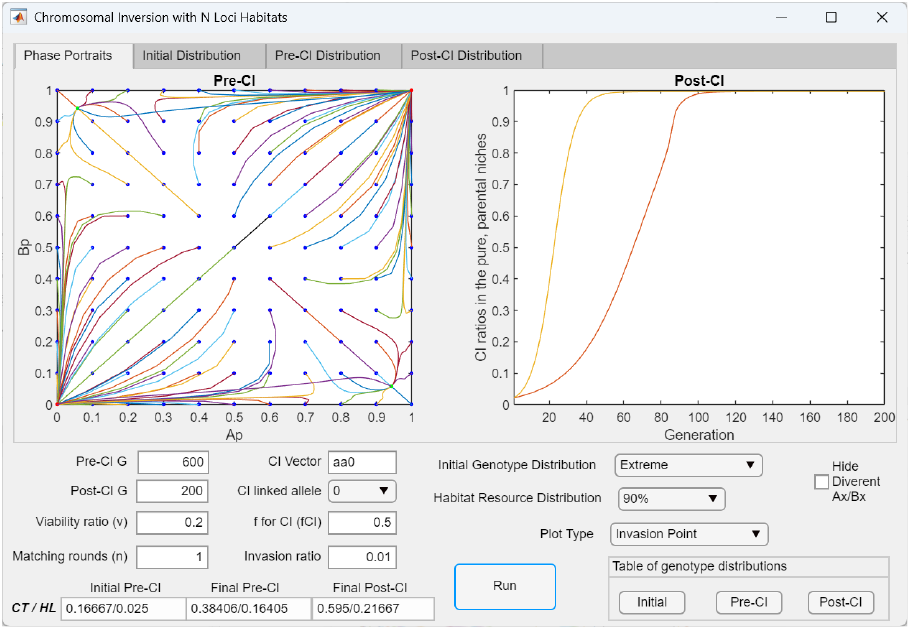
Invasion of a CI capturing only locally adaptive ecological alleles reduces disruptive ecological selection and weakens postzygotic reproductive isolation. The *Invasion Point* plot shows the successful invasion of a CI capturing two of three locally adaptive ecological alleles in niche *A* (*CI* = *aaaa*0) at the fixed point established in Fig 50a. As the CI mutant (initial invasion ratio = 0.01) successfully invades and becomes fixed in niche *A*, it reduces the effective number of ecological gene loci from three to two, thereby weakening disruptive ecological selection. Consequently, invasion of the CI eliminates the preestablished fixed-point polymorphism in the *Ap*/*Bp* phase portrait and drives the population vector to the origin, where the *P* allele is lost.

##### 3.2 Invasion of a CI capturing both ecological alleles and a hybrid-incompatibility allele

In contrast, when a CI captures locally adaptive ecological alleles and the locally prevalent hybrid-incompatibility allele in a niche, its invasion anchors the incompatibility allele in the niche where it has a numerical advantage, prevents its introgression into the opposite niche, and further increases its prevalence within the niche. As a result, the *Ap*/*Bp* fixed point shifts to the lower-right corner of the phase portrait.

Fig 52a shows an example in which invasion of a CI capturing two of three locally adaptive ecological alleles and the prevalent hybrid-incompatibility allele *P* in niche *A* (CI = *aa*0*P*) shifts the pre-established *Ap*/*Bp* fixed point to the lower-right corner of the phase portrait, resulting in stronger postzygotic RI. Fig 52b shows the same outcome following invasion of a CI capturing one of three ecological alleles and the prevalent hybrid-incompatibility allele (CI = *a*00*P*). Compared with the invasion trajectory in Fig 52a, Fig 52b shows that capturing fewer locally adaptive alleles confers lower invasion fitness, as reflected by the longer generation time required for the CI to reach fixation. In contrast, a CI capturing the less prevalent Q allele (*CI* = *aa*0*Q*) in niche *A* cannot invade and goes extinct.

**Fig 52a.**
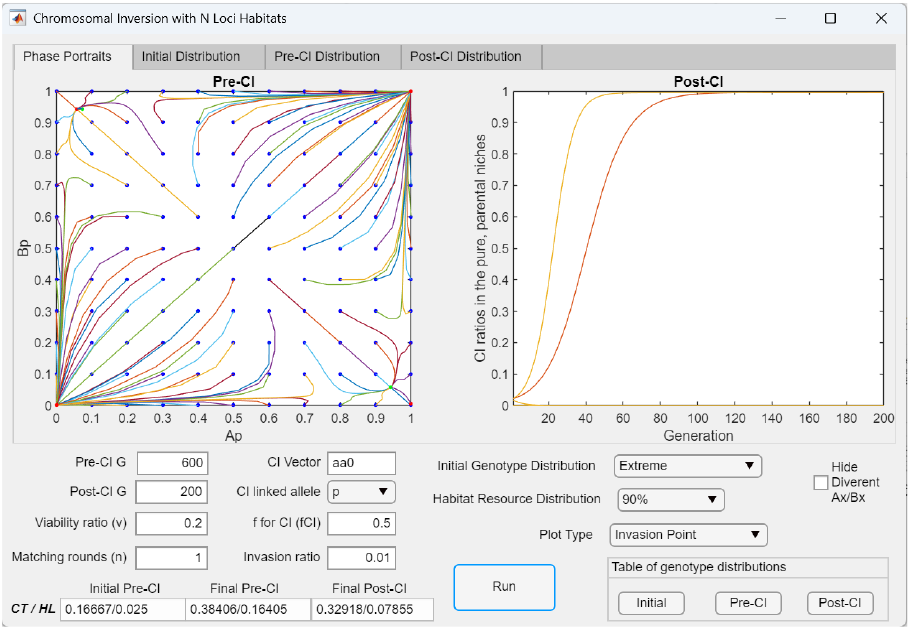
Invasion of a CI that captures two of the three locally adaptive ecological alleles and the locally prevalent incompatibility allele (CI = *aa*0*PP*) shifts the *Ap*/*Bp* fixed point to the lower-right corner of the phase portrait. When the CI shown in Fig 51 also captures the locally prevalent incompatibility allele (*CI* = *aa*0*P*), its invasion at the *Ap*/*Bp* fixed point shown in Fig 50a shifts the fixed point to the lower-right corner, resulting in the maximal level of postzygotic RI permitted by *v*. By anchoring the *P* allele to the niche-*A* ecotype, the CI prevents introgression of *P* into niche *B*, leading to fixation of *P* in niche *A*. In contrast, a CI capturing the less prevalent *Q* allele (*CI* = *aa*0*Q*) cannot invade and goes extinct.

**Fig 52b.**
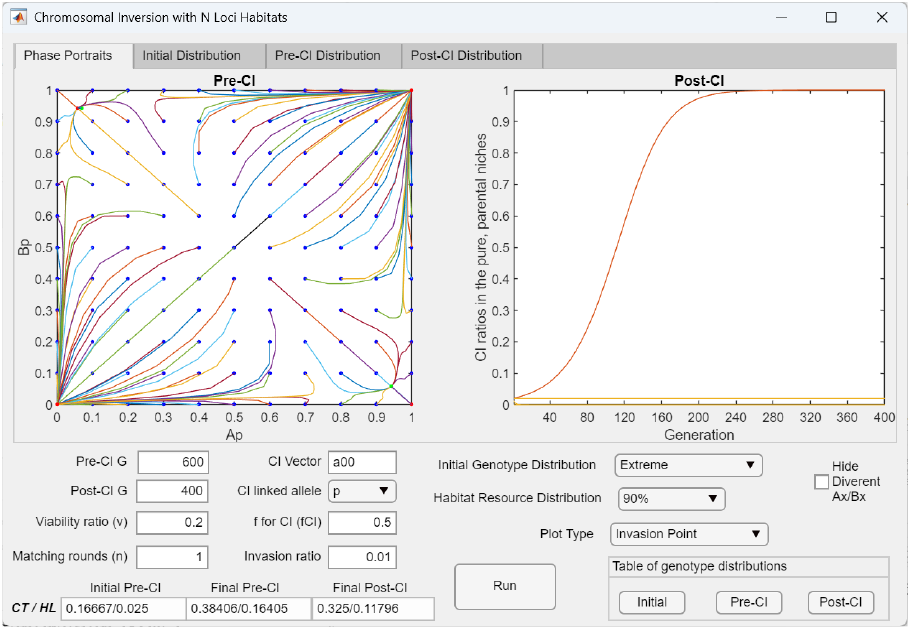
Invasion of a CI capturing one of the three locally adaptive ecological alleles and the locally prevalent incompatibility allele (*CI* = *a*00*PP*) shifts the *Ap*/*Bp* fixed point to the lower-right corner of the phase portrait but exhibits reduced invasion fitness. When the CI shown in Fig 52a captures only one locally adaptive ecological allele (CI = *aa*00*P*), its invasion at the *Ap*/*Bp* fixed point established in Fig 50a similarly displaces the fixed point to the lower-right corner of the phase portrait. However, relative to a CI capturing more locally adaptive ecological alleles (Fig 52a), this CI has reduced invasion fitness, as indicated by the longer time required to reach fixation in niche *A* (∼200 generations versus ∼100 generations).

### IV. Systems with multilocus hybrid-incompatibility genotypes

We used the GUI application MultiCompCI to examine how direct selection against mismatched hybrid-incompatibility genotypes influences system dynamics in models with multiple incompatibility loci. We then analyzed the invasion dynamics of CI variants capturing different incompatibility alleles. MultiCompCI enables simulations with an arbitrary number of hybrid-incompatibility loci. Following each mating generation, mismatched hybrid offspring were removed using a V-shaped viability function parameterized by the minimum viability value *v* (see Methodology). To isolate pre-CI dynamics, the post-CI phase was initially disabled by setting the number of post-CI generations (*Post* - *CI G*) to zero. In subsequent simulations, the post-CI phase was enabled, and CI mutations capturing different combinations of hybrid-incompatibility alleles were introduced to evaluate their invasion fitness and effects on overall system behavior.

#### 1 The effects of having multilocus hybrid-incompatibility genotypes (without CI invasion)

We first examined how varying the number of hybrid-incompatibility gene loci affects population dynamics in the *AP*/*BP* phase portrait. To do so, the post-CI phase was disabled by setting *Post* - *CI G* = 0.

Fig 53a shows that, in the model with direct selection against mismatched hybrid offspring using a V-shaped viability function, the *A*/*BP* phase portrait is divergent, even though the same parameter values produce convergence in the one-locus-equivalent model shown in Fig 46d. This difference arises because the elimination functions for mismatched hybrid offspring differ between the two models.

**Fig 53a.**
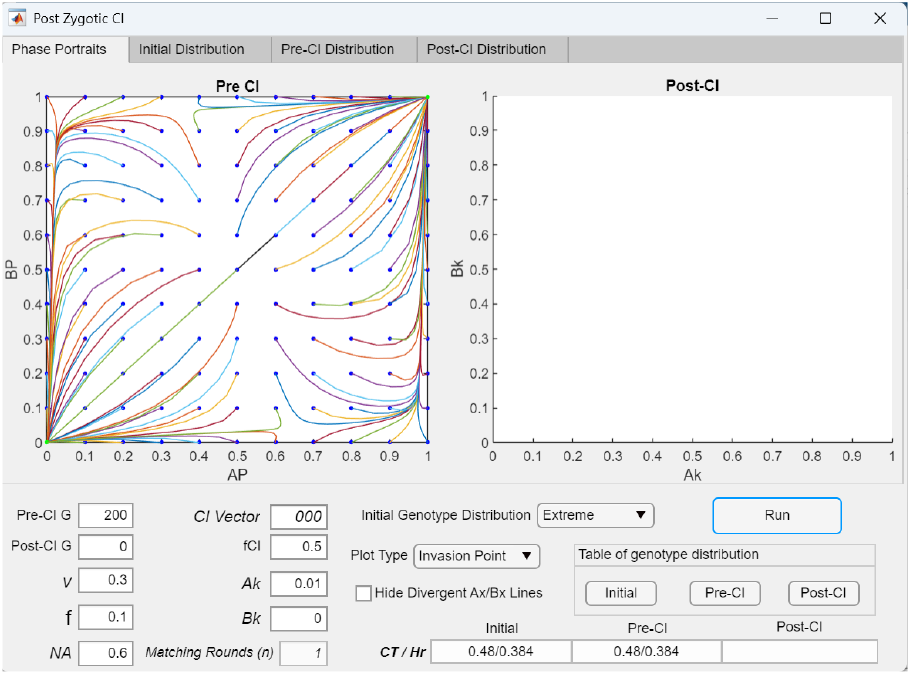
In a model with three incompatibility gene loci, the *AP*/*BP* phase portrait fails to converge without sufficiently strong disruptive ecological selection and hybrid incompatibility selection. With *v* = 0.3 and *f* = 0.1, the *AP*/*BP* phase portrait is divergent, even though these same parameter values yield convergence in the corresponding one-locus-equivalent two-allele model shown in Fig 46d. This difference arises because the two models are not directly comparable: in Fig 46d, *v* is defined assuming no viable incompatibility hybrids, whereas in the multilocus model, hybrid genotypes are eliminated via a V-shaped fitness function parameterized by *v*. In the current model, the *AP*/*BP* phase portrait remains divergent for *v* = 0.3 and *f* = 0.1 when the number of incompatibility loci is fewer than five.

**Fig 53b.**
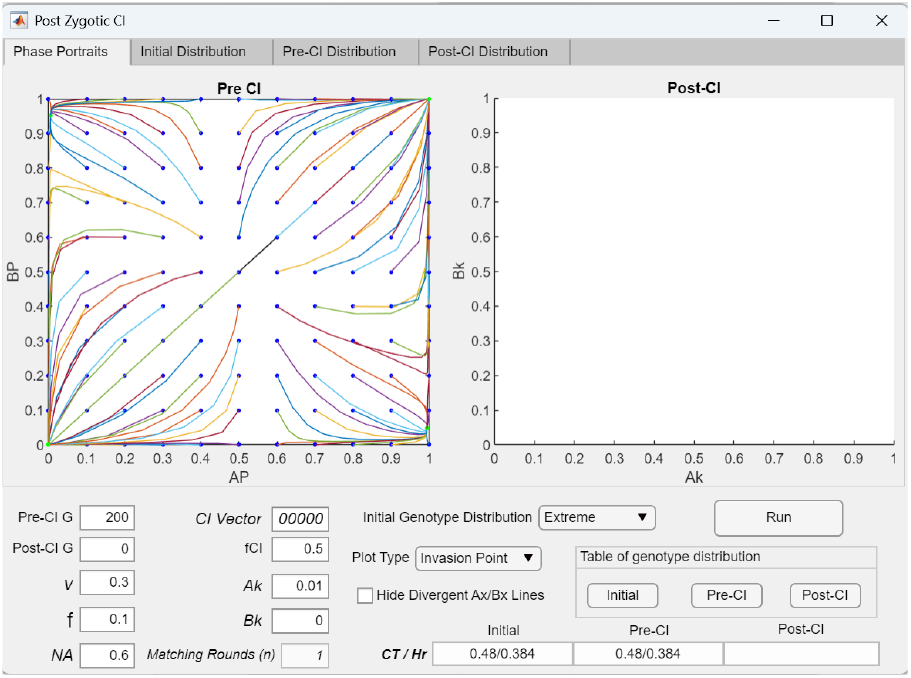
In a multilocus incompatibility model, increasing the number of gene loci promotes fixed-point convergence in the lower-right quadrant of the *AP*/*BP* phase portrait. Increasing the number of loci in Fig 53a from three to five generates a fixed point in the lower-right quadrant of the *AP*/*BP* phase portrait. This occurs because additional loci increase the proportion of hybrid offspring genotypes, which experience incompatibility selection under the V-shaped fitness function, thereby reducing the effective value of *v* and strengthening hybrid incompatibility selection.

**Fig 53c.**
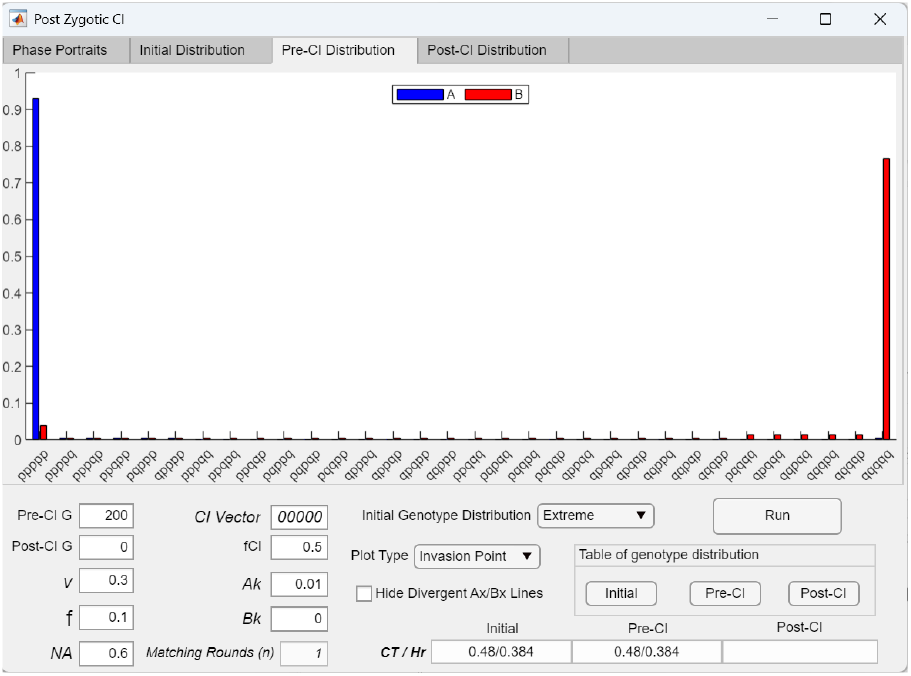
Genotype distribution at the fixed point in a model with five incompatibility gene loci. The pre-CI bar graph shows the genotype distribution at the fixed point of the *AP*/*BP* phase portrait in Fig 53b. Intermediate hybrid genotypes between the extreme-*P* and extreme-*Q* genotypes are eliminated by a V-shaped fitness function due to intrinsic genetic incompatibilities.

In the one-locus-equivalent model, elimination of mismatched hybrids is parameterized by *v*, the total offspring return ratio for the extreme parental genotypes. This measure (*v*) includes both offspring with parental genotypes (i.e., as determined by the actual number of incompatibility gene loci) and parental genotypes regenerated through matings among hybrid offspring. In contrast, in the direct elimination model, incompatible hybrid offspring are removed using a V-shaped viability function based on *v*. Although the elimination functions differ slightly and the results do not match exactly, the two models show qualitatively similar trends across parameter changes.

As demonstrated by the example shown in Figs 53b and 53c, increasing the number of hybrid-incompatibility gene loci promotes system convergence and the emergence of fixed points in the *AP*/*BP* phase portrait. This occurs because increasing the number of gene loci increases both hybrid production and hybrid loss, while decreasing the effective value of *v*. Similarly, as shown in Figs 54a and 54b, directly decreasing *v* also increases hybrid loss and promotes fixed-point convergence and stronger postzygotic RI. Notably, in Figs 53c and 54b, mismatched hybrid genotypes still exist at the fixed-point genotype distribution, although their frequencies are much lower than those of the parental genotypes.

**Fig 54a.**
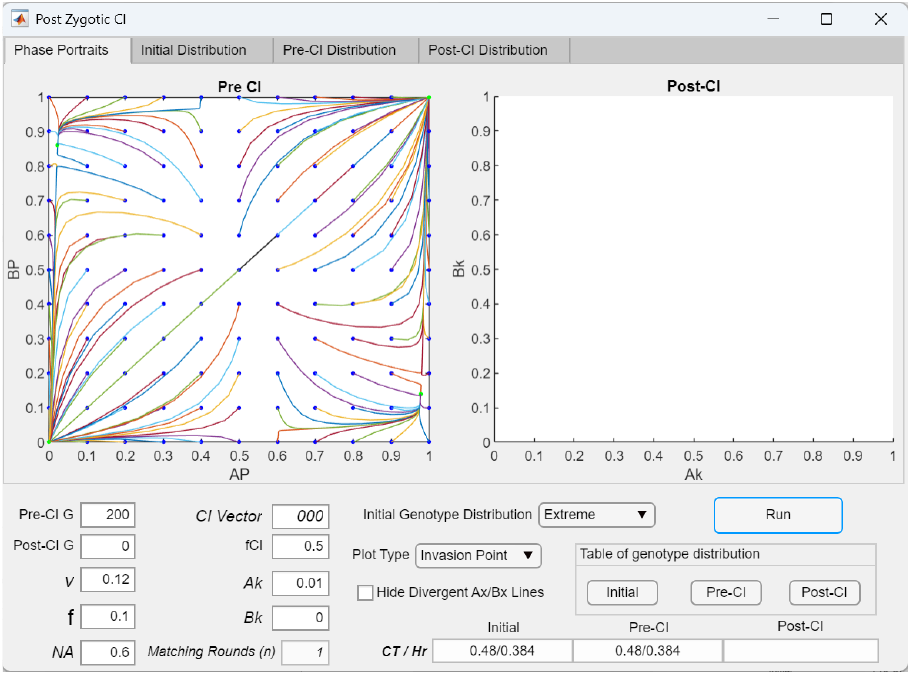
In a multilocus incompatibility model, increasing hybrid incompatibility (decreasing *v*) promotes fixed-point convergence in the lower-right quadrant of the *Ap*/*Bp* phase portrait. As shown, decreasing *v* from 0.3 to 0.12 in Fig 53a generates a fixed point in the lower-right quadrant of the *AP*/*BP* phase portrait. This occurs because lower *v* increases the elimination of hybrid offspring genotypes under the V-shaped fitness function, thereby strengthening hybrid incompatibility selection. With all other parameters held constant, convergence occurs when *v* ≤ 0.12.

**Fig 54b.**
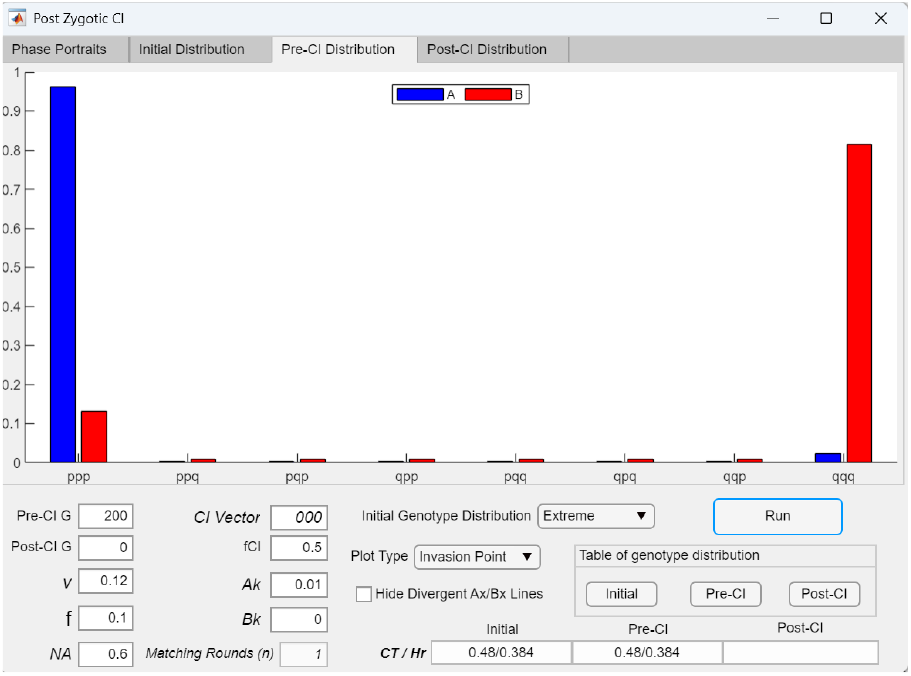
Genotype distribution at the fixed point in a model with three incompatibility gene loci. The pre-CI bar graph shows the genotype distribution at the fixed point of the *AP*/*BP* phase portrait in Fig 54a.

In Fig 55, decreasing *f* increases the strength of ecological selection, such that most offspring are produced from intra-niche rather than inter-niche matings. This shifts the pre-established *AP*/*BP* fixed point toward the lower-right corner of the phase portrait, resulting in stronger postzygotic RI.

**Fig 55.**
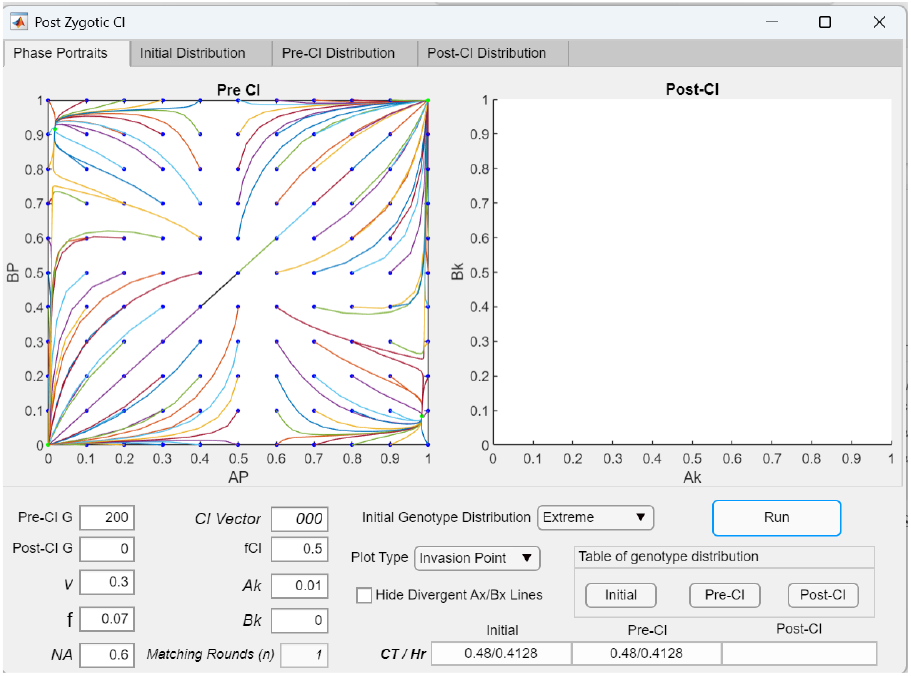
In a multilocus incompatibility model, strengthening disruptive ecological selection (decreasing *f*) promotes fixed-point convergence in the lower-right quadrant of the *AP*/*BP* phase portrait. As shown, reducing f from 0.1 to 0.07 in Fig 53a generates a fixed point in lower-right quadrant of the *AP*/*BP* phase portrait. This occurs because lower *f* reduces gene flow from inter-niche matings, allowing the locally prevalent incompatibility allele (*P* or *Q*) in each niche to more effectively eliminate the less prevalent allele. With all other parameters held constant, fixed-point convergence occurs when *f* ≤ 0.07.

In Fig 56, increasing the number of hybrid-incompatibility gene loci strengthens incompatibility selection. This also shifts the pre-established *AP*/*BP* fixed point toward the lower-right corner of the phase portrait.

**Fig 56.**
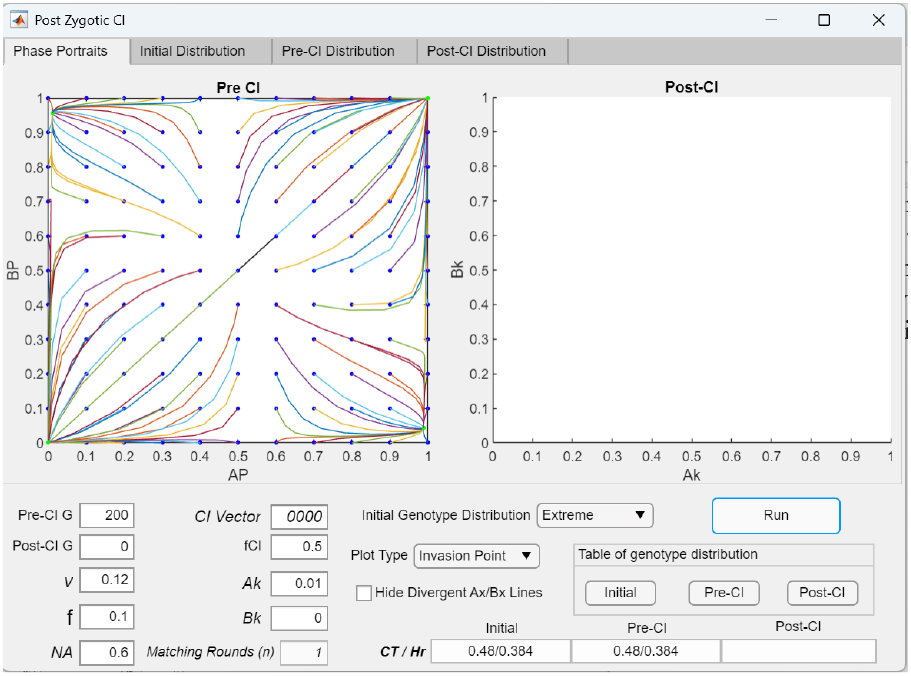
In a multilocus incompatibility model with a stable fixed point in the *AP*/*BP* phase portrait, increasing the number of incompatibility loci shifts the fixed point closer to the lower-right corner and strengthens postzygotic RI. As shown, increasing the number of loci in Fig 54a from three to four moves the fixed point toward the lower-right corner of the phase portrait, resulting in stronger postzygotic RI between niche ecotypes.

#### 2 Invasion of a CI capturing hybrid-incompatibility alleles in a two-niche system with multilocus hybrid-incompatibility genotypes

The invasion dynamics of a CI capturing hybrid-incompatibility alleles in the multilocus mating-bias model are examined by specifying the CI Vector and setting the post-CI generation (*Post*-*CI G*) to be greater than zero.

Fig 57a illustrates that, in a model with multiple hybrid-incompatibility gene loci and an established *AP*/*BP* fixed point, a CI that arises within the fixed-point population but captures only a single incompatibility allele (*CI* = *p*000) gains no fitness advantage and cannot invade.

**Fig 57a.**
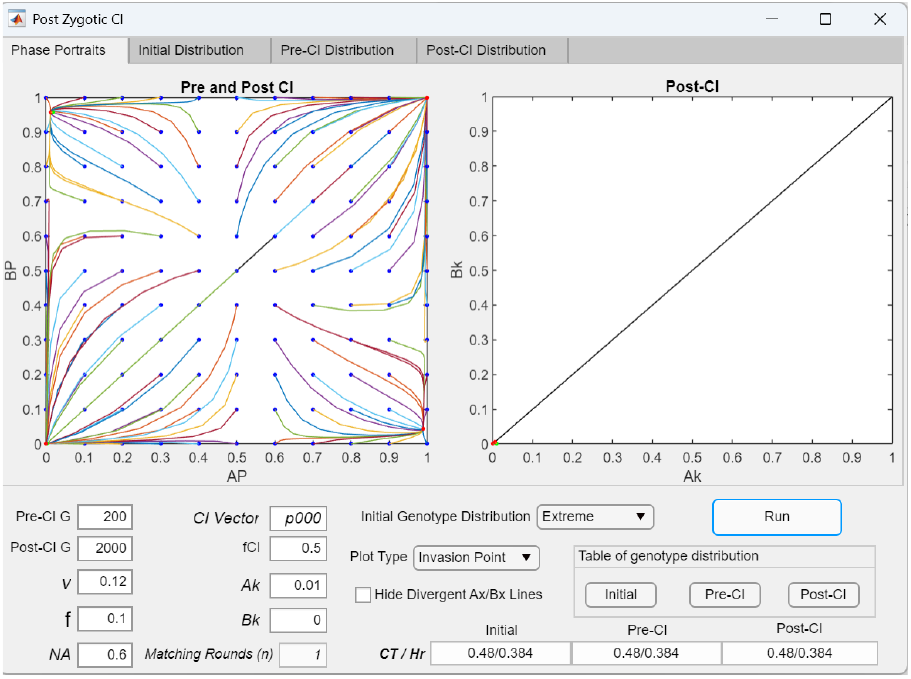
In a multilocus incompatibility model, a CI capturing a single incompatibility allele has no fitness advantage and cannot invade. As shown, a CI capturing one incompatibility allele (CI = *p*000) fails to invade the fixed-point population shown in Fig 56 and goes extinct. Consequently, the post-CI fixed point remains unchanged.

**Fig 57b.**
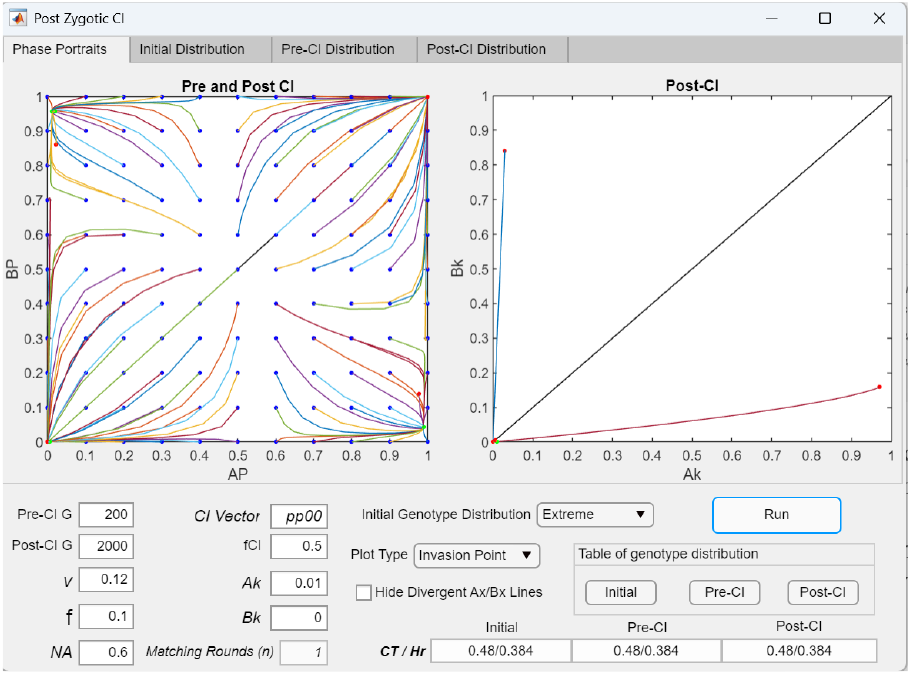
In a model with four incompatibility gene loci, successful invasion of a CI capturing two extreme-type incompatibility alleles (CI = *pp*00) leads to weaker postzygotic RI. In contrast to a CI capturing a single incompatibility allele (CI = *p*000; Fig 57a), a CI capturing two extreme-*P* alleles (CI = *pp*00) gains a fitness advantage and is able to invade. This advantage arises because matings between individuals carrying the CI and individuals with different, mismatched genotypes produce fewer hybrid offspring, which are less viable due to genetic incompatibilities. However, because successful invasion of the CI reduces the effective number of incompatibility loci experienced by individuals in the system, postzygotic RI is weakened. This is demonstrated in the *Invasion Point* plot: as the CI mutant (CI = *pp*00) invades and reaches a fixed-point polymorphism in the post-CI *Ak*/*Bk* phase portrait, the pre-CI *AP*/*BP* fixed point shifts away from the lower-right corner of the *AP*/*BP* phase portrait, indicating reduced postzygotic RI between niche ecotypes.

**Fig 57c.**
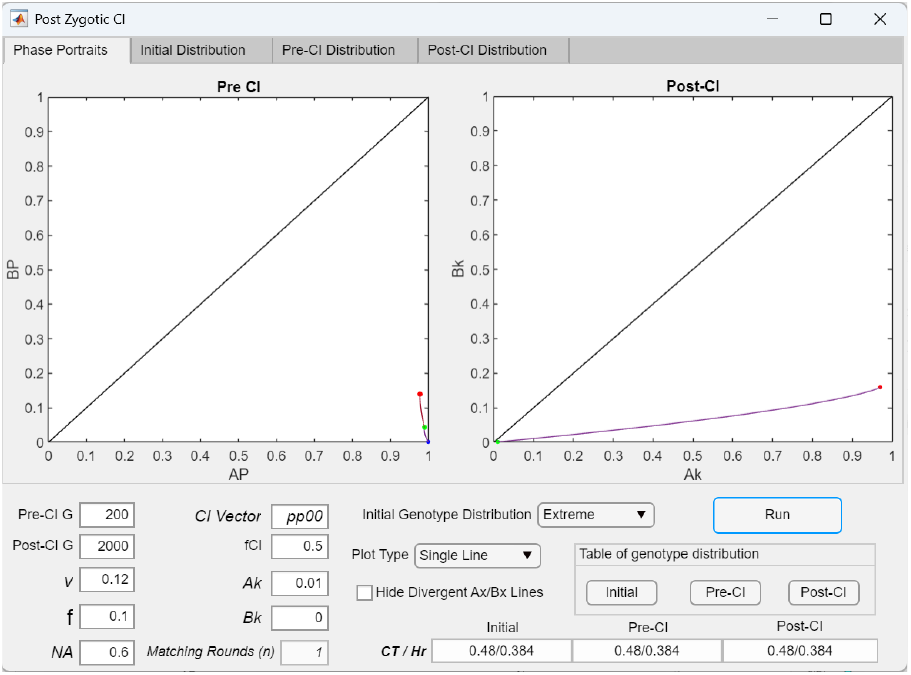
*Single Line* plot illustrating invasion of a CI capturing two extreme-type incompatibility alleles (CI =*pp*00) and the resulting shift of the *AP*/*BP* fixed point away from the lower-right corner of the phase portrait. Selecting the *Single Line* plot option in Fig 57b shows the pre-CI *AP*/*BP* fixed point (green dot) shifting away from the lower-right corner of the phase portrait (to the red dot) as the CI mutant successfully invades and reaches a fixed-point polymorphism (*Ak*/*Bk* = 0.970362/0.158427) in the post-CI *Ak*/*Bk* phase portrait.

**Fig 57d.**
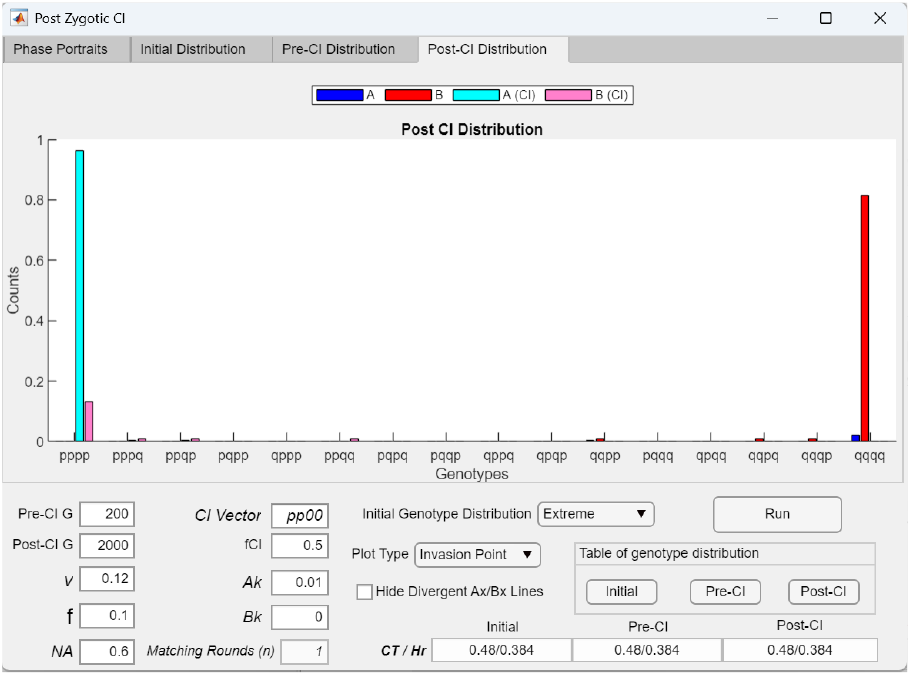
Post-CI genotype distribution following invasion of a CI capturing two extreme-P alleles (CI = *pp*00) in a model with four incompatibility gene loci. The post-CI bar graph shows the genotype distribution after successful invasion of the CI in Fig 57c.

**Fig 57e.**
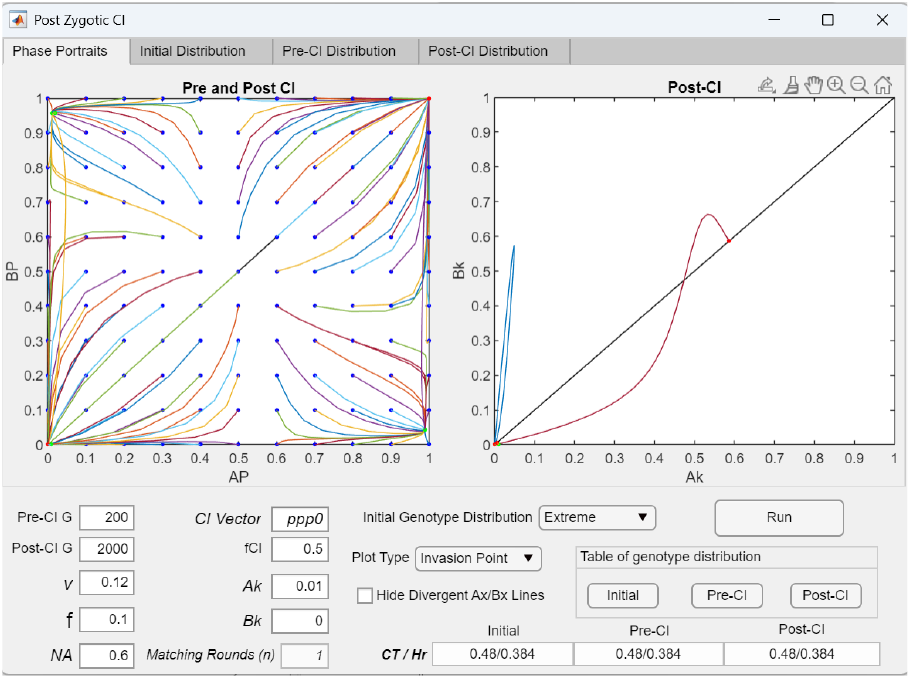
In a model with four incompatibility gene loci, invasion of a CI capturing three extreme-type incompatibility alleles (CI = *ppp*0) causes the *AP*/*BP* phase portrait to become divergent. Increasing the number of extreme-*P* alleles captured by the invading CI in Fig 57b from two to three (i.e., CI = *ppp*0) further reduces the effective number of incompatibility loci experienced by individuals in the system, to the point that fixed-point convergence in the *AP*/*BP* phase portrait is no longer possible. Consequently, following invasion of the CI (CI = *ppp*0), the pre-CI *AP*/*BP* fixed point is eliminated and the population trajectory moves to the upper-right corner of the phase portrait, where only the extreme-*P* genotype persists in both niches. The same outcome is observed for a CI capturing all extreme-*P* alleles (CI = *pppp*).

**Fig 57f.**
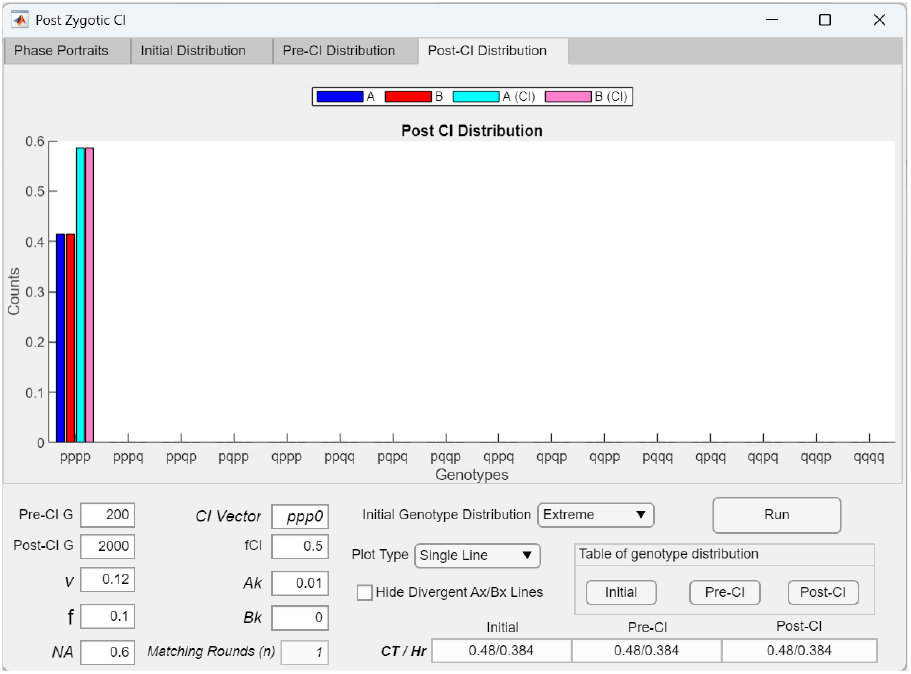
Post-CI genotype distribution following invasion of a CI capturing three extreme-*PP* alleles (CI = *ppp*0) in a model with four incompatibility gene loci. The post-CI genotype distribution bar graph shows that only the extreme-*P* genotype persists in both niches following successful invasion of the CI in Fig 57e.

Figs 57b–57d demonstrate that when the CI shown in Fig 57a captures two of the four extreme-*P* incompatibility alleles (*CI* = *pp*00), it gains a fitness advantage because individuals carrying the CI are less likely to produce mismatched offspring than individuals without the CI. This advantage enables the CI to invade and reach a stable fixed-point polymorphism in the *Ak*/*Bk* phase portrait. However, by reducing the effective number of incompatibility gene loci, successful CI invasion shifts the original *AP*/*BP* fixed point away from the lower-right corner of the phase portrait, resulting in weaker postzygotic RI.

In Figs 57e–57f, when the CI captures three of the four extreme-*P* incompatibility alleles, successful CI invasion reduces the effective number of incompatibility gene loci to such an extent that the system becomes divergent, eliminating the fixed points in the *AP*/*BP* phase portrait.

Fig 58 shows that a CI capturing incompatibility alleles cannot invade when the *AP*/*BP* phase portrait lacks a pre-established fixed point. In other words, an adaptive CI mutation capturing a favorable combination of incompatibility alleles cannot invade from near the origin and can only spread within a population already located at a fixed point in the *AP*/*BP* phase portrait.

**Fig 58.**
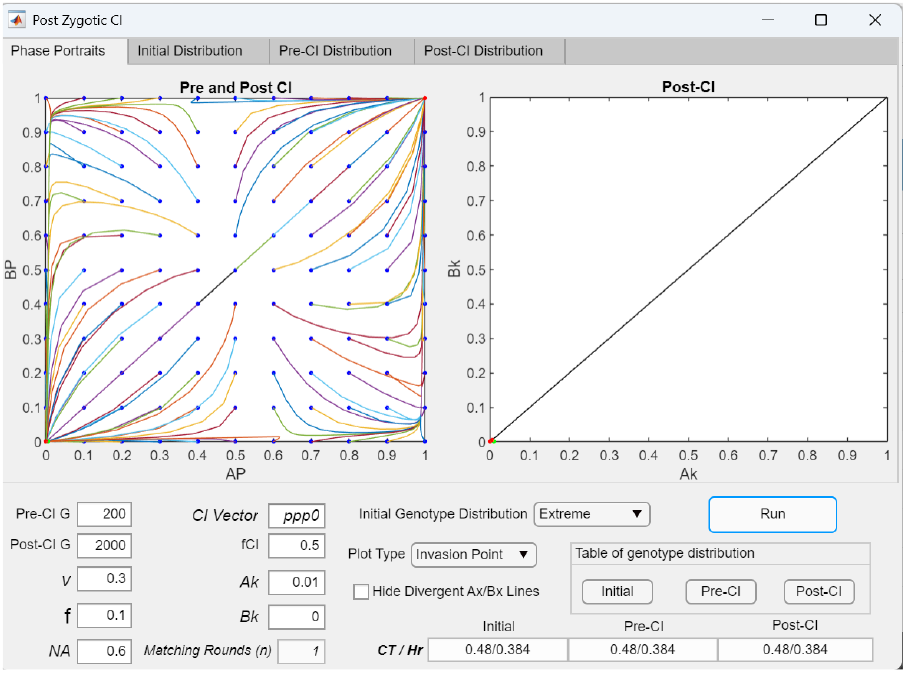
No CIs capturing incompatibility alleles can invade when no fixed point exists in the pre-CI *AP*/*BP* phase portrait. Increasing *v* in Figs 57a–57f from 0.12 to 0.3 causes the pre-CI *AP*/*BP* phase portrait to become divergent and eliminates the fixed point. Consequently, no CI capturing any number or combination of incompatibility alleles is able to invade. Therefore, invasion of an adaptive CI is possible only when a fixed point already exists in the *AP*/*BP* phase portrait.

Fig 59 shows that early CI presence in an initially segregated sympatric population can prevent the later establishment of a fixed point in the *AP*/*BP* phase portrait. This can occur when an adaptive CI (*CI* = *pp*0 in the Fig 59 example) that reduces the effective number of incompatibility gene loci is already present when niche ecotypes carrying different incompatibility alleles meet in sympatry, such as during secondary contact. In this case, CI invasion and spread reduce the effective number of incompatibility loci, thereby inhibiting fixed-point divergence and the evolution of postzygotic RI. This scenario can be simulated in the GUI application by setting *Pre*-*CI GG* to zero and *Post* - *CI G* to a value greater than one.

**Fig 59.**
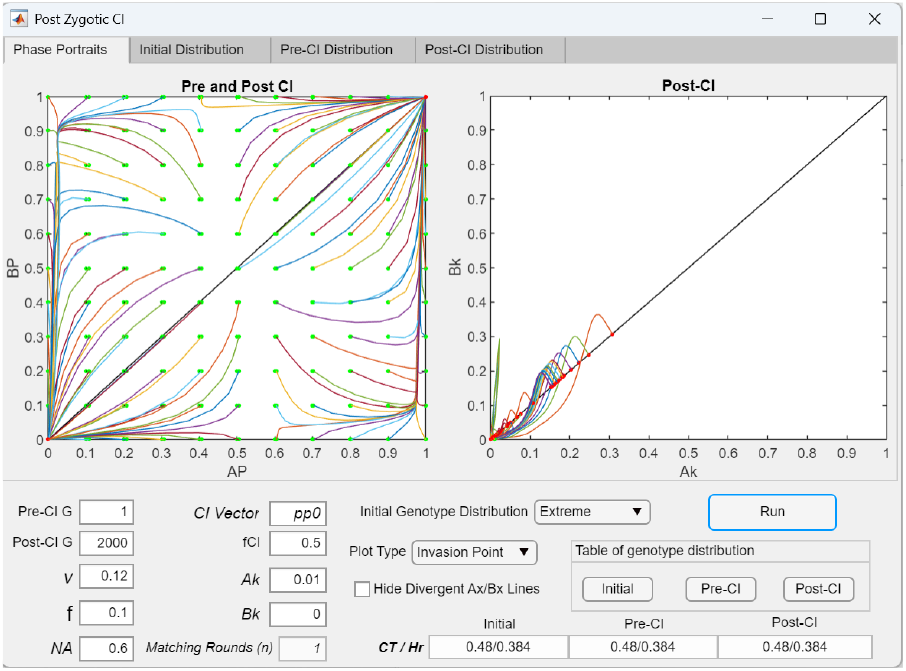
The presence of CIs capturing extreme-type incompatibility alleles reduces the effective number of incompatibility gene loci and impedes convergence to fixed points and the establishment of postzygotic RI. The parameter values used in Fig 54a permit the emergence of a fixed point in the lower-right quadrant of the *AP*/*BP* phase portrait in the absence of CIs, but this fixed point fails to form when a CI capturing extreme-*P* alleles is present in the initial population. As shown in the *Invasion Point* plot, when setting *Pre*-*CI G* to 1, the CI mutant (CI = *pp*0, *Ak* = 0.01) in the population rapidly invades, reduces the effective number of incompatibility gene loci experienced by individuals in the system, and prevents convergence to a fixed point in the lower-right quadrant of the *AP*/*BP* phase portrait.

### IV. DISCUSSION

In modern genomic research, chromosomal inversions (CIs) are recognized as ubiquitous structural features across the genomes of many organisms and are increasingly viewed as important contributors to local adaptation and speciation [1-9]. Although CIs have been studied since the 1920s and numerous theoretical models have explored their role in speciation [4, 5, 18, 28-33]—primarily through their influence on local adaptation, reduction of gene flow, and the evolution of reproductive isolation (RI) between diverging populations—their origins and subsequent interactions with other speciation processes remain insufficiently understood [22, 27]. In particular, whether CIs impede or facilitate speciation through interactions with ecological and reproductive barrier variables remains unresolved [37, 38]. In this study, we aim to narrow this knowledge gap by analyzing the potential role of CIs in sympatric speciation using mathematical modeling and computer-based simulations.

In this study, we focus primarily on the role of chromosomal inversions in speciation as recombination suppressors [17, 18]. During meiosis in heterokaryotype matings, the inverted and noninverted chromosomal segments pair through the formation of an inversion loop, which inhibits the breakdown of linked allele combinations within the inverted region [37]. Rare single crossover events within the loop can still occur and produce 50% nonviable offspring [22, 40]. Recombination resistance may also be eroded by limited gene flux resulting from rare double crossover events or gene conversion [41]. However, because these processes are rare and lack compelling empirical support for their significance in speciation, we do not incorporate them into our analyses. Instead, we model the recombination-suppressive effect of CIs by assigning a value of 0.5 to *fCI*, the offspring return ratio between a CI and its corresponding uninverted colinear region in heterokaryotype mating. While *fCI* could be set to a value below 0.5 to account for offspring loss due to single crossover events, we adopt *fCI* = 0.5 to isolate the primary effect of recombination suppression. A value of *fCI* < 0.5 may become important for preventing invasion by uninverted same-niche genotypes once the CI is the predominant genotype within a polymorphism [38]. Other potential mechanisms, such as breakpoint-induced gene disruption or reduced hybrid viability in diploid heterokaryotypes, are also excluded from our model because of limited empirical evidence supporting their importance in speciation.

A CI mutation gains its fitness advantage by capturing favorable combinations of locally adaptive alleles and preventing their dissociation through recombination. It can then effectively function as a supergene that integrates the properties of the alleles it captures. Previous studies and simulations have shown that this advantage from suppressing recombination is strongest in high–gene-flow environments where adaptive traits are mediated by many alleles of small effect rather than a few alleles of large effect [5]. Consequently, CIs are well suited to act as an effective mechanism in sympatric speciation, where gene flow remains substantial and most ecological and barrier traits are polygenic.

In a prior study, we developed a mathematical model of a sympatric population under disruptive ecological selection [39], illustrated in Fig 3. The model consists of two ecological niches (*A* and *B*), each capable of supporting a distinct adaptive ecotype, but it does not include hybrid niche resources that would allow viable ecological hybrids to exist. Individuals carry one of two mating-bias alleles (*X* or *Y*) at a single locus, which determines mating compatibility.

In the present study, we extend this model by developing GUI applications that incorporate multilocus ecological, mating-bias, and hybrid-incompatibility genotypes to investigate how polygenic trait architectures influence system dynamics. Within this polygenic framework, we evaluate the invasion fitness and evolutionary effects of CIs that capture different combinations of alleles from these genotypes.

In the model incorporating polygenic ecological traits, we introduce hybrid niche resources capable of supporting viable ecological hybrids and examine how their presence influences population dynamics. For simplicity and analytic tractability, individuals retain the original one-locus, two-allele (*X* or *Y*) representation of mating-bias genotypes to implement sexual selection. Similarly, individuals carry a modified one-locus-equivalent, two-allele (*P* or *Q*) representation of hybrid-incompatibility genotypes to implement hybrid-incompatibility selection.

In the model with polygenic mating-bias traits, we retain the ecological structure of the original framework in Fig 3 to implement ecological selection. In this case, the system contains two parental ecotypes without viable ecological hybrids, and the strength of disruptive ecological selection is determined by the offspring return ratio *f*.

Likewise, in the model with polygenic hybrid-incompatibility traits, we use the same ecological framework in Fig 3 to implement ecological selection, with the strength of disruptive ecological selection again specified by *f*. The results of our study are discussed in the following sections.

### SYSTEMS WITH MULTILOCUS ECOLOGICAL GENOTYPES

#### 1 The Effect of Having Multilocus Ecological Genotypes and Viable Ecological Hybrids (Without CI Invasion)

##### 1.1 The presence of viable ecological hybrids weakens disruptive ecological selection and reduces RI

Our GUI application enables users to specify the carrying capacities of hybrid niche resources that can sustain viable ecological hybrid genotypes, as defined in the Habitat Resource Distribution Table (Fig 9). Because the model assumes that each generation produces enough offspring to occupy all available niche resources, the normalized population ratios of the ecological genotypes are fixed by their corresponding niche sizes at the start of every mating generation. Consequently, the only changes that may occur within each genotype population over time are the ratios of the *X* and *Y* mating-bias alleles.

Our simulations consistently show that the presence of viable ecological hybrids increases the effective value of *f* (the offspring return ratio) experienced by the parental niche-*A* and niche-*B* ecotypes in Fig 3, thereby weakening the disruptive ecological selection required to produce strong RI (Fig 8 and Fig 21). This occurs because viable hybrids can mate and regenerate parental genotypes, which reduces the amount of maladaptive hybrid loss that results from inter-niche matings. When the hybrid population becomes sufficiently large, the system becomes divergent, and no *Ax*/*Bx* fixed point exists in the phase portrait.

##### 1.2 Increasing the number of ecological gene loci strengthens disruptive ecological selection and increases RI

Our GUI application (MultiEcoCI) allows users to specify any number of gene loci for the ecological genotypes. When hybrids are maladaptive or when hybrid niche resources are limited, our simulations show that increasing the number of ecological loci lowers the effective value of *f* experienced by the parental niche-*A* and niche-*B* ecotypes in Fig 3, which strengthens the disruptive ecological selection needed to produce strong RI (Fig 8 and Fig 22). This occurs because offspring genotypes are formed by the random assortment of niche-*A* (*a*-type) and niche-*B* (*b*-type) alleles at each locus. As the number of loci increases, a greater proportion of offspring become ecological hybrids rather than the parental genotypes (e.g., *aaaa* or *bbbb*). Because most maladaptive hybrid genotypes are eliminated, the offspring return ratio *f* decreases, which intensifies the disruptive ecological selection acting on the parental ecotypes. This result is consistent with previous studies [49].

Theoretically, when disruptive ecological selection is weak and most hybrids are viable, reducing the number of loci underlying ecological genotypes can limit the production of viable hybrids, thereby reducing gene flow and introgression. This effect has been demonstrated in simulations of two-island models, in which selection against maladaptive immigrants plays an important role in maintaining ecological divergence [50]. However, in our simulations of sympatric populations, almost all systems with weak ecological selection remain divergent and fail to establish premating RI even when the number of ecological loci is reduced.

#### 2 Invasion of a CI in a Two-Niche, Two-Mating-Bias-Allele System with Multilocus Ecological Genotypes under Disruptive Ecological Selection

In our models, CI invasion is considered an adaptive process driven by maladaptive hybrid loss. For a CI mutation to successfully invade a niche, individuals carrying the inversion must have higher fitness than same-niche individuals without it. A CI can acquire this fitness advantage by capturing locally adaptive ecological alleles. Because the favorable allele combinations within a CI cannot be broken up by recombination, individuals carrying the CI experience less hybrid offspring loss in inter-niche matings and gain a fitness advantage over their same-niche uninverted counterparts. However, when premating RI is already strong and inter-niche hybrid loss is minimal, this advantage disappears and a CI cannot invade.

Understanding this mechanism allows us to evaluate CI invasion outcomes in our models, both when the inversion captures only ecological alleles and when it captures ecological alleles together with a mating-bias allele.

##### 2.1 Invasion of a CI capturing only locally adaptive ecological alleles weakens RI by reducing disruptive ecological selection

In a sympatric population under disruptive ecological selection, a CI mutation that captures locally adaptive ecological alleles gains enhanced invasion fitness because individuals carrying the inversion produce fewer maladaptive hybrids in inter-niche matings than uninverted individuals from the same niche. However, when the CI captures all locally adaptive ecological alleles, maladaptive hybrid offspring are no longer produced in inter-niche heterokaryotype matings, and the effective value of *f* experienced by ecotypes with the CI becomes 0.5. Once the CI rises to near fixation, there is no hybrid loss to drive or maintain the non-random assortment of mating-bias alleles, and the *Ax*/*Bx* phase portrait becomes divergent. Consequently, one of the two mating-bias alleles (*X* or *Y*) is eliminated, and premating RI collapses (Figs 23a–23c).

In the real world, this scenario would produce two predominant ecological morphs in a sympatric population, each adapted to either the *A* or *B* niche. Invasion of a CI that captures all locally adaptive ecological alleles effectively combines those alleles into a supergene that is protected from recombination and segregates as a single Mendelian locus. The two ecotypes can interbreed freely, but no intermediate hybrid offspring are produced. The ecological alleles contained within the inversion remain isolated between the niche ecotypes, while the rest of the genome recombines freely. Because there is no maladaptive hybrid loss, high-mating-bias alleles cannot invade, making the evolution of premating RI—and thus sympatric speciation—impossible.

A likely explanation for why this outcome is not observed more frequently in nature is that genes mediating adaptive polygenic traits may be scattered far apart on the same chromosome or located on different chromosomes, making it impossible for a single CI to capture all of them [39].

When an invading CI captures only a subset of locally adaptive alleles, maladaptive hybrids are again produced in inter-niche matings. The resulting hybrid loss restores disruptive ecological selection and allows fixed points to re-emerge in the *Ax*/*Bx* phase portrait. However, because hybrid loss is reduced, the premating RI that develops is weaker (Figs 24a and 24b). This occurs because the CI reduces the effective number of ecological loci experienced by the system, making convergence more difficult and producing weaker RI. When a CI captures only a single ecological allele, it gains no fitness advantage in inter-niche matings and is unable to invade (Fig 25).

In general, capturing more locally adaptive ecological alleles gives a CI greater invasion fitness, enabling it to invade more rapidly and reach a higher steady-state frequency within the niche (Figs 23a and 24a). However, capturing more adaptive alleles also produces fewer hybrid offspring in inter-niche matings, which reduces hybrid loss and weakens RI (Figs 23b and 24a). This negative coupling effect may contribute to genomic patterns analogous to those produced by the interchromosomal effect [2, 51, 52], in which inversions suppress recombination within the inverted region and increase it elsewhere in the genome. Such a recombination landscape can also generate a genomic island of divergence, where divergence is concentrated within the inversion while the remainder of the genome remains more homogenized by gene flow.

##### 2.2 Invasion of a CI capturing all locally adaptive ecological alleles and a mating-bias alleles produces RI depending on relative niche sizes

We have shown that in a sympatric population under disruptive ecological selection, a CI that captures all locally adaptive ecological alleles acquires a fitness advantage and can successfully invade. When such a CI also captures the locally prevalent mating-bias allele in its adaptive niche (e.g., CI = *aaaX*), it can carry that mating-bias allele to near fixation within the niche through hitchhiking.

However, because inter-niche heterokaryotype matings produce no maladaptive hybrids in this scenario, hybrid loss cannot contribute to premating RI. Instead, sexual selection and the relative sizes of the two niches play key roles in determining the invasion outcome. As illustrated in Fig 26 and Fig 28, when *NA* ≥ *NB*, invasion of a CI that captures all niche-*A* ecological alleles and the *X* allele (CI = *aaaX*) causes the *Ax*/*Bx* phase portrait to become divergent, ultimately eliminating the *Y* allele and destroying the preexisting RI. A likely explanation is that because the *X* allele cannot be separated from the niche-*A* ecotype, its larger population size in the sympatric system allows it to outcompete the smaller *Y*-allele population through sexual selection.

In contrast, when niche *B* is larger than niche *A*, the outcome differs. As shown in Fig 29a, when *NB* > *NA*, the larger *Y*-allele population in niche *B* cannot eliminate the minority *X*-allele population in niche *A* because the *X* allele in niche *A* is linked to the highly fit CI genotype (CI = *aaaX*). Consequently, invasion of the CI can maintain a fixed-point polymorphism in the *Ax*/*Bx* phase portrait and preserve premating RI. Nonetheless, when the mating-bias parameter *α* is relaxed beyond a threshold (*α* > 3.1 in Fig 29), the increased inter-niche mating gives the *X* allele in niche *B* enough fitness advantage to eliminate the *Y* allele, causing the system to become divergent.

These results suggest that when a CI captures all locally adaptive ecological alleles and the locally prevalent mating-bias allele, an *Ax*/*Bx* fixed-point polymorphism—and therefore premating RI—can be maintained only when *NB* > *NA* and under a restricted range of strong mating bias.

Finally, the post-CI genotype distribution in Fig 29b indicates that the CI (*aaaX*) cannot invade the hybrid niches. As a result, viable hybrids can mate and continuously regenerate the uninverted parental ecotypes, preventing CI fixation in niche *A* and maintaining a polymorphism between inverted and uninverted genotypes.

##### 2.3 A CI capturing a partial set of locally adaptive ecological alleles and a high–mating-bias allele acquires high invasion fitness, enabling successful invasion regardless of mating bias and allowing RI to evolve independent of niche sizes

When a CI mutant captures a subset of locally adaptive ecological alleles together with a locally favored mating-bias allele, it acquires high invasion fitness and can invade to establish premating RI regardless of niche sizes. In this scenario, both the production of maladaptive hybrids and disruptive ecological selection are restored, and hybrid loss again becomes available to drive the invasion of high– mating-bias alleles, thereby generating fixed-point polymorphisms and premating RI. The anchoring of the high–mating-bias allele to locally adaptive ecological alleles effectively creates a “pseudomagic trait” that is both resistant to the homogenizing effects of gene flow—because the linked mating-bias allele cannot be transferred to the opposite ecotype—and has high fitness in its adaptive niche due to its linked ecological alleles [35, 36]. Together, these properties confer high invasion fitness to the CI.

As illustrated in Figs 30a and 30b, a CI mutation (*CI* = *aa*0*X*) can invade both when *NA* > *NB* and when *NB* > *NA*. Fig 31a provides an example in which a CI capturing only a single locally adaptive allele in niche *A* has no invasion advantage on its own; however, when it also captures the mating-bias allele *X*, this CI (CI = *a*00*X*) acquires synergistic fitness through their linkage and invades to reach fixation in all genotype niches that contain the captured ecological allele (Fig 31b).

In comparison, even though a CI capturing more ecological alleles has greater invasion fitness—i.e., a shorter *T*_95_ (Figs 31a, 31c, and 32)—its invasion produces weaker RI because it tends to persist in polymorphism with same-niche uninverted ecotypes (Fig 31c). This occurs because CIs that capture more ecological alleles can invade fewer hybrid genotypes, which allows more uninverted hybrids to continue regenerating uninverted parental ecotypes. In addition, the invasion of such CIs reduces the hybrid loss (*HL*) available to drive the invasion and coupling of other high–mating-bias alleles, since capturing more ecological alleles decreases the production of maladaptive hybrid offspring in inter-niche heterokaryotype matings.

These outcomes are predicted by the dynamics of the model in Fig 3, in which ecological selection and sexual selection act as complementary forces in generating system convergence and premating RI [39]. Lower values of *f* and *α*, corresponding to stronger disruptive ecological selection and stronger mating bias, reinforce one another to produce stronger premating RI. Consequently, invasion of a CI that captures a larger number of locally adaptive ecological alleles increases the effective value of *f* experienced by the ecotypes, thereby weakening premating RI for a given mating-bias strength *α*.

Although postmating barriers are not explicitly modeled in this system, previous work has shown that mutations causing intrinsic hybrid incompatibilities— which reduce *f* but are underdominant and cannot invade on their own—can nevertheless invade when linked to high mating-bias alleles [38]. Such linkage can create a positive feedback loop between prezygotic and postzygotic barriers to gene flow [38, 53]. In addition, increased hybrid loss can further strengthen premating RI through reinforcement, while stronger premating RI reduces inter-niche mating, slows the purging of alleles causing hybrid incompatibilities, and may allow additional or stronger hybrid incompatibilities to evolve [38]. As a result, a mutation possessing both premating and postmating barrier properties can achieve high invasion fitness and generate strong RI due to the jointly reduced values of *f* and *α* experienced by genotypes carrying the mutation.

Even when the pre-CI system is divergent, as shown in Fig 31d, both the CIs in Figs 31a and 31c (CI = *aa*0*X* and CI = *aa*0*X*) can invade and create a fixed-point *Ax*/*Bx* polymorphism and premating RI.

In our simulations, CI invasion succeeds whether the mutant arises in a system already at a pre-CI fixed point (Fig 31a) or functions as a magic trait invading a system that lacks such a fixed point (Figs 31d and 32). Remarkably, in all of these scenarios, the high invasion fitness of the CI enables successful invasion across the entire range of mating-bias strengths 0 < *α* < 1.

In summary, our simulation results show that a CI mutant capturing locally adaptive ecological alleles together with a locally favored high–mating-bias allele acquires high invasion fitness [54], which enables it to invade regardless of niche sizes or whether the system is convergent or divergent. Moreover, a CI that captures more locally adaptive alleles attains higher invasion fitness and invades more rapidly (lower *T*_95_) but ultimately reaches a lower equilibrium frequency and produces weaker RI (higher *CI*) together with reduced residual hybrid loss (lower *HL*).

##### 2.4 A CI that captures a partial set of locally adaptive ecological alleles and a locally adaptive high-mating-bias allele has higher invasion fitness than an otherwise identical CI lacking the mating-bias allele

A CI that captures a partial set of locally adaptive ecological alleles gains additional invasion advantage when it also captures a locally advantageous high-mating-bias allele. Together, these alleles increase both the CI’s invasion fitness and the steady-state frequency it ultimately achieves within the niche (Fig 34a–34d). This pattern indicates that, once a CI has acquired locally adaptive ecological alleles, positive selection favors the subsequent acquisition of a high-mating-bias mutation within that CI background.

Conversely, CI variants that capture alleles maladaptive in a given niche—such as ecological alleles originating from the wrong habitat (e.g., CI = *a*0*bX* in Fig 33) or a mating-bias allele that is not locally favored (e.g., CI = *aa*0*Y* in Fig 34e)— experience reduced fitness and fail to invade.

##### 2.5 Viable ecologically hybrids can reproduce uninverted parental ecotypes that cause an adaptive CI to exist in a polymorphism rather than fixation, while hybrid swamping can eliminate the CI entirely

In the model without viable ecological hybrids (Fig 3), a locally adaptive CI consistently rises to fixation [39]. In contrast, our current model allows viable ecological hybrids to form and persist, and these hybrid genotypes can mate and regenerate uninverted parental ecotypes that prevent the adaptive CI from reaching fixation in its niche. Instead, the invading CI persists in a polymorphism with uninverted genotypes. This polymorphism is maintained by an immigration– selection balance in which the less-fit uninverted genotypes are eliminated by selection favoring CI genotypes yet are continuously replenished by the uninverted hybrid populations (Figs 34a–34d).

In the extreme case, when the hybrid population is sufficiently large, hybrid swamping can supply so many uninverted genotypes to the polymorphism that it eliminates the CI population (Fig 35a–35c).

### SYSTEMS WITH MULTILOCUS MATING-BIAS GENOTYPES

#### 1 The Effect of Having Multilocus Mating-Bias Genotypes (Without CI Invasion)

We developed a MATLAB GUI application (MultiSexCI) capable of modeling any specified number of mating-bias gene loci to examine how multilocus mating-bias genotypes influence the dynamics of the model in Fig 3. In this polygenic framework, mating biases between genotypes vary linearly with genetic distance, as illustrated in Fig 15, and the maximum bias between extreme genotypes is specified by the parameter *α*. Ecological selection follows the same framework used in Fig 3, where the strength of disruptive ecological selection is represented by the offspring return ratio *f*, and the system contains no viable ecological hybrids.

Our previous analyses of the Fig 3 model showed that, just as viable ecological hybrids can reduce the strength of disruptive ecological selection and impede system convergence, the presence of mating-bias hybrids can likewise weaken sexual selection and hinder both convergence and the emergence of premating RI [39]. However, whereas ecological hybrid populations are constrained by the carrying capacities of hybrid-specific niches, mating-bias hybrids associated with parental ecotypes in the polygenic model are unconstrained because all such hybrids occupy the same ecological niche and are fully viable. Consequently, the population dynamics become more complex and more sensitive to initial conditions.

Despite these complications, systems with multiple mating-bias loci can still converge and generate premating RI under favorable parameter conditions, driven by selection on niche ecotypes to reduce maladaptive ecological hybrid loss arising from inter-niche matings. When convergence occurs, our simulations consistently show at most a single fixed-point peak in the multi-dimensional phase portrait of mating-bias genotypes. This fixed-point peak may or may not be visible in the lower-dimensional *AX*/*BX* phase portrait—depending on whether it involves extreme genotypes or consists solely of hybrid genotypes—but it can always be identified in the post-CI genotype distribution.

Two parameters primarily determine the breadth and shape of the post-CI genotype distribution and influence whether convergence occurs: the number of mating-bias gene loci and the maximum mating bias *α*. In the sections below, we describe how these two parameters determine system convergence and the resulting strength of RI. We also analyze the influences of additional system parameters, including the strength of ecological selection (*f*), niche size (*NA*), and the number of matching rounds (*n*).

##### 1.1 Increasing the number of mating-bias loci decreases the effective strength of sexual selection, leading to weaker RI or system divergence

In our models, the presence of mating-bias hybrids is just as detrimental to system convergence as the presence of ecological hybrids. It follows that increasing the number of mating-bias loci results in more mating-bias hybrids being produced by matings between the extreme genotypes (e.g., *xxxx* and *xxxx*). This is reflected by an increase in the number of hybrid genotypes in the post-CI genotype distribution graph, thereby hindering the establishment of premating RI. This result is consistent with previous studies [49, 55].

Our simulations confirm that in a convergent system, where a fixed-point polymorphism already exists in the *AX*/*BX* phase portrait, adding more mating-bias loci tends to shift the fixed point away from the lower-right corner of the phase portrait and results in weaker RI (Figs 36a and 36b). This reduction in RI is reflected by increases in the fixed-point values of *CI*, the ratio of total offspring produced from inter-niche matings, and *Hr*, the ratio of hybrid offspring produced from inter-niche matings. Because the model does not allow viable hybrids, *Hr* is equivalent to the ratio of maladaptive hybrid loss.

As the number of mating-bias loci continues to rise, an invasion-resistant pattern appears in the *AX*/*BX* phase portrait (Fig 36c), preventing small mutant populations near the origin from invading. Eventually, when the number of loci increases beyond a threshold, the system becomes globally divergent (Fig 36d): the *AX*/*BX* phase portrait contains no fixed points, and premating RI cannot form.

##### 1.2 Increasing the mating bias between extreme mating-bias genotypes facilitates system convergence and produces stronger RI

Increasing the mating bias between the extreme genotypes—achieved by reducing the value of *α*— facilitates convergence of the system to fixed points and helps to eliminate the invasion-resistance pattern observed in the *AX*/*BX* phase portrait (Fig 36e). In convergent systems, further decreases in *α* shift the *AX*/*BX* fixed point toward the lower-right corner of the phase portrait and produce progressively stronger RI (Figs 36c and 36e-36g).

Fig 36f illustrates a representative pre–CI genotype distribution for a system with multilocus mating-bias genotypes. The bar graph shows the normalized ratios of all genotypes ordered by genetic distance, with the two extreme genotypes positioned at opposite ends. When the *AX*/*BX* phase portrait is divergent, the niche-*A* and niche-*B* ratios of each genotype converge toward equality as the population vector moves toward the diagonal of the phase portrait. Once on the diagonal line, sexual selection subsequently drives the population ratios toward opposite ends of the line, where all but one extreme genotype are eliminated.

In contrast, when the *AX*/*BX* phase portrait converges to a stable fixed point, the genotype distribution shown in Fig 36f emerges. Under these conditions, nonrandom assortment between the two extreme genotypes causes them to segregate into different niches, where each becomes the predominant genotype. Intermediate hybrid genotypes exhibit skewed distributions toward their like-kind parental extreme within each niche, producing a concave pattern across the genotype spectrum. As the *AX*/*BX* fixed point moves closer to the lower-right corner of the phase portrait—signaling stronger RI—this concavity becomes increasingly pronounced, pushing hybrid genotype frequencies toward their corresponding parental extremes. In the limiting case where *α* = 0, mating bias is maximal and all intermediate hybrid genotypes are eliminated, leaving only the two extreme mating-bias genotypes segregated across the niches (Fig 36g).

To illustrate how such a genotype distribution might arise in nature, imagine a fish species in which mating discrimination is based on body color encoded by multiple loci. Suppose the multilocus genotype *yyyy* produces a blue phenotype, whereas *xxxx* produces a red phenotype. Crosses between these extreme genotypes yield hybrids expressing intermediate purple hues, with coloration biased toward red or blue depending on genetic proximity to the parental extremes. Mate choice is governed by color similarity through a matching-compatibility rule analogous to the table shown in Fig 15. As premating RI strengthens, assortative mating increasingly partitions the population into sharply defined red and blue ecotype groups, with reddish-purple and bluish-purple hybrids progressively displaced toward their respective parental red and blue extremes. Conversely, weak premating RI produces a broader spread of color variants and a less concave distribution of genotypes across the red–blue spectrum.

##### 1.3 Stronger disruptive ecological selection, a higher number of matching rounds, and roughly equal niche sizes facilitate system convergence and produce stronger RI

Our simulations show that strengthening ecological selection—implemented by reducing the value of *f*—has an effect similar to increasing the maximum mating bias, *α* (Fig 36h). This result is expected because, in our model (Fig 3), ecological selection (governed by *f*) and sexual selection (determined by *α* and *n*) act complementarily to promote system convergence and the establishment of premating RI [39].

Likewise, increasing the number of matching rounds (*n*) reduces intra-niche sexual selection against rare high–mating-bias mutants, contracts the invasion-resistant region in the phase portrait, and facilitates system convergence (Fig 37a). This pattern is consistent with results from our single-locus model in Fig 3 [39].

Finally, in systems with multiple mating-bias loci, convergence is most readily achieved when niche sizes are equal, i.e., when *NA* = *NB* = 0.5 (Fig 37b). When an invasion-resistant region is present in the *AX*/*BX* phase portrait, its size is minimized when *NA* = 0.5 and increases as *NA* deviates from this value. Beyond a symmetric range centered around *NA* = 0.5, the system becomes divergent. A possible explanation is that inter-niche matings and the production of maladaptive hybrids are maximized when niche sizes are equal, creating the greatest amount of hybrid loss to drive the evolution of mating-bias barriers.

#### 2 Invasion of a CI in a Two-Niche System with Multilocus Mating-Bias Genotypes

We next examine the invasion dynamics of CIs that capture mating-bias alleles in a two-niche model with multilocus mating-bias genotypes. In general, for evolution to occur, some less-fit variants in the population need to be eliminated so that other more-fit alternatives can rise in frequency in the gene pool. In our model of a sympatric population under disruptive ecological selection, the unfit variants are maladaptive hybrid offspring produced during inter-niche matings between locally adapted ecotypes. This maladaptive hybrid loss is what drives the invasion of an adaptive CI capturing a favorable combination of mating-bias alleles. Such a CI derives its invasion fitness from reducing the effective number of mating-bias loci in the system, which in turn reduces ecological hybrid loss by increasing premating RI between niche ecotypes. Consequently, if there is no hybrid loss, the CI has no invasion fitness and cannot invade.

In our simulations, we assume that an invading CI mutation arises in a small population of extreme-*X* genotypes in niche *A*, within a population otherwise composed entirely of the extreme-*Y* genotype. For instance, the initial genotype of a mutant population containing a CI mutation (CI = *xx*00) capturing the first two *x*-type mating-bias alleles in niche *A* is 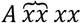, where the overbrace denotes the linked alleles. Because all mating-bias hybrids are ecologically viable, their relative frequencies within a niche population are not constrained by niche resources. This makes the population dynamics of systems with multiple mating-bias loci both more complex and more difficult to predict, and more sensitive to the initial frequency and composition of the invading CI genotype.

Finally, for comparison, we analyze the less common case in which a CI mutant arises in a population composed solely of extreme-*Y* genotypes, as could occur if such a CI is donated through adaptive introgression in a secondary contact scenario. In a system with four mating-bias loci, an invading genotype carrying such a CI that captures the first two x-type mating-bias alleles (CI = *xx*00) would then take the form 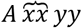 rather than 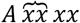.

##### 2.1 Invasion of a CI capturing favorable mating-bias alleles promotes system convergence and premating RI by reducing the effective number of mating-bias loci

A CI capturing a favorable combination of high mating-bias alleles derives its invasion fitness from reducing the effective number of mating-bias gene loci, increasing premating RI between niche ecotypes, and limiting the production of maladaptive ecological hybrids. Therefore, in the absence of hybrid loss (i.e., *Hr* = 0), such a CI cannot invade (Fig 38).

However, when premating RI is incomplete, hybrid loss may then permit a CI capturing all extreme-*X* alleles to invade a population otherwise composed entirely of extreme-*Y* genotypes, leading to the establishment of an *AX*/*BX* fixed-point polymorphism and premating RI. This can occur because invasion of the CI reduces the effective number of mating-bias gene loci to two and facilitates system convergence. This is illustrated in Figs 39a and 39b. When an invasion-resistant pattern emerges in the *AX*/*BX* phase portrait as the number of mating-bias gene loci increases from two to three (Fig 39a), invasion of a CI capturing all extreme-*X* alleles (CI = *xxx*) restores the effective number of mating-bias gene loci to two (Fig 39b), allowing invasion of the rare 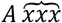 genotype and the establishment of premating RI.

##### 2.2 Increasing the maximum mating bias (α) increases the invasion fitness of CIs capturing partial sets of extreme mating-bias alleles and promotes fixation at extreme genotypes

In our simulations, an invading CI mutation captures either a complete or a partial subset of mating-bias alleles from a small population of extreme-*X* genotypes in niche *A*, within a population otherwise fixed for the extreme-*Y* genotype (Fig 40). Consequently, when a CI captures only a partial set of extreme-*X* alleles, the initial mutant population still expresses the extreme-*X* genotype (e.g., 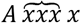 in a four-locus genotype, where the overbrace denotes the captured *x*-type alleles).

In general, our results confirm that the more extreme-*X* alleles a CI captures, the greater its invasion fitness. As demonstrated in Fig 41a, a CI capturing three of four extreme-*X* mating-bias alleles (CI = *xxx*0) can invade a divergent system and produce convergence and premating RI. This is possible because the CI reduces the effective number of mating-bias loci, thereby enabling convergence. In contrast, a CI capturing only two of four extreme-*X* mating-bias alleles (CI = *xx*00) is not able to invade the same divergent system (Fig 41d).

An invading CI’s uncaptured gene loci determine the possible CI genotypes that can exist in the post-CI genotype distribution. For instance, in Fig 41a, following the invasion of the CI (CI = *xxx*0), the only genotypes that can carry the CI are *xxxx* and *yyyy*. Invariably, the successful invasion and fixation of the CI in these two genotypes can then reduce the effective number of mating-bias loci from three to at most two (Fig 41c). Similarly, after invasion and fixation of the CI that captures two out of four gene loci (CI = *xx*00), the maximum number of effective gene loci is reduced to three, with eight possible genotypes in the post-CI distribution (Fig 41e).

As shown in Fig 41b, for CI = *xxx*0 and an initial invading genotype of 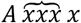, the CI does not always fix at the extreme-*X* genotype in niche *A*. Instead, when *α* = 0.3, the CI fixes at the hybrid mating-bias genotype *Axxxy*. A plausible explanation is that, because niche *A* is initially composed entirely of the extreme-*Y* genotype, CI introgression via intra-niche mating rapidly establishes a sizable *Axxxy* population in niche *A* during the early phase of invasion. As *Axxxy* rises in frequency, weak mating bias between *Axxxy* and *Axxxy* promotes increased intra-niche mating between these genotypes, reducing inter-niche matings between *Axxxx* and *Byyyy* that are essential for maximizing the invasion fitness of the extreme-*X* genotype. At *α* = 0.3, the mating bias (*α*) favoring *Axxxx* is insufficient to compensate for its reduced inter-niche matings, allowing the numerically dominant *Axxxy* genotype to achieve higher net fitness through more frequent inter-niche matings despite its weaker inter-niche mating bias. Consequently, *Axxxy* and the *x* allele at the fourth locus are eliminated early in the invasion.

In this case, although the effective number of mating-bias loci is reduced to one (Fig 41b), this reduction comes at the cost of a weaker mating bias between *Axxxy* and *Byyyy* than the maximum value *α* between the extreme-*X* and extreme-*Y* genotypes, resulting in weaker premating RI than would occur if fixation were at the extreme-*X* genotype.

Similar effects occur when the CI captures two of four extreme-*X* alleles (CI = *xx*00). Such a CI cannot invade a divergent *AX*/*BX* system when *α* = 0.3 (Fig 41d), but reducing *α* to 0.2 allows invasion and fixation at the hybrid genotype *Axxyy*, while further reducing *α* to 0.1 leads to fixation at the extreme-*X* genotype *Axxxx* and stronger premating RI (Fig 41e). Together, these results show that invasion outcomes for CIs capturing partial sets of extreme mating-bias alleles depend on both the composition of the invading CI genotype and the maximum mating-bias value *α*.

##### 2.3 A CI capturing more extreme mating-bias alleles acquires greater fitness to invade a convergent system with multiple mating-bias loci, resulting in stronger RI

A CI capturing only a single mating-bias allele gains no invasion advantage and cannot invade (Figs 42a and 42b). In contrast, in a convergent system that already exhibits a fixed point in the *AX*/*BX* phase portrait, a CI capturing more than one extreme mating-bias allele can consistently invade and further strengthen existing RI by reducing the effective number of mating-bias gene loci (Figs 42c–42e). Furthermore, CIs capturing more favorable extreme mating-bias alleles gain greater invasion fitness—as reflected by shorter invasion times (*T*_95_) and higher steady-state fixation frequencies—and produce stronger RI following invasion (Figs 42c–42e).

If the pre-CI phase is omitted (*Pre*-*CI G* = 1) and the post-CI *X*/*BX* phase portrait is globally convergent (Fig 43b), a CI capturing extreme mating-bias alleles can invade from anywhere in the phase portrait—i.e., from the origin or from any established fixed point—by reducing the effective number of mating-bias gene loci and thereby enhancing RI (Figs 43a–43c).

##### 2.4 CIs capturing mating-bias alleles in a small extreme-X population can invade a divergent system and generate fixed points and premating RI when the initial AX/BX ratios fall within a permissible region of the phase portrait

As shown in Figs 44a–44i, a wedge-shaped region extending from the origin along the *x*-axis of the *AX*/*BX* phase portrait defines the set of initial *AX*/*BX* population ratios that permit invasion by a CI arising in the *AX* population. When the initial population ratios fall within this region, the CI can invade and convert an otherwise divergent system into a convergent one with stable fixed points and premating RI. The invasion fitness of the CI increases with the number of mating-bias alleles it captures, resulting in stronger premating RI and an expansion of the region of *AX*/*BX* ratios that permit invasion. By contrast, when the initial *AX*/*BX* population ratios lie outside this region, the CI fails to invade. Consequently, the system remains divergent, and sexual selection ultimately eliminates all genotypes except the extreme-*Y* genotype (Figs 44h and 44i). Together, these results show that CI invasion dynamics in systems with multiple mating-bias loci are highly nonlinear and sensitive to initial population composition.

##### 2.5 CIs capturing x-type mating-bias alleles arising within an extreme-Y population without pre-existing extreme-X genotypes can invade a divergent system by reducing the effective number of mating-bias loci to one

In contrast to a CI arising from a small extreme-*X* subpopulation and invading as an extreme-*X* genotype (e.g., CI = *xx*00 with invading genotype 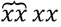), a CI capturing *NN*-type mating-bias alleles and originating within an extreme-*Y* population without pre-existing extreme-*X* genotypes necessarily carries *NN*-type alleles at its uncaptured loci (e.g., CI = *xx*00 with invading genotype 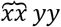). In the *AX*/*BX* phase portrait, such a CI therefore arises at the origin rather than from an *AX* population displaced from the origin. Because no x-type alleles are present at the uncaptured loci in the population, invasion by this class of CI invariably reduces the effective number of mating-bias loci to one (Figs 45a, 45b, and 45d). As a result, these CIs have very high invasion fitness, as they can always invade and reduce the effective number of mating-bias loci of any system to one, thereby facilitating convergence (Fig 45d).

When the CI captures only a partial set of the extreme-*X* alleles, the mating bias between the resulting CI hybrid genotype and the extreme-*Y* genotype is less than *α*. Consequently, the premating RI generated is weaker than that produced by CIs capturing the full complement of extreme-*X* alleles (Figs 45d and 45e). When *AX* genotypes are present in the population, invasion by such CIs also remains constrained by a permissible region of initial *AX*/*BX* population ratios in the *AX*/*BX* phase portrait (Fig 45c). From a mechanistic perspective, the origin of such CIs through mutation within an extreme-*Y* population appears unlikely and is more plausibly explained by introgression from an allopatrically diverged population during secondary contact.

### SYSTEMS WITH MULTILOCUS HYBRID-INCOMPATIBILITY GENOTYPES

We adapted previously developed GUI applications and further modified those created in this study to investigate population dynamics in a two-allele model of postzygotic hybrid incompatibility (see Methodology). These modifications also allow us to examine the invasion dynamics of CIs in models that incorporate polygenic traits.

#### 1 Single-Locus-Equivalent Two-Allele Model of Hybrid Incompatibility

##### 1.1 Low gene flow can maintain divergence in hybrid-incompatibility alleles across niches and sustain postzygotic reproductive barriers when niche ecotypes harbor pre-established genetic incompatibilities

In speciation genetics, gene flow refers to the exchange of genetic material between populations. In our two-niche models, gene flow can be viewed as the exchange of alleles between niche populations and therefore reflects the degree of RI between locally adapted ecotypes.

Gene flow ultimately depends on the production of viable, fertile hybrids capable of backcrossing with parental populations and facilitating introgression. Barriers to gene flow arise through premating or postmating RI. Premating RI prevents mating between niche ecotypes, whereas postmating RI reduces hybrid viability or fertility when premating barriers are incomplete [42].

Postmating RI may be extrinsic or intrinsic. Disruptive ecological selection represents an extrinsic form of RI mediated by the environment: hybrids expressing intermediate, maladaptive phenotypes are eliminated when no suitable ecological niche exists [47]. In contrast, intrinsic postmating RI commonly arises from postzygotic hybrid incompatibilities, in which mismatched alleles inherited from each parent reduce hybrid survival through genetic or developmental dysfunction [45, 46].

In a prior study [38], we simulated the invasion dynamics of a one-allele model of postzygotic hybrid incompatibility in a two-niche sympatric population under disruptive selection, based on the mathematical framework shown in Fig 3. The results showed that a mutation *f*(-) that decreases the viability of hybrid offspring produced by inter-niche matings cannot invade because it is underdominant, causing individuals carrying the mutation to experience a fitness disadvantage relative to non-carriers within the same niche.

The simulations in the present study also confirm that, in a two-allele model of postzygotic barriers involving hybrid-incompatibility alleles *P* and *Q*, a mutant allele *P* conferring a stronger incompatibility bias (lower *v*) cannot invade when introduced near the origin of the *Ap*/*Bp* phase portrait. However, when divergent assortment of *P* and *Q* alleles is already established between ecotypes, a fixed point may emerge in the lower-right quadrant of the *Ap*/*Bp* phase portrait when both incompatibility bias and disruptive ecological selection are sufficiently strong (Figs 46d and 46f).

Such initial divergence in incompatibility alleles across niche ecotypes could arise following secondary contact, when allopatrically diverged populations become fixed for different incompatibility alleles through the Bateson–Dobzhansky–Muller (BDM) mechanism [42-44]. Alternatively, this divergence may arise as an inevitable byproduct of ecological adaptation when emerging ecotypes specialize to different niches in a sympatric population under disruptive ecological selection [42].

The schematic diagram in Fig 60 illustrates the interaction between incompatibility alleles *P* and *Q* following initial divergence. As shown, niche *A* predominantly carries the *P* allele, whereas niche *B* predominantly carries the *Q* allele. If the *P* and *Q* alleles are fixed in their respective niches before secondary contact, gene flow during secondary contact will cause the *Q* allele from niche *B* to introgress into niche *A*, and conversely the *P* allele from niche *A* to introgress into niche *B*. If no barriers to gene flow exist, individuals from niches *A* and *B* effectively form a single panmictic population, where individuals encounter one another randomly and mate freely. In this scenario, the most prevalent allele in the total population will eliminate the less prevalent allele through their attritive interactions (Fig 46a).

**Fig 60.**
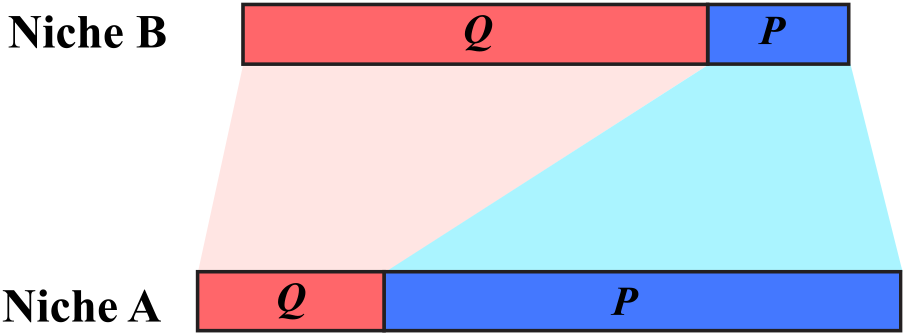
Schematic diagram illustrating interactions between incompatibility alleles (*P* and *Q*) following initial divergence between niche ecotypes. In the example shown, the *P* allele predominates in the niche-*A* ecotype, whereas the *Q* allele predominates in the niche-*B* ecotype. Because matings between genotypes carrying mismatched alleles produce less viable hybrid offspring, under panmixia the allele most numerous in the population as a whole (across niches *A* and *B*) eliminates the less common allele through their attritive interactions. By contrast, when gene flow between niche populations is restricted—owing to disruptive ecological selection or premating RI—most matings occur within niches rather than between niches. Consequently, increased inter-niche RI allows the locally prevalent allele (*P* in niche *A*; *Q* in niche *B*) to eliminate the less frequent, incompatible allele (*Q* in niche *A*; *P* in niche *B*) within each niche. This process enhances divergence of incompatibility alleles across niches and strengthens postzygotic RI between niche populations, reflected by a shift of the *Ap*/*Bp* fixed point toward the lower-right quadrant of the *Ap*/*Bp* phase portrait.

In contrast, when isolation is complete and gene flow is effectively zero—for example, when niches are separated by an insurmountable geographic barrier— the more prevalent incompatibility allele in each niche eliminates the less prevalent allele locally, producing the *Ap*/*Bp* phase portraits shown in Fig 46c. In this respect, hybrid incompatibility selection behaves analogously to sexual selection, in which numerically dominant alleles eliminate rarer incompatible alleles.

Between these two extremes, an intermediate but low level of gene flow can maintain the *P*–*Q* allele polymorphism shown in Fig 60. In this scenario, the minority *Q* allele is eliminated by the predominant *P* allele in niche *A* during intra-niche matings but is continuously replenished through net introgression from inter-niche matings. A selection–migration balance then maintains the *P*–*Q* polymorphism in niche *A*. The same mechanism operates in niche *B*, maintaining reciprocal polymorphism. This dynamic results in the emergence of a stable fixed point in the lower-right quadrant of the *Ap*/*Bp* phase portrait (Fig 46d).

These results have implications for the outcome of secondary contact. High gene flow—resulting from weak premating and/or postmating barriers—tends to eliminate postzygotic incompatibilities between diverged ecotypes fixed for different hybrid-incompatibility alleles, potentially leading to fusion into a single population. In contrast, low gene flow can maintain stable divergence in hybrid-incompatibility alleles, preserve postzygotic RI, and sustain distinct species boundaries.

Conventional theory holds that mutations causing intrinsic postzygotic incompatibilities are unlikely to be favored by selection because they reduce hybrid offspring viability and therefore incur a direct fitness cost [45]. Under this view, postzygotic barriers arise incidentally—most commonly through the accumulation of BDM incompatibilities during periods of low gene flow, such as in allopatric speciation—rather than through direct positive selection for reduced hybrid viability.

Nevertheless, a theoretical pathway may exist by which positive selection could favor the invasion of a mutation that reduces hybrid viability. Consider a scenario of secondary contact in which gene flow is already limited, owing to partial premating isolation or persistent geographic constraints. Under such conditions, the fitness cost of producing inviable hybrid offspring may be relatively small because inter-niche mating is infrequent. The primary direct cost would instead arise from incompatibilities expressed in intra-niche matings between carriers and non-carriers within the same ecotype, which are likely weaker than those between diverged ecotypes.

Now consider a scenario in which maladaptive introgression constitutes the primary selective pressure. If foreign lethal alleles entering from the opposite niche disrupt locally adapted genomic networks and substantially reduce the fitness of resident genotypes, then a mutation that limits such introgression—by decreasing the viability of inter-niche hybrid offspring—could confer a net selective advantage on its carriers. By restricting the introgression and spread of deleterious foreign alleles, the mutation would help preserve the integrity of locally adapted gene complexes.

For such a mutation to invade and spread, its indirect benefit in maintaining genomic integrity must outweigh its direct cost of reduced hybrid viability. If this condition is met, positive selection could, in principle, drive the invasion and eventual fixation of an underdominant incompatibility allele. Although this hypothesis is theoretically plausible, it remains to be validated by empirical evidence and formal modeling.

##### 1.2 Complementary action of hybrid-incompatibility selection and disruptive ecological selection promotes and reinforces divergence of hybrid-incompatibility alleles across niches, whereas roughly equal niche sizes are most conducive to such divergence

When incompatibility selection is complete (*v* = 0 and all mismatched hybrids are inviable; Fig 46e), the *Ap*/*Bp* phase portrait resembles that of complete isolation produced by complete ecological disruption (*f* = 0 and all ecological hybrids are inviable; Fig 46c). Gene flow between niches is absent, and the more prevalent incompatibility allele eliminates the less prevalent allele within each niche.

In the absence of complete incompatibility selection and disruptive ecological selection, weak disruptive ecological selection and weak hybrid-incompatibility selection (high values of *f* and *v*) may still be insufficient to produce fixed-point divergence in the *Ap*/*Bp* phase portrait (Fig 46b).

However, increasing either the strength of ecological selection (low *f*) or hybrid-incompatibility selection (low *v*) often leads to the emergence of a fixed point in the lower-right quadrant of the *Ap*/*Bp* phase portrait (Figs 46d and 46f). When a fixed point already exists, increasing the strength of either type of selection shifts the fixed point closer to the lower-right corner, resulting in stronger postzygotic RI between niche ecotypes.

The actions of hybrid-incompatibility selection and disruptive ecological selection appear to be complementary: weak hybrid-incompatibility selection can be compensated by strong disruptive ecological selection, and vice versa. This complementarity can be predicted from the mathematical model shown in Fig 3 by replacing the mating-bias variable *α* with the hybrid viability variable *v*, substituting alleles *X* and *Y* with *P* and *Q*, and setting the number of matching rounds (*n*) to one. In this model, both forms of selection, mediated by the parameters *f* and *v*, generate hybrid loss through interactions between genotype groups. Such interactions increase the frequency of the larger genotype group and decrease that of the smaller genotype group, thereby promoting divergence of hybrid-incompatibility alleles across niche ecotypes. Because hybrid-incompatibility selection eliminates mismatched hybrids from both intra-niche and inter-niche matings, it operates more like disruptive ecological selection (which eliminates maladaptive ecological hybrids) than like mating-bias selection (in which mating-bias hybrids produced within the same ecological niche remain viable).

The actions of these two forms of selection can also be explained by the schematic diagram in Fig 60. Increasing disruptive ecological selection (lower *f*) eliminates more hybrid offspring produced by inter-niche matings and results in a greater proportion of offspring being produced from intra-niche matings. This shift from inter-niche to intra-niche matings reduces gene flow and introgression of alleles from the opposite niche and increases elimination of the minority allele by the majority allele within each niche. Consequently, the *Ap*/*Bp* fixed point shifts toward the lower-right corner of the phase portrait, leading to stronger postzygotic RI between niche ecotypes.

Similarly, increasing the strength of hybrid-incompatibility selection (lower *v*) allows the majority allele to more effectively eliminate the minority allele, further shifting allele frequencies in favor of the majority allele within each niche.

Finally, our simulations demonstrate that fixed-point convergence is most likely when the two niches are equal in size, which maximizes inter-niche mating encounters and the production of mismatched hybrid offspring from inter-niche matings. Outside a symmetric range of *NA* values centered on *NA* = 0.5, fixed-point convergence becomes impossible (Fig 46g).

##### 1.3 Reciprocal reinforcement among premating barriers, postmating barriers, and ecological divergence strengthens reproductive isolation via positive feedback

Our analysis of interaction dynamics between niche ecotypes carrying divergent hybrid-incompatibility alleles (Fig 60) indicates that reducing gene flow between niche populations can drive further divergence of incompatibility alleles and increase postmating RI between niche populations. Therefore, premating barriers that limit inter-niche mating and the production of viable hybrid offspring can act to reinforce and strengthen existing postzygotic barriers.

In our models, these premating barriers can take the form of mating-bias barriers (Figs 47a–47c) [39] or habitat-preference barriers (Figs 48a and 48b) [48]. Their reciprocal interactions with postzygotic barriers generate a positive feedback loop in which premating and postmating barriers mutually reinforce one another, resulting in stronger overall RI.

The examples in Figs 47d and 47e further demonstrate that an established postzygotic incompatibility barrier can add to existing disruptive ecological selection, thereby intensifying selection against hybrid offspring from inter-niche matings. In our mathematical model (Fig 3), ecological selection and sexual selection act complementarily to promote fixed-point convergence and premating RI. Consequently, increased hybrid loss strengthens overall selection against hybrids (lowering the effective value of *f*) and relaxes the parametric conditions required for premating barriers to evolve via reinforcement.

Our simulation results therefore extend the classic concept of reinforcement. Traditionally, reinforcement refers to the evolution of premating barriers driven by selection to reduce postzygotic hybrid loss [56-58]. Our results suggest that reinforcement may also operate in a reciprocal or bidirectional manner, whereby premating RI can contribute to the strengthening of existing postzygotic RI between niche ecotypes. In this scenario, increases in postmating RI are ultimately driven by selection arising from inter-niche hybrid loss, either due to extrinsic disruptive ecological selection or intrinsic genetic incompatibility.

Previous studies have demonstrated the potential for enhanced mutual reinforcement and positive feedback between premating and postmating barriers when they are linked—either on autosomes or on the Y chromosome, where the absence of recombination ensures tight linkage—under a secondary contact scenario [53]. In contrast, our results indicate that such physical linkage is not necessary. Instead, effective mutual reinforcement and positive feedback between premating and postmating barriers can arise purely from population dynamic processes in a two-niche model involving ecotypes that carry divergent hybrid incompatibility alleles.

Such positive-feedback reinforcement can also occur among different postzygotic barriers (Figs 49a and 49b). Our simulations show that extrinsic disruptive ecological selection, by eliminating maladaptive hybrids, can reinforce intrinsic postzygotic hybrid-incompatibility selection. From the perspective of any specific postzygotic barrier, hybrid loss caused by extrinsic ecological selection or intrinsic genetic incompatibility is functionally equivalent and indistinguishable. Consequently, the establishment of one postzygotic barrier can facilitate the emergence and coupling of additional weaker barriers (Fig 49b). More generally, our results suggest that, after multiple postzygotic barriers accumulate in isolated populations during extended periods of allopatric divergence, selection against hybrids during secondary contact may cause these barriers to synergistically reinforce and strengthen one another, producing strong and potentially irreversible overall postmating RI.

Irreversibility is further reinforced because post-zygotic barriers are weakened or eroded by ongoing gene flow [39]. In the absence of gene flow, geographically separated populations are free to accumulate diverse postzygotic barriers—for example through mutation-order divergence, Dobzhansky– Muller incompatibility (DMI) accumulation, or the evolution of genetic conflicts [45-47, 59-62]—as they adapt to distinct environments. The accumulation of multiple postzygotic barriers can increase the resistance of reproductive isolation to erosion by gene flow. During secondary contact, individual post-zygotic barriers may be gradually weakened, in part due to their underdominance, but the presence of a large number of diverse, mutually reinforcing barriers can slow their collective elimination. This slower erosion of postzygotic isolation effectively extends the time window during which selection can favor the evolution or strengthening of premating barriers. As premating barriers become established, they reduce inter-niche mating and gene flow, helping to preserve existing postzygotic barriers and stabilizing reproductive isolation between diverging populations.

In contrast, young or incipient species pairs that possess only a small number of postzygotic barriers may be more vulnerable to fusion during secondary contact. With few barriers present, gene flow can rapidly erode postzygotic isolation before premating barriers have time to evolve or strengthen. As a result, interactions between postzygotic and premating barriers may play an important role in determining whether diverging populations complete speciation or collapse back into a single gene pool following secondary contact.

In our current model, each niche (*A* or *B*) is occupied by the most locally adaptive ecotype (ecotype *A* or ecotype *B*), which competitively excludes less adaptive variants within the same niche. However, fixation of the locally optimal ecotype may be slowed by soft selection arising from factors such as environmental heterogeneity, genetic drift, or pleiotropy. Under these conditions, ongoing gene flow can continuously introduce foreign alleles into each niche via introgression and maintain ecotype polymorphism within each niche. A selection– immigration balance maintains ecotype polymorphism within each niche, analogous to how hybrid-incompatibility allele polymorphisms are maintained in Fig 60.

Strong postzygotic RI shifts this balance by reducing gene flow and limiting the introgression of maladaptive alleles from the opposite niche, thereby allowing the most locally adaptive alleles to outcompete less-adaptive variants within each niche and increase in frequency through intra-niche ecological selection. In this way, postzygotic RI can facilitate divergent ecological adaptation between niche populations [63]. Reciprocally, reduced gene flow and enhanced ecological adaptation to distinct niche environments promote the accumulation of additional postzygotic hybrid incompatibilities between niches. As ecotype populations become more homogeneous within each niche while increasingly distinct across niches, inter-niche matings yield a greater proportion of genetically mismatched hybrids, strengthening intrinsic postzygotic isolation. Together, these processes may generate a positive feedback loop in which ecological divergence and postmating RI mutually reinforce one another.

Premating barriers can also generate the same positive feedback effect when ecotype polymorphism exists within distinct niches. By reducing inter-niche gene flow, premating barriers disrupt the selection– immigration balance that maintains polymorphism in favor of the most locally adaptive ecotype in each niche. This causes ecotype populations within each niche to become more homogeneous while becoming increasingly distinct across niches. The resulting increase in ecological divergence reinforces postzygotic RI by increasing the production of incompatible inter-niche hybrid offspring, which in turn further reinforces premating RI, completing the positive feedback loop.

Finally, theoretical models and simulations predict that during the intermediate and late stages of speciation, strong assortative mating can further strengthen postmating RI through a mechanism analogous to ancestry bundling [64]. In our model, when ecotypes are polygenic, individuals sharing the same ecotype also share similar sets of niche-adaptive alleles; thus, assortative mating by ecotype effectively bundles these alleles in their offspring. Because individuals preferentially mate with partners of the same ecotype, hybrids carrying mixed sets of ecotype-specific alleles become concentrated in a subset of individuals, increasing the variance among individuals in ecotype composition. This increased variance allows disruptive ecological selection to purge maladaptive hybrid genotypes more efficiently, thereby strengthening genetic isolation between populations.

#### 2 Systems with Multilocus Ecological Genotypes and Single-Locus-Equivalent Hybrid-Incompatibility Genotypes

##### 2.1 Increasing the number of ecological gene loci or reducing viable ecological hybrids strengthens disruptive ecological selection and reinforces postzygotic RI between niche ecotypes

Consistent with our analyses of the interaction dynamics between two niche populations carrying divergent hybrid-incompatibility alleles (Fig 60), our simulations show that strengthening disruptive ecological selection in models with multilocus ecological genotypes—either by increasing the number of ecological gene loci or by reducing the proportion of viable ecological hybrids—leads to stronger postmating RI (Figs 50a–50c).

This pattern arises because reduced inter-niche gene flow decreases the introgression of the locally rare hybrid-incompatibility allele from the opposite niche. As a result, the locally prevalent allele in each niche (e.g., the *P* allele in niche *A*) can more effectively remove the less prevalent allele (the *Q* allele in niche *A*), thereby increasing the divergence of incompatibility alleles across niches.

##### 2.2 Invasion of a CI capturing only locally adaptive ecological alleles weakens both disruptive ecological selection and postmating RI, whereas invasion of a CI capturing locally adaptive ecological alleles together with the locally prevalent hybrid-incompatibility alleles enhances postmating RI

Invasion of a CI capturing only locally adaptive ecological alleles reduces the effective number of ecological gene loci and weakens disruptive ecological selection, leading to increased inter-niche gene flow and weaker postmating RI (Fig 51).

In contrast, when the CI captures both locally adaptive ecological alleles and the locally prevalent hybrid-incompatibility allele, it anchors the prevalent incompatibility allele within its adaptive niche and limits its introgression into the opposite niche. This results in increased divergence of hybrid-incompatibility alleles across niches and stronger postmating RI (Figs 52a and 52b).

In both scenarios, despite their different outcomes, the more locally adaptive ecological alleles a CI captures, the greater its invasion fitness. This occurs because individuals carrying the CI possess a favorable combination of locally adaptive ecological alleles that cannot be broken up by recombination, giving them a fitness advantage over same-niche individuals lacking the CI.

Our simulations also show that CIs capturing alleles that are not locally adaptive (e.g., niche-*B* ecological alleles linked with alleles from the niche-*A* ecotype) experience a fitness disadvantage, cannot invade, and ultimately go extinct.

#### 3 Systems with Multilocus Hybrid Incompatibility Genotypes

##### 3.1 Increasing the number of hybrid-incompatibility loci, reducing hybrid viability, or strengthening disruptive ecological selection promotes the emergence of postzygotic barriers and reinforces existing postmating RI

We developed a GUI application (MultiCompCI) that allows users to model any specified number of hybrid-incompatibility loci within the mathematical framework illustrated in Fig 3. In this application, hybrid offspring carrying mismatched incompatibility alleles are directly eliminated by applying a V-shaped viability function to the offspring genotype distribution in each generation. Genotypes with the greatest number of mismatches—located at the center of the distribution—are reduced by the viability parameter *v*, whereas parental genotypes without mismatches, located at the extremes of the distribution, remain unaffected (see Fig A5 in the Appendix). Genotypes with intermediate levels of mismatch are reduced proportionally according to a linear function that scales from *v* at the center to 1 at the extremes (see Methodology for details).

In general, the simulation results of the multilocus model align closely with those of the one-locus-equivalent hybrid-incompatibility model. In both frameworks, disruptive ecological selection and hybrid-incompatibility selection act in a complementary manner to promote the emergence and strengthening of postzygotic RI. Weak disruptive ecological selection (high value of *f*) can be compensated by strong hybrid-incompatibility selection (low value of *v*), and vice versa (Figs 54a– 54b and 55).

In multilocus models, hybrid-incompatibility selection operates analogously to sexual selection within niches, in that the locally prevalent allele tends to eliminate the less prevalent incompatible allele. However, unlike mating-bias hybrids—which remain ecologically viable within the same niche— incompatibility hybrids are always eliminated due to genetic mismatch. As a result, hybrid-incompatibility selection behaves more similarly to disruptive ecological selection than to mating-bias selection within the mathematical framework illustrated in Fig 3.

Accordingly, increasing the number of hybrid-incompatibility gene loci strengthens hybrid-incompatibility selection. As the number of loci increases, more mismatched hybrid genotypes are produced and subsequently eliminated, thereby reducing the effective value of *v*. This effect parallels disruptive ecological selection, in which increasing the number of ecological loci strengthens selection. In contrast, in mating-bias selection, fewer loci produce stronger selection because fewer viable mating-bias hybrids are generated. Consistent with these expectations, our simulations show that increasing the number of hybrid-incompatibility loci promotes fixed-point convergence (Figs 53a–53c) and strengthens an existing *AP*/*BP* fixed point by shifting it toward the lower-right corner of the phase portrait (Fig 56).

##### 3.2 CIs capturing multiple locally prevalent hybrid-incompatibility alleles gain an invasion advantage but reduce postmating RI

In our model, hybrid incompatibility selection operates analogously to disruptive ecological selection, in which hybrid offspring have reduced viability. Increasing the number of incompatibility loci therefore strengthens both forms of selection and promotes convergence toward stable fixed points. This contrasts with mating-bias selection, where fewer loci facilitate convergence because mating-bias hybrids remain viable within the same niche.

Based on this reasoning, we expect the invasion of CIs capturing multiple locally prevalent hybrid-incompatibility alleles to reduce the effective number of incompatibility loci, hinder system convergence, and weaken existing postmating RI.

Our simulation results support this prediction. Locally advantageous alleles captured within a CI cannot be separated by recombination, giving individuals carrying the CI a fitness advantage over same-niche individuals lacking the inversion. Whereas a CI capturing a single incompatibility allele confers no such advantage and fails to invade (Fig 57a), a CI capturing multiple locally prevalent incompatibility alleles gains positive invasion fitness. However, by reducing the effective number of loci, the successful invasion of such a CI either weakens existing prezygotic RI (Figs 57b–57d) or causes system divergence (Figs 57e–57f).

Because new mutations causing hybrid incompatibility are underdominant, they cannot invade from the origin of the phase portrait. Such mutations can persist only in the lower-right quadrant of the *AP*/*BP* phase portrait, where divergence in hybrid-incompatibility alleles is already established. Consequently, a CI capturing hybrid-incompatibility alleles can invade only from an existing fixed-point population in this region and cannot invade when no fixed point exists in the phase portrait (Fig 58). If a CI capturing locally prevalent hybrid-incompatibility alleles is already present at the onset of secondary contact, its invasion can likewise reduce the effective number of incompatibility loci and impede convergence (Fig 59).

Nevertheless, as demonstrated in our previous study, an underdominant incompatibility mutation may successfully invade if it becomes associated with other processes that confer sufficient fitness benefits to offset its selective disadvantage [38]. For example, tight linkage to advantageous alleles through capture by CIs, location within chromosomal regions of reduced recombination, pleiotropic effects, or its occurrence as an inevitable byproduct of local adaptation can compensate for the costs of underdominance. By hitchhiking with these processes, the mutation may attain a net fitness advantage and invade from the origin of the phase portrait.

### INTERACTIONS AMONG PREMATING AND POSTMATING ISOLATION, ECOLOGICAL DIVERGENCE, AND CHROMOSOMAL INVERSIONS

Fig 61 summarizes the three speciation processes investigated in this study: premating RI, postmating RI, and divergent ecological adaptation. Premating RI may arise from mating-bias barriers or habitat-preference barriers. Postmating RI may be either extrinsic or intrinsic. Extrinsic postmating RI results from disruptive ecological selection, in which environmental conditions eliminate maladaptive ecological hybrids. Intrinsic postmating RI arises from reduced hybrid viability due to genetic incompatibilities. Ecological divergence occurs when ecotypes adapt to distinct environmental conditions.

**Fig 61.**
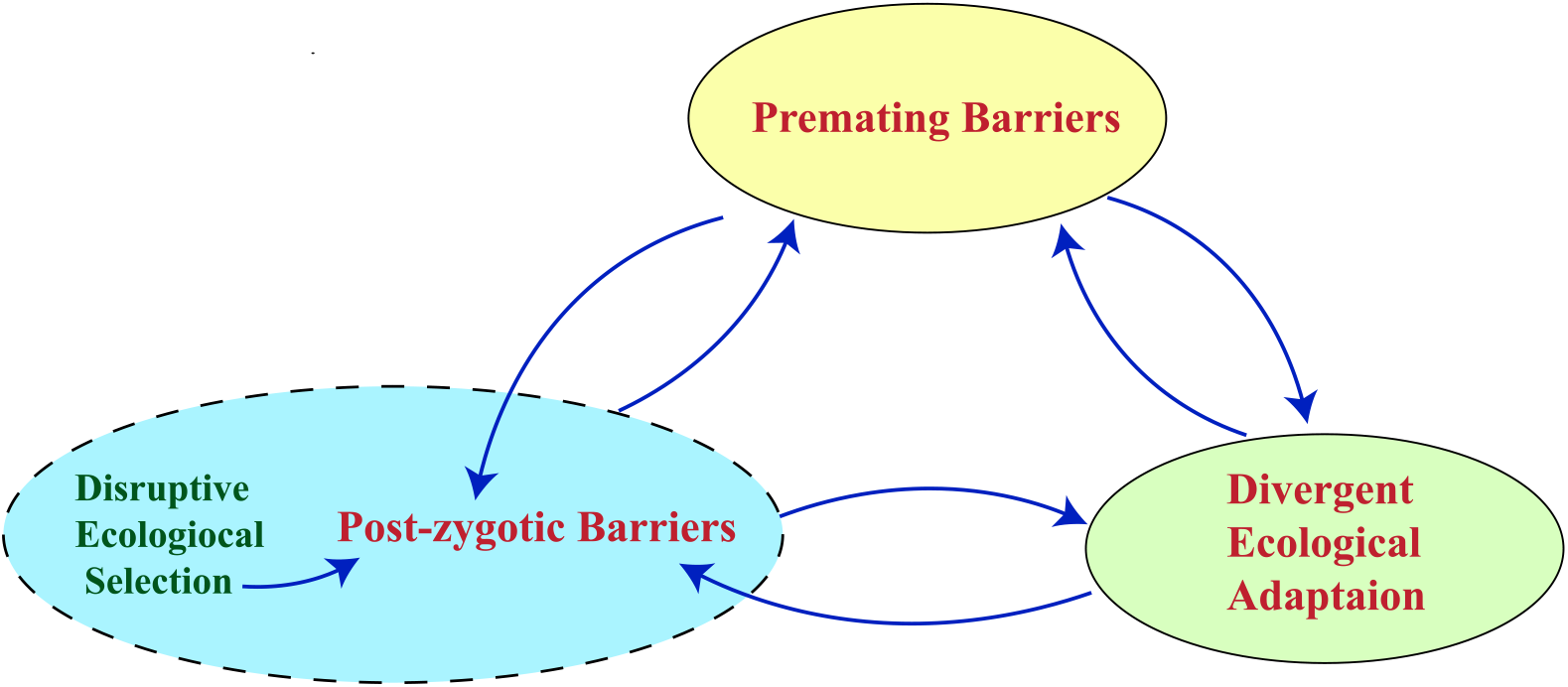
Schematic diagram illustrating the interactions among premating barriers, postmating barriers, and ecological divergence. Premating RI may result from mating-bias barriers or habitat-preference barriers. Postmating RI may be either extrinsic, driven by disruptive ecological selection, or intrinsic, caused by postzygotic hybrid incompatibilities. Reciprocal arrows indicate mutual reinforcement among these processes, forming a positive feedback loop that strengthens the interacting mechanisms.

The reciprocal arrows indicate mutual reinforcement between processes within a positive feedback loop. In our simulations, disruptive ecological selection strengthens postzygotic barriers (Fig 46d), and distinct types of postzygotic barriers can reinforce one another (Figs 49a–49b). Reciprocal reinforcement also occurs between premating and postzygotic barriers, generating a positive feedback loop that enhances both forms of barriers (Figs 47a–47e and 48a–48b).

Although not explicitly modeled or simulated in the present study, we propose that analogous reciprocal reinforcement and positive feedback are likely to occur between premating barriers and ecological divergence, as well as between postzygotic barriers and ecological divergence, when ecotype polymorphisms are maintained within each niche. This proposition is supported by the following considerations:

1. Based on the interaction dynamics of our mathematical model (Fig 3), disruptive ecological selection (postmating RI) and premating mating-bias RI act complementarily to generate a stable fixed-point polymorphism of mating-bias alleles in the phase portrait and promote ecotype divergence [39].
2. Simulation results from previous work have shown that differences in fitness of ecotypes across distinct niches cause divergent adaptations and can reinforce premating RI [38].
3. Theory predicts reciprocal reinforcement and positive feedback between postzygotic barriers and divergent ecological adaptation when polymorphisms of ecotypes exist in distinct niches (see Discussion, subsection 1.3, “Systems with Multilocus Hybrid-incompatibility Genotypes”). By similar reasoning, premating RI can also restrict gene flow and reduce the variance of ecotype polymorphisms in distinct niches.
4. Theoretical models predict that differential adaptation of ecotypes inherently generates postzygotic barriers as an inevitable consequence of gene network divergence (see the gene network theory of speciation in [65]).
5. In intermediate or late stages of speciation, when premating RI due to assortative mating is already established, increased ecological divergence can reinforce postzygotic barriers through ancestral bundling [64].

Our GUI applications examine the effects of multilocus polygenic traits in each of the processes shown in Fig 61. In a sympatric population under disruptive ecological selection, increasing the number of gene loci underlying ecological traits promotes fixed-point convergence and the evolution of RI. Because more maladaptive ecological hybrids are generated and eliminated, ecological selection is strengthened (lower *f*) [49].

Similarly, in postzygotic hybrid-incompatibility selection, increasing the number of incompatibility loci produces more genetically mismatched hybrids that are removed by intrinsic incompatibilities, promoting fixed-point convergence and strengthening postzygotic selection (lower *v*).

In contrast, in premating mating-bias selection, all mating-bias hybrids remain viable within the same ecological niche. In this case, fewer mating-bias loci facilitate fixed-point convergence, because fewer viable mating-bias hybrids are produced, thereby strengthening premating mating-bias selection (lower *α*) [49, 55].

Understanding the effects of multiple gene loci in each of the processes shown in Fig 61 helps clarify the consequences of invasion by a CI capturing locally adaptive alleles. In general, a CI that captures multiple gene loci reduces the effective number of loci in the system, and its impact depends on whether more loci strengthen or weaken the process in question.

Consequently, invasion by a CI capturing multiple locally adaptive ecological alleles reduces the effective number of ecological loci and weakens ecological selection, leading to system divergence or weaker RI (Figs 24a and 24b). Notably, invasion by a CI capturing all locally adaptive ecological alleles eliminates the production of ecological hybrids, and the system invariably becomes divergent because there is no hybrid loss to drive the evolution of premating RI (Figs 23a–23c). Similarly, invasion by a CI capturing locally prevalent hybrid-incompatibility alleles reduces the effective number of incompatibility loci and weakens existing postzygotic RI (Figs 57b– 57d). Invasion by a CI capturing all locally prevalent hybrid-incompatibility alleles eliminates the production of incompatibility hybrids and invariably causes the system to become divergent (Figs 57e and 57f). In contrast, invasion by a CI capturing locally advantageous high-mating-bias alleles reduces the effective number of mating-bias loci, which promotes system convergence and strengthens existing premating RI (Figs 39–45).

Lastly, our GUI applications allow us to examine the invasion of CIs capturing locally adaptive alleles from different processes. In general, invasion by CIs capturing locally adaptive alleles from different processes can couple those processes and anchor the captured advantageous alleles in their adaptive niches, resulting in enhanced nonrandom assortment of alleles across niches and synergistically strengthening overall RI [66, 67]. An exception occurs when the invading CIs capture all locally adaptive ecological or hybrid-incompatibility alleles, thereby eliminating the production of unfit hybrids and depriving the system of the hybrid loss necessary to drive the evolution of RI. In this case, invasion by such CIs, instead of facilitating system convergence and strengthening RI, can cause system divergence and weaken existing RI Figs 26 and 28).

Our simulation results show that invasion by a CI capturing a partial set of locally adaptive ecological alleles (e.g., in niche *A*) together with the locally prevalent mating-bias allele (e.g., the *X* allele) anchors the mating-bias allele within its adaptive niche and shifts the *Ax*/*Bx* fixed point toward the lower-right corner of the phase portrait, resulting in stronger RI (Figs 30a–30b, 31a–31d, and 32).

Although capturing more adaptive ecological alleles increases the invasion fitness of the CI, the resulting premating RI tends to be weaker. This occurs because capturing additional locally adaptive ecological alleles reduces the production of maladaptive hybrids and, consequently, the hybrid loss necessary to drive the evolution of strong premating RI (Figs 31a– 31c).

Our results further show that the strongest RI is produced when the CI captures only a single locally adaptive ecological allele together with the locally prevalent mating-bias allele (Figs 31a–31c). This combination maximizes hybrid loss to drive the evolution of strong premating RI while simultaneously reducing the homogenizing effect of gene flow by anchoring the mating-bias allele in its adaptive niche. Because increasing the number of ecological loci strengthens ecological selection, whereas decreasing the number of mating-bias loci strengthens mating-bias selection, we expect that an invading CI capturing the fewest ecological loci and the greatest number of mating-bias loci would be most effective at promoting system convergence and generating strong RI, although this prediction has not been directly verified by simulations in the present study.

Similarly, our simulations show that invasion by a CI capturing a partial set of ecological alleles together with the locally prevalent hybrid-incompatibility allele anchors the incompatibility allele within its adaptive niche by preventing its introgression into the opposite niche, thereby strengthening postzygotic RI between niches (Figs 52a and 52b).

Because increasing the number of gene loci strengthens hybrid-incompatibility selection, we expect that an invading CI capturing the fewest ecological loci and the fewest hybrid-incompatibility loci would be most effective at promoting system convergence and increasing RI. This combination maximizes ecological hybrid loss, thereby strengthening ecological selection, and maximizes incompatibility hybrid loss, thereby enhancing hybrid-incompatibility selection. However, this prediction has not been directly verified.

Our current study does not simulate the invasion of CIs capturing both hybrid-incompatibility alleles and mating-bias alleles. However, simulation results from a separate study demonstrate that a mutant allele combining mating-bias and hybrid-incompatibility properties can achieve positive invasion fitness by offsetting the cost of incompatibility with the fitness advantage of high mating bias [38]. Invasion by such a CI can generate synergistically stronger RI through positive reinforcement. Another modeling study also shows that when premating and postmating barriers are physically linked—either on an autosome or on a Y chromosome—reciprocal reinforcement and positive feedback can produce strong RI [53].

Based on our reasoning regarding how the number of gene loci influences barrier strength, we expect that invasion by a CI capturing the fewest hybrid-incompatibility alleles and the greatest number of mating-bias alleles would be most effective at promoting system convergence and strengthening RI. This prediction is consistent with a prior modeling study of diploid, gonochoric organisms, which found that partial linkage of barriers can be more effective than complete linkage in generating strong RI [35, 36].

Lastly, all three processes shown in Fig 61 can be captured within a single CI to generate strong overall RI. This scenario was examined in a separate study, in which the properties of premating RI, postmating RI, and divergent ecological adaptation were incorporated into a single mutant allele to investigate the evolution of BDM incompatibilities [38].

### STAGE-DEPENDENT EFFECTS OF POLYGENIC ARCHITECTURE ON SYMPATRIC DIVERGENCE

Our model assumes that, within each niche, the strength of local selection exceeds the homogenizing effect of migration, thereby ensuring the establishment and persistence of two distinct ecotype populations in a sympatric system under disruptive ecological selection. Under this assumption, the system is effectively constrained to a bimodal ecological landscape in which intermediate genotypes lack access to niche resources. Consequently, the formation of ecotypes is not a limiting step, and the dynamics we analyze primarily reflect the subsequent processes governing the maintenance and reinforcement of divergence.

Mechanistically, this is implemented by assuming that sufficient numbers of offspring carrying locally adaptive genotypes are produced each generation, allowing these ecotypes to outcompete less adaptive hybrid genotypes and increase in number to fully saturate each niche’s carrying capacity. As a result, the ecotype populations—bounded by the niche-specific carrying capacities *NA* and *NB*—are effectively fixed at the beginning of each mating generation. This assumption simplifies the analytical framework and is biologically plausible in systems with high reproductive output, such as many fish species that can spawn millions of offspring with diverse genotypes within a single breeding season.

However, there are situations in which local selection is insufficient to overcome the homogenizing effect of migration and gene flow, preventing the formation of distinct ecotype populations. In such cases, our prior individual-based simulations [39] indicate that a minority niche population may be eliminated due to incumbent selection (i.e., demographic swamping). Although the minority ecotype may experience increased ecological and reproductive fitness due to access to underutilized resources, this advantage may not be enough to compensate for the loss of maladaptive hybrid offspring resulting from inter-niche mating. Consequently, the minority population may fail to establish and persist.

Under certain conditions, however, the minority ecotype may persist at a reduced but stable population size. This occurs when the ecological advantage conferred by access to an underutilized niche allows the minority population to produce more offspring and partially offset hybrid losses, resulting in a stable equilibrium with the majority ecotype population occupying the alternative niche. In such cases, the effective population size of the minority ecotype— rather than the full niche carrying capacity—provides a more appropriate parameter for model calculations [39]. Once premating isolation evolves between niche ecotypes, the minority population, freed from incumbent selection, may then increase in number to reach its niche’s carrying capacity.

In natural systems, environmental heterogeneity and stochastic processes may further complicate these dynamics. Even under disruptive ecological selection, when selection is weak and migration is strong, sympatric populations may remain centered around a single intermediate phenotype rather than splitting into two specialized ecotypes. This intermediate phenotype can be interpreted as a suboptimal generalist capable of utilizing resources in both niches, although less efficiently than niche-specific specialists.

Numerous modeling studies have investigated the conditions under which populations may undergo ecological branching during the early stages of sympatric speciation. In contrast to our results, a recent simulation study [5] shows that the emergence of local adaptation under gene flow is facilitated when ecological traits are controlled by a small number of loci with relatively large effects. Nonetheless, this result is indirectly supported by our finding that chromosomal inversions capturing locally adaptive alleles can acquire a fitness advantage and successfully invade.

This prior study further shows that, under high gene flow and highly polygenic architectures composed of many small-effect loci, most loci are individually susceptible to swamping (swamping-prone) [5]. In such cases, chromosomal inversions can facilitate local adaptation by capturing and linking multiple locally adaptive alleles into a single genomic block, effectively forming a supergene of larger effect. This reduces the effective number of independently segregating ecological loci and increases the overall strength of selection acting on the combined genotype, thereby promoting local adaptation and ecotype divergence.

Once two distinct ecotype populations have been established—either through sufficiently strong disruptive selection or local selection, or through the action of other barrier mechanisms—the role of polygenicity changes fundamentally. In this later stage, our results show that increasing the number of ecological loci enhances multilocus mismatch in hybrid and recombinant genotypes. This mismatch amplifies hybrid maladaptation and reduces hybrid offspring fitness, thereby strengthening disruptive ecological selection and promoting the maintenance of local adaptation. In this regime, polygenicity facilitates divergence rather than hindering it.

Consequently, chromosomal inversions that capture locally adaptive alleles can have stage-dependent effects. In the early stage, by reducing the effective number of loci, inversions facilitate the emergence of ecotype divergence. In the later stage, however, this reduction in locus number may limit the accumulation of multilocus ecological mismatch and thereby weaken the reinforcement of reproductive isolation.

This stage dependence raises the question of what constitutes an “optimal” level of polygenicity for sympatric speciation. Our results suggest that a single static level of polygenicity may not be optimal across all stages of divergence. Instead, different genetic architectures may be favored at different stages. In particular, architectures dominated by a few loci of larger effect may facilitate the initial establishment of ecotypic divergence, whereas architectures involving many loci of small effect may contribute to multilocus mismatch and the strengthening of reproductive isolation once divergence has begun [49]. Mechanisms such as chromosomal inversions or other forms of linkage could facilitate this process by preserving advantageous combinations of alleles and promoting the coupling of barrier mechanisms, although the extent to which such effects are necessary or widespread in natural systems remains uncertain.

Lastly, it is important to note that, in our sympatric model, migration between ecotypes is effectively maximal under panmixia. If migration between niches is allowed to vary—such as in a two-island model where gene flow is governed by a migration parameter—then reducing the migration rate can shift the migration–selection balance in favor of local selection. Under such conditions, sufficiently low migration can facilitate the establishment and maintenance of two stable ecotype populations.

In summary, the role of genetic architecture in sympatric speciation is inherently dynamic. Fewer ecological loci favor the initiation of local adaptation and ecological divergence, whereas a greater number of ecological loci strengthens the maintenance and reinforcement of divergence once distinct ecotype populations are established. This stage-dependent shift highlights a fundamental tension in determining the optimal level of polygenicity and underscores the complex role that chromosomal inversions play in facilitating sympatric speciation.

## V. LIMITATIONS

Modeling the mechanisms of speciation reminds us of the parable of a group of blind men trying to describe an elephant: each man perceives only a part of the elephant and may arrive at a partial, sometimes seemingly conflicting, description, yet each captures a real aspect of the underlying object. The same limitation applies to the modeling of biological processes. All models necessarily rely on simplified assumptions and constrained methodologies, and thus can represent only part of the complex dynamics involved. We nevertheless assume that an objective truth exists in nature. Although no single model can fully describe it, they are ultimately all describing the same underlying reality. Thus, different models could be viewed as complementary rather than contradictory, and integrating insights across approaches can help reveal a more complete understanding of the process.

The same limitations apply to our models. For every parametric variable included in our models, many others are excluded for the sake of simplicity and analytical tractability. These choices reflect practical constraints rather than judgments about biological importance, and in no way imply that the omitted variables are unimportant or would not produce surprising or contradictory results.

With respect to mating strategy, our models use phenotype matching among hermaphroditic haploid individuals instead of the more traditional approach of modeling compatibility between gonochoric individuals with male traits and female preferences [68]. Theoretically, we expect the results to be the same because natural systems tend to maintain equal sex ratios through various feedback mechanisms. Because individuals of the same sex cannot mate and are effectively invisible to one another in the mating pool, the computations should ultimately yield equivalent results. This expectation is supported by previous modeling studies [49, 69], although we have not explicitly implemented or tested it in our study.

Similarly, extending the model to diploid individuals would require explicit consideration of dominance hierarchies and recessivity in phenotype expression, leading to additional layers of complexity and nonlinear interactions among genotypes. While such extensions may yield important insights, they fall outside the scope of the present study.

In our modeling of CIs, we focus primarily on their ability to suppress recombination among captured alleles and therefore largely ignore rare events such as single crossovers within inverted regions [22, 40] and gene flux arising from double crossovers or gene conversion [41, 70], all of which can erode this suppression. In reality, although the effects of single crossovers are typically small, they can nevertheless contribute to hybrid loss in heterokaryotype matings and may play an important role in preventing invasion by noninverted genotypes once a CI has become predominant within a niche [38]. Similarly, gene flux can undermine CI persistence by restoring recombination among the captured alleles.

To simplify the analysis, we model polygenic extensions of ecological traits and mating-bias traits separately rather than simultaneously. To examine polygenic ecological traits, we extend the model in Fig 3 by allowing multilocus ecological genotypes and hybrid niche resources, while retaining the original one-locus, bi-allelic framework for mating bias. Conversely, to study polygenic mating-bias traits, we retain the ecological framework of the model in Fig 3, with the offspring return ratio *f* specifying the strength of disruptive ecological selection, and allow mating bias to be determined by multiple loci. This modular approach limits the number of interacting variables and maintains analytical tractability, while still capturing the essential dynamics expected in a model that includes both multilocus ecological and multilocus mating-bias genotypes. Nonetheless, we cannot exclude the possibility that jointly modeling both extensions may yield additional or unexpected results. The same approach is used to model multilocus hybrid-incompatibility genotypes, and the same limitations apply.

Additional limitations arise from the design choices in the models. For example, in our treatment of polygenic ecological traits, each gene locus is restricted to two alleles that are randomly assorted from parents to offspring with equal probability. Epistatic interactions are not modeled explicitly, but can be approximated by modifying the distribution of niche resources, which effectively serves as a simplified genotype–phenotype fitness landscape. We also assume that, in each generation, sufficient offspring are produced to fully occupy all niche carrying capacities, thereby fixing initial mating population ratios and avoiding incumbent selection [39]. In natural systems, however, these assumptions are likely oversimplifications and may fail to capture the full complexity, nonlinearity, and multivariate nature of ecological and evolutionary interactions.

Similarly, in modeling multilocus mating-bias traits, we assume that mating biases vary linearly with genetic distance between genotypes, as illustrated by the Matching Compatibility Table in Fig 15. This structure promotes smooth hill-climbing dynamics toward a global fitness maximum for an invading CI capturing high-mating-bias alleles. In nature, however, mating compatibilities are unlikely to follow such a strictly monotonic pattern. Instead, mating compatibilities are likely shaped by nonlinear pleiotropic and epistatic interactions among alleles, creating rugged genotype–phenotype landscapes in which high-fitness CIs may become trapped at local fitness peaks. In modeling multilocus hybrid-incompatibility genotypes, we likewise assume that viability varies linearly with the degree of allele mismatch according to a V-shaped fitness function—an assumption that overlooks nonlinearities in genotype–phenotype interactions.

Finally, our models do not explicitly simulate direct competition among CIs with different fitness values arising from the specific alleles they capture. Instead, we evaluate the invasion fitness of hypothetical CIs and implicitly assume that evolutionary processes—such as de novo mutation, adaptive introgression, or standing variation [71]—will generate these variants and allow fitter CIs to replace less-fit ones over time. While this approach isolates the fundamental invasion dynamics of CIs, it does not capture the full competitive and historical processes through which multiple CIs may arise and interact in natural populations.

## VI. CONCLUSION

Modern genomic studies have revealed that chromosomal inversions (CIs) are widespread across the genomes of many organisms and are increasingly recognized as important contributors to adaptation and speciation. Their significance is commonly attributed to their ability to suppress recombination among favorable combinations of adaptive alleles captured within inverted regions, thereby packaging these alleles into a supergene that can be transmitted and inherited as a single Mendelian locus. This property makes CIs particularly effective in polygenic, high– gene-flow environments, where adaptive traits are determined by many loci of small effect that would otherwise be vulnerable to recombination. By consolidating these loci into a single linkage block, inversions can generate large phenotypic effects and maintain local adaptation despite ongoing gene flow. These features suggest an important role for CIs in sympatric speciation, where gene flow remains substantial and adaptive traits are often polygenic. However, the precise mechanisms by which CIs promote or constrain speciation—and how they interact with other barrier mechanisms—remain incompletely understood.

In a previous study, we developed a two-niche, two-mating-bias-allele model of a sympatric population under disruptive ecological selection to examine the mechanism of sympatric speciation [39]. In the present study, we extend this framework to a polygenic context by modeling multilocus ecological, mating-bias, and hybrid-incompatibility genotypes and allowing for viable hybrids. Using simulations, we show how polygenic trait architectures can reshape system dynamics and how the invasion of adaptive CIs can further modify these dynamics by altering linkage relationships, reproductive isolation (RI), and longterm evolutionary outcomes.

In sympatric speciation, CIs are considered a late-stage, adaptive mechanism. They are late-stage because CIs are more likely to capture locally adaptive alleles once divergence and nonrandom assortment have occurred across niches [31, 38]. CIs are adaptive because their invasion fitness is derived from selection to reduce unfit hybrid loss [38]. CI invasion is maximized under intermediate gene flow. In early-stage speciation, high gene flow prevents locally adaptive alleles from becoming sufficiently differentiated for effective capture; when speciation is nearly complete, RI prevents sufficient hybrid loss to drive CI invasion [38].

Overall, our polygenic model confirms that viable hybrids—whether ecological, mating-bias, or hybrid-incompatibility hybrids—impede system convergence and the evolution of RI. Consequently, RI is more readily established with more ecological loci and hybrid-incompatibility loci and fewer mating-bias loci.

By suppressing recombination among linked alleles, the invasion of adaptive CIs influences convergence and the establishment of RI by reducing the effective number of ecological, mating-bias, or hybrid-incompatibility loci contributing to hybrid formation. In general, inversions capturing a greater number of locally adaptive alleles have higher invasion fitness. However, the consequences of invasion are determined by how the number of trait loci modifies the strength of the underlying barrier mechanisms. Accordingly, an inversion capturing locally adaptive alleles can either promote or impede the evolution of RI, depending on whether this reduction in the effective number of loci increases or decreases the strength of selection generated by the underlying isolating mechanisms.

Although prior studies have shown that local adaptation under gene flow may be facilitated by architectures with fewer ecological loci of larger effect during early divergence [5], this apparent contrast does not contradict our findings but instead reflects stage-dependent roles of polygenic architecture. By reducing the effective number of loci, inversions may facilitate early-stage ecotype divergence, but this same reduction may limit multilocus mismatch and weaken the reinforcement of reproductive isolation once distinct ecotype populations are established..

A key mechanism by which CIs can promote speciation is through the capture of adaptive alleles from distinct isolating mechanisms, thereby synergistically increasing their selection strength and anchoring them within their adaptive niches (Fig 61). In contrast to inversions that capture only alleles associated with a single barrier process, inversions capturing an optimal number of locally adaptive alleles across multiple barrier processes consistently promote the evolution of strong RI between niche ecotypes.

Our simulation results show that inversions capturing locally prevalent mating-bias or hybrid-incompatibility alleles together with a subset of locally adaptive ecological alleles can anchor these barrier alleles within their adaptive niche and limit their introgression into alternative niches. By physically linking barrier alleles to ecological alleles, the invasion of such inversions reduces the homogenizing effects of gene flow and promotes the divergence of barrier alleles across niches. This coupling and anchoring of distinct barrier mechanisms promote system convergence and facilitates the evolution of stronger RI. Although not directly simulated here, previous studies indicate that CIs capturing adaptive mating-bias and hybrid-incompatibility alleles similarly exhibit elevated invasion fitness and contribute to the evolution of strong RI [38, 53].

A key caveat is that, for ecological and hybrid-incompatibility barriers, the invasion of a CI that captures all locally adaptive alleles underlying either mechanism eliminates the production of unfit hybrids and the associated hybrid loss necessary to drive the evolution of RI. This constraint does not apply to mating-bias barriers, as all mating-bias hybrids are viable within the same ecological niche. This is consistent with our simulation results showing that more loci increase the strength of disruptive ecological and hybrid-incompatibility selection, but weaken mating-bias selection. Therefore, capturing a partial subset—rather than the entirety—of adaptive ecological or hybrid-incompatibility alleles is necessary to preserve selection against unfit hybrids and allow the evolution of RI.

Taken together, our simulations and prior theoretical work suggest that, in the coupling of different barrier mechanisms, the invasion of a CI capturing the least number of ecological or hybrid-incompatibility alleles but the greatest number of mating-bias alleles produces the strongest RI. This configuration maximizes the selection strength of the coupled barrier mechanisms, even though it may reduce the invasion fitness of the CI and prolong its establishment time.

Our analyses also reveal reciprocal reinforcement and positive feedback among the three principal speciation processes examined in this study: premating isolation, postmating isolation, and divergent ecological adaptation (Fig 61). Modeling each process as polygenic enabled us to evaluate how the number of loci underlying these traits modulates the strength of selection generated by these RI mechanisms. We also assessed how inversions capturing alleles within and across these processes influence invasion dynamics and barrier evolution in a two-niche model of sympatric speciation.

Overall, our findings reinforce the view that CIs play an important role in speciation. An inversion capturing adaptive alleles within a single barrier mechanism may either facilitate or impede the evolution of RI, depending on the mechanism involved. In contrast, inversions that couple alleles across distinct barrier mechanisms consistently gain an invasion advantage and can synergistically enhance barrier strength, thereby accelerating the evolution of robust RI and promoting speciation.

It is worth noting that chromosomal inversion represents but one mechanism within a broader class of processes that suppress recombination and preserve favorable allele combinations. Similar effects may arise through pleiotropy [72], genomic clustering [5], alternative chromosomal rearrangements [27], physical proximity, or localization within regions of reduced recombination [73]. The pervasive presence and influence of recombination and recombination suppression across genomes underscore the fundamental importance of these processes in adaptation and speciation. We hope that the theoretical insights developed here will motivate further empirical and modeling efforts to elucidate how genomic architecture governs the origin and maintenance of species and biological diversity.

## VII. FUTURE RESEARCH

In this study, we have focused on CIs as recombination suppressors to investigate their role in speciation. However, many important aspects of CIs remain poorly understood. Fundamental questions regarding how CIs originate, how they are maintained in stable polymorphisms [30, 33, 74, 75], and how they contribute to local adaptation and interact with other processes in speciation remain unresolved.

A central and enduring question concerns how adaptive alleles come to reside in sufficient physical proximity for a CI to capture them. For instance, in our simulations, the invasion of CIs capturing only adaptive ecological alleles weakens rather than strengthens premating RI and impedes speciation. In contrast, when a CI captures locally adaptive ecological alleles together with locally prevalent mating-bias alleles, it gains invasion fitness and can synergistically enhance both processes, thereby promoting speciation (Fig 34c) [54, 67, 76]. Accordingly, CIs capturing both locally adaptive ecological and mating-bias alleles have higher invasion fitness and are expected to displace comparable CIs lacking the mating-bias allele. However, a key mechanistic puzzle remains: how can a CI assemble multiple classes of adaptive alleles when those alleles initially reside on different chromosomes or are widely separated along the same chromosome?

Addressing this question likely requires consideration of multiple interacting factors, including inversion size, the potential for nested or sequential inversions [5], adaptive competition among alternative CIs, the asexual versus sexual mode of evolution of a CI as determined by its frequency within a polymorphism, pleiotropic regulatory effects, and the modular organization of the genome that may facilitate evolvability, among other factors [33, 76-78]. In a separate review, we explored several of these possibilities and proposed potential mechanisms [79]; however, these hypotheses remain speculative and await formal modeling and empirical validation.

Although the present study models polygenic traits for several speciation processes, many potential interactions were not directly simulated. For example, we have not explicitly simulated the coupling effects of CIs linking polygenic mating-bias barriers with polygenic hybrid-incompatibility barriers. While theoretical reasoning and indirect evidence allow us to generate predictions regarding such interactions, these predictions have not yet been tested through direct simulation.

In sum, our analyses likely illuminate only a small portion of the complex evolutionary dynamics associated with CIs and, more broadly, recombination-suppressing mechanisms [1, 80]. The field remains rich with unanswered questions and offers fertile ground for future theoretical and empirical research aimed at clarifying the origin, maintenance, and evolutionary roles of CIs and other recombination-suppressing mechanisms in adaptation and speciation.

## VIII. ACKNOWLEDGMENTS

We thank Karyn How, Lauren Gieck, and James Lin for reviewing the manuscript and providing valuable comments. ChatGPT was used to assist with copyediting the manuscript.

## IX. APPENDIX

### Two-allele model of hybrid incompatibility

In the two-niche model of a sympatric population shown in Fig 3, we assume two background genotypes, *P* and *Q*, with matings between them producing less viable hybrid offspring due to negative epistasis. Each individual in the population carries either genotype *P* or *Q*. In the absence of premating barriers (i.e., *α* = 1), the probabilities of encounters between individuals are determined by the Encounter Probability Matrix shown in Fig A1. This matrix is identical to the encounter matrix in Fig 5, except that the mating-bias allele *X* is replaced by the incompatibility allele *P*, and allele *Y* is replaced by allele *Q*.

Because *α* = 1, all genotypes match successfully in the first matching round (*n* = 1) and produce offspring. Fig A2 shows the Unit Offspring Matrix of the different matched genotype pairs, analogous to the matrix shown in Fig 6, before the offspring are subject to disruptive ecological selection. In each cell are the ratios of different offspring genotypes produced by the corresponding parental genotype pairs on the vertical and horizontal axes. In the matrix, 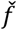 is the offspring return ratio under hybrid incompatibility selection, analogous to the offspring return ratio *f* in ecological selection. The parameter 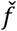 indicates the ratio of offspring with the same *P* or *Q* parental genotype that are regenerated by mating between *P* and *Q* parental genotypes (i.e., nonhybrid offspring). The value of 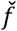 ranges from 0 to 0.5, the same range as *f* in the ecological selection model. The ratios of hybrid genotypes *Ah* and *Bh* are multiplied by *δ* = *c* + (1 − *c*), where *c* is the fraction of hybrid offspring that are viable and 1 − *c* is the fraction that are inviable due to incompatibility. Because *δ* = 1, all ratios in each cell of the matrix sum to one.

If we are only interested in analyzing changes in the population dynamics of the parental genotypes *Ap, Aq, Bp*, and *Bq*, and not what happens in hybrid populations, we can simplify the model by constructing an equivalent one-locus, two-allele model that reproduces the outcome shown in Fig A2, in the same way that we simplify the ecological selection model. In this equivalent model, we assume complete hybrid incompatibility, such that no hybrid offspring are viable (i.e., *c* = 0). Under this assumption, the maximum value that 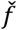 can attain is 0.5 when the genotypes *P* and *Q* are determined by two alleles (*p* and *q*) at a single gene locus.

Within this modeling framework, the baseline value of 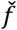 depends on the number of loci underlying the incompatibility. For example, with three loci, 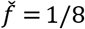, whereas as the number of loci increases toward infinity, 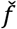 approaches zero. As in our simplified models of ecological and sexual selection, the presence of viable hybrids would increase 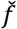 above these baseline values. Nonetheless, the value of 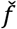 is always constrained to lie between 0 and 0.5. Consequently, in this equivalent one-locus, two-allele model, 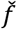 can serve as a measure of the strength of incompatibility selection, analogous to *f* as a measure of disruptive ecological selection and *α* as a measure of sexual selection. Lower values of 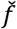 indicate a greater fraction of inviable hybrid offspring due to incompatibility and therefore correspond to stronger incompatibility selection.

If we define a viability ratio 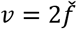, then *v* represents the total fraction of hybrid offspring that regenerate either the *P* or *Q* parental genotypes, while 1 − *v* represents the fraction of hybrid offspring eliminated due to incompatibilities. The value of *v* therefore ranges from 0 to 1 and represents the strength of postzygotic incompatibility selection: smaller values of *v* correspond to stronger postzygotic barriers.

In the equivalent one-locus *P*-*Q* model without viable hybrids, the offspring genotypes that survive incompatibility selection are shown in Fig A3. These surviving offspring are then subjected to disruptive ecological selection, after which the offspring genotype ratios in each cell are renormalized to sum to one, yielding the Unit Offspring Matrix shown in Fig A4.

Notably, the Unit Offspring Matrix in Fig A4 is identical to that obtained from the corresponding mating-bias model shown in Fig 3 when the parameters governing hybrid incompatibility are replaced by those governing mating-bias selection— i.e., by substituting *P* with *X, Q* with *Y*, and *v* with *α*—and the number of matching rounds is restricted to one (*n* = 1). This equivalence demonstrates that the existing GUI applications developed for mating-bias barriers can also be used to simulate the dynamics of postzygotic incompatibility barriers by setting *n* = 1 and relabeling the mating-bias alleles *X* and *Y* as the incompatibility alleles *P* and *Q*. Under this formulation, the fraction of unmatched individuals eliminated after the first matching round in sexual selection is formally equivalent to the fraction of inviable hybrid offspring eliminated due to intrinsic genetic incompatibility.

We acknowledge that our simplified models intentionally ignore the dynamics within viable hybrid populations. We address this limitation by assuming that the presence of such viable hybrid populations— whether arising from ecological divergence, mating bias, or incompatibility—would primarily increase the effective values of *f, α*, and *v* within their respective allowable ranges. As a result, despite these limitations, the simplified framework can nonetheless provide a qualitative yet informative description of the system’s evolutionary dynamics.

Still, the effects of multilocus incompatibility genotypes can be simulated directly using our existing GUI application for multilocus mating-bias genotypes. This is achieved by eliminating premating barriers (i.e., setting *α* = 1) and using *v* to specify the minimum viability of mismatched hybrid genotypes. Under this implementation, offspring with the extreme parental genotypes *P* and *Q* (e.g., *pppp* and *qqqq*) are assumed to be fully viable, whereas the viability of intermediate hybrid offspring depends on the number of mismatched alleles inherited from the parental genotypes. Hybrids carrying the greatest number of mismatched alleles experience the lowest survival probability, reflecting maximal incompatibility as determined by *v*. Survival probability then increases linearly from *v* to 1 as the number of mismatched alleles decreases and the hybrid genotypes approach the parental genotypes.

In the multilocus mating-bias GUI application, this is implemented by setting *α* = 1, thereby removing premating mating-bias barriers, and restricting the number of matching rounds (*n*) to one. The resulting offspring genotype distribution is then modified according to the number of mismatched alleles in each genotype, with genotypes carrying more mismatches experiencing greater elimination due to intrinsic hybrid incompatibility. Offspring genotypes that survive intrinsic incompatibility selection then proceed to extrinsic disruptive ecological selection.

As an illustration, Fig A5 shows the offspring genotype distribution of a four-locus system prior to disruptive ecological selection. A V-shaped function, *f*(*v*), is applied to this distribution to model intrinsic hybrid incompatibility by reducing the frequencies of mismatched hybrid genotypes. The extreme parental genotypes pppp and qqqq remain unaffected, whereas the frequencies of the most mismatched hybrid genotypes at the center of the distribution are multiplied by *v*, the minimum viability ratio representing maximal hybrid incompatibility. Hybrid genotypes between the center and the extremes experience intermediate reductions that scale linearly from *v* to 1, according to their genetic distance from the extreme genotypes.

**Fig A1.**
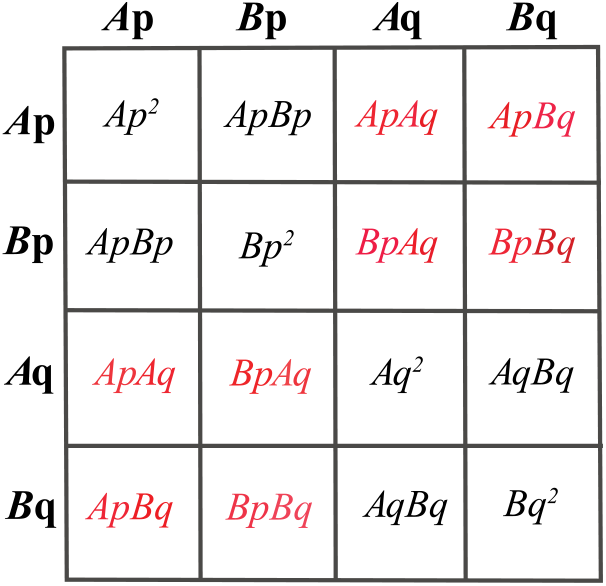
Encounter Probability Matrix for niche-*A* and niche-*B* ecotypes with the *P* and *Q* genotypes. Assuming no premating barriers and random encounters, all matrix elements sum to 1. Encounter probabilities between *P* and *Q* genotypes are highlighted in red.

**Fig A2.**
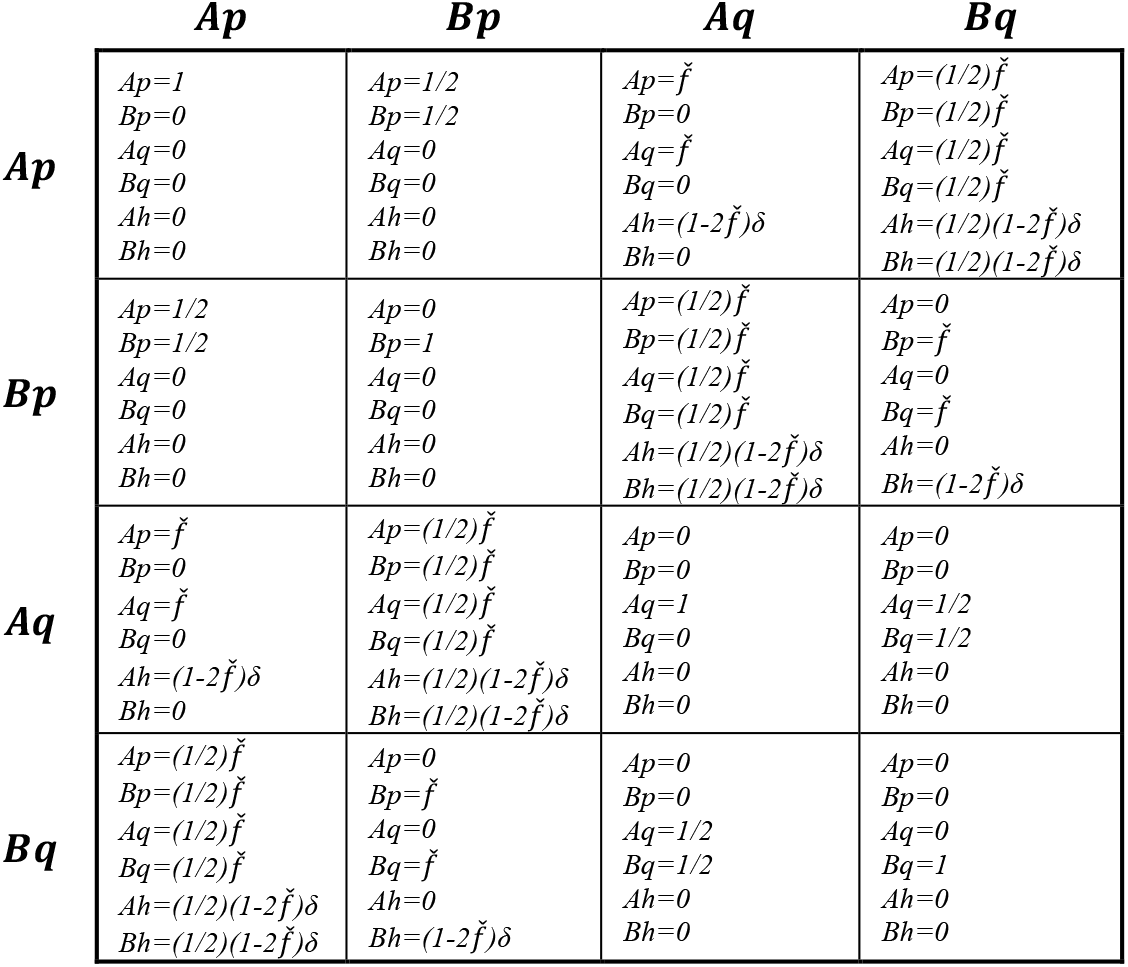
Unit Offspring Matrix for matched *Ap, Aq, Bp*, and *Bq* parental pairs. The matrix shows the offspring genotype ratios produced by each matched parental pair derived from the Encounter Probability Matrix in Fig A1. *Ah* denotes the niche-*A* ecotype with *P*-*Q* hybrid genotypes; similarly, *Bh* denotes the niche-*B* ecotype with *P*-*Q* hybrid genotypes. Hybrid offspring ratios are weighted by *δ* = *c* + (1 − *c*) = 1, where *v* is the fraction of hybrid offspring that are viable and 1 − *c* is the fraction that are inviable due to incompatibility. All genotype ratios within each matrix cell sum to 1.

**Fig A3.**
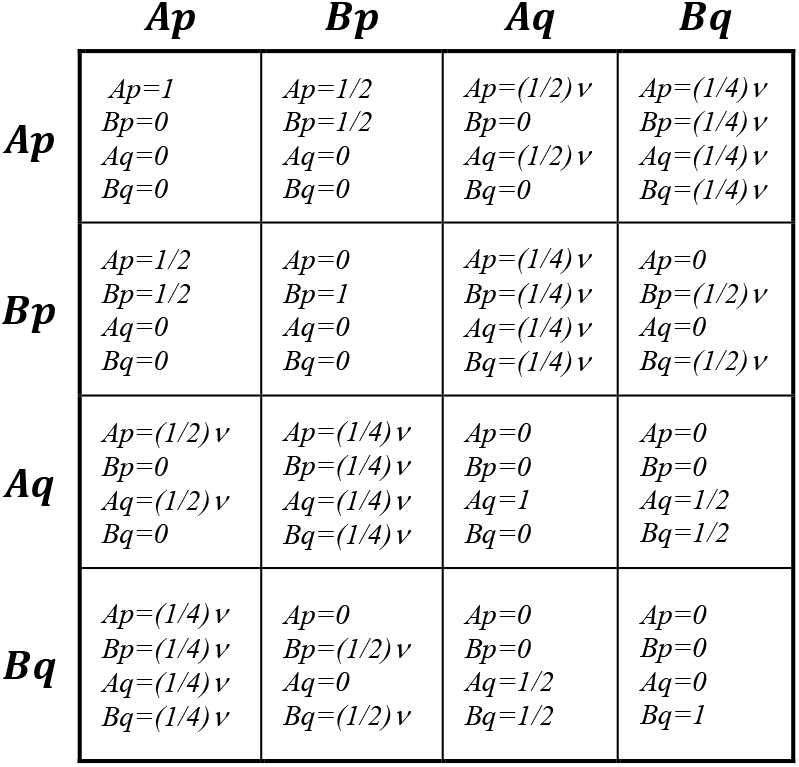
Unit Offspring Matrix for an equivalent one-locus *P*-*Q* model without viable hybrids. The model assumes no viable *P*-*Q* hybrids and uses the parameter *v* to represent the strength of intrinsic hybrid incompatibility selection. The value of *v* denotes the fraction of hybrid offspring that regenerate either the *P* or *Q* parental genotypes, whereas 1 − *v* denotes the fraction eliminated due to incompatibilities. The value of *v* ranges from 0 to 1, and the model implicitly assumes that the effect of having viable *P*-*Q* hybrids is to increase *v* within this allowable range. The matrix therefore shows the offspring genotype ratios that survive intrinsic hybrid incompatibility selection (determined by *v*) before undergoing extrinsic disruptive ecological selection (governed by *f*).

**Fig A4.**
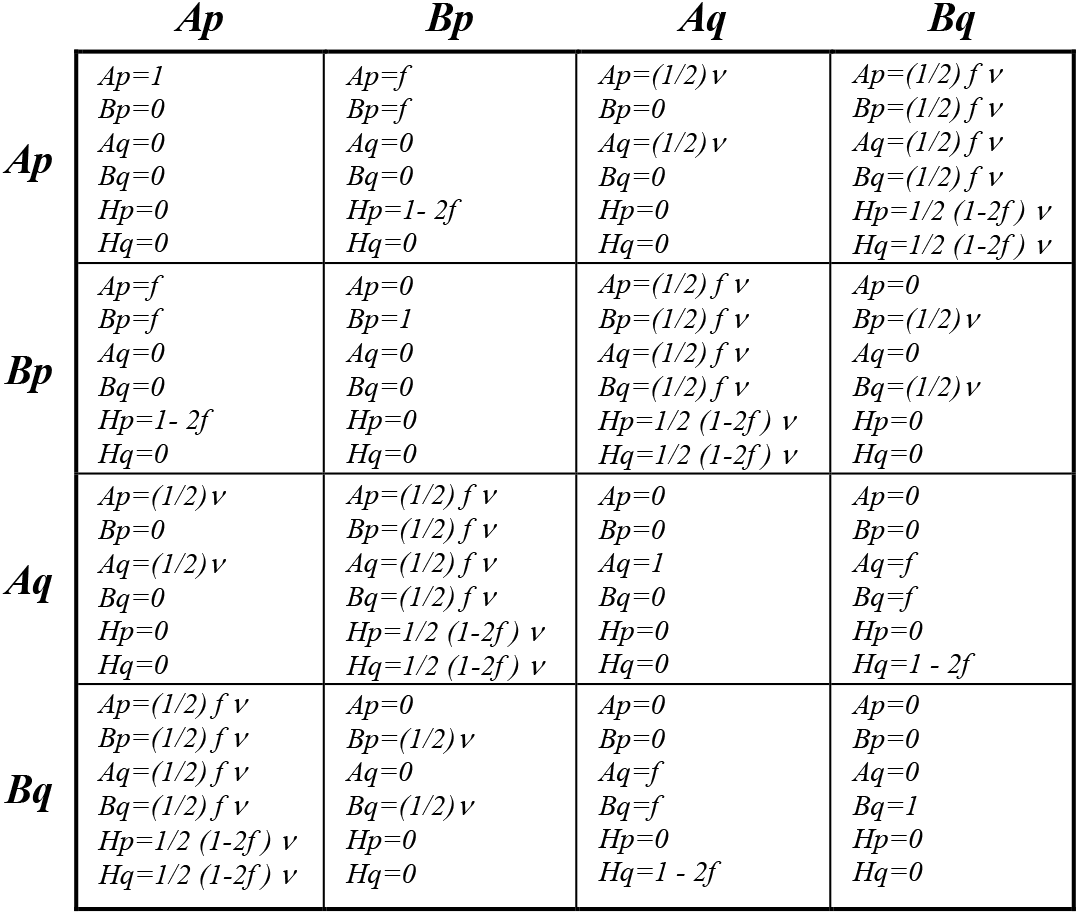
Unit Offspring Matrix for offspring genotypes that survive intrinsic hybrid incompatibility selection and subsequently undergo disruptive ecological selection. The offspring genotypes that survived incompatibility selection in Fig A3 undergo disruptive ecological selection to yield the unit offspring matrix shown. Notably, substituting *v* with *α* yields an identical matrix to that of a single-locus mating-bias model with mating bias *α* and a single mating round (*n* = 1).

**Fig A5.**
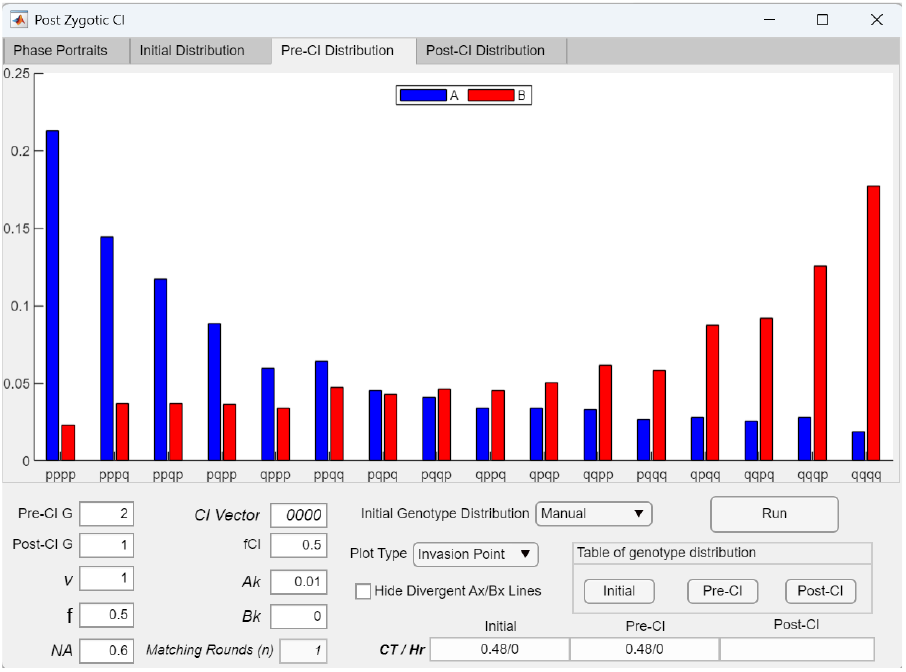
Offspring genotype distribution for a model with four-locus incompatibility genotypes. The bar graph shows a representative offspring genotype distribution for incompatibility genotypes with four gene loci. During intrinsic hybrid incompatibility selection, a V-shaped function, *f*(*v*), is applied to the distribution: genotype frequencies at the center of the distribution are reduced by a factor *v*; the extreme-*P* and extreme-*Q* genotypes remain unchanged; and the frequencies of intermediate hybrid genotypes between the center and the extremes are reduced by factors that scale linearly from *v* to 1.

